# JUMP Cell Painting chemical dataset: Morphological map of 115,796 compounds in human cells

**DOI:** 10.1101/2023.03.23.534023

**Authors:** Srinivas Niranj Chandrasekaran, Srijit Seal, Jeanelle Ackerman, Lakshmi B. Akella, David Alencar Araripe, Eric Alix, D. Michael Ando, John Arevalo, Melissa Bennion, Nicolas Boisseau, Adriana Borowa, Justin D. Boyd, Laurent Brino, Patrick J. Byrne, Hugo Ceulemans, Carolyn Ch’ng, Beth A. Cimini, Djork-Arne Clevert, Nicole Deflaux, John G. Doench, Thierry Dorval, Regis Doyonnas, Vincenza Dragone, Ola Engkvist, Patrick W. Faloon, Johan Fredin Haslum, Briana Fritchman, Florian Fuchs, Sakshi Garg, Tamara J. Gilbert, David Glazer, David Gnutt, Jonathan T. Goldstein, Amy Goodale, Jeremy Grignard, Judith Guenther, Yu Han, Zahra Hanifehlou, Santosh Hariharan, Joshua A. Harrill, Desiree Hernandez, Shane R. Horman, Gisela Hormel, Michael Huntley, Ilknur Icke, Makiyo Iida, Christina B. Jacob, Steffen Jaensch, Isabel S. Jerchel-Furau, Alexandr A. Kalinin, Jawahar Khetan, Maria Kost-Alimova, Tomasz Krawiec, Daniel Kuhn, Charles-Hugues Lardeau, Amanda Lembke, Francis Lin, Kevin D. Little, Gang Liu, Kenneth R. Lofstrom, David J. Logan, Sofia Lotfi, Yi Luo, Franck Madoux, Paula A. Marin Zapata, Brittany A. Marion, Glynn Martin, Nicola J. McCarthy, Lewis Mervin, Lisa Miller, Haseeb Mohamed, Tiziana Monteverde, Elizabeth Mouchet, Alán F. Muñoz, Barbara Nicke, Arnaud Ogier, Anne-Laure Ong, Marc Osterland, Magdalena Otrocka, Constantia Pantelidou, Pieter J. Peeters, James Pilling, Stefan Prechtl, Chen Qian, Krzysztof Rataj, David E. Root, Sylvie K. Sakata, Simon Scrace, Rebecca A. Senft, Runxi Shen, Hajime Shimizu, David Simon, Suganya Sivagurunathan, Peter Sommer, Craig Spruiell, Iffat Sumia, Susanne E. Swalley, Stephanie Tang, Hiroki Terauchi, Amandine Thibaudeau, Amy Unruh, Jelle Van de Waeter, Michiel Van Dyck, Carlo van Staden, Gerard J. P. van Westen, Michał Warchoł, Erin Weisbart, Amélie Weiss, Nicolas Wiest-Daessle, Guy Williams, Shan Yu, Bolek Zapiec, Igor Zlobine, Marek Żyła, Shantanu Singh, Anne E. Carpenter

## Abstract

Image-based profiling decodes complex cellular states for high-impact biological and therapeutic applications, yet a large, public reference dataset of chemical perturbations has been lacking. We present Cell Painting images and 1.53 billion single-cell profiles for 115,796 compounds in human osteosarcoma cells, generated by the Joint Undertaking for Morphological Profiling (JUMP)-Cell Painting Consortium of 18 industry and non-profit partners. Despite technical variability across 12 data-generating laboratories, more than 15% of compounds show statistically significant phenotypic activity, reproducibility drops only 34% versus single-laboratory data, and compounds sharing biological targets often show similar image-based profiles. JUMP establishes a validated public reference resource that brings image-based profiling on par with genomics and transcriptomics. The dataset, shared alongside previously released genetic perturbation data in the Cell Painting Gallery, is a powerful predictive engine for the impact, mechanism, targets and activities of compounds, fundamentally accelerating chemical and cell biology and early-stage drug development.

## Introduction

The image-based microscopy profiling assay, Cell Painting, is useful in many applications in biological research and drug discovery, including disease phenotype identification, hit identification, assay activity prediction, toxicity detection, mechanism of action determination, and functional genomics. Cell Painting multiplexes six inexpensive dyes across eight cellular components, capturing five imaging channels by fluorescence microscopy ^1^. Image analysis software identifies individual cells and extracts a few thousand features from each, producing single-cell profiles. Cell images contain a vast amount of quantifiable information about the status of the cell: for example, whether it is diseased, responding to drug treatment, or impacted by a genetic mutation ^2^. In fact, image profiles yield information density comparable to transcriptomics and proteomics, but at higher throughput and a fraction of the cost ^3–5^.

As with genomics and transcriptomics, a public reference database is critical for interpreting image-based profiles. For a query sample (a disease phenotype, chemical, or genetic perturbation), matching image-based profiles to reference entries yields immediate mechanistic hypotheses through guilt-by-similarity or chemical starting points for drug discovery campaigns. Such queries can discern a target or mechanism of action when compounds are matched with known compounds ^6^. Further, matching compounds to a genetic perturbation’s profile can identify potential regulators of a given gene’s pathway ^7^. Researchers interested in particular compounds can directly discover their phenotypic impact by exploring image features and images. A large structured reference database can also be used to train machine learning models to predict compounds’ assay activity ^8,9^ to identify promising compounds to test physically ^10,11^ or uncover toxicity ^12–14^. Image data can also be used in representation learning, to teach deep learning models biologically useful embeddings ^11,15^.

Large, general-purpose image repositories offer tremendous value to the scientific community (e.g. ^16–19)^ but contain few image sets using a standardized assay like Cell Painting. Before JUMP, the largest public Cell Painting datasets testing chemical perturbations included 30,616 tested compounds from the Broad Institute ^20^ and several datasets of ∼1,500 chemical perturbations each, often in dose-response and/or multiple cell types (https://www.rxrx.ai). These standardized datasets have proven powerful in many applications, including identifying candidate therapeutics and targets. They have demonstrated the value and provided the motivation to create a much larger public dataset, while also raising the question of whether large datasets created at multiple experimental laboratories could be productively combined.

Here, we present the largest freely-available Cell Painting dataset of chemical perturbations, systematically acquired in five replicates across 12 pharmaceutical and non-profit JUMP Consortium partner laboratories, with the support of six technology companies. We characterize data quality, demonstrate major applications of the resource, and resolve technical questions to power future research using this unique data type.

## Results

### Data production and organization

The Joint Undertaking for Morphological Profiling (JUMP)-Cell Painting Consortium comprised 10 pharmaceutical/biotechnology companies (Amgen, AstraZeneca, Bayer, Biogen, Eisai, Johnson & Johnson, Beerse, Belgium, Merck KGaA, Darmstadt, Germany, Pfizer, Servier, Takeda), two non-profit organizations (Broad Institute of MIT and Harvard, Ksilink), and six supporting partners contributing technologies (Ardigen, Google Research, Revvity, Dharmacon, a Revvity, Inc. Company, Nomic, Verily). The consortium aimed to create the largest publicly available Cell Painting dataset, in terms of the number of genetic and chemical perturbations tested, and to determine best practices for running experiments, generating profiles within and across laboratories, and analyzing, aligning, and storing community data.

JUMP produced multiple subsets of data (Figure 1A), including the already-published JUMP genetic perturbation dataset, including over 15,000 genes overexpressed using Open Reading Frames (ORFs) or knocked out by Clustered Regularly Interspaced Short Palindromic Repeats (CRISPR) guides (a subset of cpg0016-jump) ^21^. Three additional pilot datasets include cells perturbed under various experimental and imaging conditions: testing alternative cell lines, genetic perturbation types, time points, and imaging conditions (cpg0000-jump-pilot) ^22^, optimizing staining concentrations and conditions (cpg0001-cellpainting-protocol) ^1^ and comparing multiple microscopes and their settings (cpg0002-jump-scope) ^23^.

**Figure 1:**
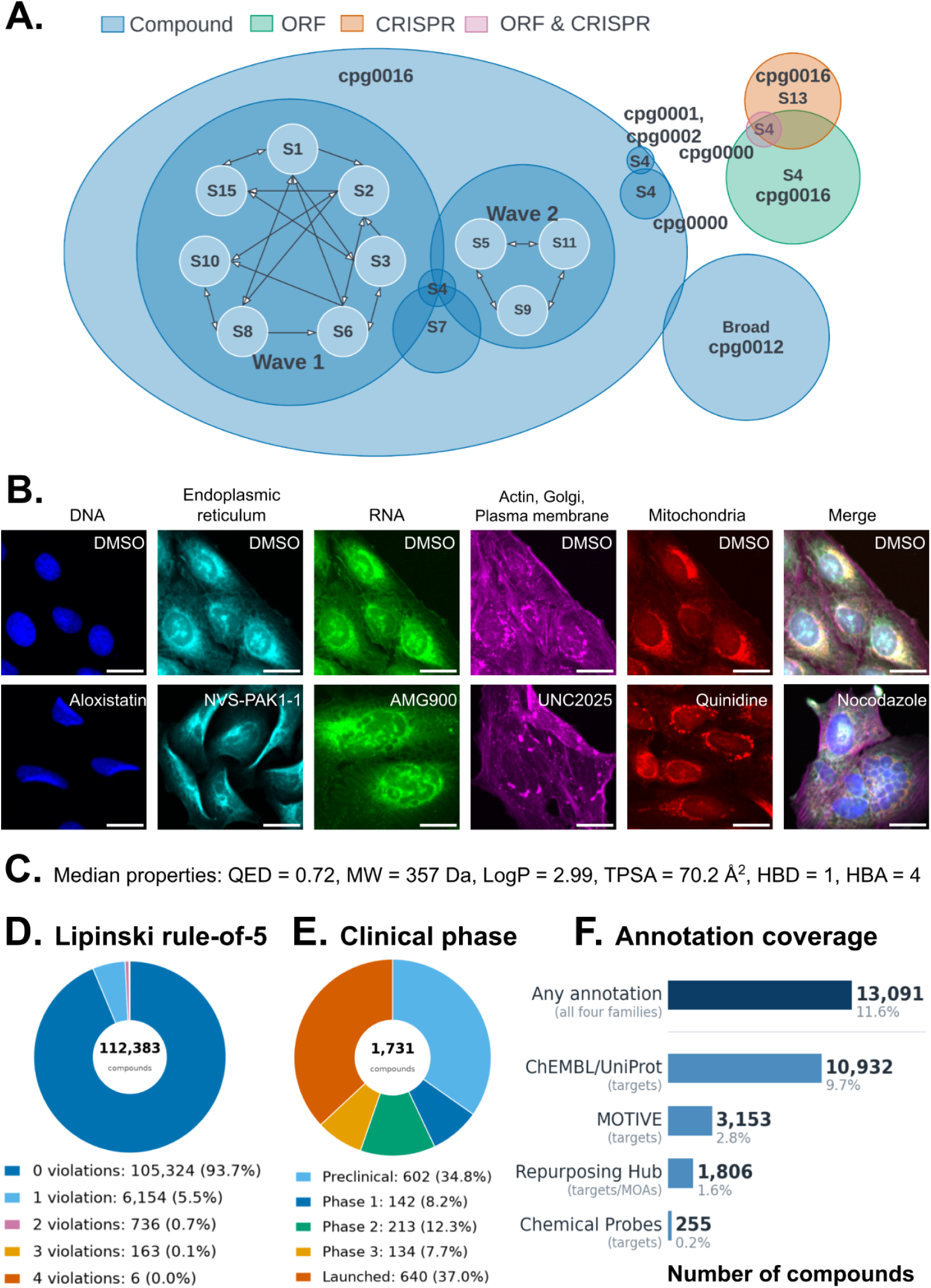
The JUMP consortium generated Cell Painting data for 115,796 compounds in cpg0016-jump. (A) The JUMP Cell Painting dataset includes cpg0000, cpg0001, cpg0002, and cpg0016 (Methods). This study presents the 115,796 chemical compounds of cpg0016, generated across the 12 partner sources/laboratories (note: Source 7 and 13 correspond to the same laboratory). Typically, five replicates of each compound were generated at two to four laboratories besides the one that nominated the compound. The arrows denote the compound exchange logistics (Supplementary Table 1) and apply only to the ten wave 1/wave 2 nominated-compound sources; S4, S7, and S13 did not participate in that exchange. cpg0016 genetic perturbations were previously published ^21^: CRISPR data generated entirely at S13 and ORFs at S4, the Broad. The size and overlaps in the schematic are not precise to scale. Overlap is also shown for cpg0012 ^20^, Broad’s existing public dataset of 30,616 bioactive compounds, reprocessed using the JUMP processing pipeline (Methods). (B) Example Cell Painting images. Top: One negative control field of view (DMSO, dimethyl sulfoxide) showing the five imaged channels, pseudocolored, plus their merge. Bottom: visually strong phenotypes in each channel for various compounds. Single-channel and merged images were individually scaled to the 99th intensity percentiles and full intensity range of the image for visibility, respectively. All images are from a single source (S5) for consistency. Scale bars = 25 μm. (C-F) Chemical characterization of the 112,462 analyzed JUMP compounds: (C) Median physicochemical properties: QED (Quantitative Estimate of Drug-likeness ^24^); MW (molecular weight); LogP (calculated octanol-water partition coefficient); TPSA (topological polar surface area); HBD (hydrogen-bond donors); HBA (hydrogen-bond acceptors). (D) Lipinski rule-of-five (Ro5) compliance. (E) Clinical development stage for the 1,731 compounds with such annotation. (F) Number and percentage of compounds with a target or mechanism of action (MOA) annotation from the Drug Repurposing Hub, ChEMBL, the Chemical Probes Portal, and MOTIVE.

Here, we present the chemical perturbations in cpg0016-jump and explore the dataset’s value as a reference. The full dataset comprises 1,151,808 sample wells of chemical and genetic perturbations, yielding 43,072,410 field-of-view single-channel images (127 TB). It includes 115,796 chemical perturbations and several versions of their extracted image profiles; this paper uses these compounds’ 1,533,879,336 CellProfiler single-cell profiles (56 TB) and well-level profiles (103 GB, for the raw aggregated profiles alone). Images were generated across 12 data-producing laboratories (also called “sites” or “sources”). Each source exchanged its nominated compounds with either two or four other sources, where they were assayed and imaged in human osteosarcoma U2OS cells (Figure 1B), using different instruments and microscopes (Methods - Experimental Design, and Supplementary Table 1). This study analyzed the 112,462 compounds that were tested at 10 μM plus positive control and Target-2 compound plates at 5 μM; Source 7’s bioactive compounds were tested at 0.625 μM and excluded (discussed later). JUMP’s design captures real-world technical variability, allowing studies of data alignment and batch effect correction and improving the chances of external laboratories aligning their data successfully. The consortium established standard segmentation and feature extraction procedures and pipelines (Methods) ^1^. We followed consistent platemaps (Supplementary Figures 1, 2) and included JUMP-Target-2-Compound (Supplementary Figure 3), a positive control plate of 301 diverse compounds annotated as targeting 160 proteins ^22^ in most experimental batches (except Source 1 and 9, see Methods), to enable alignment of data within the JUMP dataset and with future datasets generated outside the consortium. The structure of the experiment, its sources, batches, microscopes, plate format (mostly 384-well but some 1536-well), and control plates are mapped in Supplementary Figure 4. JUMP data are shared in the Cell Painting Gallery ^18^, along with dozens of other Cell Painting datasets, including Broad’s existing public dataset of 30,616 bioactive compounds ^20^, reprocessed using the JUMP processing pipeline.

### The JUMP compound collection spans broad drug-like chemical space

Each partner nominated and provided compound samples and structures based on their own, undisclosed criteria (Supplementary Figure 5). The 112,462 analyzed compounds are typically drug-like (Figure 1C, D), reflecting their origin from pharmaceutical screening libraries (105,324 Lipinski rule-of-5 compliant; 93.7%), and structurally diverse (73,208 unique Bemis-Murcko scaffolds). They contain 640 annotated as launched drugs, 602 preclinical, 142 Phase 1, 213 Phase 2, 134 Phase 3 (Figure 1E) and 5,021 pan-assay interference (PAINS)-flagged compounds ^25^. Across the full 115,796-compound collection, 105,789 (91.4%) compounds can be mapped against 39 external databases via UniChem and MyChem; 59,904 (53.3%) are commercially available (per MolPort), 13,091 (11.6%) have target or mechanism of action (MOA) annotations (Figure 1F), and 8,852 (7.9%) had two separate samples of the same compound provided by different partners. Functional use categorization using the U.S. Environmental Protection Agency (EPA) CompTox pipeline ^26^ identified 427 compounds with a curated Functional Use Categorization and 7,651 with predicted Functional Use Probabilities, with the most common classifications being antimicrobials, skin conditioners, colorants, and antioxidants (Supplementary Figure 5F, G). This diversity across drug-like chemical space makes JUMP suitable as a broadly useful reference database.

### Phenotypic signal is detectable and reproducible across laboratories

While a public dataset can be used for many purposes, important applications are unlocked if it can be queried with future data produced by external laboratories, despite technical variation. We therefore explored data quality in this multi-laboratory dataset, asking whether biological signals survive the technical variability introduced by different laboratories, instruments, microscopes, filter sets, and staff across the Consortium (Supplementary Tables 2-5 and Supplementary File PROTOCOL).

The hit rate was, as expected for a profiling assay, much higher than typical single-readout assays: 15.2% of tested compounds (17,069 of 112,462) show statistically significant phenotypic activity, distinguishable from negative controls (false discovery rate, FDR < 0.10) (Figure 2A). The true active fraction is certainly even higher: phenotypic activity requires replicate consistency ^27^, which is eroded by batch effects. For this study, we used profiles of engineered features extracted by CellProfiler software ^28^ which are more readily interpretable and which performed comparably to Cell Painting convolutional neural network (CP-CNN) deep learning features ^11^ for both phenotypic activity and phenotypic consistency (matching compounds sharing an annotated protein target) (Supplementary Figure 6). CellProfiler features also showed 3.5-fold less batch effect after correction (using Harmony ^29^; Supplementary Figure 7A, Supplementary Figure 8A). Source 7 was excluded because its compounds were screened at a lower concentration than the rest of the dataset (Methods).

**Figure 2:**
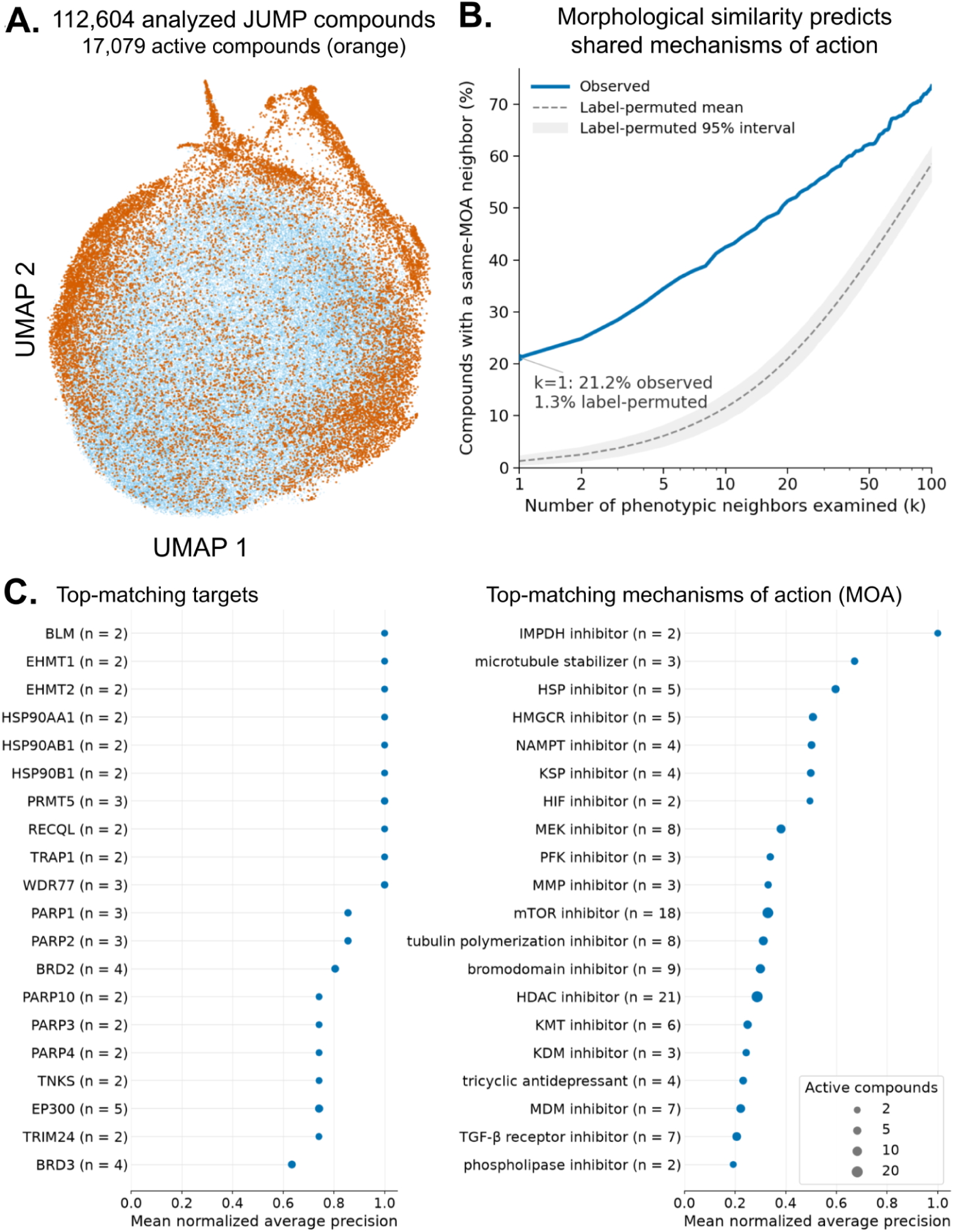
Cell Painting profiles of compounds sharing the same target or mechanism of action tend to cluster. (A) Cosine Uniform Manifold Approximation and Projection (UMAP) gives a global view of phenotypically active JUMP compounds (orange) relative to those lacking Cell Painting activity (light blue, Benjamini–Hochberg-corrected p-value above 0.10). UMAP is for display only; phenotypic activity was calculated from high-dimensional profiles. Gray denotes 142 profiles (0.13%) without an activity result: 8 positive controls, 1 negative control, 124 Target-2-plate-only compounds excluded from this analysis, and 9 compounds with only 1 replicate. Alternate views (sources, cell count, MOA classes) are shown in Supplementary Figure 8. (B) Similarity of Cell Painting profiles among compounds annotated with the same MOA was evaluated by computing the cumulative percentage of eligible compounds with at least one same-MOA compound among their k nearest eligible phenotypic neighbors. Here, similar scaffolds were excluded but similar results were found when included (see Methods). (C) Top target (Chemical Probes) and MOA (Drug Repurposing Hub) classes in terms of phenotypic consistency, with the number of compounds in that class shown as n. Seven other annotations’ top classes are provided in Supplementary File PHENOTYPIC_CONSISTENCY.

No correction for plate layout effect was needed (Supplementary Figure 7B), nor did regressing cell density out of the morphological features improve performance (an ordinary-least-squares regression of each feature on the per-well cell density; data not shown). Cell density (cells per mm^2^) negatively correlates with but does not fully explain phenotypic activity (Spearman rho = −0.33; Supplementary Figure 7C, Supplementary Figure 8B). Still, phenotypically active compounds tend to have ∼10% fewer cells (median 411 cells/mm^2^ versus 457).

We next explored variability within and across sources directly, using the Target-2 plate run repeatedly at most laboratories. The 301 unique compounds on this plate are derived from the Drug Repurposing Hub collection and thus known to impact biological systems generally, but were not known to be phenotypically active by Cell Painting. Thus, the true upper bound is unknown but certainly below 100%. Still, 77% of Target-2 compounds were phenotypically active using replicates within-laboratory (comparable to the 68% of known bioactive molecules in our prior study ^30^, albeit with quite different compound selection, analysis methodologies, and plate-layout confounding). The signal dropped to 51% when using replicates across laboratories. This 34% reduction in signal represents the impact of multi-laboratory data generation, even after batch correction (Supplementary Figure 9). We conclude that meaningful biological signals persist despite multi-laboratory technical variability.

### Image-based profiles can predict the targets and mechanisms of chemical compounds

One of the most compelling and well-proven applications of profiling, particularly Cell Painting, is generating hypotheses about chemicals’ mechanisms of action (MOA) based on the principle that compounds with similar mechanisms or targets should yield similar cell phenotypes ^2^. Consistent with prior work and smaller datasets, we observed and quantified this for MOA-annotated compounds; nearest-neighbor compounds had the same MOA at a 16-fold higher rate than expected by chance (Figure 2B, 21.2% observed versus 1.3% under label permutation). Quantifying phenotypic consistency ^27^ across nine annotation sources (Figure 1F; Methods), many targets and mechanisms do indeed cluster, as do compounds associated with the treatment of particular disease classes such as neurology/psychiatry and allergy (Figure 2C, Supplementary Figure 8C, Supplementary File PHENOTYPIC_CONSISTENCY). We confirmed this isn’t an artifact of similarly annotated compounds being co-located (Supplementary Figure 8A); in fact, some member compounds falling below our activity thresholds nevertheless cluster near their annotated group members (Supplementary File PHENOTYPIC_CONSISTENCY). The JUMP dataset is therefore a rich resource for capturing compound targets and mechanisms using guilt-by-association analyses for novel compounds. This also implies JUMP data can power lead-hopping, identifying novel chemical structures producing a known compound’s desired phenotypic effect but lacking undesirable structural or *in vivo* properties.

Evaluating more compounds by loosening the phenotypic activity threshold did not improve phenotypic consistency overall (Supplementary Figure 10A), nor did alternative strategies such as calculating phenotypic consistency using medians instead of means, nor absolute value of cosine similarity (to allow annotation groups like target and disease area to have compounds with opposite phenotypes) (Supplementary Figure 10B,C). To help design future datasets, we found that 2-3 replicates (when matching within a given laboratory’s data) and 4-5 replicates (when including other laboratories) provide 90% of the maximal MOA phenotypic consistency (Supplementary Figure 11).

### What kinds of compounds are phenotypically active and consistent

We wondered what properties make compounds more likely to be phenotypically active and consistent in the Cell Painting assay. Generally, phenotypic activity and consistency are correlated, at the individual compound level and especially when aggregated at the shared-target-group level (Figure 3A), though this correlation can only be calculated for target-annotated compounds, which tend to be optimized and thus are not representative.

**Figure 3:**
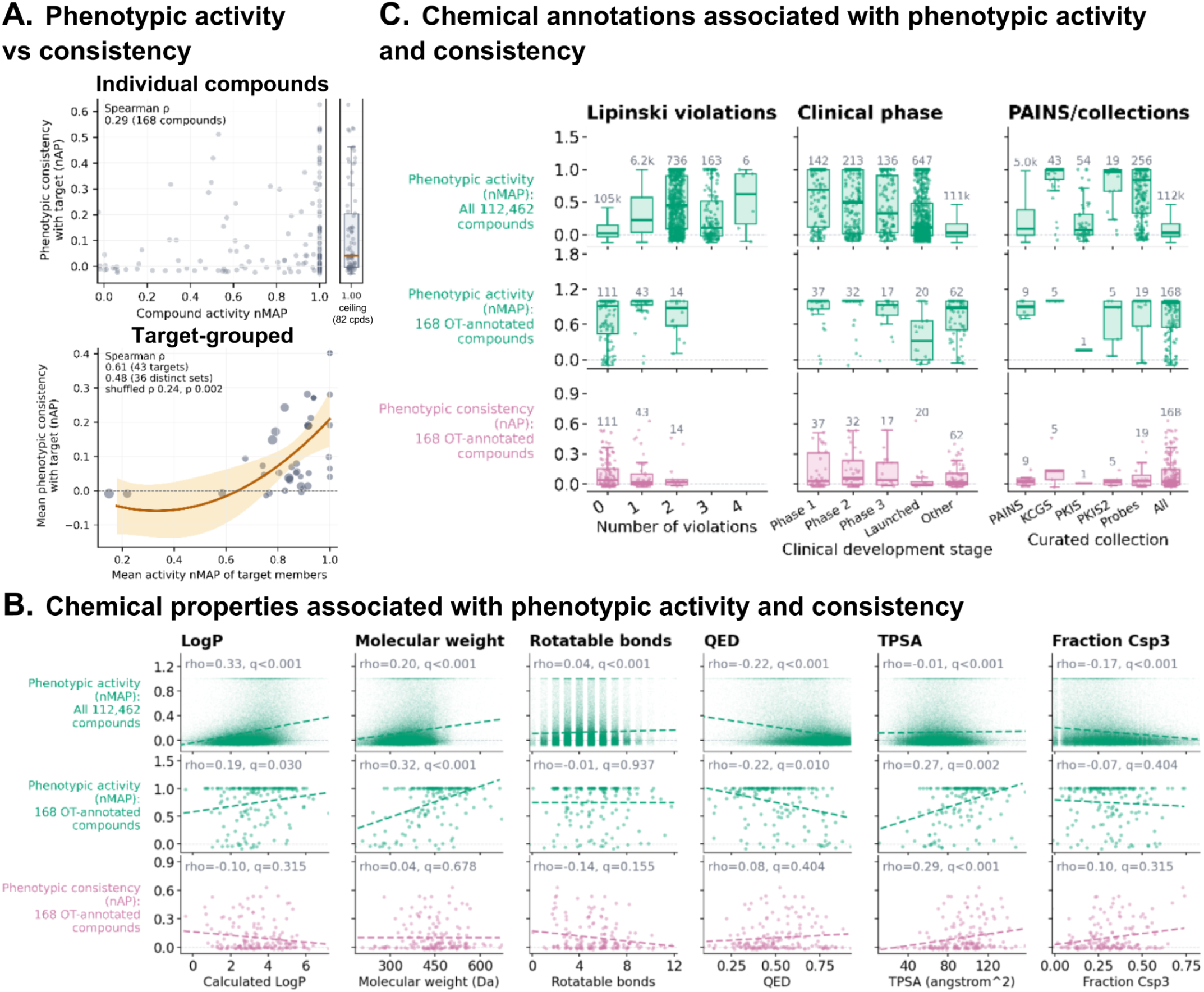
Properties of compounds associated with phenotypic activity and phenotypic consistency in the Cell Painting assay. (A, top) Phenotypic activity (normalized mean average precision, nMAP) measures a compound’s replicate retrieval and is plotted here against phenotypic consistency for a compound (normalized average precision, nAP), which measures a compound’s retrieval of other compounds annotated to the same target (versus active compounds annotated to unrelated targets), using its best-matching target. The plot shows the 168 JUMP compounds with OpenTargets annotations for targets with five or more analyzed compounds; an nAP of zero is the random expectation. Transparency reveals density. The vertical accumulation (82 of 168 compounds) at activity nMAP = 1.0 is the ceiling; adjacent, the same points are jittered horizontally over a box showing the median and interquartile range and whiskers extending to 1.5 times the interquartile range. (A, bottom) Similar analysis, but each data point is a target; data point area represents the number of analyzed compounds annotated to that target (range: 5-21). The orange curve is an across-target quadratic model adjusted for target-member annotation degree and active-reference count; the shaded band is its 95% cluster-robust interval by exact membership/reference set. (B) Six properties of chemical structures are analyzed for association with phenotypic activity across all 112,462 analyzed compounds (top), 168 target-annotated compounds as in (A) (middle), and with phenotypic consistency (bottom), for each compound’s best-matching annotated target. Dashed unweighted least-squares lines are drawn. Integer properties are jittered and rotatable bonds excludes one 18-bond compound, being clipped at 12. Labels report Spearman rho with a Benjamini-Hochberg q. (C) Chemical annotations are analyzed for association with phenotypic activity and consistency as in (B). Boxes plot quartiles, with whiskers reaching 1.5 interquartile ranges, outliers are not drawn, and the count above each box is the number of compounds (individual data points not shown for categories with more than 1,000). Collections are defined in Methods.

Among physicochemical properties (Figure 3B, top and middle rows), LogP is the primary predictor of phenotypic activity (ρ = 0.33); lipophilicity may drive phenotypic response through increased intracellular bioavailability, impacts on organelle membranes, and non-specific target promiscuity. Compounds whose structure violates the Lipinski rule of five ^31^ were also more likely to be phenotypically active (Figure 3C, left column). These trends did not typically hold for phenotypic consistency (Figure 3B,C, bottom row), although results varied depending on which compound annotations are used (Supplementary Figure 12A). This implies that although certain physicochemical properties increase the likelihood of a Cell Painting phenotype, those phenotypes are not distinctive enough to yield strong phenotypic consistency within target classes.

Next, we examined various annotations for the compounds. Clinically tested compounds were more likely to be phenotypically active (Figure 3C, middle column). Even so, compounds further along in testing, particularly those launched, were *less* likely to be phenotypically active than earlier phases, perhaps indicating successful compounds induce fewer side effects. Pan-assay interference (PAINS) nuisance compounds were more likely to be phenotypically active than those not annotated as such (Figure 3C, right column). Compounds annotated as selective kinase inhibitors and curated Structural Genomics Consortium (SGC) chemical probes are even more likely (Figure 3C, right column). Again, trends seen for phenotypic activity do not hold for phenotypic consistency.

We also examined structural elements: compounds with certain functional groups, BRICS (Breaking Retrosynthetically Identified Chemical Substructures) fragments, and Bemis-Murcko scaffolds were more (or less) likely to be phenotypically active (Supplementary Figure 12B-D; 58 of 74 functional groups, 293 of 868 BRICS groups, and 34 of 93 Bemis-Murcko classes, Supplementary File STRUCTURAL_CLASSES). Some of these results mirror those seen for intrinsic chemical properties: for example, particular substructures reflect lipophilicity. Other results are also expected; among BRICS, tight kinase-hinge-like motifs are more phenotypically active than average whereas polar heterocycles are less.

The JUMP dataset is also relevant to studying chemicals beyond drug discovery. Among the six EPA-curated functional-use categories (FUCs) ^26^ that met the testing criteria, none were statistically more or less likely to be phenotypically active than the others (Supplementary Figure 12E). Structure-based predicted functional use probabilities (FUPs) can be calculated for many more compounds; of 24 testable FUP classes, six were more likely to be phenotypically active (FDR < 0.05; Supplementary Figure 12F), including some whose inherent structure or intended function relies on biological reactivity, such as heat stabilizers (36.1% phenotypically active), antimicrobials, antioxidants, and fragrances. The four classes less likely to be phenotypically active included predicted buffers (0.4% active), skin- and hair-conditioners, and preservatives. Thus, phenotypic activity associates with a chemical’s intended impact on biological reactivity.

Finally, we explored polypharmacology. Although the number of appropriately-annotated compounds was small, when compounds annotated as hitting two different targets yielded a *similar* profile, profiles of a compound annotated as hitting both targets were usually also similar (Supplementary Figure 13). When compounds annotated as hitting two different targets yielded *distinct* profiles, profiles of a compound annotated as hitting both targets most often strongly matched one of their annotated targets rather than a blend of the two, with a few exceptions such as compounds targeting EGFR and ERBB2 (Supplementary Figure 13); advanced methods and improved annotations might improve the detection of polypharmacology ^32,33^.

### Image-based profiles reflect structure-activity relationships and reveal activity cliffs

A chemical’s structure determines which cell components it may bind and impact, which may in turn impact a cell’s morphology, as quantified by the Cell Painting assay. 60.3% (158/262) of Bemis-Murcko scaffolds (with ≥5 active compounds) were more morphologically self-similar than size-matched random groups (Supplementary Figure 14A), and, conversely, 87.5% (70/80) of morphology-based clusters contained at least one significantly overrepresented scaffold. At the level of individual phenotypically active compounds, however, structural and morphological neighborhoods overlapped only modestly. Their mean Jaccard overlap was 0.010, compared with 0.0015 expected by chance, and only 43.8% of compounds shared any neighbor between the two spaces (Supplementary Figure 14B). Compounds in certain mechanism-of-action classes were somewhat more likely to overlap, including p38 MAPK, HDAC, PI3K, EGFR, and mTOR inhibitors (Supplementary Figure 14C). Morphological similarity increases with structural similarity across the 145,530,330 pairs among 17,061 phenotypically active compounds with a nonzero Morgan ECFP4 fingerprint (Supplementary Figure 14D).

We next identified activity cliffs: compounds that share a core chemical structure but with one yielding a Cell Painting phenotype and the other with no activity, potentially indicating key structural elements that impact properties like protein binding and membrane permeability. 19% of structurally similar pairs were cliffs (Supplementary Figure 14E, Supplementary File ACTIVITY_CLIFF_PAIRS). Others have analyzed cliffs based on scaffolds ^34^.

We also identified *phenotypic cliff* pairs where both compounds are phenotypically active but produce *different* Cell Painting profiles. This type of cliff is only feasible for assays with multiple readouts. 4,089 pairs (32% of the 12,920 structurally similar phenotypically active pairs) show divergent Cell Painting phenotypes (cosine similarity < 0.3) (Supplementary Figure 14F, Supplementary File PHENOTYPIC_CLIFFS). Phenotypic cliffs can arise from engagement with multiple targets, among other mechanisms. 159 pairs (out of 4,089) had target annotations for both compounds from the same source; of these, 59.7% had no annotated protein target in common, as compared to 29.5% of structurally similar pairs with concordant morphology (odds ratio 3.5, p = 1.5×10⁻⁸), and this enrichment held using ChEMBL targets alone. The lists we provide are valuable resources for medicinal chemists studying cliffs.

### Image-based profiles predict only certain chemical activities

We next explored whether JUMP Cell Painting data could be used to train predictors for various chemical activities. Predicting PAINS compounds is useful because they appear frequently as false positives in other assays. Using 5,032 PAINS-flagged compounds ^25^ in JUMP, we found that Cell Painting profiles indeed predict PAINS status (scaffold-split ROC-AUC 0.634 ± 0.014) better than chance (ROC-AUC 0.500 ± 0.007) and better than simple cell count alone (ROC-AUC 0.547 ± 0.014) (Supplementary Figure 15A, B), consistent with our prior work ^35^. However, there is no improvement over a predictor built on chemical structures’ molecular properties alone (MW/LogP/TPSA; ROC-AUC 0.634 ± 0.007), so there is no practical utility except, speculatively, for chemicals or biologics in regions of chemical space less well captured by these properties. We do note that combining molecular properties and morphology seems to provide a boost (ROC-AUC 0.660 ± 0.018).

Cell Painting profiles showed minimal predictive signal for chemical activity at the organism level, including drug-induced human cardiotoxicity (DICT, consistent with our prior finding of AUC 0.52 ^36^), human drug-induced liver injury (DILI), and continuous pharmacokinetic (PK) properties (Supplementary Figure 15C). Chemical structure fingerprints (Extended-Connectivity Fingerprint, diameter 4, ECFP4) outperformed Cell Painting profiles for all of these endpoints and combining the two data sources did not improve predictions (Supplementary Figure 15D). These results likely reflect the U2OS cell line’s limited relevance to organ-specific metabolism and toxicity; our ‘Omics for Assessing Signatures for Integrated Safety (OASIS) Consortium aims to improve hepatotoxicity predictions using more compounds and specialized cell models ^37^.

Cell Painting profiles had only slightly better predictive power for cell-based and biochemical assays, such as mitotoxicity (consistent with our prior finding ^38^), where it beats structure-based chemical fingerprints (Supplementary Figure 15C). Across a broader external activity-prediction benchmark, ToxCast (on 95 assays with at least 100 compounds, including at least 20 active and 20 inactive compounds), Cell Painting profiles achieved a median AUC of 0.611, compared with 0.674 for chemical fingerprints and 0.538 for cell count (Supplementary Figure 15E). On 51 ChEMBL single-protein targets, morphology achieved a median AUC of 0.563 versus 0.584 for fingerprints (Supplementary Figure 15E). A few particular assays were better predicted by morphology than by chemical structure alone, including some cell-based nuclear-receptor and stress-response assays such as CAR (Constitutive Androstane Receptor; morphology AUC 0.854 vs 0.607 for fingerprints), FXR (Farnesoid X Receptor; 0.807 vs 0.687), and p53 (encoded by TP53, 0.821 vs 0.764). ToxCast assays for which hits yielded phenotypically consistent Cell Painting profiles are listed in Supplementary File PHENOTYPIC_CONSISTENCY.

### Image profile matching identifies PPARG inverse agonist mechanisms

To evaluate whether JUMP is an effective resource for external experiments, we tested whether profiling a small compound series in a distinct cell type could predict mechanism of action purely by querying the pre-existing reference database, without needing to concurrently screen a full reference library. We profiled two recently discovered novel chemical series of PPARG inverse agonists that directly bind and modulate PPARG ^39,40^ alongside the JUMP-Target-2-compound reference plate of annotated compounds, in duplicate for each compound-concentration. One pair of plates used U2OS cells for consistency with the JUMP dataset and the other used UM-UC-9 cells, which have high PPARG expression and a strong phenotypic and antiproliferative response to PPARG perturbation ^41^. The novel compounds yielded a dose-dependent, consistent Cell Painting phenotype (Supplementary Figure 16).

We first sought phenotypic matches among compounds within this stand-alone experiment, including the well-annotated compounds of the Target-2 plate processed concurrently to the PPARG inverse agonist plate, which includes selected compounds with various cell proliferation-related mechanisms. In U2OS and UM-UC-9 cells, apart from known PPARG inverse agonists, we saw other strong matches for the PPARG inverse agonists, such as GSK2110183 (an AKT inhibitor inducing G1 arrest) and several CDK4/6 inhibitors including palbociclib (Figure 4A and Supplementary Figure 17). Visual inspection of the images revealed similar morphology between cells treated with PPARG inverse agonists and palbociclib, with cells adopting an enlarged, fried egg-like shape (Figure 4B and Supplementary Figure 18).

**Figure 4:**
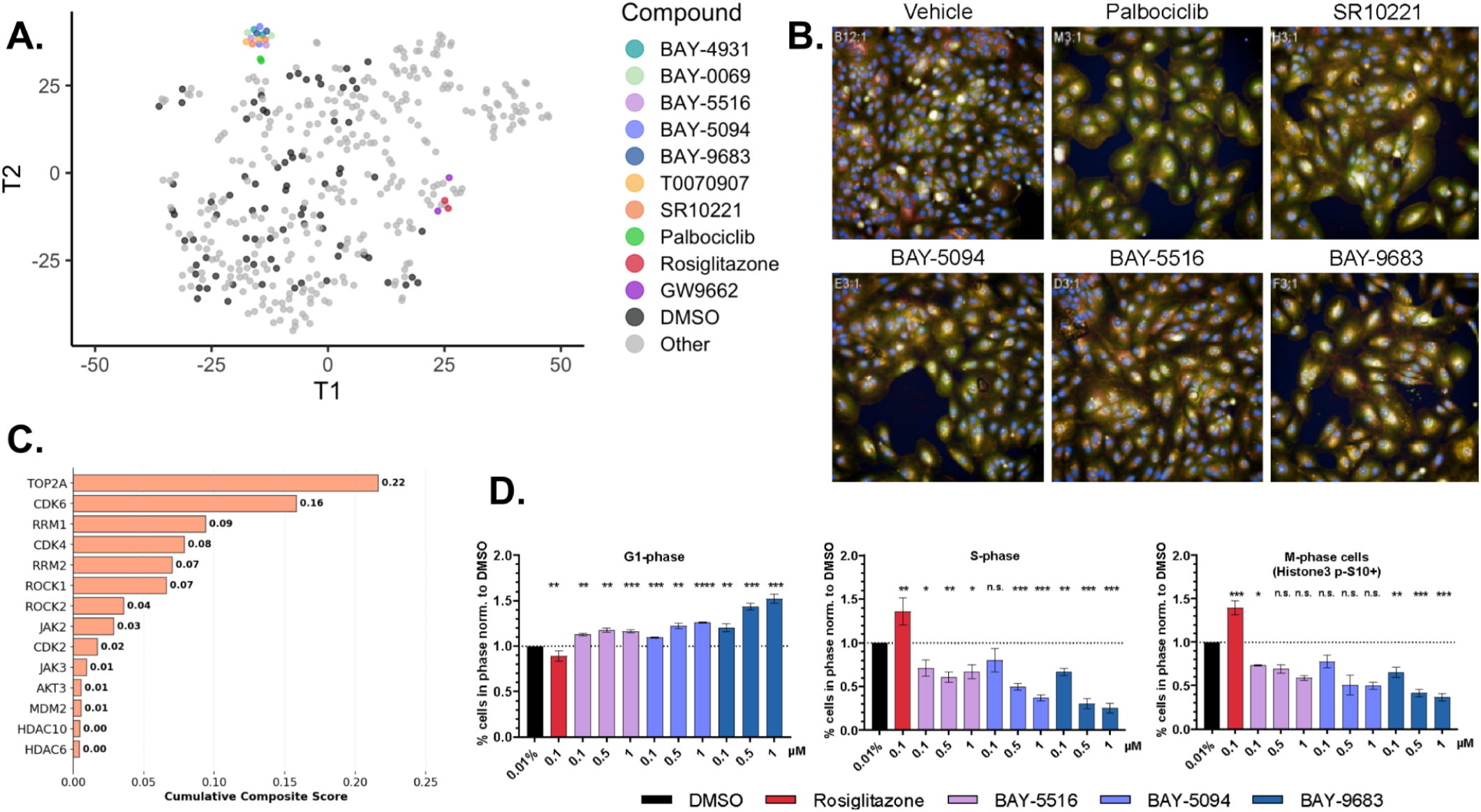
PPARG inverse agonists induce G1 accumulation and senescence in UM-UC-9 cells. (A) t-Distributed Stochastic Neighbor Embedding (t-SNE) visualization of Cell Painting morphological profiles of UM-UC-9 cells, showing only the highest concentration for simplicity. The plot is colored by mechanism of action, with the BAY-compounds, T0070907, and SR10221 being PPARG inverse agonists or antagonists. (B) Merged fluorescence images for five Cell Painting channels of UM-UC-9 cells treated with indicated compounds at 900nM for 72 hours. Representative image shown (see Supplementary Figure 18 for full images with scale bars). (C) Annotated targets for the compounds from the JUMP dataset that best match to the set of five query BAY-compounds in UM-UC-9 cells. See Methods for details. (D) Cell cycle analysis of PPARG-amplified UM-UC-9 bladder cancer cells treated with indicated compounds for 24 hours and measuring the proportion of cells in G1- or S-phase (n ≥ 4) as well as M-phase (n ≥ 2); BAY-compounds generally increase cells in G1 and decrease cells in S and M while for comparison, Rosiglitazone shows the opposite pattern.

We then aligned these plates with the JUMP dataset (see Methods) and examined the annotated targets of compounds with Cell Painting profile similarity to the PPARG inverse agonists. PPARG itself was not represented among the top targets; most PPARG-targeting compounds were filtered out of the analysis due to low phenotypic activity, likely because U2OS cells are less responsive to PPARG modulation as compared to UM-UC-9 cells. Instead, the top three target families that phenotypically match the query profile of the novel PPARG inverse agonists in UM-UC-9 cells were CDK, RRM, and TOP2A (Figure 4C and Supplementary Figure 19 and Supplementary Figure 20A). All are involved in DNA replication during S phase and would therefore cause G1 arrest; they have known interactions and relationships to cancer ^42–44:^ most CDKs (Cyclin-Dependent Kinases) regulate the cell cycle at S-phase initiation, ribonucleotide reductase subunits (RRMs) provide the building blocks for DNA synthesis, and TOP2A (DNA topoisomerase II alpha) resolves topological constraints. We note that matching worked better *across* cell types; having a stronger phenotypic response in a different cell type (UM-UC-9) provided better matching than weaker phenotypes in the same cell type (U2OS).

Given the phenotypic match between the BAY PPARG inverse agonists and CDK4/6 inhibitors, we explored JUMP genetic perturbation datasets ^21^. Analysis revealed matching phenotypes among various CDK4/6 and PPARG gene family members (by CRISPR-Cas9 knockout and overexpression), confirming their similar mechanisms (Supplementary Figure 21); matching compounds to genes directly awaits better data alignment methods ^45^.

The CDK inhibitor palbociclib is known to induce G1 arrest through blocking the phosphorylation of retinoblastoma protein ^46^ and is also known to induce cellular senescence ^47^. To evaluate whether PPARG inverse agonists similarly induce cell cycle arrest, we treated UM-UC-9 cells and observed an increase in cells in G1 phase and a concomitant decrease in cells in S and M phase by flow cytometry (Figure 4D). We also found evidence of PPARG inverse agonists causing a reduction of phosphorylated Retinoblastoma Protein (p-Rb) (S780) levels in cells, consistent with G1 arrest (Supplementary Figure 20B), increasing expression of the senescence-associated secretory phenotype (SASP) genes TNF, CXCL10, CCL5 and IL6 (Supplementary Figure 20C), and increasing the proportion of cells positive for senescence-associated β-galactosidase activity (Supplementary Figure 20D). We therefore conclude that these PPARG inverse agonists induce G1 arrest, cellular senescence, and morphology similar to that induced by the CDK4/6 inhibitor palbociclib.

More broadly, this analysis confirmed the practical utility of JUMP as a universal reference atlas: a low-replicate query (n=2) generated in an external laboratory can successfully retrieve correct biological targets and cell-cycle mechanisms through simple profile-similarity matching against the static database, even across differing cell types. This eliminates the need to run a large reference set of compounds concurrently with novel compounds of unknown mechanism.

### Public portals enable community access and exploration

The web-based portal JUMP Cell Painting Hub serves as a central entry point for JUMP tools and data (http://broad.io/jump; unpublished communication, Alán Fernando Muñoz González). It provides interactive tools to query images and similarity results, how-to guides for common analyses, and guidance to access the latest versions of raw images, single-cell profiles, and all levels of well-level profiles (aggregated, annotated, normalized, feature-selected); from *the Cell Painting Gallery (‘registry.opendata.aws/cellpainting-gallery’).* JUMP_rr (http://broad.io/compound) allows interactive search for compounds with the most similar profiles to a query compound, and displays images and distinctive features (relative to negative controls). Instructions are also provided for clustering compounds’ profiles using the web-based data visualization and analysis software Morpheus (https://software.broadinstitute.org/morpheus/).

JUMP Consortium partner Ardigen developed a free web-based application, phenAID JUMP-CP Data Explorer (https://phenaid.ardigen.com/jumpcpexplorer/), for viewing images and metadata and searching for 100 nearest neighbors. Unlike JUMP_rr, whose similarities are based on cosine similarities between image profiles, phenAID projects compound profiles to the same two-dimensional space using UMAP (uniform manifold approximation and projection) with Euclidean distance and color-coded by clusters (BIRCH clustering algorithm: Balanced Iterative Reducing and Clustering using Hierarchies).

Finally, we made lists of commercially available compounds aiming for maximal Cell Painting profile diversity ^30^ to provide the community with enriched libraries with diverse functionality to use as landmarks (Supplementary File ENRICHED). Overall, of the 17,069 phenotypically active compounds, 10,498 (61.5%) are commercially available (per MolPort).

## Discussion

The JUMP Cell Painting Consortium project answered a crucial question: multiple laboratories using different microscopes across the world can productively combine their Cell Painting data into a unified reference, avoiding the need for each laboratory to run their own landmark or discovery compounds. For most researchers, the 34% loss of signal compared to within-lab datasets (51% versus 77% phenotypic activity for clinical/preclinical compounds) is offset by access to 100,000+ compounds’ profiles in a free, public database. We found that over 15% of compounds show significant phenotypic activity, and because profiles from compounds sharing targets are usually similar, chemical mechanisms of action can be discovered and lead hopping can reveal useful chemical starting points for drug discovery. We also show structure-activity relationships can be studied and activity cliffs identified in the data and some chemical activities (albeit a limited set) can be predicted from image profiles. Already, dozens of researchers have used the JUMP Cell Painting dataset to make discoveries relating to mechanisms of action ^48^, library enrichment and lead hopping ^34^, chemical activity prediction ^49,50^, functional genomics ^51–53^, and virtual phenotypic screening ^54^. The dataset is also popular for improving computational methods, including batch effect correction ^55–57^, phenotype detection ^27^ and prediction ^58–60^, and representation learning ^59,61–70^. JUMP’s data organization makes it navigable by computational biologists and AI agents alike ^71^.

The dataset has limitations which will motivate future data generation: (1) A single cell type, U2OS, was used, apart from pilots ^22^. Although image profiles can sometimes ‘translate’ across cell types (as here for PPARG, plus our unpublished data) and the JUMP Target-2 plate of reference chemicals can assist in alignment, a cell line lacking any of the proteins impacted by the chemical cannot produce a phenotype. (2) Chemicals were tested at a single concentration, 10 μM, except for controls and Source 7. Dose responses can better capture subtle phenotypes and avoid cytotoxic outcomes but increase cost dramatically; JUMP chose to test a larger number of compounds, versus a smaller number studied in dose-response. The OASIS Consortium is creating Cell Painting data alongside transcriptomics and proteomics; its goal of predicting hepatotoxicity mechanisms dictates use of liver-relevant cell models and an 8-point dose response ^37^.

Apart from using JUMP data as-is, as described in the applications here, there are also many opportunities to develop and test improved analytical methodology. The CellProfiler features and Harmony batch correction used here were optimized as independent steps for a single representation, whereas the 34% cross-source signal loss might be reduced by jointly optimizing the full workflow (feature extraction, normalization, and batch correction) across representations including deep learning embeddings, which would be eased by end-to-end automation. Batch correction, in particular, seems to be a major opportunity for improvement ^55^. Single-cell profiles were median-averaged here; methodologies that leverage single-cell data could also improve performance ^72–74^. Quality control methods could be improved, perhaps using machine learning, as each partner performed their own procedures and no methods are proven reliable nor standard yet. The majority of JUMP laboratories captured brightfield images (Supplementary Table 2); these could augment profiles and/or be used to test and improve stain inference methods ^75^. The availability of the JUMP dataset also opens the door for widespread integration of Cell Painting data with-omics data types. Predictions of cultured-cell-omics profiles from image profiles have proven fruitful ^4,76–79^, as has integrating data modalities to make discoveries about chemical or genetic perturbations. Cell Painting provides information complementary to both transcriptomics ^4,5,80,81^ and proteomics ^3^.

The major untapped opportunity afforded by JUMP is the combined analysis of its compound profiles with the previously released genetic perturbations ^21^. Identifying genes that match compounds can identify a novel compound’s target or mechanism, and identifying compounds that match genes enables phenotype-driven virtual screening ^7^. However, the workflows that maximized signals differ between the JUMP genetic and chemical perturbations, yielding different feature spaces and thus impeding direct matching. So far, our explorations in the full JUMP dataset show little successful matching of genes to their annotated chemical inhibitors (results not shown), consistent with our prior analyses of JUMP subsets: a similarity metric learned on top of classical image features (via a bilinear transformation) did not rank known gene-compound pairs better than random chance in a large subset of JUMP ^45^, nor did our attempt to align the two profile types in a shared space in a smaller JUMP pilot dataset ^22^. Message-passing graph neural networks show improvement over simple baselines under some conditions ^45^, but gene-compound matching remains an open challenge.

The JUMP Cell Painting consortium brought together 10 pharmaceutical/biotechnology companies, two non-profit organizations, and six technology partners to create a freely-available reference resource with ∼127 TB of images, plus profiles, covering 1.81 billion cells and 115,796 compounds, 73,208 unique chemical scaffolds, and 15,000+ genetic perturbations ^21^, plus processing pipelines. The JUMP-Target-2 plate provides standard landmark compounds for any laboratory to align new experiments to the JUMP reference, extending the resource. Future datasets that explore disease-relevant cell types, multiple concentrations per compound, and additional compounds and biologic perturbations would be most welcome.

JUMP is a validated, publicly available reference for identifying compounds with useful behavior and exploring compound mechanisms. We invite the scientific community to make maximal use of it for drug discovery, basic biological research, and the development of new computational methods for image-based profiling.

## Supporting information

Supplementary_File_ACTIVITY_CLIFF_PAIRS

Supplementary_File_ACTIVITY_CLIFF_SCAFFOLD_GROUPS

Supplementary_File_CHEMBL

Supplementary_File_ENRICHED

Supplementary_File_FUC

Supplementary_File_FUP

Supplementary_File_PHENOTYPIC_CLIFFS

Supplementary_File_PHENOTYPIC_CONSISTENCY

## Methods

The JUMP Cell Painting dataset is a collection of several datasets that were either generated or reprocessed by the JUMP Cell Painting Consortium. The primary dataset (cpg0016; production data) was generated during the data production phase of the JUMP-CP project. It comprises a compound dataset (compound production data, analyzed in this study), an Open Reading Frame (ORF) overexpression dataset (ORF production data) and a CRISPR knockout dataset (CRISPR production data). In addition, three datasets were generated during the pilot phase of the project (pilot data). These include a previously released compound dataset to test different experimental/staining conditions (cpg0001; CPPilot data ^1^), another previously released dataset to test different perturbation modalities (cpg0000; CPJUMP1 data ^22^) and a compound dataset to test different microscopes (cpg0002; CPJUMP-Scope1 data ^23^). In the following sections, unless specified explicitly, all the details about experimental conditions and analysis pertain to the cpg0016 data. Details about the other datasets are described in other manuscripts describing those datasets.

## Data and code availability

The chemical perturbation data analyzed in this study are part of cpg0016-jump and are publicly available in the Cell Painting Gallery ^18^ at https://registry.opendata.aws/cellpainting-gallery/. The Gallery holds the field-of-view images, the CellProfiler single-cell profiles, the aggregated well-level profiles, and the deep-learning embeddings, although not every data product is available for every source. The JUMP datasets repository (https://github.com/jump-cellpainting/datasets provides metadata, profile manifests, and download instructions, and the Cell Painting Gallery folder-structure documentation (https://github.com/broadinstitute/cellpainting-gallery/blob/main/folder_structure.md) describes how the files are organized.

For the primary compound analyses, we used the v1.0 compound_no_source7 CellProfiler profiles (https://cellpainting-gallery.s3.amazonaws.com/cpg0042-chandrasekaran-jump/source_all/workspace/profiles_assembled/compound_no_source7/v1.0/profiles_var_mad_int_featselect_harmony.parquet), which exclude source 7 and were corrected for batch effects with Harmony. We also compared the v1.0 compound_DL_CPCNN_no_source7 profiles (https://cellpainting-gallery.s3.amazonaws.com/cpg0042-chandrasekaran-jump/source_all/workspace/profiles_assembled/compound_DL_CPCNN_no_source7/v1.0/profiles_dropna_var_mad_int_featselect_harmony.parquet), computed with the Cell Painting CNN (CP-CNN), with the CellProfiler profiles for phenotypic activity, phenotypic consistency, and the batch effects that remain between sources after correction. EfficientNet profiles (https://cellpainting-gallery.s3.amazonaws.com/cpg0042-chandrasekaran-jump/source_all/workspace/profiles_assembled/compound_DL_EFNET/v1.0/profiles_dropna_var_mad_int_featselect_harmony.parquet) are also public, but we did not analyze them in this study. Cellpose segmentations and cell centers are available in the Cell Painting Gallery, together with the workflow that produced them (https://github.com/theislab/jump-cpg0016-segmentation), but we did not evaluate them in this paper.

The JUMP_rr tables computed from the same compound_no_source7 profiles used in this paper are archived ^82^. The archive contains compound similarity matches, CellProfiler feature rankings, metadata, checksums, and a record of how each file was produced. A separate archive ^83^ contains consensus profiles for phenotypically active compound, ORF, and CRISPR perturbations. The archives do not include all of the annotation tables used in the analyses, and the sources of those tables are listed under Compound annotation.

The jump-profiling-recipe (https://github.com/broadinstitute/jump-profiling-recipe) contains the workflow that generated the profiles from the images.

The JUMP Hub (https://broad.io/jump) and JUMP_rr provide access to current JUMP resources and tools. Terra is a scalable and secure platform for biomedical researchers to access data, run analysis tools, and collaborate. It is co-developed by the Broad Institute of MIT and Harvard, Microsoft, and Verily. Instructions for accessing JUMP profiles through Terra (https://github.com/broadinstitute/jump_hub/blob/195d51d710addbd62e7a08a5eaffcede3c5a5993/misc/terra_workspace.md) are available from the JUMP Hub. The Hub and JUMP_rr are updated over time. The manifests and archives cited above record the data versions used in this paper.

### Chemical perturbations

#### Chemical perturbation selection

For generating the production data, the JUMP consortium partners nominated, provided, and profiled 115,796 unique compounds, including controls, which are described in a later section. These nominated compounds meet the following criteria:

- Structure of the compound was in the public domain or could be released by the company.
- The purity of the compound is either greater than 90% or is at least 80% with additional quality control data.
- Lastly, the nominated compounds should not include any controlled substances, to facilitate sharing the compounds across international borders.

#### Controls

We included several controls to identify and/or correct for different experimental artifacts. The controls can be broadly categorized as within-plate controls (those that are on the same plate as the treatments), and control plates (entire plates that are run alongside the treatment plates).

#### Within-plate controls

Within-plate controls include negative controls, positive controls, and untreated wells.

### Negative control wells

Dimethyl Sulfoxide (DMSO) wells are used as a vehicle negative control for detecting and correcting plate to plate variations and as a baseline for identifying perturbations with a detectable morphological signal.

### Positive control wells

We selected eight compound controls using profiles from the standard U2OS, 48-hour compound condition of the CPJUMP1 JUMP-Target-1 experiment ^22^. Candidate compounds were required to have a mean cell count greater than one-quarter of the mean for DMSO, a DMSO-scaled replicate-correlation score (sim_scaled_mean_ref_i_mean_i, termed replication grit) greater than the mean plus three standard deviations for DMSO wells, and a measured purity of at least 75% in the pilot sample. DMSO wells occupying the same position across replicate plates were treated as replicates when calculating the reference distribution. These criteria yielded 53 candidate compounds.

Normalized, feature-selected, outlier-trimmed well profiles were median-aggregated across replicate wells for each candidate, and Pearson correlations were calculated between the resulting compound consensus profiles. We applied a modified correlation-pruning procedure over signed Pearson-correlation cutoffs from 0.05 to 0.30 in increments of 0.001, successively removing compounds from pairs whose correlation exceeded the cutoff. A cutoff of 0.28 was the first to retain exactly eight compounds; thus, every retained pair had a signed Pearson correlation of 0.28 or less. Because signed rather than absolute correlations were used, anticorrelated profiles were considered distinct phenotypes. We then inspected images and feature patterns for the selected compounds and verified compound identities and commercial availability before finalizing the panel. The controls were therefore selected for reproducible, mutually distinct Cell Painting phenotypes rather than uniformly strong activity and are not conventional strong positive controls (see Supplementary Figure 9C, D). Their identities are provided in Supplementary Table 6.

### Untreated wells

Some wells were treated neither with treatments nor with controls; they contained only cells. These wells are unfortunate practical limitations, primarily useful for recognizing the plate orientation. They might also be used for normalizing samples, but they are non-ideal negative controls, given that they are not mock-treated with any reagents.

### Control plates

#### Negative control plates

Some data acquisition partners ran negative control plates periodically (e.g., one per batch, or at the beginning and end of each batch). These plates can help disentangle staining and imaging artifacts, including batch effects and well-position effects. Examples include:

- An entire plate of untreated cells.
- An entire plate where all wells were treated with DMSO.

#### JUMP-Target-2-Compound plates

Most partners ran one or more plates of JUMP-Target-2-Compound compounds in each batch to correct batch effects (unrelated plates from the same batch correlate strongly with each other compared to related plates in other batches). Source 1 did not run any Target-2 plates and analysis of Source 9 indicates an accidental lack of compound treatment for Target-2 plates only.

#### JUMP-Target-1-Compound plates

To align the production data (cpg0016) with the CPJUMP1 experiment in the pilot data (cpg0000), we ran four JUMP-Target-1-Compound plates in one batch of cpg0016 from source_4 (Broad). These plates were not used in the analyses reported in this paper.

#### JUMP-Target-2-Compound plates with polybrene

To quantify the effects of adding polybrene, a viral transfection agent that is added to the ORF experiment, on the morphology of the cell, we ran two JUMP-Target-2-Compound plates with polybrene in one batch. These plates were not used in the analyses reported in this paper.

### Plate layout design

#### Compound plates

The 384-well plate layouts (Supplementary Figure 1) were designed such that the controls are in the outer columns and the treatment wells are in the inner columns of the plate, to minimize well-position effects (also called plate-layout effects, whereby identical treatments in outer wells of a plate show different results than the same treatment positioned in inner ones) for the treatment samples, and to be compatible with partners’ instrumentation and logistics. Four replicates of all eight compound positive controls are added to the outermost columns 1 and 24, while DMSO is added to the next-innermost columns, 2 and 23. The remaining wells contain one replicate of compound treatments. In the case of 1536-well plates (Supplementary Figure 2), each quadrant of the plate follows the same layout as a 384-well plate.

#### JUMP-Target-1-Compound plates

The plates contain 301 compounds and DMSO; all but 14 compounds are in singlicates. There are 64 DMSO wells that are spread across the plate. The 14 compounds with two replicates are diverse positive controls that are different from the positive controls on the production plate. Additional details about these positive controls and the criteria met by the compounds of the plate have been described ^22^.

#### JUMP-Target-2-Compound plates

The contents of the plate are the same as that of JUMP-Target-1-Compound plates, but the layout is different (Supplementary Figure 3). JUMP-Target-2-Compound plates meet all the criteria met by JUMP-Target-1-Compound plates ^1^. Both layouts are provided in our GitHub repository – https://github.com/jump-cellpainting/JUMP-Target) and the compound list is provided also as Supplementary File TARGET2.

### Perturbation identifier

All compound, ORF, and CRISPR perturbations are assigned a unique JUMP identifier. These identifiers start with the code JCP2022_.

### Compound annotation

Compound annotations were assembled from the frozen sources below. Unless stated otherwise, counts refer to the canonical 112,462-compound analyzed roster.

**ChEMBL** protein target annotations were generated using bioactivity data from ChEMBL release 33. On execution, the accompanying reproducible workflow automatically downloads the official ChEMBL 33 SQLite distribution when it is not already available locally and queries this release-pinned database. JUMP compounds were mapped to ChEMBL by exact structure matching: JUMP SMILES were standardized with the ChEMBL structure-standardization pipeline and matched to ChEMBL structures in SmallWorld’s chembl_31.anon collection using a topological search restricted to identical graphs, disregarding stereochemistry ^84^. For matched compounds, recorded bioactivities were filtered to high-confidence, well-defined measurements: ChEMBL confidence score ≥ 6 (homologous or direct protein-complex subunits, or homologous or direct single-protein targets), binding and functional assays only, exact (non-censored) activity relations, and standard activity types of IC_50_, EC_50_, AC_50_, K_i_, K_d_, or Potency. Records carrying a data-validity comment or flagged as potential duplicates were discarded, and pChEMBL values were computed from the reported standard activity in nM, μM, or uM when not supplied by ChEMBL. ChEMBL canonical structures were additionally standardized, recognized salt and solvent fragments were removed, and unresolved multicomponent structures were excluded. Targets were mapped to UniProt accessions, and successful single-accession mappings defined the set treated as single-protein targets. For each compound–target pair, pChEMBL values were averaged across all retained activity records, and a pair was labelled active when the mean pChEMBL exceeded 5 (10 μM) and inactive otherwise; both classes were retained so that inactive compounds could serve as negatives. Only targets with at least 15 active and 15 inactive compounds were kept. Target eligibility was determined before duplicate standardized structure-target pairs were resolved. The resulting dataset (targetannotations_singleproteins_jumpcompounds_all_5.csv) comprised 68,211 compound–target annotations (24,029 active, 44,182 inactive) spanning 17,892 distinct compounds and 102 single-protein targets, 77 of which are human. This ChEMBL target set, represented by UniProt accession columns in the supplied wide table and corresponding to Metadata_Uniprot_target in downstream study tables, was used for all target-based analyses in this study. Extraction and processing code together with the required inputs and generated outputs, is available at src/jump_production/downloading/chembl_targets_sqlite.py in the paper’s GitHub repository (code) and archived as Supplementary File CHEMBL and at targetannotations_singleproteins_jumpcompounds_all_5.csv in https://zenodo.org/records/21800106 (data).

**Clinical development stage and Drug Repurposing Hub target/MOA annotations** were obtained from the 2020-03-24 Drug Repurposing Hub samples and drugs exports (Corsello et al., 2017), joined on the Hub compound name and matched to JUMP compounds by the first 14 characters of the InChIKey. Only Hub records carrying a target annotation were retained, so clinical phase and MOA derive from target-annotated records.

**Chemical Probes Portal** target annotations were taken from a Chemical Probes Portal export (Sanfelice et al., 2025). Compound and target tables were joined, structures were standardized with RDKit, entries flagged as probes were retained, duplicate standardized InChIKeys were resolved by probe quality, and full InChIKeys were exact-matched to JUMP compounds.

**MOTIVE** (Arevalo, Su, Carpenter, et al., 2024) compound-gene annotations (Zenodo record 18197517), integrating BioKG, DGIdb, DrugRep, Hetionet, OpenBioLink, OpenTargets, PharmeBiNet, and PrimeKG, were matched to JUMP compounds by the first 14 InChIKey characters. A compound was counted as having a target or MOA annotation (Figure 1F) if it carried a ChEMBL/UniProt target, a Drug Repurposing Hub target or MOA, a Chemical Probes Portal target gene, or any MOTIVE gene annotation.

**External database mapping** used a frozen UniChem/MyChem InChIKey pointer table from the same Zenodo record, spanning 39 external chemical databases; the live services were not queried. A compound was mapped if its full InChIKey appeared in the pointer table with at least one database.

**Commercial availability** was assessed against a frozen MolPort batch quote archive (Jan 24, 2026). Submitted SMILES were deduplicated, Perfect and Isomer hit types were accepted, the minimum quoted price was required to be at or below $200, and quotes were joined to JUMP compounds by exact match between the submitted SMILES and the metadata SMILES.

### Experimental conditions

We optimized several experimental parameters in the pilot experiments before embarking on data production; therefore, those experiments used a variety of parameters and did not all use the optimized final parameters. Experimental conditions of the pilot experiments and their optimization have been described ^1,22^. The following are the experimental parameters used for generating the production data (cpg0016).

- Cell Painting dyes: We modified the concentrations of many dyes to improve their performance while simultaneously making the Cell Painting assay cheaper; the final choices are as described ^1^.
- Cell type: We compared A549 and U2OS cell lines in pilot experiments (cpg0000) ^22^ and settled on using U2OS because the performance of both cell lines were similar and there are previous datasets generated using U2OS, which allows comparison across experiments.
- Time point: We compared different time points for each perturbation modality (cpg0000) ^22^ and settled on the following time points: Compound – 48 hours, ORF – 48 hours, CRISPR – 96 hours.
- Reagent vendor: Given the similar performance of the Cell Painting dyes from both vendors, Thermo Fisher and Revvity, Inc. (cpg0001) ^1^, we performed our experiments with dyes from the latter, also known as PhenoVue^TM^ Cell Painting Kit, 2.0 (part number PING22, Revvity, Inc., Waltham, MA). Revvity, Inc. donated PhenoVue kits to the consortium.
- Microtiter plates: After comparing different plates for their ability to minimize evaporation in the outer wells (cpg0000) ^1^, we used Revvity, Inc. Cell Carrier Ultra for our data production. All laboratories used 384-well plates, except for source_1 and source_9 that used 1536-well plates.
- Microscope settings: Microscope settings are described in the sample preparation and image acquisition section below.
- Number of fields of view: We captured six (source_2 and source_10) or nine (source_3 through source_8, source_11, source_13, and source_15) fields of view in the 384-well plates and four fields of view in 1536-well plates (source_1 and source_9).
- Concentration of compounds: The treatment compounds were assayed at 10 μM at all laboratories, apart from Source 7 where the compounds were assayed at 0.625 μM (the goal being to assay some of the compounds at a low concentration in addition to the higher concentration used for most of data production). The positive control compounds in compound, ORF and CRISPR plates were assayed at 5 μM. JUMP-Target-1-Compound and JUMP-Target-2-Compound plates were also assayed at 5 μM. In all cases, DMSO was 0.2%.

### Experiment design

The production data was generated at 12 laboratory sites, typically called sources in this paper, around the world (Figure 1A and Supplementary Table 1). Compound data-producing partners are divided into two groups, wave 1 (7 partners) and wave 2 (3 partners). Additionally, there are two other data producing partners, one (source_4, the Broad) generated the 5-replicate ORF data and the other (source_13) generated a 5-replicate compound dataset and the 5-replicate CRISPR dataset. Each wave 1 partner shared their ∼12,000 nominated compounds with four other wave 1 partners and received compounds from four other wave 1 partners. Thus, each partner screened roughly five times the number of compounds they nominated. Thus, each compound was typically screened in five replicates across five laboratories. Each wave 2 partner similarly shared their nominated compounds with the other two partners and received compounds from the other two partners. Each wave 2 partner then screened three replicates of the compounds they nominated along with a single replicate of the compounds they received. Thus, each compound was screened in five replicates across three laboratories. Nearly 6000 compounds were common between wave 1 and wave 2 partners, to help align wave 1 and wave 2 compound production datasets. The plates were grouped into batches. The number of plates in each batch is different for different sources. In each batch, along with the treatment plates, most partners ran at least one untreated or DMSO plate and one JUMP-Target-2-Compound plate, so that the different batches can be aligned within a source and across sources.

### Sample preparation and image acquisition

In the Cell Painting assay, eight cellular components are stained with six fluorescent dyes and imaged in five channels: 1) mitochondria (MitoTracker; Mito), 2) nucleus (Hoechst; DNA), 3) nucleoli and cytoplasmic RNA (PhenoVue 512 Nucleic Acid; RNA), 4) endoplasmic reticulum (concanavalin A; ER), 5) Golgi and plasma membrane (wheat germ agglutinin (WGA); AGP) and the actin cytoskeleton (phalloidin; AGP). We followed the optimized Cell Painting assay ^1^; the differences between the v2 ^85^ and v3 protocols and any future updates are available at https://github.com/carpenter-singh-lab/2023_Cimini_NatureProtocols/wiki. Five types of microscope were used to acquire images at the 12 data production laboratories. These included the Revvity, Inc. Opera Phenix, ImageXpress Micro confocal, Yokogawa CV8000, Yokogawa CV7000, and Revvity, Inc. Operetta. Images were acquired in either widefield or confocal modes, and one or three brightfield planes were also acquired at some laboratories, apart from the fluorescent channels. The settings for each microscope and the filter configuration are provided in Supplementary Tables 2-5.

### Image processing

We used CellProfiler bioimage analysis software (versions 4.0.7, 4.1.3 or 4.2.1; see Supplementary Table 7) to process the images. CellProfiler uses classical algorithms to measure cellular features. We corrected for variations in background illumination intensity and then segmented cells, with each partner adjusting segmentation parameters as needed for their image set. We then measured several categories of features such as intensity, granularity, texture, and density for each cell in each fluorescent channel and each brightfield plane, if captured. Similarly, we measured features at the whole-image level. For more details, see http://broad.io/cellprofilermanual and the image analysis pipeline https://github.com/broadinstitute/imaging-platform-pipelines/tree/master/JUMP_production#production-pipelines). Ultimately, we measured up to 7648 features (including features measured at the per-cell level and those measured only at the per-image level) for fields of view that included three brightfield planes (each measured separately in the pipeline); for sources that collected fluorescence images only, the profiles contained 4762 features for each cell). CellProfiler was executed in headless mode, using either Amazon Web Services (AWS) via Distributed-CellProfiler ^86^ or standard high-performance computing clusters.

### Profile processing

Single-cell CellProfiler features were aggregated to mean well-level profiles and processed using the jump-profiling-recipe pipeline (https://github.com/broadinstitute/jump-profiling-recipe), which uses a bundled copy of pycytominer ^87^ operations for feature selection. Low-variance features were removed, and each remaining feature was normalized per plate by subtracting the median and dividing by the median absolute deviation computed across negative control wells (DMSO). Features were then transformed to a standard normal distribution via rank-based inverse normal transformation. Redundant features (Pearson correlation > 0.9), blocklisted features, and features containing missing values were removed. Batch effects across plates were corrected using Harmony ^29^ with Metadata_Batch as the batch key (300 clusters, 20 iterations).

### Chemical space characterization

All cheminformatics analyses were performed using RDKit (https://www.rdkit.org). For each compound, a panel of physicochemical descriptors was computed from SMILES representations: molecular weight, octanol-water partition coefficient (LogP), topological polar surface area, hydrogen bond donor and acceptor counts, rotatable bond count, ring count, aromatic ring count, heavy atom count, fraction of sp3-hybridized carbons, and the quantitative estimate of drug-likeness (QED) score ^24^. Lipinski Rule of Five compliance ^31^ was assessed by counting violations of the four criteria: molecular weight > 500 Da, LogP > 5, hydrogen bond donors > 5, and hydrogen bond acceptors > 10. Bemis-Murcko scaffolds ^88^ were extracted to assess structural diversity across the compound collection. Compounds were screened for PAINS substructures ^25^ using the RDKit FilterCatalog implementation.

Morgan circular fingerprints (radius 2, 2048 bits) were computed for each compound. To visualize the structural chemical space, fingerprint vectors were L2-normalized and embedded via UMAP ^89^ using cosine distance (n_neighbors = 15, min_dist = 0.1), which approximates the Tanimoto coefficient for binary fingerprints. The L2-normalization and cosine approximation to Tanimoto similarity are described in detail in the Structure-phenotype analysis section. A complementary property-space UMAP was computed from the 11 continuous descriptors listed above after z-score standardization, using Euclidean distance with the same neighbor and minimum distance parameters (n_neighbors = 15, min_dist = 0.1). GPU-accelerated neighbor search and UMAP embedding were performed via rapids_singlecell ^90^ when available; otherwise scanpy ^91^ was used on CPU.

### Functional use classification

Functional use categories were obtained using the EPA functional use classification pipeline ^26^. JUMP compounds were mapped to DSSTox substance identifiers (DTXSIDs), the unique chemical identifiers assigned by the US EPA’s Distributed Structure-Searchable Toxicity (DSSTox) database, which curates chemical structures linked to toxicity and exposure data. Of 112,462 JUMP compounds (excluding source 7), 10,451 were mapped to DTXSIDs and submitted for classification, yielding 917 compounds with curated functional use categories (FUCs; 107 categories) and 9,680 compounds with predicted functional use probabilities (FUPs; 35 classes). After restricting to compounds present in the JUMP analysis set and filtering by copairs phenotypic activity calls, 8,300 compounds remained, of which 427 carried curated FUCs (across 97 represented categories) and 7,651 carried FUPs; these are the sets used for the enrichment analyses. The pipeline returns two outputs per compound: (i) FUCs, discrete labels assigned by expert review of reported industrial/consumer uses, and (ii) FUPs, predicted probabilities from random-forest quantitative structure-use relationship (QSUR) models ^26^ trained on structural and physicochemical descriptors.

Associations between functional use and Cell Painting activity (copairs corrected p < 0.10) were analyzed separately for FUCs and FUPs. FUCs were tested by two-sided Fisher’s exact tests among compounds with at least one curated FUC, requiring at least 20 compounds per group and five observations per table cell. FUPs were tested by logistic regression on continuous probabilities among compounds with non-null endpoint values; odds ratios represent an interquartile-range increase. FUP endpoints required at least 50 observations, five active and five inactive compounds, two unique probabilities, and a positive interquartile range. The probability >= 0.5 threshold was used only descriptively. Across the 8,300 overlapping compounds, the mean was 1.99 classes per compound (median 2, maximum 9), and 4,790 (57.7%) had at least two classes. Benjamini-Hochberg correction was applied separately to FUCs and FUPs; FDR < 0.05 indicated significance.

### Phenotypic activity and consistency metrics

Phenotypic activity and target consistency were quantified using the mean Average Precision (mAP) framework implemented in the copairs library ^27^. All similarity computations used cosine distance over the full morphological feature space.

For phenotypic activity, Average Precision was computed per replicate by defining positive pairs as profiles sharing the same compound identity (Metadata_JCP2022) and negative pairs as DMSO controls on the same plate. Plate-level blocking ensured that positive and negative pairs were drawn from comparable imaging conditions. Average Precision scores were then averaged across all replicates of each compound to yield a single mAP value. Statistical significance was assessed by permuting compound labels 100,000 times to generate a null distribution, and the resulting p-values were corrected for multiple testing using the Benjamini-Hochberg procedure ^92^. Compounds with a corrected p-value below 0.10 were classified as phenotypically active.

Target consistency was evaluated using the same mAP framework but with a different pairing structure. Positive pairs comprised compounds sharing a given annotation (e.g., protein target, mechanism of action), while negative pairs comprised compounds differing on that annotation. Because individual compounds may map to multiple targets, the multilabel formulation in copairs was used, splitting pipe-separated annotation values and computing Average Precision accordingly. Prior to consistency analysis, replicates were aggregated to a single median profile per compound, and only compounds classified as phenotypically active (corrected p < 0.10) were retained. The consistency significance threshold was set at corrected p < 0.05 with the same 100,000-permutation null. Consistency was evaluated across nine annotation label sets drawn from five resource families: Drug Repurposing Hub targets ^93^, UniProt targets from ChEMBL ^94^, Chemical Probes portal targets ^95^, mechanism of action and disease area from the Drug Repurposing Hub, three knowledge-graph-based target sets from MOTIVE ^45^ using BioKG, OpenTargets ^96^, and PrimeKG ^97^, and ToxCast active assay annotations ^98^.

Note that four related scores are used: Average precision (AP) scores one query profile; normalized average precision (nAP) rescales AP so that the chance expectation is zero; mean average precision (mAP) averages raw AP over the queries in a compound or annotation group; and mean normalized average precision (nMAP) averages nAP over those queries. Normalization is applied per query before averaging.

#### Associations of chemical properties with phenotypic activity and consistency

Continuous properties were related to phenotypic activity by Spearman correlation, with Benjamini-Hochberg correction within each panel; categorical annotations were tested on the binary activity call by Fisher’s exact tests (logistic regression for continuous traits), with Benjamini-Hochberg correction within each family. For consistency, each compound was scored by target-alignment nAP: its retrieval of phenotypically active compounds sharing its OpenTargets target (via MOTIVE) against active compounds annotated only to unrelated targets, excluding the query, for targets with enough analyzed and active compounds; each compound is shown at its best-scoring target. The across-target curve in Figure 3A is a least-squares fit of target-mean consistency on target-mean activity and its square, adjusted for annotation degree and active-reference count.

Chemical collections analyzed include KCGS (https://www.sgc-unc.org/kcgs), PKIS ^99^, and Probes (https://www.thesgc.org/chemical-probes).

### Structural class enrichment

Three families of structural classes were tested for association with phenotypic activity among analyzed compounds with a valid structure: RDKit functional groups, BRICS (Breaking Retrosynthetically Identified Chemical Substructures) fragments, and the Bemis-Murcko scaffolds described under Chemical space characterization. For each class, phenotypic activity was compared between compounds carrying the class and all other valid-structure compounds by a two-sided Fisher’s exact test, reported as an odds ratio with a confidence interval; classes with too few compounds in any cell of the table were not tested. Benjamini-Hochberg correction was applied separately within each family.

### Morphology UMAP visualization

Global structure of the morphological space was visualized with UMAP ^89^ applied to the Harmony batch-corrected, feature-selected CellProfiler profiles described in *Profile processing* (source 7 excluded). Embeddings were computed at two levels: a well-level embedding that retains each profile’s data-generating source (used for the source-colored view in Supplementary Figure 8A) and a perturbation-level embedding in which replicates are mean-aggregated to one profile per compound (used for Figure 2A and the per-compound views). In both cases, profiles were reduced to 50 principal components, a nearest-neighbor graph was built on that PCA embedding with n_neighbors = 15 using cosine distance, and the graph was embedded with min_dist = 0.1 and a fixed random seed (42). GPU-accelerated PCA, neighbor search, and UMAP were performed via rapids_singlecell ^90^ when available (approximate Inverted File with Flat quantization (IVF-Flat) neighbor search for the cosine metric); otherwise scanpy ^91^ was used on CPU. GPU UMAP is not bitwise reproducible, but the global cosine layout was stable across random seeds (mean Procrustes correlation 0.97, minimum 0.93 across five seeds at the perturbation level).

The embeddings are used for display only: phenotypic activity, consistency, and all other quantitative results were computed from the high-dimensional profiles, not from UMAP coordinates. For visualization, extreme outliers were excluded from the plotted axis range by an interquartile-range rule (axis bounds set three interquartile ranges below the first quartile and above the third quartile on each axis); no points were removed from the embedding itself.

### Within-source and pooled-source reproducibility analysis

To assess whether morphological signatures were reproducible across laboratories, the mAP framework was applied to compounds profiled on JUMP-Target-2 plates, which were shared across contributing sources. We used well-level, harmonized imaging profiles from JUMP-Target-2 plates to estimate how target- and MOA-level phenotypic consistency changed with replicate depth (count). Seven usable sources were included (source_2, source_3, source_5, source_6, source_8, source_10, and source_11); source_9 plates were excluded because of anomalously low treatment activity, and source_7 was excluded upstream from the main compound analysis. source_4 and source_13 profiled genetic perturbations and therefore all the plates were excluded from the harmonized compound profile, although JUMP-Target-2 plates are present in the sources. Source_15 was excluded entirely because of a missing Cell Painting channel. A replicate was defined as one treatment well from one experimental batch. For each replicate depth R=1-8, one quality control-passing plate was selected per sampled batch, one well per compound was selected from that plate, and imaging profiles from the selected wells for each compound were median-aggregated.

All 293 Target-2 treatment compounds (301 unique compounds - 8 positive controls) were eligible at every replicate depth, with no phenotypic-activity filter. For each annotation type, consistency was computed over compounds carrying the corresponding target or MOA annotation. Phenotypic consistency was recomputed using the same multilabel copairs average-precision framework used for the main target/MOA consistency analysis, with cosine distance, positive pairs defined by shared target or MOA labels, and negative pairs defined by no shared labels. For each target or MOA group, phenotypic consistency was summarized as group-level nMAP. For each replicate depth, the saturation curve was computed as the mean nMAP across the selected groups.

We evaluated two group-level reporting views. The all-group view included all scored target or MOA groups represented by at least two Target-2 compounds. The significant-group view included the subset of groups that were significant in the main consistency_no_target2 analysis at corrected p < 0.05 and were represented by at least two Target-2 compounds (Figure 2C). In the within-source analysis, batches were sampled separately within each source. All R-batch combinations were enumerated when the number of possible combinations was ≤100; otherwise, 100 random R-batch subsets were sampled. In the pooled-source analysis, 100 random (source, batch) subsets were sampled for each R. The 90% knee was defined as the smallest R at which the mean nMAP across the selected groups reached at least 90% of the value at the largest available R for that curve.

### Batch-effect quantification

Residual cross-source batch effects after Harmony correction were quantified with two local neighborhood-integration metrics from the single-cell integration benchmarking (scIB benchmark) ^100^ on the 50-dimensional PCA embedding of each profile set (CellProfiler and CP-CNN deep-learning features, with and without Source 7; ∼750,000 wells each). For every well, the k = 90 nearest neighbors were identified by cosine distance (IVF-Flat approximate search on GPU). The graph integration Local Inverse Simpson’s Index (graph iLISI, a kNN-graph extension of iLISI ^29^, reports the effective number of distinct sources among a well’s neighbors (higher indicates better mixing); k-nearest-neighbour batch effect test (kBET) ^101^ tests whether each local neighborhood’s source composition matches the global composition, reported here as the acceptance rate (the fraction of neighborhoods consistent with the global mix). We report both for CellProfiler and CP-CNN features (CellProfiler iLISI 0.318 / kBET 0.0071; CP-CNN 0.092 / 0.0003).

### CP-CNN deep-learning feature comparison

We compared the Harmony-corrected CellProfiler well profiles with Cell Painting CNN (CP-CNN) well profiles (Moshkov et al., 2024). The same copairs activity analysis was applied to both representations, and concordance is the fraction called active by both or by neither; consistency was compared on the union of compounds called active by either representation.

### Cell density and phenotypic activity

Per-well cell density (cells per imaged area) was regressed on source, robustly standardized within plate (median/MAD), and median-aggregated per compound; its Spearman correlation with activity nMAP was computed over all the analyzed compounds.

### Structure-phenotype analysis

Chemical structures were represented as 2048-bit Morgan fingerprints with radius 2, computed using RDKit (https://www.rdkit.org), corresponding to the extended-connectivity fingerprint ECFP4 ^102^. To enable efficient nearest-neighbor computation, fingerprint vectors were L2-normalized and compared using cosine similarity, which closely approximates the Tanimoto coefficient for binary fingerprints (Spearman rho = 0.998).

The relationship between chemical structure and morphological phenotype was assessed by comparing neighborhoods in the two spaces. For each phenotypically active compound, the k = 50 nearest neighbors were identified independently in structural space (L2-normalized fingerprints) and in morphological space (L2-normalized CellProfiler feature profiles), both using cosine distance. Neighborhood overlap for each compound was quantified as the Jaccard index of the two k-nearest-neighbor sets - that is, the number of neighbors shared between structural and morphological spaces divided by the size of their union.

To assess whether specific mechanisms of action exhibited stronger structure-phenotype coherence than expected by chance, compounds were annotated with mechanism-of-action labels from the Drug Repurposing Hub. For each mechanism represented by at least 10 compounds, the mean neighborhood overlap across its members was computed and expressed as a fold enrichment over the random expectation, defined as k / (2N) where N is the total number of compounds. This expected value reflects the probability that two independently drawn k-nearest-neighbor sets of size k from N compounds share a given element.

Scaffold coherence was tested for each Bemis-Murcko scaffold with at least 5 phenotypically active compounds: the mean pairwise cosine similarity among member profiles was compared with 1,000 size-matched random groups drawn from all active profiles. Conversely, active profiles were reduced to 50 principal components and clustered by MiniBatchKMeans (80 clusters); over-representation of each scaffold in each cluster was tested by a one-sided Fisher exact test.

For the related analysis in Figure 2B, each eligible phenotypically active, MOA-annotated compound, compounds with the same Bemis-Murcko scaffold were excluded before the remaining compounds were ranked by cosine similarity between high-dimensional perturbation profiles. To define the null expectation, complete multilabel annotation sets were permuted among compounds 10,000 times, preserving the number and prevalence of MOA labels, including multilabel combinations, while breaking their relationship to phenotype; the gray band is the resulting 95% interval. Excluding compounds with the same scaffold focuses the evaluation on cases where structural-similarity approaches struggle, though results were similar either way: the maximum difference along the curve was 1.81 percentage points at k=4; mean difference was 1.15 points, with the permutation null remaining essentially unchanged.

### Polypharmacology analysis

To analyse whether a compound annotated as engaging two different targets produces a profile resembling a combination of two compounds that have one target each, we assembled trios from Drug Repurposing Hub annotations, each comprising one compound annotated as hitting exactly two targets (A and B), one annotated as hitting only A, and one as hitting only B, requiring all three to be phenotypically active. Within each trio we computed cosine similarities between the dual-target profile and each single-target profile, between the two single-target profiles [sim(A,B)], and between the dual-target profile and the element-wise mean (“blend”) of the two single-target profiles, using one perturbation-level profile per compound. Retaining the highest-activity trio per target pair and excluding any trio in which the least-active of the three members had nMAP < 0.5 left 28 target pairs (of 32). Each pair was classified into one of five classes (convergent targets, divergent-from-shared, additive polypharmacology, dominant target, or emergent phenotype) using a cosine threshold of 0.3, based on whether the two single-target phenotypes were similar [sim(A,B) ≥ 0.3] and whether the dual-target profile resembled one/both single targets or their blend. Additivity was summarized per pair as the difference between the dual-to-blend similarity and the larger of the two dual-to-single similarities.

### Activity cliff detection

Two related analyses were performed: an exhaustive screen for on/off activity cliffs, structurally similar pairs in which one compound is phenotypically active and the other is not; an active-only nearest-neighbor screen for phenotypic cliffs, structurally similar pairs of active compounds with divergent Cell Painting profiles. The first uses the 112,524 profiled compounds (excluding Source 7) with a valid Morgan fingerprint. Relative to the 112,462 compounds in the canonical activity analysis, this set omits the few assessed compounds whose fingerprint is empty and retains 141 profiled compounds without an activity assessment, mainly compounds present only on Target-2 plates plus the positive controls. The second uses the 17,061 phenotypically active compounds with a valid fingerprint.

For the on/off screen, all compound pairs with Morgan-fingerprint Tanimoto similarity of at least 0.70 were enumerated, and pairs in which exactly one compound was phenotypically active were designated activity cliffs (Supplementary File ACTIVITY_CLIFF_PAIRS). The concordance of activity status among structurally similar pairs was compared with a null in which activity labels were permuted among the participating compounds, across a range of similarity thresholds (Supplementary Figure 14E).

In the pairwise approach to identify phenotypic cliffs, the 100 nearest structural neighbors of each active compound were identified in Morgan fingerprint space using cosine distance on L2-normalized vectors. For each pair, structural similarity (cosine, approximating Tanimoto) and morphological similarity (cosine on L2-normalized CellProfiler profiles) were computed. A pair was classified as an activity cliff if its structural similarity was at least 0.7 and its morphological similarity was below 0.3, yielding 4,089 phenotypic cliffs among the 12,920 structurally similar pairs. The strength of the overall structure-morphology relationship was assessed by permutation: morphological similarities were shuffled 200 times to build a null distribution for the Spearman correlation (observed ρ = 0.111, roughly 100 standard deviations above the null).

To test whether phenotypic cliffs were enriched for pairs with different targets, annotated targets were compared within each source: a pair was counted as annotated when both compounds carried targets from the same source and as divergent when they shared no target in any such source, and phenotypic cliffs were compared with concordant pairs (morphological similarity ≥ 0.7) by a two-sided Fisher’s exact test.

### Predicting Activities from Cell Painting profiles

We evaluated whether Cell Painting profiles predict binary toxicity, continuous pharmacokinetic (PK), and external assay-activity endpoints, benchmarked against chemical-structure fingerprints alone. Across all tasks, compounds were matched to JUMP by the first 14 characters of the InChIKey (connectivity layer), and four feature sets were compared on an identical compound set: Cell Painting morphology profiles (758 features), Morgan fingerprints (ECFP4; 2,048-bit, radius 2), their concatenation (“Combined”), and a single median-cell-count feature. All models were histogram gradient-boosting (scikit-learn HistGradientBoostingClassifier/Regressor; max_iter = 100, max_depth = 6, learning_rate = 0.1; class-weight balanced for classification), evaluated by 5-fold cross-validation (ensuring each fold maintains the active/inactive distribution) with fold assignments shared across feature sets (random_state = 42). Two reference baselines were included: a label-shuffled stratified dummy classifier and the cell-count-only model above (a mean-predictor DummyRegressor for regression).

Drug-induced cardiotoxicity labels were taken from DICTrank ^36,103^ and drug-induced liver-injury labels from DILIrank(v2) ^104,105^, with compounds of “less” or “most” concern treated as positive and “no” concern as negative (ambiguous entries excluded); mitochondrial-toxicity labels were obtained from the MitoTox database (mitotox.org). After InChIKey matching and restriction to compounds with both profiles and fingerprints, 366 compounds were available for DICT, 356 for DILI, and 246 for MitoTox. Performance was reported as mean ± s.d. ROC-AUC across folds, together with the lift of the morphology model over the stratified baseline.

Five human PK parameters, steady-state volume of distribution (VDss), clearance (CL), fraction unbound in plasma (fup), mean residence time (MRT), and terminal half-life (t½), were obtained from a curated human-PK dataset ^106^ and matched to JUMP as above, yielding 396 (VDss), 404 (CL), 320 (fup), 392 (MRT), and 396 (t½) compounds. VDss, CL, MRT and t½ were modeled on a log scale and fup on its original scale; performance was reported as cross-validated coefficient of determination (R²) and Spearman rank correlation (ρ) between predicted and observed values, against a mean-predictor baseline.

We additionally screened per-endpoint binary activity across ToxCast assays (invitrodb v4.3 hit-calls) and ChEMBL single-protein targets. Each endpoint was retained only if, after intersecting with compounds having both a profile and a fingerprint, it contained ≥ 100 compounds with ≥ 20 active and ≥ 20 inactive, yielding 95 eligible ToxCast assays and 51 eligible ChEMBL targets. For each endpoint, the four feature sets were trained and evaluated by 5-fold stratified cross-validation as above, and results were summarized as survival curves showing, per feature set, the number of endpoints whose mean cross-validated ROC-AUC exceeds a given threshold.

### PAINS prediction

Each compound was labeled as PAINS-positive if its SMILES matched at least one substructure in the combined Baell PAINS-A/B/C catalogue as implemented in RDKit’s FilterCatalog ^25^; 5,032 / 112,524 compounds (4.5 %) were flagged (see Activity cliff detection for explanation of the set of compounds analyzed). We trained HistGradientBoosting classifiers to predict this label from (i) the 758 normalised Cell Painting morphology features, (ii) three physicochemical properties (MW, LogP, TPSA), and (iii) per-well median_cell_count alone, and compared ROC-AUC under 5-fold scaffold-split cross-validation. Note that the 5,021 PAINS compounds mentioned earlier in the text derives from the canonical analyzed set of 112,462 compounds.

### PPARG inverse agonists experiments

#### Cell Painting for PPARG plates

Cell Painting was performed as described ^85^ with minor modifications. UM-UC-9 cells (Sigma-Aldrich) were plated at 3,500 cells/well in either a pre-dispensed assay-ready plate (ARP) containing a panel of compounds from the Drug Repurposing Hub (https://clue.io/repurposing) or a blank 384-well Revvity, Inc. CellCarrier Ultra plate (ThermoFisher Scientific) in MEM-α media containing 10% fetal bovine serum (media as recommended by the supplier). UM-UC-9 cells in the blank plates were dosed with a custom set of vehicle, tool compounds, and PPARG inverse-agonist compounds using an HP-D300e Digital Dispenser (Tecan) and allowed to incubate at 37 °C with 5% CO2 and 90–95% humidity for 72 hours until reaching 80-90% confluence. All plates were stained and imaged as described ^85^.

#### PPARG plates Cell Painting image processing and feature extraction

The PPARG plates were imaged following the same protocol as for JUMP compounds at the Broad. Similarly, the same CellProfiler feature extraction pipeline and settings were used, with some modifications as captured in GitHub: https://github.com/broadinstitute/imaging-platform-pipelines/tree/master/cellpainting_UMUC9_20x_phenix_bin2. The features were then processed and aligned using the jump_profiling_recipe https://github.com/broadinstitute/jump-profiling-recipe/tree/main. The data from the PPARG plates were processed and aligned together with the JUMP compound data using the pipeline setting *profiles_var_mad_int_featselect_harmonyrsc,* see recipe repository description for details.

#### PPARG plates Pearson correlation

The processed profiles from the JUMP recipe were collapsed from replicate level down to consensus profiles, per cell line and compound-concentration, by averaging across replicates. The consensus profile for the highest dose for each of the compounds on the PPARG Cell Painting plates was selected on a per cell line basis. The Pearson correlation between each of the consensus profiles was then calculated using Morpheus (https://software.broadinstitute.org/morpheus). The resulting correlation matrix’s columns were filtered for the five *BAY-* compounds. The mean Pearson correlation between each compound and the five *BAY-* compounds was calculated and used to rank the rows of the matrix. This was done for the two cell lines UM-UC9 and U2OS separately.

#### PPARG plates phenotypic activity assessment

Following feature processing and alignment, the phenotypic activity of each compound-concentration pair was assessed using the *copairs* library ^27^. This calculates the similarity between replicates per compound-concentration pair compared to the negative control, measured using mAP. This strategy is used to identify compound-concentration pairs with distinct phenotypes compared to the negative control (DMSO). This is done within each of the cell lines in the PPARG plate data individually and similarly done for the JUMP compound data separately.

#### PPARG plates target association

Using annotations of each compound’s target from the Repurposing hub ^93^, we assessed the phenotypic consistency of phenotypically active compounds in the JUMP compound set using the *copairs* library ^27^. This evaluates how consistent each compound annotated as hitting a particular target is to others hitting the same target by comparing to those not having the same annotations, for targets with at least five compounds. Phenotypically consistent targets are defined as those with a positive nMAP score and corrected *p-value* less than 0.05 FDR. The analysis was done on compound-concentration pairs individually, where phenotypically active is defined as mAP score with a corrected p-value less than FDR = 0.05. A composite score for each target and compound was calculated based on the mean AP values across all phenotypically active concentrations for each compound. A cumulative score was calculated for each annotation group by summing the composite score of each compound in that group for each target separately.

#### JUMP CRISPR and ORF gene similarity

Features were extracted and processed from the JUMP ORF and CRISPR sets using the same exact settings as described above for PPARG plate data. After processing and alignment, consensus profiles were created by averaging across replicates of each *gene-reagent.* Phenotypic activity was calculated for each gene-reagent as described above for JUMP compounds. The processed data was filtered for gene symbols matching *CDK4, CDK6, PPARG (with any number or letter following)*. This resulted in 12 unique gene-reagents including both ORF and CRISPR samples. Cosine similarity was calculated between each of these 12 and all the unique gene-reagents in ORF and CRISPR. The similarity score was then converted to percentile ranking, individually for each of the 12 gene-reagents, resulting in a matrix of 12×23,393. The second dimension of this was then filtered down to each of these 12 gene-reagents, resulting in a 12×12 percentile matching ranking. Hierarchical clustering was applied using Morpheus https://github.com/cmap/morpheus.js. The default setting of *1-Pearson* correlation, Euclidean distance and average linkage was used.

##### Chemicals in PPARG experiments

1. UM-UC-9 Custom Compounds: BR00128250

a. DMSO (Sigma Cat# D5879-100, Lot SHBN0786)
b. LXH254 (RAF inhibitor) Selleck Chemicals Cat# S8745, lot # S874501
c. Palbociclib/PD-0332991 (Selleck Chemicals)
d. Rosiglitazone (Tocris)
e. GW1929 (Sigma Cat# G5668)
f. GW9662 (Tocris)
g. T0070907 (Tocris)
h. SR10221 (synthesized in house as described previously ^107^)
i. BAY-4931 (synthesized in house as described previously ^39,40^)
j. BAY-0069 (synthesized in house as described previously ^39,40^)
k. BAY-5516 (synthesized in house as described previously ^39,40^)
l. BAY-5094 (synthesized in house as described previously ^39,40^)
m. BAY-9683 (synthesized in house as described previously ^39,40^)
2. UM-UC-9 JUMP Target-2 ARP: BR00128252

a. See Supplementary File TARGET2

#### t-SNE (Morpheus)

t-SNE was performed using the Similarity Matrix tool in Morpheus (Broad Institute; https://software.broadinstitute.org/morpheus/), operating on columns with a distance metric of 1 minus the Pearson correlation, epsilon (learning rate) = 10, and perplexity = 15. The analysis was run on a filtered dataset (UMUC9_annotated_highconcentration_tSNE.gct) prepared as follows: (i) all features were normalized against DMSO negative controls; (ii) features with a high proportion of extreme outliers were removed; (iii) only the highest dose of each compound was retained (5 μM for ARP Target-2 compounds and 2.5 μM for custom compounds). Code for reproducing the t-SNE visualization is available in the R notebook PPARG_cellpainting_figures.Rmd.

#### Cell Cycle Analysis

UM-UC-9 cells were treated with the indicated compounds for 24 hours, trypsinized, spun at 300rcf for 5 min, and washed in PBS. Cells were incubated with LIVE/DEAD™ Fixable Near-IR Dead Cell Stain (#L34975 ThermoFisher Scientific), according to the manufacturer’s instructions. After washing in PBS, cells were fixed with ice-cold 70% ethanol in water and incubated for 30 minutes on ice. Fixed cells were washed in PBS twice and cell pellet was resuspended in propidium iodide (PI)/RNase staining buffer (#550825 BD Biosciences). Cells were acquired 15 minutes later using an Attune flow cytometer (ThermoFisher Scientific). Cell debris was excluded using an SSC-A vs FSC-A plot and viability dye-negative live cells were gated. Doublets were excluded by plotting PI-A vs PI-H and 20,000 single cell events were acquired. Data analysis was performed using FlowJo v10.8.1 software and cell cycle phases were determined using the FlowJo cell cycle analysis tool (Watson model).

#### Senescence β-gal Staining

UM-UC-9 cells were plated in 6-well Corning plates using MEM-α with 10% fetal bovine serum and treated with compounds at 500nM for a total of 8 days to induce senescence. Cells were passaged at 4 days based on normalization to vehicle cell density to maintain equivalent cell counts in each treatment condition. CellEventTM Senescence Green Detection Kit (ThermoFisher Scientific) was used to stain for β-galactosidase activity according to the manufacturer’s recommendation and plates were imaged using IncuCyte S3 Essen Bioscience/Sartorius.

#### RT-qPCR (reverse transcriptase quantitative polymerase chain reaction) for in vitro and ex vivo analyses of PPARG target gene expression

RNA was isolated using the RNeasy Kit (Qiagen) and first strand cDNA was synthesized using the LunaScript® RT SuperMix (New England Biolabs) according to the manufacturer’s recommended protocols. Quantitative PCR was performed on a QuantStudio 6 or 6 Pro, using TaqMan™ Universal PCR Master Mix (Applied Biosystems) and commercially available probes (Life Technologies). Gene expression probes were as follows:

| **Gene** | **DESCRIPTION** | **Probe ID** | **Reporter** |
|---|---|---|---|
| TNF | Human target gene | Hs00174128_m1 | FAM |
| CXCL10 | Human target gene | Hs00171042_m1 | FAM |
| CCL5 | Human target gene | Hs00982282_m1 | FAM |
| IL6 | Human target gene | Hs00985639_m1 | FAM |
| B2M | Human housekeeping gene | Hs00187842_m1 | VIC |
| PPIA | Human housekeeping gene | Hs99999904_m1 | FAM or VIC |

#### Western Blotting

UM-UC-9 cells were plated in 15cm dishes and treated for 8 days with compounds at 500nM for approximately 20 x 10^6^ cell density at final collection. Cells were passaged with seeding normalization to vehicle cell density after 4 days. Cells were collected in 1% SDS in PBS with protease and phosphatase inhibitors (Roche Applied Science) and lysed by sonication using a Branson Digital Sonifier Model 250/450 (Branson Ultrasonics Corporation) on a 30 seconds on/off cycle at 15% amplitude for 1 minute with samples kept on ice. Sample loading concentrations were normalized to lowest concentration. Western blots were performed using Novex 8-16% Tris-Glycine Plus Midi Gels (ThermoFisher Scientific) with semi-dry transfer using Trans-Blot SD (Bio-Rad Laboratories). Blots were probed with antibodies as indicated (see table for details) and imaged using the LI-COR Odyssey CLx Infrared Imaging System. For quantification of the detected signals of senescence markers, each lane of the specific senescence marker signal was normalized to the corresponding vinculin signal and subsequently normalized to DMSO vehicle via Image Studio software.

Antibodies for Western blot:

| Target | Vendor/Cat# | Dilution for Western blot |
|---|---|---|
| p-Rb (S780) | Cell Signal 8180T | 1:1000 |
| Rb | Cell Signal 9309S | 1:1000 |
| Cyclin D1 | Cell Signal 2978S | 1:1000 |
| Beta-actin | Millipore-Sigma | 1:1000 |
| Goat Anti-Mouse IgG Antibody, IRDye; 680LT Conjugated | LI-COR Biosciences 926-68020 | 1:10,000 |
| Goat Anti-Rabbit IgG Antibody, IRDye; 800CW | LI-COR Biosciences 926-32211 | 1:10,000 |

## Acknowledgments

The authors appreciate the scientists and administrative personnel who contributed to the organization, scientific direction, and execution of the JUMP Cell Painting Consortium’s project. In particular, we thank Justyna Różycka, Barbara Ściborowska, and Bartosz Szabucki, for work on the development of PhenAID JUMP-CP Data Explorer application, Piotr Gaiński and Michał Koziarski for their efforts in preprocessing and analysis of the data presented in the application, the Pistoia Alliance for coordinating assembly of the final shared compound library, and Niveditha Iyer, Daniel Michael, S-Y Gordon Chi, and Pranav Rajpurkar for foundational work that revealed fundamental challenges of image-based gene-compound matching.

## Author contributions

**Conceptualization**: J.D.B., H.C., D.A.C., T.D., R.D., O.E., P.W.F., S.G., S.H., D.K., F.M., A.-L.O., C.P., J.P., P.S., S.E.S., S.S.₄, A.E.C.; **Data curation**: S.N.C., S.S.₁, D.A.A., J.A., N.B., P.J.B., B.A.C., D.A.C., V.D., J.F.H., J.G.₁, J.G.₂, Y.H., Z.H., S.H., G.H., I.I., C.J., S.J., M.A., D.K., C.H.L., F.L., K.R.L., D.J.L., S.L., Y.L., P.A.M.Z., L.M.₁, M.O.₁, C.Q., S.K.S., I.S., H.T., G.J.P.v.W., E.W., N.W.D., G.W., S.Y., B.Z., S.S.₄; **Analysis**: S.N.C., S.S.₁, L.A., J.A., T.D., J.F.H., J.T.G., J.G.₁, J.G.₂, Y.H., Z.H., S.H., J.A.H., I.I., S.J., A.A.K., D.K., F.L., G.L., D.J.L., P.A.M.Z., L.M.₁, H.M., A.F.M., A.O., M.O.₁, C.P., R.A.S., R.S., D.S., H.T., M.V.D., E.W., G.W., B.Z., I.Z., M.Z., S.S.₄, A.E.C.; **Funding acquisition**: J.D.B., H.C., D.A.C., R.D., O.E., S.G., D.G.₁, N.J.M., P.J.P., S.S.₄, A.E.C.; **Experiments/Investigation**: S.N.C., S.S.₁, E.A., D.M.A., M.B., N.B., J.D.B., P.J.B., V.D., J.F.H., B.F., F.F., S.G., T.G., D.G.₂, J.T.G., A.G., S.H., D.H., G.H., M.I., C.J., I.J.F., J.K., M.A., D.K., C.H.L., K.D.L., K.R.L., D.J.L., Y.L., B.A.M., G.M., N.J.M., L.M.₁, L.M.₂, T.M., E.M., B.N., A.-L.O., C.P., S.P., S.S.₂, H.S., I.S., S.T., A.T., J.V.d.W., C.v.S., E.W., A.W., G.W., S.Y., B.Z., I.Z., S.S.₄; **Methodology**: S.N.C., S.S.₁, A.B., H.C., B.A.C., T.D., R.D., J.F.H., B.F., S.G., T.G., A.G., J.G.₁, S.H., J.A.H., S.R.H., M.I., S.J., A.A.K., M.A., D.K., C.H.L., A.L., D.J.L., F.M., B.A.M., G.M., L.M.₁, L.M.₂, E.M., J.P., D.E.R., G.W., B.Z., I.Z., M.Z., S.S.₄, A.E.C.; **Project administration**: S.N.C., J.L.A., B.A.C., J.G.D., R.D., O.E., P.W.F., B.F., S.G., A.G., S.R.H., C.J., S.J., M.A., D.K., C.H.L., F.M., P.A.M.Z., B.A.M., L.M.₂, E.M., A.O., A.-L.O., D.E.R., S.S.₂, S.E.S., M.W., S.S.₄, A.E.C.; **Resources**: E.A., N.B., C.C., D.A.C., J.G.D., R.D., V.D., P.W.F., B.F., S.G., T.G., D.G.₂, A.G., S.H., D.H., S.R.H., G.H., M.H., I.I., M.I., M.A., C.H.L., A.L., D.J.L., P.A.M.Z., B.A.M., G.M., H.M., E.M., A.-L.O., M.O.₁, S.P., D.E.R., S.K.S., S.S.₂, H.S., C.S., I.S., H.T., M.V.D., B.Z.; **Software**: S.N.C., S.S.₁, D.A.A., D.M.A., J.A., N.B., B.A.C., N.A.D., J.F.H., Y.H., A.A.K., T.K., F.L., G.L., D.J.L., L.M.₁, A.F.M., M.O.₁, C.Q., S.S.₃, H.T., A.U., M.V.D., E.W., N.W.D., G.W., B.Z., S.S.₄; **Supervision**: S.N.C., M.B., A.B., L.B., H.C., B.A.C., D.A.C., J.G.D., T.D., R.D., O.E., P.W.F., B.F., S.G., D.G.₂, A.G., S.R.H., C.J., S.J., M.A., T.K., D.K., C.H.L., D.J.L., F.M., P.A.M.Z., N.J.M., H.M., E.M., A.O., A.-L.O., M.O.₂, D.E.R., H.S., P.S., I.S., G.J.P.v.W., M.W., S.S.₄, A.E.C.; **Validation**: S.N.C., S.S.₁, J.A., P.J.B., J.F.H., Y.H., Z.H., I.J.F., J.K., C.H.L., K.R.L., D.J.L., B.A.M., L.M.₁, L.M.₂, C.Q., S.S.₃, A.T., J.V.d.W., A.W., N.W.D., G.W., B.Z., S.S.₄; **Visualization**: S.N.C., S.S.₁, J.F.H., K.R., R.A.S., D.S., S.S.₄, A.E.C.; **Writing – original draft**: S.N.C., S.S.₁, D.A.A., A.B., B.A.C., B.F., A.G., L.M.₂, M.O.₂, C.P., S.S.₄, A.E.C.; **Writing – review & editing**: S.N.C., S.S.₁, B.A.C., S.G., J.T.G., J.A.H., S.R.H., I.J.F., M.A., D.J.L., M.O.₂, S.T., M.W., E.W., S.S.₄, A.E.C.

## Declaration of generative AI and AI-assisted technologies in the manuscript preparation process

During the preparation of this work, some authors used Claude Code and Codex for data analysis, and various LLMs to inspect claims in the text, provide critical feedback, and to identify papers relevant to the work. The authors reviewed and edited the content and take full responsibility for the content of the published article.

## References

1. Cimini, B.A., Chandrasekaran, S.N., Kost-Alimova, M., Miller, L., Goodale, A., Fritchman, B., Byrne, P., Garg, S., Jamali, N., Logan, D.J., et al. (2023). Optimizing the Cell Painting assay for image-based profiling. Nat. Protoc. 18, 1981–2013.

2. Seal, S., Trapotsi, M.-A., Spjuth, O., Singh, S., Carreras-Puigvert, J., Greene, N., Bender, A., and Carpenter, A.E. (2025). Cell Painting: a decade of discovery and innovation in cellular imaging. Nat. Methods 22, 254–268.

3. Dagher, M., Ongo, G., Robichaud, N., Kong, J., Rho, W., Teahulos, I., Tavakoli, A., Bovaird, S., Merjaneh, S., Tan, A., et al. (2025). nELISA: a high-throughput, high-plex platform enables quantitative profiling of the inflammatory secretome. Nat. Methods 22, 2375–2385.

4. Haghighi, M., Caicedo, J.C., Cimini, B.A., Carpenter, A.E., and Singh, S. (2022). High-dimensional gene expression and morphology profiles of cells across 28,000 genetic and chemical perturbations. Nat. Methods 19, 1550–1557.

5. Way, G.P., Natoli, T., Adeboye, A., Litichevskiy, L., Yang, A., Lu, X., Caicedo, J.C., Cimini, B.A., Karhohs, K., Logan, D.J., et al. (2022). Morphology and gene expression profiling provide complementary information for mapping cell state. Cell Syst. 13, 911–923.e9.

6. Perlman, Z.E., Slack, M.D., Feng, Y., Mitchison, T.J., Wu, L.F., and Altschuler, S.J. (2004). Multidimensional drug profiling by automated microscopy. Science 306, 1194–1198.

7. Rohban, M.H., Fuller, A.M., Tan, C., Goldstein, J.T., Syangtan, D., Gutnick, A., DeVine, A., Nijsure, M.P., Rigby, M., Sacher, J.R., et al. (2022). Virtual screening for small-molecule pathway regulators by image-profile matching. Cell Syst. 13, 724–736.e9.

8. Hofmarcher, M., Rumetshofer, E., Clevert, D.-A., Hochreiter, S., and Klambauer, G. (2019). Accurate Prediction of Biological Assays with High-Throughput Microscopy Images and Convolutional Networks. J. Chem. Inf. Model. 59, 1163–1171.

9. Moshkov, N., Becker, T., Yang, K., Horvath, P., Dancik, V., Wagner, B.K., Clemons, P.A., Singh, S., Carpenter, A.E., and Caicedo, J.C. (2023). Predicting compound activity from phenotypic profiles and chemical structures. Nat. Commun. 14. 10.1038/s41467-023-37570-1.

10. Cuccarese, M.F., Earnshaw, B.A., Heiser, K., Fogelson, B., Davis, C.T., McLean, P.F., Gordon, H.B., Skelly, K.-R., Weathersby, F.L., Rodic, V., et al. (2020). Functional immune mapping with deep-learning enabled phenomics applied to immunomodulatory and COVID-19 drug discovery. bioRxiv, 2020.08.02.233064. 10.1101/2020.08.02.233064.

11. Moshkov, N., Bornholdt, M., Benoit, S., Smith, M., McQuin, C., Goodman, A., Senft, R.A., Han, Y., Babadi, M., Horvath, P., et al. (2024). Learning representations for image-based profiling of perturbations. Nat. Commun. 15, 1594.

12. Trapotsi, M.-A., Mouchet, E., Williams, G., Monteverde, T., Juhani, K., Turkki, R., Miljković, F., Martinsson, A., Mervin, L., Pryde, K.R., et al. (2022). Cell morphological profiling enables high-throughput screening for PROteolysis TArgeting chimera (PROTAC) phenotypic signature. ACS Chem. Biol. 17, 1733–1744.

13. Seal, S., Yang, H., Trapotsi, M.-A., Singh, S., Carreras-Puigvert, J., Spjuth, O., and Bender, A. (2023). Merging bioactivity predictions from cell morphology and chemical fingerprint models using similarity to training data. J. Cheminform. 15, 56.

14. Nyffeler, J., Willis, C., Lougee, R., Richard, A., Paul-Friedman, K., and Harrill, J.A. (2020). Bioactivity screening of environmental chemicals using imaging-based high-throughput phenotypic profiling. Toxicol. Appl. Pharmacol. 389, 114876.

15. Cross-Zamirski, J.O., Williams, G., Mouchet, E., Schönlieb, C.-B., Turkki, R., and Wang, Y. (2022). Self-supervised learning of phenotypic representations from cell images with weak labels. arXiv [cs.CV].

16. Williams, E., Moore, J., Li, S.W., Rustici, G., Tarkowska, A., Chessel, A., Leo, S., Antal, B., Ferguson, R.K., Sarkans, U., et al. (2017). The Image Data Resource: A Bioimage Data Integration and Publication Platform. Nat. Methods 14, 775–781.

17. Ljosa, V., Sokolnicki, K.L., and Carpenter, A.E. (2012). Annotated high-throughput microscopy image sets for validation. Nat. Methods 9, 637.

18. Weisbart, E., Kumar, A., Arevalo, J., Carpenter, A.E., Cimini, B.A., and Singh, S. (2024). Cell Painting Gallery: an open resource for image-based profiling. Nat. Methods. 10.1038/s41592-024-02399-z.

19. Hartley, M., Kleywegt, G.J., Patwardhan, A., Sarkans, U., Swedlow, J.R., and Brazma, A. (2022). The BioImage Archive - building a home for life-sciences microscopy data. J. Mol. Biol. 434, 167505.

20. Bray, M.-A., Gustafsdottir, S.M., Rohban, M.H., Singh, S., Ljosa, V., Sokolnicki, K.L., Bittker, J.A., Bodycombe, N.E., Dancík, V., Hasaka, T.P., et al. (2017). A dataset of images and morphological profiles of 30 000 small-molecule treatments using the Cell Painting assay. Gigascience 6, 1–5.

21. Chandrasekaran, S.N., Alix, E., Arevalo, J., Borowa, A., Byrne, P.J., Charles, W.G., Chen, Z.S., Cimini, B.A., Deng, B., Doench, J.G., et al. (2025). Morphological map of under- and overexpression of genes in human cells. Nat. Methods 22, 1742–1752.

22. Chandrasekaran, S.N., Cimini, B.A., Goodale, A., Miller, L., Kost-Alimova, M., Jamali, N., Doench, J.G., Fritchman, B., Skepner, A., Melanson, M., et al. (2024). Three million images and morphological profiles of cells treated with matched chemical and genetic perturbations. Nat. Methods 21, 1114–1121.

23. Tromans-Coia, C., Jamali, N., Abbasi, H.S., Giuliano, K.A., Hagimoto, M., Jan, K., Kaneko, E., Letzsch, S., Schreiner, A., Sexton, J.Z., et al. (2023). Assessing the performance of the Cell Painting assay across different imaging systems. Cytometry A 103, 915–926.

24. Bickerton, G.R., Paolini, G.V., Besnard, J., Muresan, S., and Hopkins, A.L. (2012). Quantifying the chemical beauty of drugs. Nat. Chem. 4, 90–98.

25. Baell, J.B., and Holloway, G.A. (2010). New substructure filters for removal of pan assay interference compounds (PAINS) from screening libraries and for their exclusion in bioassays. J. Med. Chem. 53, 2719–2740.

26. Phillips, K.A., Wambaugh, J.F., Grulke, C.M., Dionisio, K.L., and Isaacs, K.K. (2017). High-throughput screening of chemicals as functional substitutes using structure-based classification models. Green Chem. 19, 1063–1074.

27. Kalinin, A.A., Arevalo, J., Serrano, E., Vulliard, L., Tsang, H., Bornholdt, M., Muñoz, A.F., Sivagurunathan, S., Rajwa, B., Carpenter, A.E., et al. (2025). A versatile information retrieval framework for evaluating profile strength and similarity. Nat Commun 16, 5181.

28. Stirling, D.R., Swain-Bowden, M.J., Lucas, A.M., Carpenter, A.E., Cimini, B.A., and Goodman, A. (2021). CellProfiler 4: improvements in speed, utility and usability. BMC Bioinformatics 22, 433.

29. Korsunsky, I., Millard, N., Fan, J., Slowikowski, K., Zhang, F., Wei, K., Baglaenko, Y., Brenner, M., Loh, P.-R., and Raychaudhuri, S. (2019). Fast, sensitive and accurate integration of single-cell data with Harmony. Nat. Methods 16, 1289–1296.

30. Wawer, M.J., Li, K., Gustafsdottir, S.M., Ljosa, V., Bodycombe, N.E., Marton, M.A., Sokolnicki, K.L., Bray, M.-A., Kemp, M.M., Winchester, E., et al. (2014). Toward performance-diverse small-molecule libraries for cell-based phenotypic screening using multiplexed high-dimensional profiling. Proc. Natl. Acad. Sci. U. S. A. 111, 10911–10916.

31. Lipinski, C.A., Lombardo, F., Dominy, B.W., and Feeney, P.J. (1997). Experimental and computational approaches to estimate solubility and permeability in drug discovery and development settings. Adv. Drug Deliv. Rev. 23, 3–25.

32. Pahl, A., Schölermann, B., Lampe, P., Rusch, M., Dow, M., Hedberg, C., Nelson, A., Sievers, S., Waldmann, H., and Ziegler, S. (2023). Morphological subprofile analysis for bioactivity annotation of small molecules. Cell Chem. Biol. 30, 839–853.e7.

33. Chow, Y.L., Singh, S., Carpenter, A.E., and Way, G.P. (2022). Predicting drug polypharmacology from cell morphology readouts using variational autoencoder latent space arithmetic. PLoS Comput. Biol. 18, e1009888.

34. Sanchez, M., Bourriez, N., Bendidi, I., Cohen, E., Svatko, I., Del Nery, E., Tajmouati, H., Bollot, G., Calzone, L., and Genovesio, A. (2026). Large scale compound selection guided by cell painting reveals activity cliffs and functional relationships. Commun. Biol. 9, 225.

35. Dahlin, J.L., Hua, B.K., Zucconi, B.E., Nelson, S.D., Jr, Singh, S., Carpenter, A.E., Shrimp, J.H., Lima-Fernandes, E., Wawer, M.J., Chung, L.P.W., et al. (2023). Reference compounds for characterizing cellular injury in high-content cellular morphology assays. Nat. Commun. 14, 1364.

36. Seal, S., Spjuth, O., Hosseini-Gerami, L., García-Ortegón, M., Singh, S., Bender, A., and Carpenter, A.E. (2024). Insights into Drug Cardiotoxicity from Biological and Chemical Data: The First Public Classifiers for FDA Drug-Induced Cardiotoxicity Rank. J. Chem. Inf. Model. 64, 1172–1186.

37. Rouquié, D., Bender, A., Cheah, J., Crute, C.E., Dalmas, D., Ewald, J., Fullerton, A., Harrill, J.A., Kadri, S., Kleinstreuer, N., et al. (2025). The OASIS Consortium: integrating multi-omics technologies to transform chemical safety assessment. Toxicological Sciences. 10.1093/toxsci/kfaf128.

38. Seal, S., Carreras-Puigvert, J., Trapotsi, M.-A., Yang, H., Spjuth, O., and Bender, A. (2022). Integrating cell morphology with gene expression and chemical structure to aid mitochondrial toxicity detection. Commun. Biol. 5, 858.

39. Orsi, D.L., Pook, E., Bräuer, N., Friberg, A., Lienau, P., Lemke, C.T., Stellfeld, T., Brüggemeier, U., Pütter, V., Meyer, H., et al. (2022). Discovery and structure-based design of potent covalent PPARγ inverse-agonists BAY-4931 and BAY-0069. J. Med. Chem. 65, 14843–14863.

40. Orsi, D.L., Ferrara, S.J., Siegel, S., Friberg, A., Bouché, L., Pook, E., Lienau, P., Bluck, J.P., Lemke, C.T., Akcay, G., et al. (2023). Discovery and characterization of orally bioavailable 4-chloro-6-fluoroisophthalamides as covalent PPARG inverse-agonists. Bioorg. Med. Chem. 78, 117130.

41. Goldstein, J.T., Berger, A.C., Shih, J., Duke, F.F., Furst, L., Kwiatkowski, D.J., Cherniack, A.D., Meyerson, M., and Strathdee, C.A. (2017). Genomic activation of PPARG reveals a candidate therapeutic axis in bladder cancer. Cancer Res. 77, 6987–6998.

42. Jin, M., Li, J., Hu, R., Xu, B., Huang, G., Huang, W., Chen, B., He, J., and Cao, Y. (2021). Cyclin A2/cyclin-dependent kinase 1-dependent phosphorylation of Top2a is required for S phase entry during retinal development in zebrafish. J. Genet. Genomics 48, 63–74.

43. Ibrahim, L., Mohamed, R.H., Tolba, M.M., Radwan, S.M., Hamdy, N.M., and Elhefnawi, M. (2025). Unveiling the therapeutic potential of targeting RRM2 in hepatocellular carcinoma: An integrated in silico and in vitro study. Funct. Integr. Genomics 25, 123.

44. Nguyen, T.B., Do, D.N., Nguyen-Thanh, T., Tatipamula, V.B., and Nguyen, H.T. (2021). Identification of five hub genes as key prognostic biomarkers in liver cancer via integrated bioinformatics analysis. Biology (Basel) 10, 957.

45. Arevalo, J., Su, E., Carpenter, A.E., and Singh, S. (2024). MOTIVE: A drug-target interaction graph for inductive link prediction. Proceedings of The Thirty-Eighth Annual Conference on Neural Information Processing Systems.

46. Finn, R.S., Dering, J., Conklin, D., Kalous, O., Cohen, D.J., Desai, A.J., Ginther, C., Atefi, M., Chen, I., Fowst, C., et al. (2009). PD 0332991, a selective cyclin D kinase 4/6 inhibitor, preferentially inhibits proliferation of luminal estrogen receptor-positive human breast cancer cell lines in vitro. Breast Cancer Res. 11, R77.

47. Jost, T., Heinzerling, L., Fietkau, R., Hecht, M., and Distel, L.V. (2021). Palbociclib induces senescence in melanoma and breast cancer cells and leads to additive growth arrest in combination with irradiation. Front. Oncol. 11, 740002.

48. Dee, W., Sequeira, I., Lobley, A., and Slabaugh, G. (2024). Cell-vision fusion: A Swin transformer-based approach for predicting kinase inhibitor mechanism of action from Cell Painting data. iScience 27, 110511.

49. Nguyen, C.Q., Pertusi, D., and Branson, K.M. (2023). Molecule-Morphology Contrastive Pretraining for transferable molecular representation. arXiv [q-bio.BM]. 10.1101/2023.05.01.538999.

50. Wang, X., Fan, Y., Guo, Y., Fu, C., Lee, K., Dallakyan, K., Li, Y., Yin, Q., Li, Y., and Song, L. (2025). Prediction of cellular morphology changes under perturbations with a transcriptome-guided diffusion model. Nat. Commun. 16, 8210.

51. Paysan, D., Radhakrishnan, A., Zhang, X., Shivashankar, G.V., and Uhler, C. (2025). Image2Reg: Linking chromatin images to gene regulation using genetic and chemical perturbation screens. Cell Syst. 16, 101293.

52. Lazar, N.H., Celik, S., Chen, L., Fay, M.M., Irish, J.C., Jensen, J., Tillinghast, C.A., Urbanik, J., Bone, W.P., Gibson, C.C., et al. (2024). High-resolution genome-wide mapping of chromosome-arm-scale truncations induced by CRISPR-Cas9 editing. Nat. Genet. 56, 1482–1493.

53. Tegtmeyer, M., Arora, J., Asgari, S., Cimini, B.A., Nadig, A., Peirent, E., Liyanage, D., Way, G.P., Weisbart, E., Nathan, A., et al. (2024). High-dimensional phenotyping to define the genetic basis of cellular morphology. Nat. Commun. 15, 347.

54. Wu, A.P., Yao, H., Hoeckendorf, B., Gaskins, G., Kosaisawe, N., Liu, Z., Hanslovsky, P., Mayba, O., Skelton, N., Scalia, G., et al. (2026). Virtual phenotypic screening discovers novel scaffolds inhibiting the PI3K/mTOR pathway. bioRxiv. 10.64898/2026.06.10.731476.

55. Arevalo, J., Su, E., Ewald, J.D., van Dijk, R., Carpenter, A.E., and Singh, S. (2024). Evaluating batch correction methods for image-based cell profiling. Nat. Commun. 15, 6516.

56. Yan, C., Zhang, Y., Feng, J., Hua, H., Ruan, Z., Li, Z., Li, S., Yan, C., Li, P., Liu, J., et al. (2025). Triple-effect correction for Cell Painting data with contrastive and domain-adversarial learning. Nat. Commun. 16, 6886.

57. Caruzzo, C., and Ye, J.C. (2025). CellPainTR: Generalizable representation learning for cross-dataset Cell Painting analysis. arXiv [cs.CV]. 10.48550/arXiv.2509.06986.

58. Tomkinson, J., Kern, R., Mattson, C., and Way, G.P. (2024). Toward generalizable phenotype prediction from single-cell morphology representations. BMC Methods 1. 10.1186/s44330-024-00014-3.

59. Li, B., Wang, Q., Wang, S., Zhang, B., Peng, Y., Zeng, P., Liu, C., Li, M., Tang, Z., Yao, X., et al. (2026). MVCBench: A multimodal benchmark for drug-induced virtual cell phenotypes. bioRxiv, 2026.04.22.720110. 10.64898/2026.04.22.720110.

60. Peng, R., Liu, Z., Gao, Y., and Wang, J. (2026). A generative framework for predicting cellular morphological and transcriptomic perturbation responses. Cell Rep. Methods 6, 101459.

61. Yu, Y., Hayir, E., Tenenholtz, N., Mackey, L., Wei, Y., Alvarez-Melis, D., Amini, A.P., and Lu, A.X. (2026). Integrating chemical structures as treatments improves representations of microscopy images for morphological profiling. PLoS Comput. Biol. 22, e1014483.

62. Kim, V., Adaloglou, N., Osterland, M., Morelli, F.M., Halawa, M., König, T., Gnutt, D., and Marin Zapata, P.A. (2025). Self-supervision advances morphological profiling by unlocking powerful image representations. Sci. Rep. 15, 4876.

63. Moutakanni, T., Couprie, C., Yi, S., Doron, M., Chen, Z.S., Moshkov, N., Gardes, E., Caron, M., Touvron, H., Joulin, A., et al. (2025). Cell-DINO: Self-supervised image-based embeddings for cell fluorescent microscopy. PLoS Comput. Biol. 21, e1013828.

64. Kraus, O., Kenyon-Dean, K., Saberian, S., Fallah, M., McLean, P., Leung, J., Sharma, V., Khan, A., Balakrishnan, J., Celik, S., et al. (2024). Masked autoencoders for microscopy are scalable learners of cellular biology. In 2024 IEEE/CVF Conference on Computer Vision and Pattern Recognition (CVPR) (IEEE), pp. 11757–11768.

65. Bilodeau, A., Beaupré, F., Chabbert, J., Thériault, K., Deschênes, A., Bellavance, J.-M., Lessard, K., Bernatchez, R., De Koninck, P., Gagné, C., et al. (2025). A self-supervised foundation model for robust and generalizable representation learning in STED microscopy. bioRxiv. 10.1101/2025.06.06.656993.

66. Hayir, E., Crawford, L., and Lu, A.X. (2026). MorphoHELM: A comprehensive benchmark for evaluating representations for microscopy-based morphology assays. arXiv [cs.CV]. 10.48550/arXiv.2605.15383.

67. Agrawal, V., Peters, J., Thompson, T.N., Sanian, M.V., Pham, C., Moshkov, N., Kazi, A., Pillai, A., Freeman, J., Kang, B., et al. (2025). CHAMMI-75: pre-training multi-channel models with heterogeneous microscopy images. arXiv [cs.CV]. 10.48550/arXiv.2512.20833.

68. Watkinson, G., Cohen, E., Bourriez, N., Bendidi, I., Bollot, G., and Genovesio, A. (2024). Weakly supervised cross-modal learning in high-content screening. In 2024 IEEE International Symposium on Biomedical Imaging (ISBI) (IEEE), pp. 1–5.

69. Borowa, A., Rymarczyk, D., Żyła, M., Kańdula, M., Sánchez-Fernández, A., Rataj, K., Struski, Ł., Tabor, J., and Zieliński, B. (2024). Decoding phenotypic screening: A comparative analysis of image representations. Comput. Struct. Biotechnol. J. 23, 1181–1188.

70. Yun, S., Yao, H., Hoeckendorf, B., Richmond, D., Regev, A., and Littman, R. (2025). Sparse Mixture-of-experts for multi-channel imaging: Are all channel interactions required? arXiv [cs.CV]. 10.48550/arXiv.2511.17400.

71. Kohnstamm, S. (2026). Claude Science Meets the Quilt MCP: From Public Cell Painting Data to 25 Tested Hypotheses. https://blog.quilt.bio/claude-science-quilt-mcp-cell-painting.

72. Rohban, M.H., Abbasi, H.S., Singh, S., and Carpenter, A.E. (2019). Capturing single-cell heterogeneity via data fusion improves image-based profiling. Nat. Commun. 10, 2082.

73. van Dijk, R., Arevalo, J., Babadi, M., Carpenter, A.E., and Singh, S. (2024). Capturing cell heterogeneity in representations of cell populations for image-based profiling using contrastive learning. PLoS Comput. Biol. 20, e1012547.

74. Shpigler, A., Kolet, N., Golan, S., Weisbart, E., and Zaritsky, A. (2025). Anomaly detection for high-content image-based phenotypic cell profiling. Cell Syst. 16, 101429.

75. Kreiss, L., Jiang, S., Li, X., Xu, S., Zhou, K.C., Lee, K.C., Mühlberg, A., Kim, K., Chaware, A., Ando, M., et al. (2023). Digital staining in optical microscopy using deep learning - a review. Photonix 4. 10.1186/s43074-023-00113-4.

76. Mehrizi, R., Mehrjou, A., Alegro, M., Zhao, Y., Carbone, B., Fishwick, C., Vappiani, J., Bi, J., Sanford, S., Keles, H., et al. (2023). Multi-omics prediction from high-content cellular imaging with deep learning. arXiv [q-bio.QM]. 10.48550/arXiv.2306.09391.

77. Chlis, N.-K., Rausch, L., Brocker, T., Kranich, J., and Theis, F.J. (2020). Predicting single-cell gene expression profiles of imaging flow cytometry data with machine learning. Nucleic Acids Res. 48, 11335–11346.

78. Yang, K.D., Belyaeva, A., Venkatachalapathy, S., Damodaran, K., Katcoff, A., Radhakrishnan, A., Shivashankar, G.V., and Uhler, C. (2021). Multi-domain translation between single-cell imaging and sequencing data using autoencoders. Nat. Commun. 12, 31.

79. Naidoo, R., Hu, J., Tripodi, G., Bakal, C., and Chakraborti, T. (2026). Cell Painting Generates Single-Cell Transcriptomics via Conditional Diffusion.

80. Lejal, V., Rouquié, D., and Taboureau, O. (2025). Cell morphology and gene expression: tracking changes and complementarity across time and cell lines. Toxicol. Appl. Pharmacol. 504, 117530.

81. Ha, S.V., Jaensch, S., Kańduła, M.M., Herman, D., Czodrowski, P., and Ceulemans, H. (2025). Cross modality learning of cell painting and transcriptomics data improves mechanism of action clustering and bioactivity modelling. Sci. Rep. 15, 23010.

82. Muñoz, A.F. (2026). JUMP production paper: jump_rr-compatible compound tables. (Zenodo). 10.5281/ZENODO.21515641.

83. Muñoz, A.F. (2026). Phenotypically active ORF, CRISPR and compound consensus profiles. (Zenodo). 10.5281/ZENODO.20142386.

84. Tingle, B.I., Tang, K.G., Castanon, M., Gutierrez, J.J., Khurelbaatar, M., Dandarchuluun, C., Moroz, Y.S., and Irwin, J.J. (2023). ZINC-22─A free multi-billion-scale database of tangible compounds for ligand discovery. J. Chem. Inf. Model. 63, 1166–1176.

85. Bray, M.-A., Singh, S., Han, H., Davis, C.T., Borgeson, B., Hartland, C., Kost-Alimova, M., Gustafsdottir, S.M., Gibson, C.C., and Carpenter, A.E. (2016). Cell Painting, a high-content image-based assay for morphological profiling using multiplexed fluorescent dyes. Nat. Protoc. 11, 1757–1774.

86. McQuin, C., Goodman, A., Chernyshev, V., Kamentsky, L., Cimini, B.A., Karhohs, K.W., Doan, M., Ding, L., Rafelski, S.M., Thirstrup, D., et al. (2018). CellProfiler 3.0: Next-generation image processing for biology. PLoS Biol. 16, e2005970.

87. Serrano, E., Chandrasekaran, S.N., Bunten, D., Brewer, K.I., Tomkinson, J., Kern, R., Bornholdt, M., Fleming, S.J., Pei, R., Arevalo, J., et al. (2025). Reproducible image-based profiling with Pycytominer. Nat. Methods. 10.1038/s41592-025-02611-8.

88. Bemis, G.W., and Murcko, M.A. (1996). The properties of known drugs. 1. Molecular frameworks. J. Med. Chem. 39, 2887–2893.

89. McInnes, L., Healy, J., and Melville, J. (2018). UMAP: Uniform Manifold Approximation and Projection for Dimension Reduction. arXiv [stat.ML].

90. Dicks, S., Heumos, L., May, L., Jimenez, S., Angerer, P., Gold, I., Virshup, I., Fischer, F., Gill, M., Boerries, M., et al. (2026). GPU-accelerated single-cell analysis at scale with rapids-singlecell. arXiv [q-bio.GN]. 10.48550/arXiv.2603.02402.

91. Wolf, F.A., Angerer, P., and Theis, F.J. (2018). SCANPY: large-scale single-cell gene expression data analysis. Genome Biol. 19, 15.

92. Benjamini, Y., and Hochberg, Y. (1995). Controlling the false discovery rate: A practical and powerful approach to multiple testing. J. R. Stat. Soc. Series B Stat. Methodol. 57, 289–300.

93. Corsello, S.M., Bittker, J.A., Liu, Z., Gould, J., McCarren, P., Hirschman, J.E., Johnston, S.E., Vrcic, A., Wong, B., Khan, M., et al. (2017). The Drug Repurposing Hub: a next-generation drug library and information resource. Nat. Med. 23, 405–408.

94. Zdrazil, B., Felix, E., Hunter, F., Manners, E.J., Blackshaw, J., Corbett, S., de Veij, M., Ioannidis, H., Lopez, D.M., Mosquera, J.F., et al. (2024). The ChEMBL Database in 2023: a drug discovery platform spanning multiple bioactivity data types and time periods. Nucleic Acids Res. 52, D1180–D1192.

95. Sanfelice, D., Antolin, A.A., Crisp, A., Chen, Y., Bellenie, B., Brennan, P.E., Edwards, A., Müller, S., Al-Lazikani, B., and Workman, P. (2025). The Chemical Probes Portal - 2024: update on this public resource to support best-practice selection and use of small molecules in biomedical research. Nucleic Acids Res. 53, D1663–D1669.

96. Buniello, A., Suveges, D., Cruz-Castillo, C., Llinares, M.B., Cornu, H., Lopez, I., Tsukanov, K., Roldán-Romero, J.M., Mehta, C., Fumis, L., et al. (2025). Open Targets Platform: facilitating therapeutic hypotheses building in drug discovery. Nucleic Acids Res. 53, D1467–D1475.

97. Chandak, P., Huang, K., and Zitnik, M. (2023). Building a knowledge graph to enable precision medicine. Sci. Data 10, 67.

98. Judson, R.S., Houck, K.A., Kavlock, R.J., Knudsen, T.B., Martin, M.T., Mortensen, H.M., Reif, D.M., Rotroff, D.M., Shah, I., Richard, A.M., et al. (2010). In vitro screening of environmental chemicals for targeted testing prioritization: the ToxCast project. Environ. Health Perspect. 118, 485–492.

99. Tamir, T.Y., Drewry, D.H., Wells, C., Major, M.B., and Axtman, A.D. (2020). PKIS deep dive yields a chemical starting point for dark kinases and a cell active BRSK2 inhibitor. Sci. Rep. 10, 15826.

100. Luecken, M.D., Büttner, M., Chaichoompu, K., Danese, A., Interlandi, M., Mueller, M.F., Strobl, D.C., Zappia, L., Dugas, M., Colomé-Tatché, M., et al. (2022). Benchmarking atlas-level data integration in single-cell genomics. Nat. Methods 19, 41–50.

101. Büttner, M., Miao, Z., Wolf, F.A., Teichmann, S.A., and Theis, F.J. (2019). A test metric for assessing single-cell RNA-seq batch correction. Nat. Methods 16, 43–49.

102. Rogers, D., and Hahn, M. (2010). Extended-connectivity fingerprints. J. Chem. Inf. Model. 50, 742–754.

103. Qu, Y., Li, T., Liu, Z., Li, D., and Tong, W. (2023). DICTrank: The largest reference list of 1318 human drugs ranked by risk of drug-induced cardiotoxicity using FDA labeling. Drug Discov. Today 28, 103770.

104. Seal, S., Williams, D., Hosseini-Gerami, L., Mahale, M., Carpenter, A.E., Spjuth, O., and Bender, A. (2024). Improved detection of drug-induced liver injury by integrating predicted in vivo and in vitro data. Chem. Res. Toxicol. 37, 1290–1305.

105. Olubamiwa, A.O., Qu, Y., Connor, S., Tong, W., Li, D., and Chen, M. (2025). DILIrank 2.0: An updated and expanded database for drug-induced liver injury risk based on FDA labeling and a literature review. Drug Discov. Today 30, 104485.

106. Seal, S., Trapotsi, M.-A., Mahale, M., Subramanian, V., Greene, N., Spjuth, O., and Bender, A. (2025). PKSmart: an open-source computational model to predict intravenous pharmacokinetics of small molecules. J. Cheminform. 17, 147.

107. Marciano, D.P., Kuruvilla, D.S., Boregowda, S.V., Asteian, A., Hughes, T.S., Garcia-Ordonez, R., Corzo, C.A., Khan, T.M., Novick, S.J., Park, H., et al. (2015). Pharmacological repression of PPARγ promotes osteogenesis. Nat. Commun. 6, 7443.

