## Supplementary materials for "JUMP Cell Painting chemical dataset: Morphological map of 115,796 compounds in human cells"

#### Supplementary File PROTOCOL: Protocol variations for the Cell Painting assay across JUMP Cell Painting Consortium partners

This Excel file details the methodological parameters and operational variations of the Cell Painting protocol as implemented at each of the sources.

#### Supplementary File PHENOTYPIC_CONSISTENCY: Top phenotypically consistent classes, for multiple annotation types

This PDF file shows phenotypic consistency analysis: how consistently compounds sharing the same annotation (target, mechanism, disease area, etc.) cluster together based on Cell Painting profiles, using normalized mean Average Precision (mAP) as the consistency metric and corrected p-values for significance. The nine annotation sources tested are: Chemical Probes Portal (chemical_probes), disease area (disease_area), MOA (moa), MOTIVE bio KG (Knowledge Graph) (motive_biokg), MOTIVE open targets (motive_opentargets), MOTIVE Prime KG (motive_primekg), Drug Repurposing Hub (repurposing), ToxCast assays (toxcast_assay), and UniProt (uniprot). For each we show: (1) A volcano-style scatter plot ("Target Consistency") of nMAP versus -log₁₀(corrected p-value). Points are sized by the number of compounds with the annotation and colored blue if significant (p<0.05) or grey if not, with a red dashed line at p=0.05. Notable high-consistency annotations are labeled directly. (2) A lollipop/dot plot ("Top Consistent Targets") listing of top-ranked classes in each annotation type by nMAP, with dot size reflecting the number of compounds with the annotation (also printed on the right) and blue/grey coloring for significance. (3) A grid of small UMAP-style embedding panels labeled “all” showing where each of the top-ranked annotations’ compounds fall in the global phenotypic embedding (grey background = all compounds), with active compounds in blue and inactive in pink, plus per-panel active counts and mAP. (4) A second grid of UMAP-style panels labeled "active," showing the same as (3) but plotted only over the active-compound embedding (the differently shaped grey landscape), again colored by active/inactive.

#### Supplementary File STRUCTURAL_CLASSES: Structural classes associated with Cell Painting phenotypic activity

This file lists all 385 structural classes significantly associated with Cell Painting phenotypic activity at 5% FDR (58 of 74 RDKit functional groups, 293 of 868 BRICS fragments, and 34 of 93 Bemis-Murcko scaffolds), listed with per-class compound counts, activity percentages, Fisher odds ratios with 95% confidence intervals, and Benjamini-Hochberg-corrected p-values, as described in Methods.

#### Supplementary File FUP: Predicted functional use probability classes’ phenotypic activity

Predicted functional use probability classes are listed with several properties, including Benjamini–Hochberg-corrected p-values indicating whether compounds in that class were statistically significantly enriched or depleted for having a Cell Painting phenotype, as described in Methods.

#### Supplementary File FUC: Functional use categories’ phenotypic activity

Functional use categories are listed with several properties, including Benjamini–Hochberg-corrected p-values indicating whether compounds in that category were statistically significantly enriched or depleted for having a Cell Painting phenotype, as described in Methods.

#### Supplementary File ACTIVITY_CLIFF_PAIRS: Activity-cliff pairs

This file contains all 32,366 unordered compound pairs with exact Morgan ECFP4 Tanimoto similarity >= 0.70 among compounds with valid source-7-excluded Cell Painting profiles and nonzero Morgan fingerprints. Both compounds had a phenotypic activity assessment for 32,335 pairs; 6,149 of these pairs (19.0%) contained exactly one compound detected as phenotypically active at 10% FDR and one assessed compound for which phenotypic activity was not detected. These pairs were designated activity cliffs. Morphological similarity and scaffold membership were not used to select the pairs. The file also retains 31 structurally close pairs containing at least one unassessed compound.

#### Supplementary File ACTIVITY_CLIFF_SCAFFOLD_GROUPS: Same-scaffold activity-cliff groups

This file summarizes the subset of candidate activity-cliff pairs that share a Bemis-Murcko scaffold. It contains one row for each of 1,419 scaffolds represented among 3,855 same-scaffold candidate pairs. For each scaffold, the file reports the number of pairs, unique compounds, active compounds, compounds for which activity was not detected, Tanimoto-similarity summaries, a representative pair, and the participating compound identifiers. Scaffold membership was assigned post hoc and was not used to select candidate pairs. Compounds may participate in more than one pair.

#### Supplementary File PHENOTYPIC_CLIFFS: Phenotypic cliffs

This file lists 4,089 phenotypic-cliff pairs among 17,061 phenotypically active compounds. The pairs were selected from the 1,202,120 unique unordered pairs in the union of each compound's 100 nearest structural neighbors. Each phenotypic cliff has Morgan-fingerprint cosine similarity, used as a Tanimoto proxy, >= 0.70 and Cell Painting profile cosine similarity < 0.30. The 4,089 pairs represent 31.6% of the 12,920 pairs above the structural-similarity cutoff and 0.34% of the complete k-nearest-neighbor pair universe. Both compounds in each pair are phenotypically active, distinguishing these pairs from activity cliffs, which are defined by one active and one not-detected-as-active compound.

#### Supplementary File ENRICHED: Enriched compounds

This Excel file contains lists of commercially available compounds (Molport, ≤ $200 per compound) drawn from the JUMP Cell Painting dataset that show statistically significant, reproducible morphological phenotypes (FDR-corrected p < 0.05 in the copairs activity test). Compounds were clustered in the 758-dimensional Cell Painting feature space by K-means and the top-N nearest to each cluster centroid were selected, yielding four configurations that trade off phenotypic diversity (number of clusters) against within-cluster replication (1 versus 2 compounds per cluster). 320 was chosen to fit on a 384-well plate with room for controls or test samples of interest.

#### Supplementary File CHEMBL: Compound-target activity annotations from ChEMBL 33 for JUMP Cell Painting compounds

Wide ChEMBL release 33 compound-target activity matrix for 17,892 JUMP compounds and 102 single-protein targets identified by UniProt accession. The file (original filename targetannotations_singleproteins_jumpcompounds_all_5.csv) contains the JUMP compound ID and standardized SMILES, with 1 for active, 0 for inactive, and blank for an untested or unretained compound-target pair. It contains 24,029 active and 44,182 inactive pairs.

#### Supplementary File TARGET2: Plate map for the JUMP Target-2 compound plates

The plate map file (original name Target_2_platemap.txt) provides compound names, IDs, plate well positions, and concentrations.

| **source** | **1** | **3** | **2** | **8** | **15** | **6** | **10** | **5** | **9** | **11** | **4** | **7** | **13** |
| --- | --- | --- | --- | --- | --- | --- | --- | --- | --- | --- | --- | --- | --- |
| 1 | c1 | c1 | c1 | c1 | c1 |  |  |  |  |  |  |  |  |
| 3 |  | c3 | c3 | c3 | c3 | c3 |  |  |  |  |  |  |  |
| 2 |  |  | c2 | c2 | c2 | c2 | c2 |  |  |  |  |  |  |
| 8 | c8 |  |  | c8 | c8 | c8 | c8 |  |  |  |  |  |  |
| 15 | c15 | c15 |  |  | c15 | c15 | c15 |  |  |  |  |  |  |
| 6 | c6 | c6 | c6 |  |  | c6 | c6 |  |  |  |  |  |  |
| 10 | c10 | c10 | c10 | c10 |  |  | c10 |  |  |  |  |  |  |
| 5 |  |  |  |  |  |  |  | c5 | c5 | c5 |  |  |  |
| 9 |  |  |  |  |  |  |  | c9 | c9 | c9 |  |  |  |
| 11 |  |  |  |  |  |  |  | c11 | c11 | c11 |  |  |  |
| 4 |  |  |  |  |  |  |  |  |  |  | c4 |  |  |
| 7 |  |  |  |  |  |  |  |  |  |  |  | c7 |  |
| 13 |  |  |  |  |  |  |  |  |  |  |  |  | c13 |

#### Supplementary Table 1: **Compound exchange logistics**.

Each of the wave 1 pharmaceutical partner sources (1, 2, 3, 6, 8, 10, 15) exchanged their ~12,000 nominated compounds (c1, c2, …) with four other sources while each of the wave 2 pharmaceutical partner sources (5, 9, 11) exchanged with two other sources. Source 4 (Broad Institute, mainly ORFs), Source 7 (chemical library at lower dose), and Source 13 (mainly CRISPRs, at the same laboratory as Source 7) did not exchange perturbations with other sources. Numbering is non-sequential for historical reasons as data was uploaded and omits Source 12 (a number held for a dataset that did not materialize). The 12,000 compounds were selected by the wave 1 and 2 partners to minimally overlap, with the exception of ~6,000 compounds bridging the two waves.

| **source** | **microscope_name** | **Widefield_vs_Confocal** | **excitation_type** | **objective_na** | **n_brightfield_planes_min** | **n_brightfield_planes_max** | **distance_between_z_microns** | **sites_per_well** | **filter_configuration** |
| --- | --- | --- | --- | --- | --- | --- | --- | --- | --- |
| 1 | Opera Phenix | Widefield | Laser | 1.0 | 1 | 1 |  | 4 | H |
| 2 | CV8000 | Confocal | Laser | 1.0 | 3 | 3 | 8 | 6 | A |
| 3 | Opera Phenix | Widefield | Laser | 1.0 | 0 | 3 | 5 | 9 | B |
| 4 | Opera Phenix | Widefield | Laser | 1.0 | 3 | 3 | 5 | 9 | B |
| 5 | CV8000 | Confocal | Laser | 0.75 | 3 | 3 | 5 | 9 | C |
| 6 | CV8000 | Confocal | Laser | 0.75 | 3 | 3 | 5 | 9 | A |
| 7 | CV7000 | Confocal | Laser | 0.75 | 0 | 0 |  | 9 | D |
| 8 | ImageXpress Micro Confocal | Confocal | LED | 0.75 | 0 | 0 | 3 | 9 | E |
| 9 | Opera Phenix | Widefield | Laser | 0.8 | 0 | 0 |  | 4 | H |
| 10 | CV8000 | Confocal | Laser | 0.75 | 3 | 3 | 5 | 6 | A |
| 11 | Operetta | Widefield | LED | 1.0 | 1 | 1 |  | 9 | F |
| 13 | CV7000 | Confocal | Laser | 0.75 | 0 | 0 |  | 9 | D |
| 15 | Opera Phenix | Widefield | Laser | 0.4 | 3 | 3 | 5 | 9 | G |

#### Supplementary Table 2: **Microscope settings across all sources**.

Column names are described in Supplementary Table 3. CSV version of this table can be downloaded from <https://github.com/jump-cellpainting/datasets/blob/main/metadata/microscope_config.csv>.

| **Column name** | **Description** |
| --- | --- |
| source | Data-generating center ID |
| microscope_name | Microscope model name |
| Widefield_vs_Confocal | One of: Widefield, Confocal |
| excitation_type | One of: Laser, LED |
| objective_na | Objective numerical aperture |
| n_brightfield_planes_min | Min number of brightfield planes taken |
| n_brightfield_planes_max | Max number of brightfield planes taken |
| distance_between_z_microns | Distance between Z planes in μm (only if > 1μm) |
| sites_per_well | Number of sites per well |
| filter_configuration | Filter configuration ID |

#### Supplementary Table 3: **Microscope settings columns**.

This table describes the Supplementary Table 2 columns.

| **filter_configuration** | **excitation_low_DNA** | **excitation_low_ER** | **excitation_low_RNA** | **excitation_low_AGP** | **excitation_low_Mito** | **excitation_high_DNA** | **excitation_high_ER** | **excitation_high_RNA** | **excitation_high_AGP** | **excitation_high_Mito** | **emission_low_DNA** | **emission_low_ER** | **emission_low_RNA** | **emission_low_AGP** | **emission_low_Mito** | **emission_high_DNA** | **emission_high_ER** | **emission_high_RNA** | **emission_high_AGP** | **emission_high_Mito** | **FPBase_config** |
| --- | --- | --- | --- | --- | --- | --- | --- | --- | --- | --- | --- | --- | --- | --- | --- | --- | --- | --- | --- | --- | --- |
| A | 405 | 488 | 488 | 561 | 640 | 405 | 488 | 488 | 561 | 640 | 422.5 | 500 | 581.5 | 581.5 | 661.5 | 467.5 | 550 | 618.5 | 618.5 | 690.5 | <http://broad.io/JUMPProductionConfigCV8000> |
| B | 405 | 488 | 488 | 561 | 640 | 405 | 488 | 488 | 561 | 640 | 435 | 500 | 570 | 570 | 650 | 480 | 550 | 630 | 630 | 760 | <http://broad.io/JUMPProductionConfigPhenix405> |
| C | 405 | 488 | 488 | 561 | 640 | 405 | 488 | 488 | 561 | 640 | 422.5 | 500 | 661.5 | 581.5 | 661.5 | 467.5 | 550 | 690.5 | 618.5 | 690.5 | <http://broad.io/JUMPProductionConfigCV8000> |
| D | 405 | 488 | 488 | 561 | 640 | 405 | 488 | 488 | 561 | 640 | 422.5 | 504.5 | 581.5 | 581.5 | 661.5 | 467.5 | 539.5 | 618.5 | 618.5 | 690.5 | <http://broad.io/JUMPProductionConfigCV7000> |
| E | 350 | 458 | 511 | 544 | 617 | 404 | 492 | 551 | 576 | 645 | 417 | 516 | 573 | 604 | 672 | 477 | 556 | 613 | 644 | 712 | <http://broad.io/JUMPProductionConfigImageXpress> |
| F | 365 | 475 | 510 | 550 | 630 | 365 | 475 | 510 | 550 | 630 | 430 | 500 | 525 | 600 | 655 | 500 | 550 | 580 | 640 | 760 | <http://broad.io/JUMPProductionConfigOperetta> |
| G | 375 | 488 | 488 | 561 | 640 | 375 | 488 | 488 | 561 | 640 | 435 | 500 | 500 | 570 | 650 | 480 | 550 | 550 | 630 | 760 | <http://broad.io/JUMPProductionConfigPhenix375> |
| H | 375 | 488 | 488 | 561 | 640 | 375 | 488 | 488 | 561 | 640 | 435 | 500 | 570 | 570 | 650 | 480 | 550 | 630 | 630 | 760 | <http://broad.io/JUMPProductionConfigPhenix375> |

#### Supplementary Table 4: **Filter configuration across all sources**.

Column names are described in Supplementary Table 5. CSV version of this table can be downloaded from <https://github.com/jump-cellpainting/datasets/blob/main/metadata/microscope_filter.csv>.

| **Column name** | **Description** |
| --- | --- |
| filter_configuration | Filter configuration ID |
| excitation_low_DNA | Excitation wavelength min, DNA channel |
| excitation_low_ER | Excitation wavelength min, ER channel |
| excitation_low_RNA | Excitation wavelength min, RNA channel |
| excitation_low_AGP | Excitation wavelength min, AGP channel |
| excitation_low_Mito | Excitation wavelength min, Mito channel |
| excitation_high_DNA | Excitation wavelength max, DNA channel |
| excitation_high_ER | Excitation wavelength max, ER channel |
| excitation_high_RNA | Excitation wavelength max, RNA channel |
| excitation_high_AGP | Excitation wavelength max, AGP channel |
| excitation_high_Mito | Excitation wavelength max, Mito channel |
| emission_low_DNA | Emission wavelength min, DNA channel |
| emission_low_ER | Emission wavelength min, ER channel |
| emission_low_RNA | Emission wavelength min, RNA channel |
| emission_low_AGP | Emission wavelength min, AGP channel |
| emission_low_Mito | Emission wavelength min, Mito channel |
| emission_high_DNA | Emission wavelength max, DNA channel |
| emission_high_ER | Emission wavelength max, ER channel |
| emission_high_RNA | Emission wavelength max, RNA channel |
| emission_high_AGP | Emission wavelength max, AGP channel |
| emission_high_Mito | Emission wavelength max, Mito channel |
| FPBase_config | Fluorescence spectra config URL |

#### Supplementary Table 5: **Filter configuration columns**.

This table describes the Supplementary Table 4 columns.

| **ID** | **jcp2022_id** | **broad_sample** | **pert_iname** |
| --- | --- | --- | --- |
| C1 | JCP2022_085227 | BRD-K91188791-001-17-5 | aloxistatin |
| **C2** | **JCP2022_037716** | **BRD-K21728777-001-02-3** | **AMG900** |
| C3 | JCP2022_025848 | BRD-K38775274-001-22-1 | dexamethasone |
| C4 | JCP2022_046054 | BRD-K58550667-001-08-7 | FK-866 |
| **C5** | **JCP2022_035095** | **BRD-K47557313-001-02-7** | **LY2109761** |
| C6 | JCP2022_064022 | BRD-K28132190-001-02-0 | NVS-PAK1-1 |
| **C7** | **JCP2022_050797** | **BRD-K59632282-052-03-1** | **quinidine** |
| **C8** | **JCP2022_012818** | **BRD-K89517477-001-01-4** | **TC-S-7004** |

#### Supplementary Table 6: Compound positive controls.

Eight positive control compounds were chosen based on the diversity of their profiles (distinct from each other and DMSO). All eight compounds were included in each compound plate, and the four in bold were included in ORF plates. See Methods for details about how they were chosen, as they are not conventional strong positive controls.

| **Metadata_Source** | **Metadata_CellProfiler_Version** |
| --- | --- |
| source_1 | 4.1.3 |
| source_2 | 4.0.7 |
| source_3 | 4.2.1 |
| source_4 | 4.1.3 |
| source_5 | 4.1.3 |
| source_6 | 4.1.3 |
| source_7 | 4.2.1 |
| source_8 | 4.1.3 |
| source_9 | 4.1.3 |
| source_10 | 4.1.3 |
| source_11 | 4.1.3 |
| source_13 | 4.2.1 |
| source_15 | 4.1.3 |

#### Supplementary Table 7: CellProfiler version across sources.

CSV version of this table can be downloaded from https://github.com/jump-cellpainting/datasets/blob/main/metadata/cellprofiler_version.csv

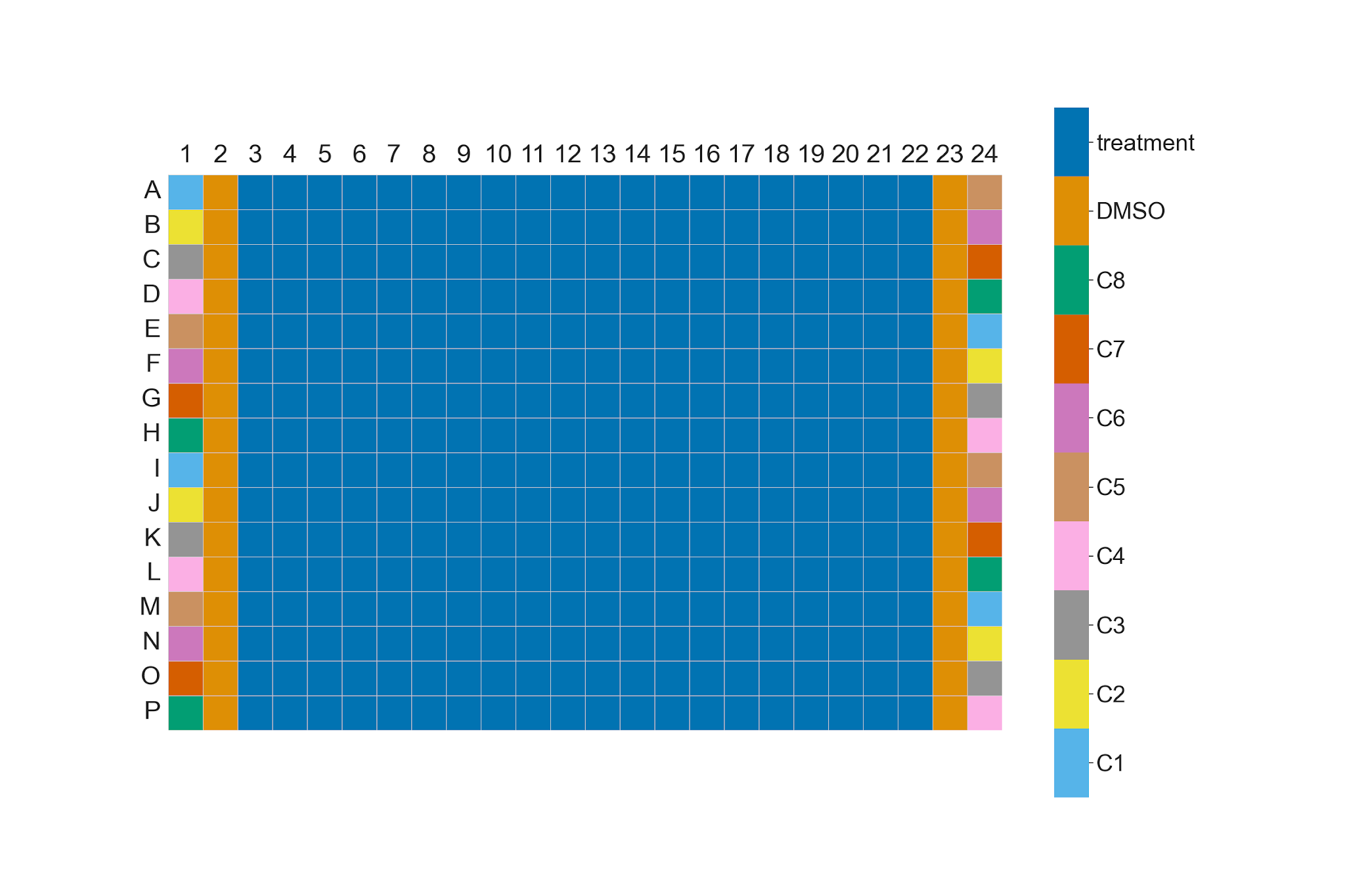

#### [Supplementary Figure](https://drive.google.com/file/d/1cSK-mw_HyMhxadSyniO2Tkp6THJoUIwT/view?usp=drive_link) 1: **Compound plate layout in a 384-well plate**.

Columns 1 and 24 contain the positive control compounds, while columns 2 and 23 contain the negative control, DMSO. The inner columns contain compound treatments. The identity of the positive control compounds is provided in Supplementary Table 6.

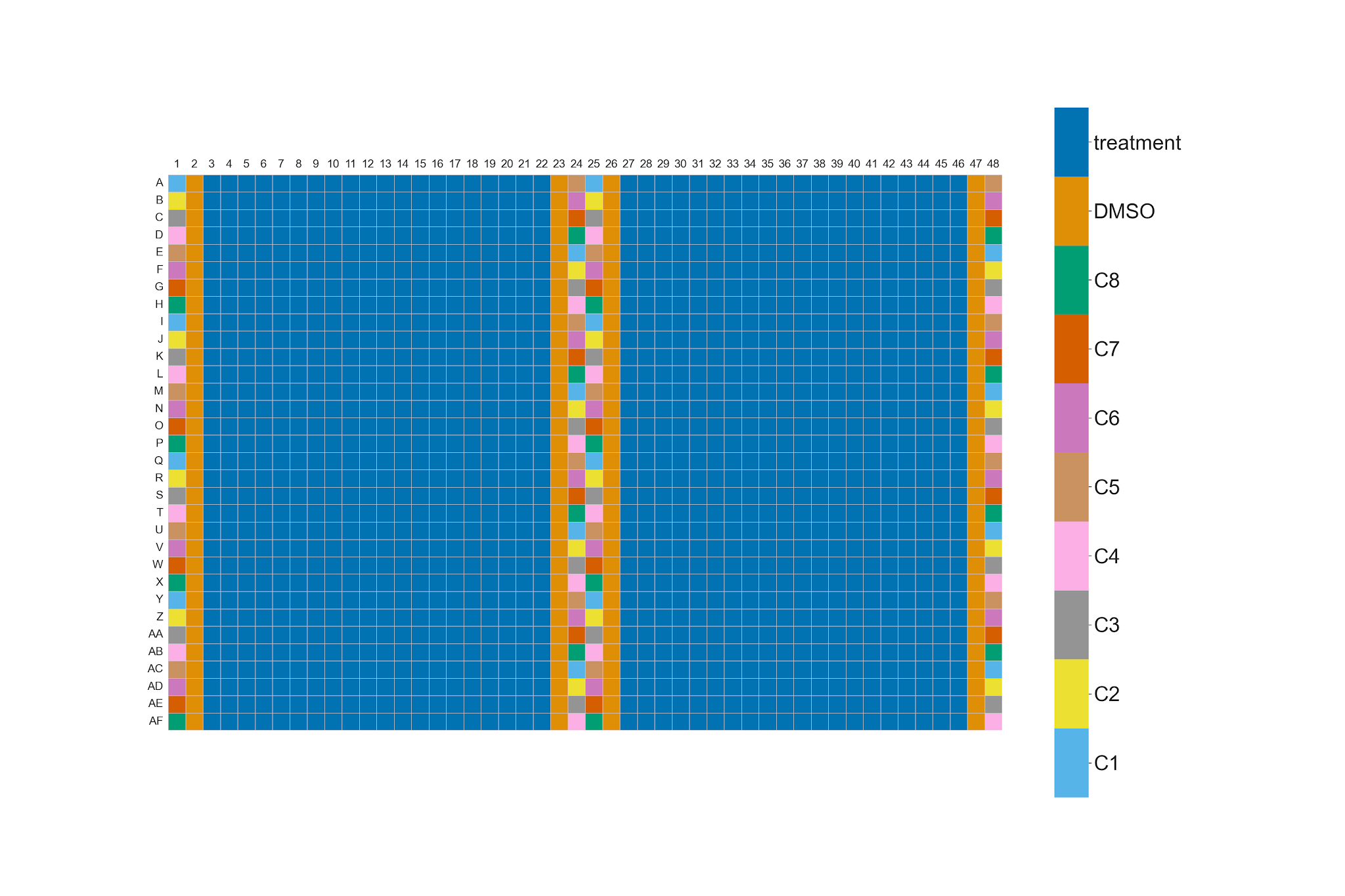

#### [Supplementary Figure](https://drive.google.com/file/d/1OnTivoyQTZajbhkbFK50DAwyY3b9fC-I/view?usp=drive_link) 2: **Compound plate layout in a 1536-well plate**.

A 1536-well compound plate is made of a 384-well plate in each of its quadrants. Hence, it contains positive control and DMSO wells, both in the outer columns and in the inner columns. The identity of the positive control compounds is provided in Supplementary Table 6.

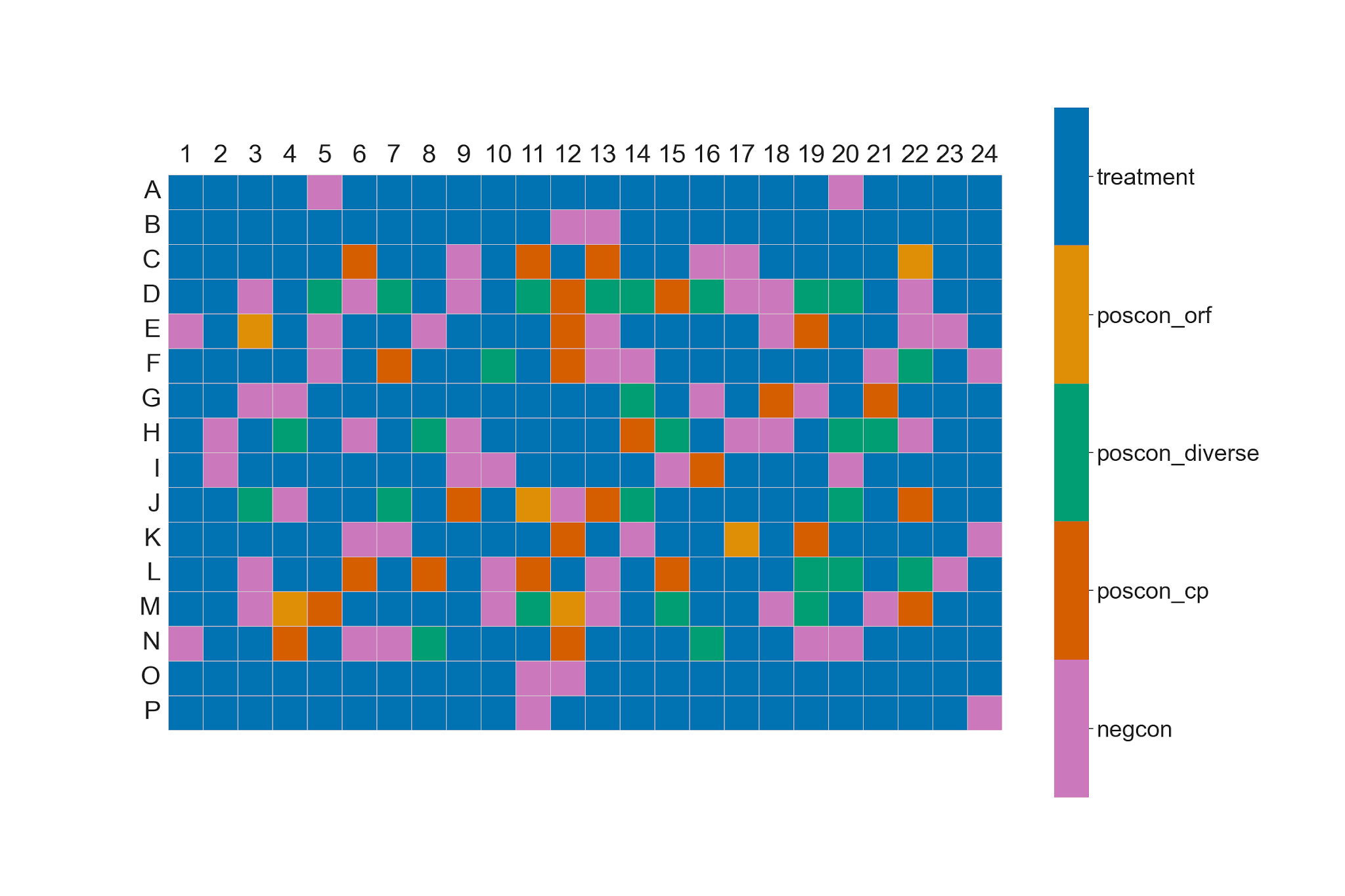

#### [Supplementary Figure](https://drive.google.com/file/d/1GjBVICv6r8i_zszvdl1i4wks3bT-p22l/view?usp=drive_link) 3: **JUMP-Target-2-Compound plate map**.

The plate consists of 301 compounds, including three different types of positive controls (poscon_orf, poscon_diverse and poscon_cp). It also contains 64 DMSO (negcon) wells. Additional details about the plate map, and how the number of unique compounds changed from 306 to 301, are available at <https://github.com/jump-cellpainting/JUMP-Target>.

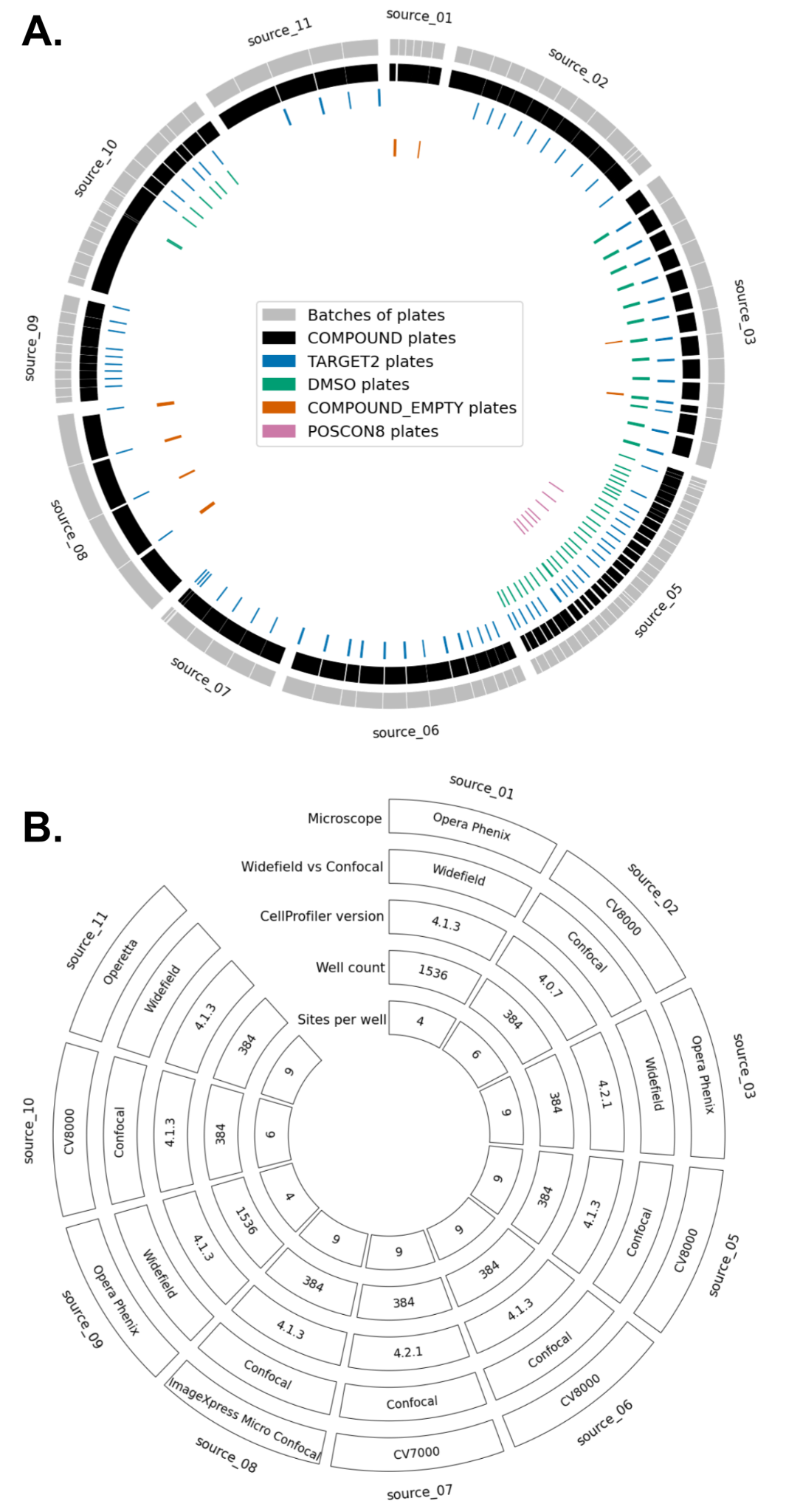

#### Supplementary Figure 4: Overview of samples processed and technical parameters at the 10 sources generating chemical perturbation data.

Sources 4 and 13 are omitted because neither ran a nominated-compound screen; their compound data were reference plates only. (A) Map showing the various plates and batch structure for compounds tested at 10 of the partners with usable data. Source 15 used alternative imaging channels and is excluded; the nine remaining pharmaceutical company partners are shown, plus one non-profit. (B) Map showing the technical parameters for data from each partner.

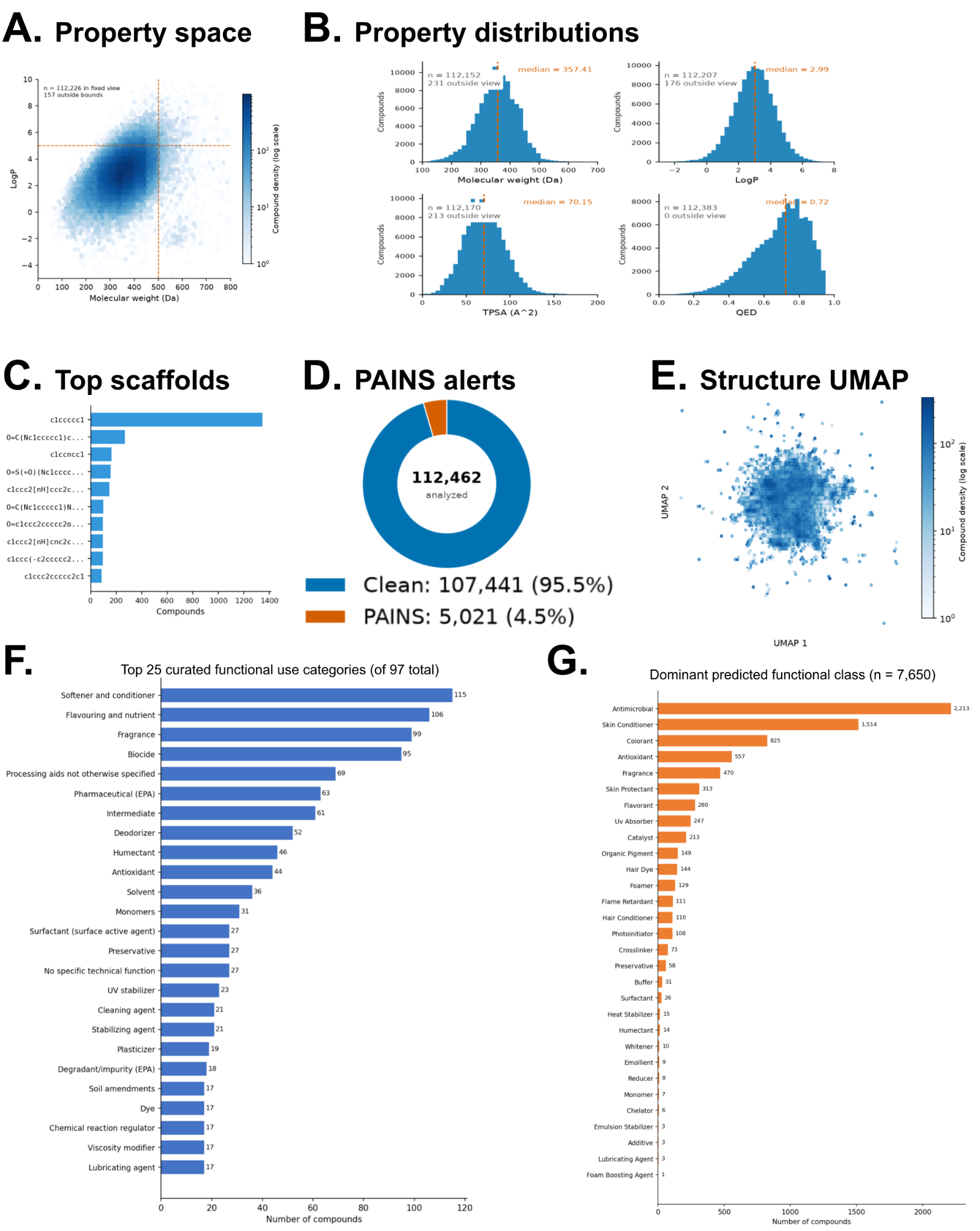

#### Supplementary Figure 5: Overview of the chemical properties of the 112,462 analyzed compounds in the JUMP compound data set.

(A) Property space - joint distribution of molecular weight and calculated LogP, the two axes of oral drug-likeness. (B) Distributions of four core physicochemical properties (molecular weight (MW), calculated octanol-water partition coefficient (lipophilicity, LogP), topological polar surface area (TPSA), and Quantitative Estimate of Drug-likeness (QED)). (C) Top Bemis-Murcko scaffolds by compound count. (D) Pan-assay interference compounds (PAINS) exist in the compound set. (E) Structure UMAP (Uniform Manifold Approximation and Projection) of Morgan-fingerprint chemical space. (F-G) Of 112,462 compounds (source 7 excluded), 8,559 mapped to DSSTox substance identifiers (DTXSIDs) and were classified by the EPA functional use pipeline. (F) Distribution of the top 25 curated functional use categories (FUCs) assigned by expert review, covering 427 compounds across 97 categories in the analysis set. Bars show the number of compounds with each curated label; a compound may carry multiple FUC labels. (G) Distribution of dominant predicted functional use class for 7,651 compounds with model-derived functional use probabilities (FUPs) across 35 classes. Each compound is assigned to a single class by argmax over its predicted probability vector.

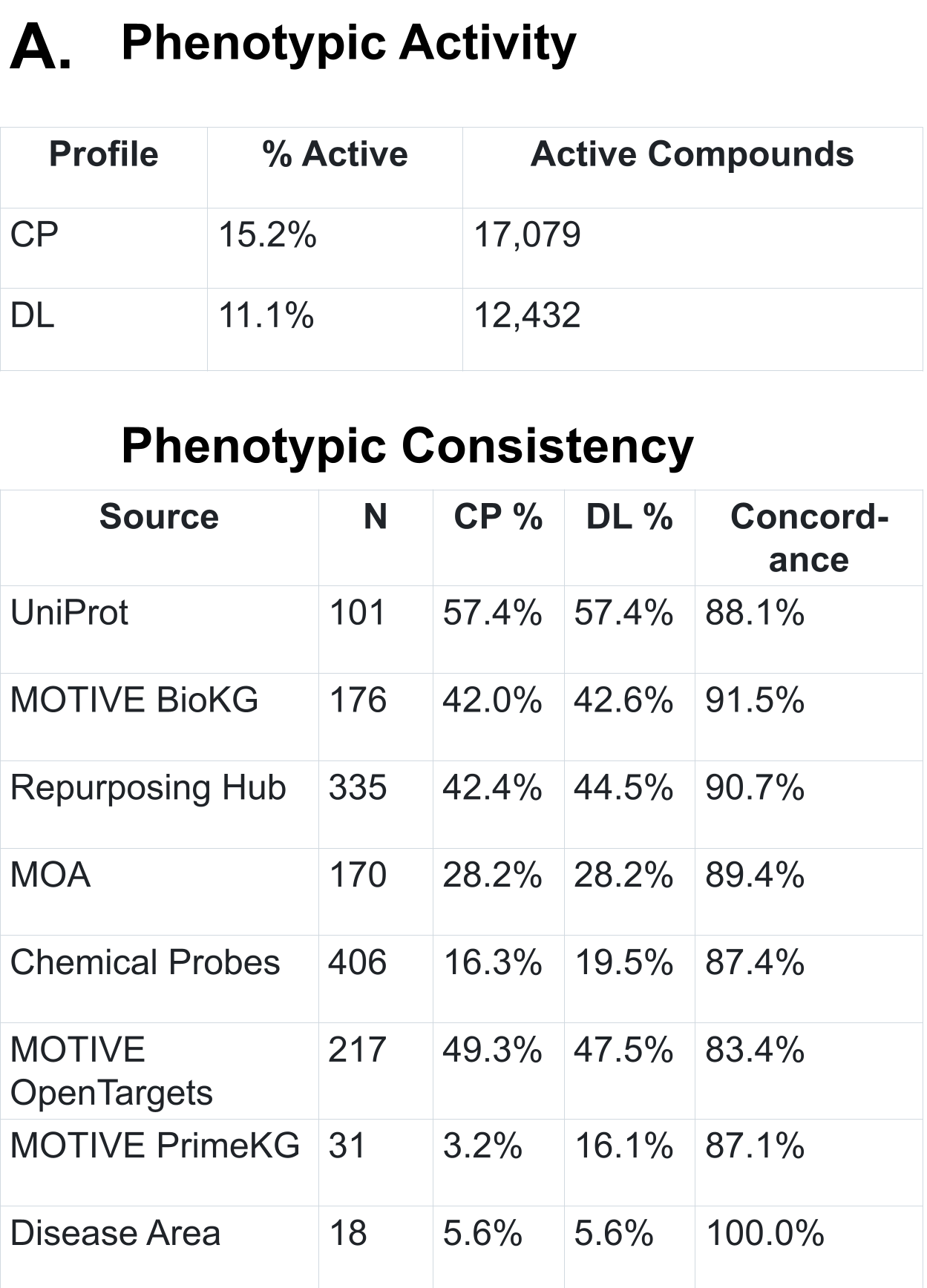

#### Supplementary Figure 6: Phenotypic activity and consistency analysis shows deep learning based features (DL) perform similarly to CellProfiler (CP)-based engineered features.

(A) Comparison of engineered CellProfiler features versus deep learning embeddings in terms of phenotypic activity (top) and phenotypic consistency (bottom). N = number of classes analyzed for the annotation type. CP and DL % refers to the percentage of classes that are phenotypically consistent (above the threshold). Concordance = (both_consistent + neither_consistent) / n_targets. Phenotypic activity uses FDR < 0.10, 112,462 compounds. Overlap: 10,893 both, 6,176 CP-only, 1,532 DL-only (93.1% concordance). Phenotypic consistency uses p < 0.05, shared pool (union of actives from either method = 18,601 perturbations).

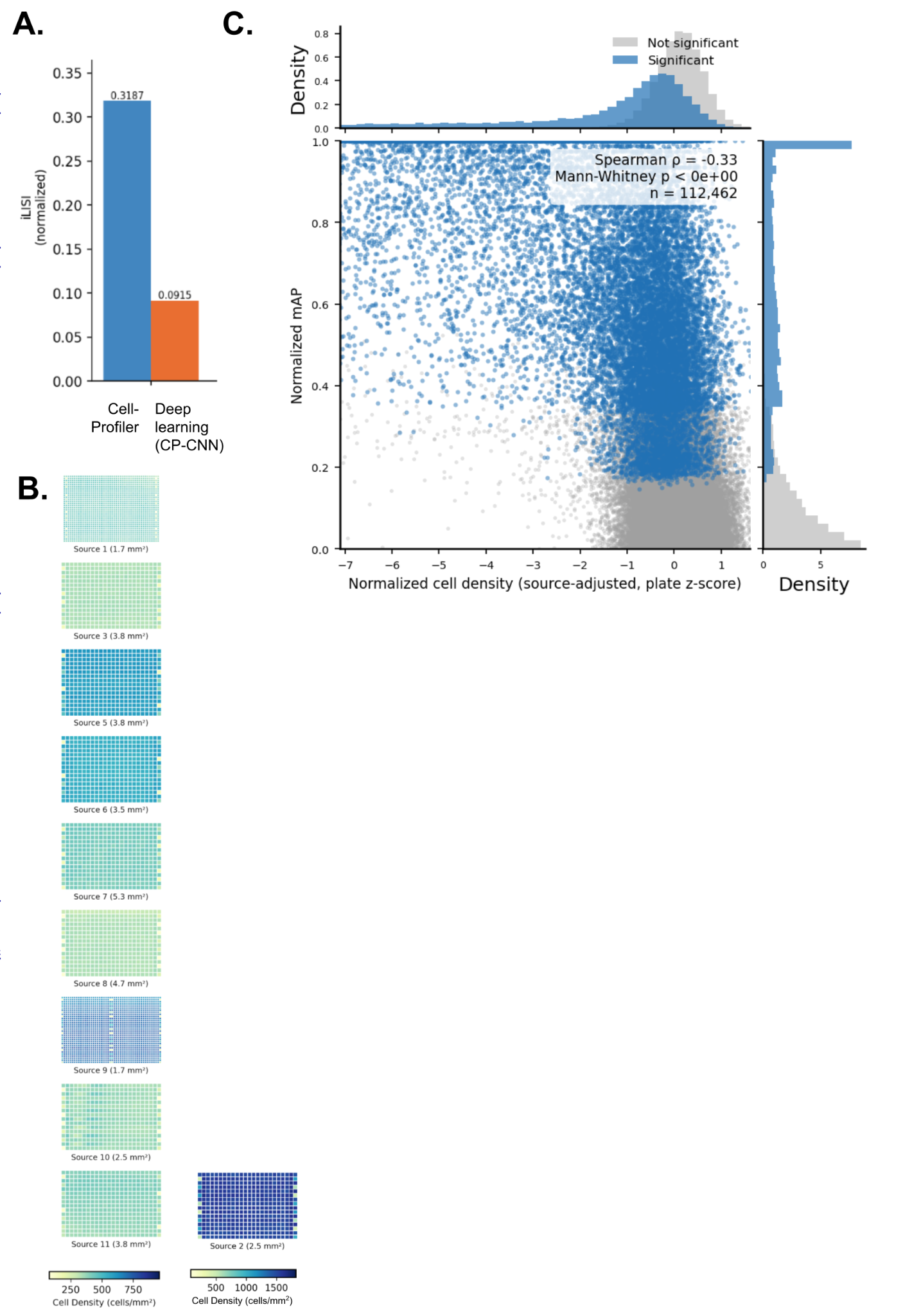

#### Supplementary Figure 7: Batch effects and plate layout effects in relationship to c**ell density** and phenotypic activity**.**

(A) Cross-source batch mixing after Harmony correction, measured by iLISI (higher = better mixing). CellProfiler features integrate across sources 3.5x better than deep-learning CP-CNN features (normalized iLISI 0.32 vs 0.09); not shown: including Source 7 does not change this (0.30 vs 0.09). (B) No major plate layout effects (well position effects, including row or column patterns) were observed in cell density across sources, other than what can be explained by the position of controls across plates. Source 2 is shown separately given its dramatically different color scale. (C) Post-treatment cell density versus phenotypic activity for 112,462 compounds. Cell density (cells per mm2) was source-adjusted and robustly z-scored within plate (0 = plate median; negative = fewer cells), then median-aggregated per compound. Points are colored by activity call (significant, FDR < 0.10, versus not significant); marginal density histograms show each group's distribution. Compounds with fewer cells after treatment tend to have higher phenotypic activity (Spearman rho = -0.33).

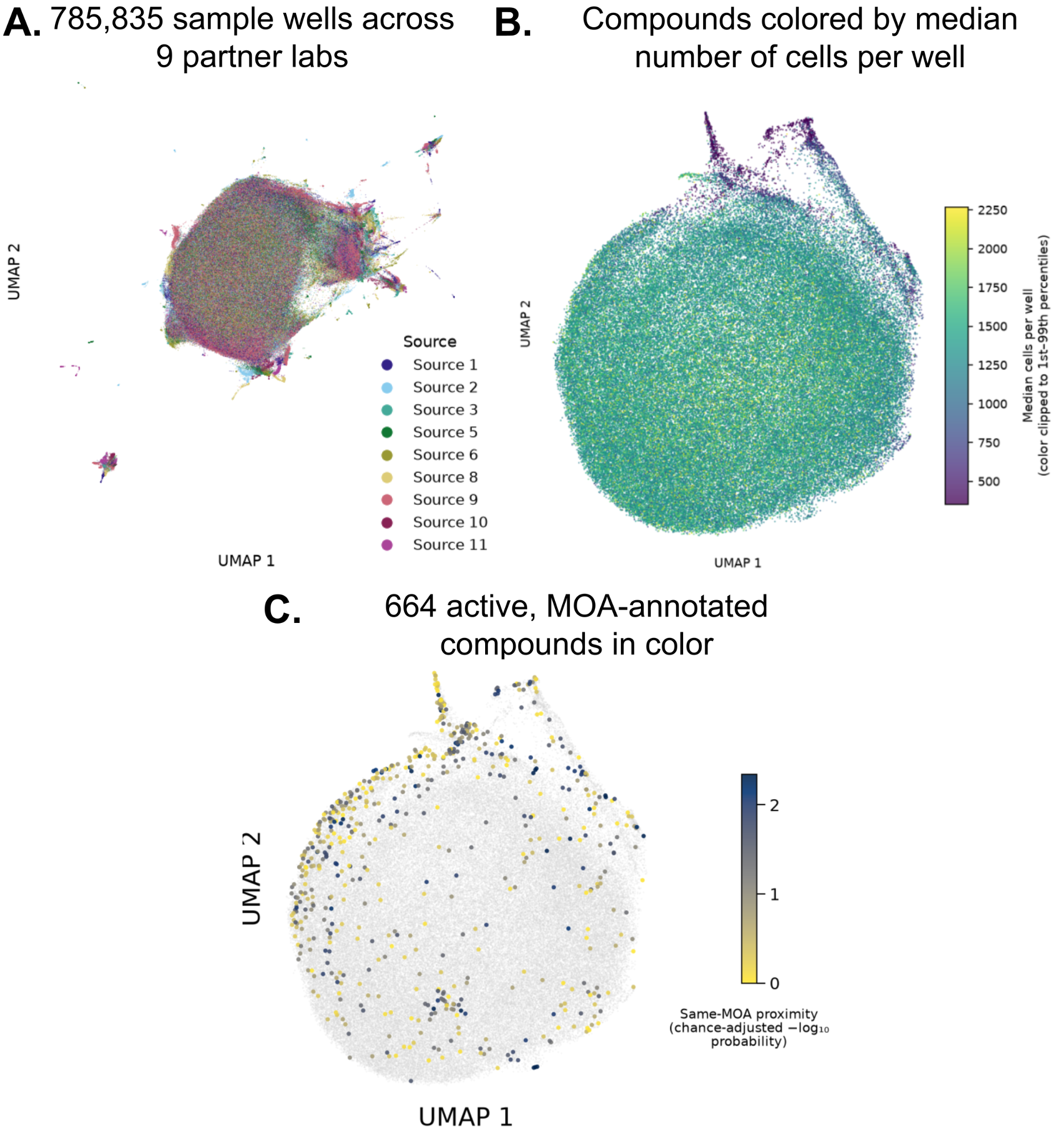

#### Supplementary Figure 8: Qualitative UMAP visualizations by source, cell count, and same-MOA similarity

(A) UMAP of the well-level profiles colored by sources suggests reasonable integration of the sources shown. Source 7 is excluded from this figure and from the main analysis under the concentration/design rule (Methods)..

(B) Cosine UMAP gives the same global view of JUMP compounds as in Figure 2A but colored here by the median number of cells per well (clipped to the 1st and 99th percentiles for visualization).

(C) Cosine UMAP gives the same global view of JUMP compounds as in Figure 2A but only the compounds annotated with an MOA are colored, and the intensity of the color corresponds to how close the compound is to its same-MOA class members (in high-dimensional space, not UMAP space).

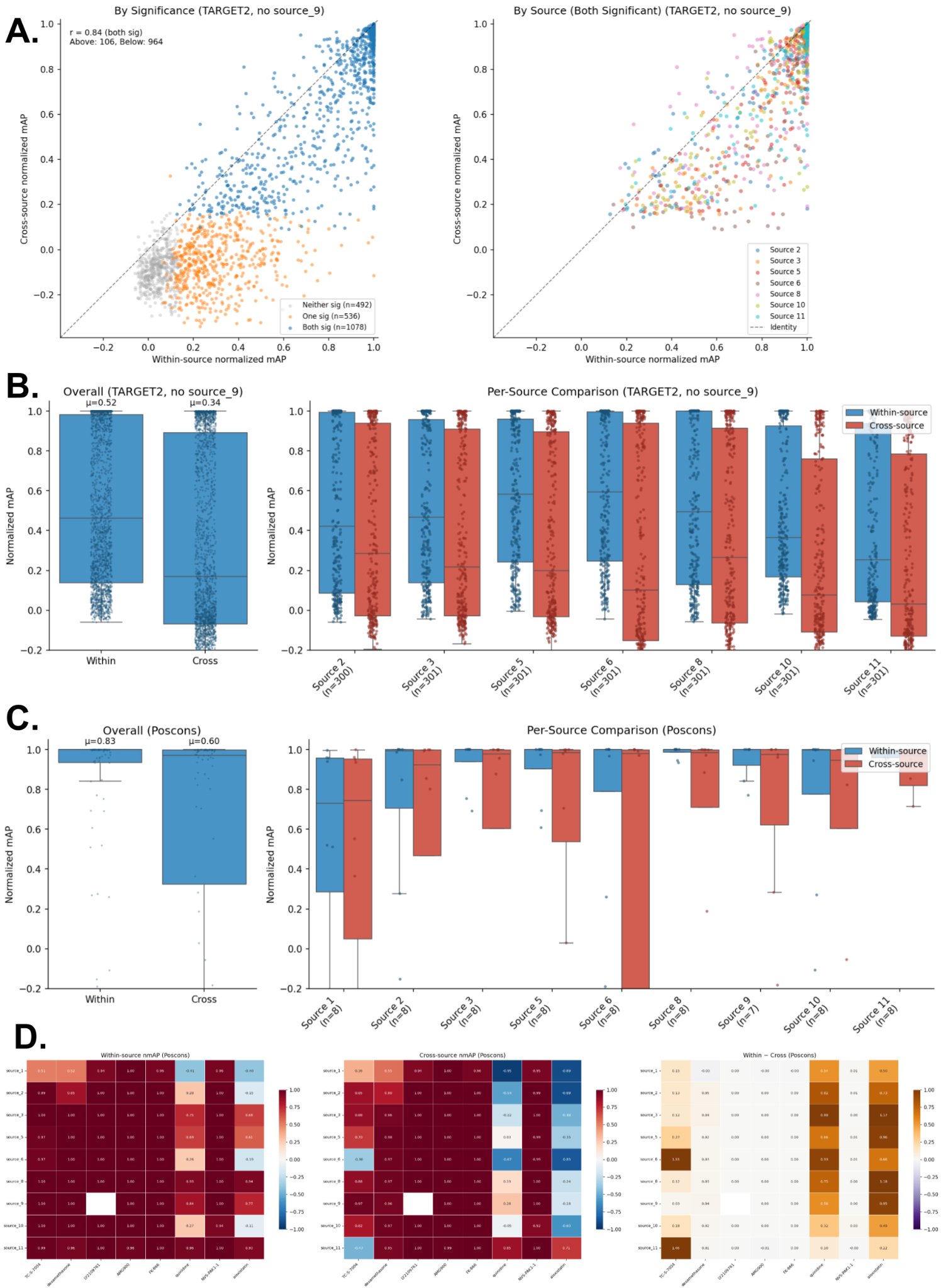

#### Supplementary Figure 9: Phenotypic activity, as judged by replicate retrieval, decreases across sources versus within a given source, as expected.

(A) The within-source nMAP is generally higher than cross-source nMAP for Target-2 compounds. Each point is one compound-source pair; Pearson r = 0.84 across n = 1,078 pairs with significant activity in both the within-source and cross-source analyses. (B) Analysis of each source individually shows the same trends; μ denotes the mean nMAP, with some sources better aligned (showing higher cross-source phenotypic activity, measured by nMAP) than others. Excluded sources: Source 1 did not include Target-2 plates in their batches; Source 9’s Target-2 plates showed near-zero phenotypic activity despite strong poscon performance, indicating an accidental lack of compound treatment, which we confirmed only impacted Target-2 plates; Sources 4 and 13 contributed primarily genetic perturbation data and were excluded from compound harmonization, precluding analysis of their Target-2 plates; Source 15 used alternative imaging channels; Source 12 does not exist (the number was held for a dataset that could not be contributed). (C) The experiment included eight so-called positive controls (C1: aloxistatin, C2: AMG900, C3: dexamethasone, C4: FK-866, C5: LY2109761, C6: NVS-PAK1-1, C7: quinidine, C8: TC-S-7004). These are not conventional positive controls (see Methods for details). (D) nMAP of the eight positive controls are shown for each source individually. All are phenotypically active under the right conditions: all yield mAP>0.50 in at least half the sources, but they show varying degrees of phenotypic activity across sources and batches. In particular, quinidine and aloxistatin are less strong than the rest (left), and less robust across sources (middle, right).

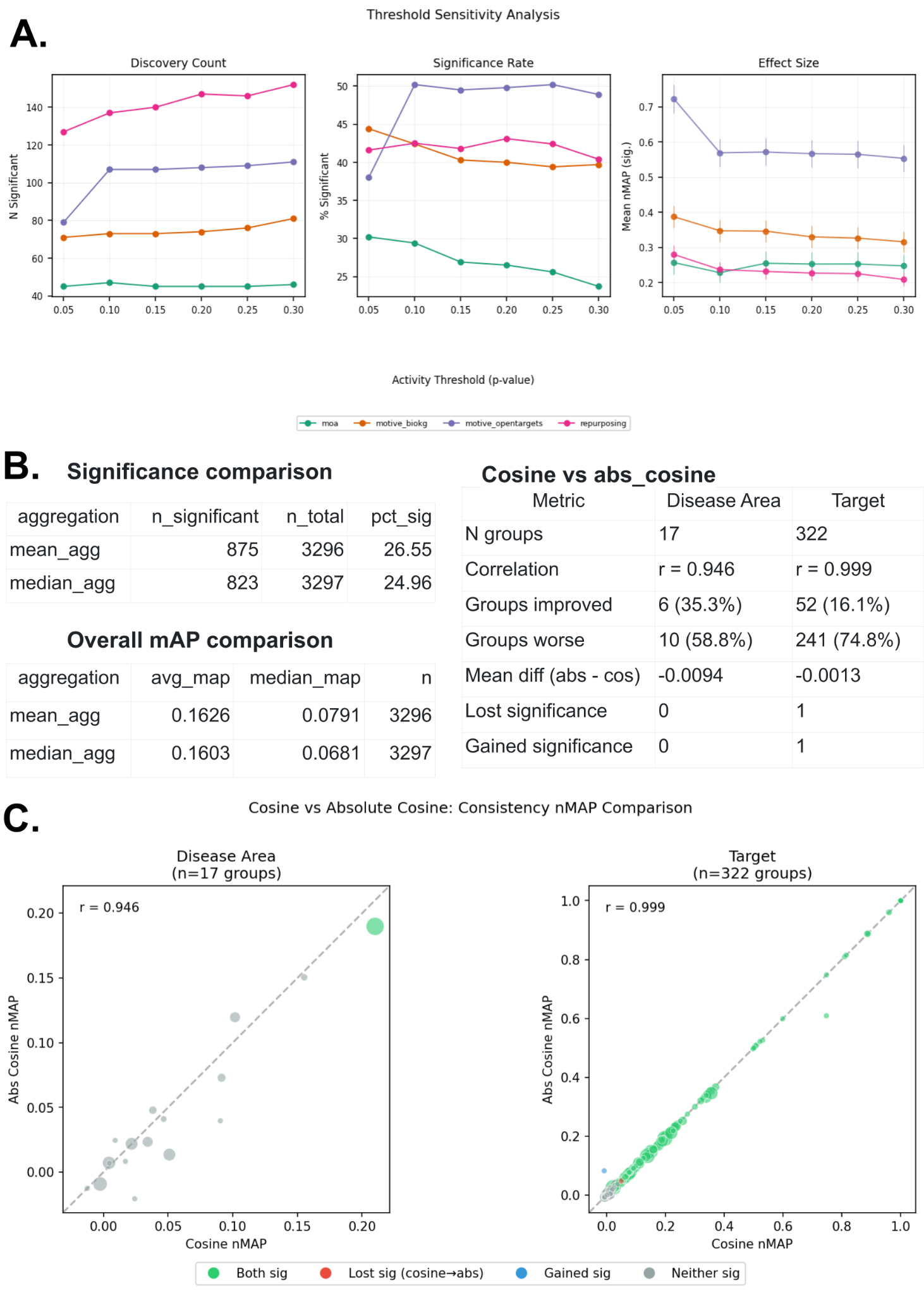

#### Supplementary Figure 10: Alternative analytics did not improve phenotypic consistency measures.

(A) Threshold sensitivity analysis across annotation types. Impact of activity p-value threshold on consistency results for four annotation types. Left: Number of annotation groups with statistically significant consistency (corrected p < 0.05). Middle: Percentage of tested groups reaching significance. Right: Mean normalized average precision (nMAP) computed over significant groups only, with error bars showing standard error of the mean. (B) Left: Phenotypic consistency does not improve when replacing mean average precision with median average precision (across compounds in a class). Likely, median suppresses cases where a few compounds in the class show strong similarity but most do not. Right: Phenotypic consistency does not improve when using absolute values for the cosine similarity, that is, we don’t distinguish between a perturbation that has exactly the same effect on a cell versus one that has exactly the opposite effect. (C) When plotting phenotypic consistency using cosine similarity (x-axis) and absolute cosine similarity (y-axis), most annotated groups remain unchanged (or worsen) with absolute values, indicating that opposite profiles are rare in this dataset.

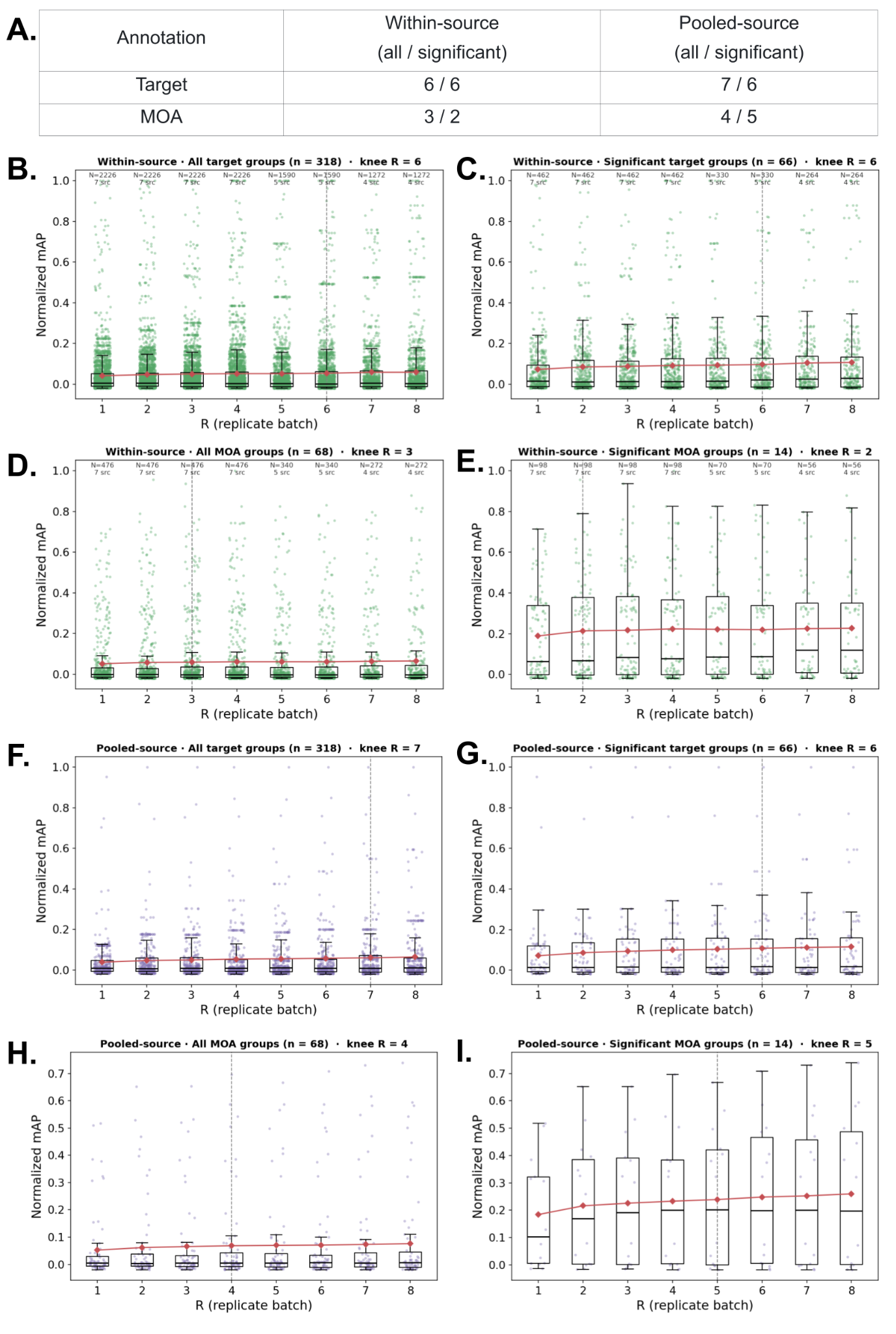

#### Supplementary Figure 11: Exploration of the number of replicates needed to achieve 90% of the maximal image profile signal.

(A) Summary table showing the number of replicates required to recover 90% of the full-depth phenotypic consistency signal for each annotation type (targets, MOA), group subset (within-source, pooled-source), and sampling scheme (all, significant); defined below. Compounds from the JUMP-Target-2 plate were analyzed here because they were repeatedly sampled across most laboratories. For each replicate count (R), one well per compound was selected from each of R sampled batches, compound profiles were median-aggregated, and the phenotypic consistency metric was recomputed. For each annotation type, consistency was computed over compounds carrying the corresponding target or MOA annotation group. Group-level nMAP summarizes phenotypic consistency: how well compounds sharing at least one target or MOA annotation retrieve one another relative to compounds sharing no such annotation. Shown in the table is the number of replicates (R) required to achieve 90% of the maximal signal. (B-E) Within-source combined analysis, in which replicate batches were sampled from one source (laboratory) at a time, and then summarized across eligible sources. Labels above each replicate count indicate the number of source-by-group observations and the number of contributing sources with at least R eligible batches. Analyses include all annotated groups with at least two Target-2 compounds (318 target groups and 68 MOA groups) in (B,D) and “significant” annotation groups in (C, E), which are just those groups showing statistically significant phenotypic consistency in the main analysis at corrected p < 0.05 (66 target groups and 14 MOA groups). (F–I) Pooled-source analysis, in which replicate batches were randomly sampled from the combined set of eligible Target-2 batches across all usable sources. For (B-I), box plots and jitter plots show the distribution of group-level nMAP values at each replicate count. In the within-source combined panels, points represent source-by-annotation-group observations. In the pooled-source panels, points represent individual annotation groups. Red diamonds and connecting lines show the mean nMAP across the plotted observations. The dashed vertical line marks the 90% knee, defined as the smallest replicate count at which the mean nMAP for that panel reached at least 90% of the value at the largest available replicate count.

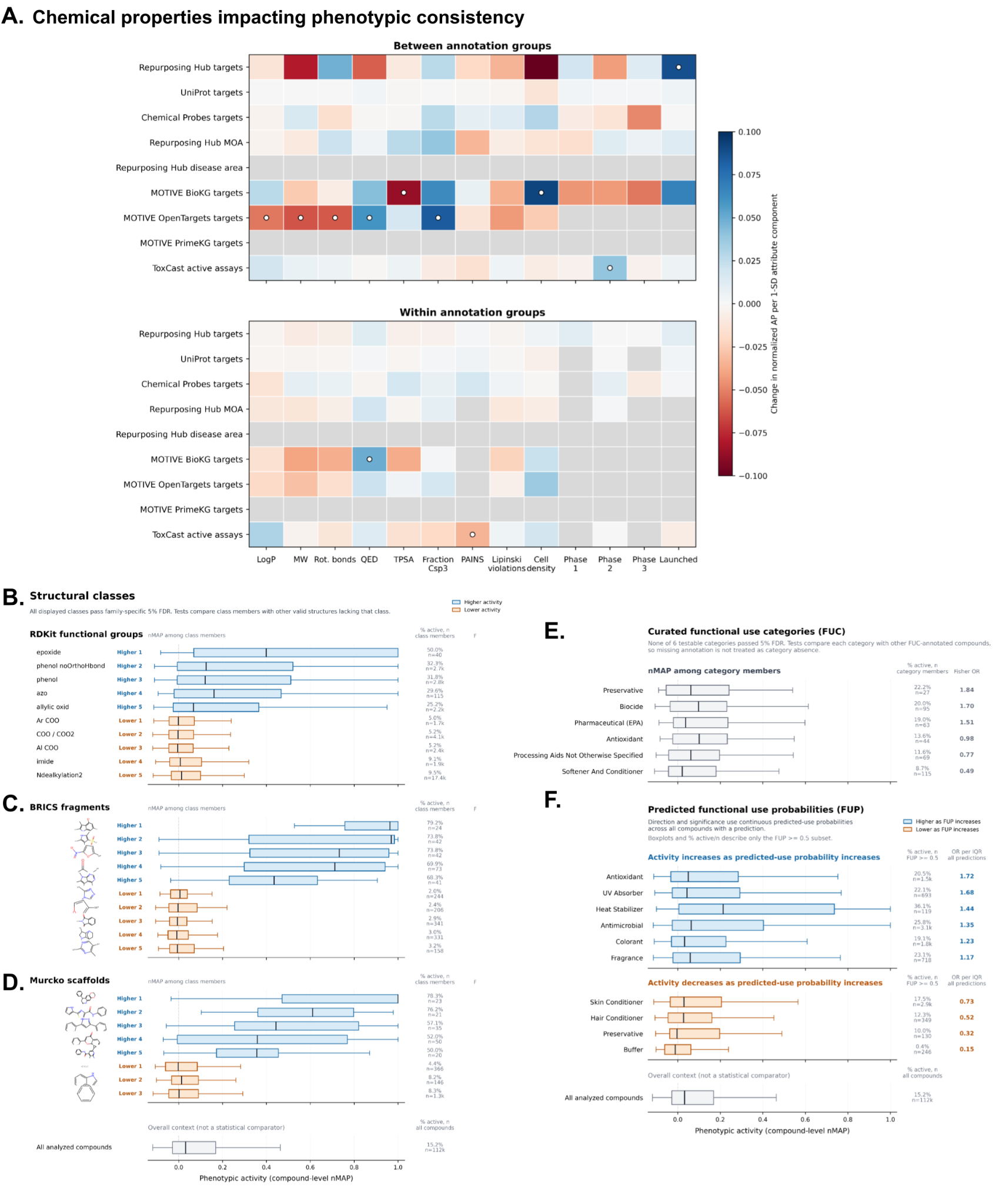

#### Supplementary Figure 12: Chemical properties associated with phenotypic activity and consistency

(A) Associations between compound attributes (columns) and phenotypic consistency in nine annotation sources (rows), among compounds passing the activity filter (corrected p < 0.1). Upper: whether groups with higher mean attribute values are more (blue) or less (red) consistent; for example, for *OpenTarget targets*, target groups with a higher mean *Fraction CSP3* have higher phenotypic consistency. Lower: whether compounds above their group's mean are more or less consistent than their group peers. Color gives the change in normalized AP per 1-SD increase. White dots mark global Benjamini-Hochberg q < 0.05, corrected separately within each panel. Undotted colored cells were tested but not significant; gray cells were not tested because fewer than 30 annotation groups or contributing compounds were available.

(B-D) Box plots and activity percentages are shown for compounds with each functional group (B), BRICS fragment (C), or Bemis-Murcko scaffold (D). Fisher odds ratios compare class members with other compounds lacking that class. Up to five statistically significant enriched (blue) or de-enriched (orange) structural components are shown; all 385 are provided in Supplementary File STRUCTURAL_CLASSES. Benjamini-Hochberg correction is applied separately across these three tested structure types (5% FDR). (E-F) Curated functional use categories (FUC, E) and predicted functional use probabilities (FUPs, F) box plots show compound categories that are more (blue) or less (orange) likely to be phenotypically active than other FUC-annotated compounds (missing FUC annotation is not treated as a category absence in Fisher tests). None of the six testable FUCs (gray) were significant at 5% FDR. FUP direction, color, OR per IQR (odds ratio per interquartile range), and significance come from logistic regression using continuous predicted use probability for every compound with a non-null value. All 10 endpoints passing the 5% FDR among 24 tested endpoints are shown. FUP blocks, plots, and displayed activity percentages summarize only compounds with probability ≥ 0.5.

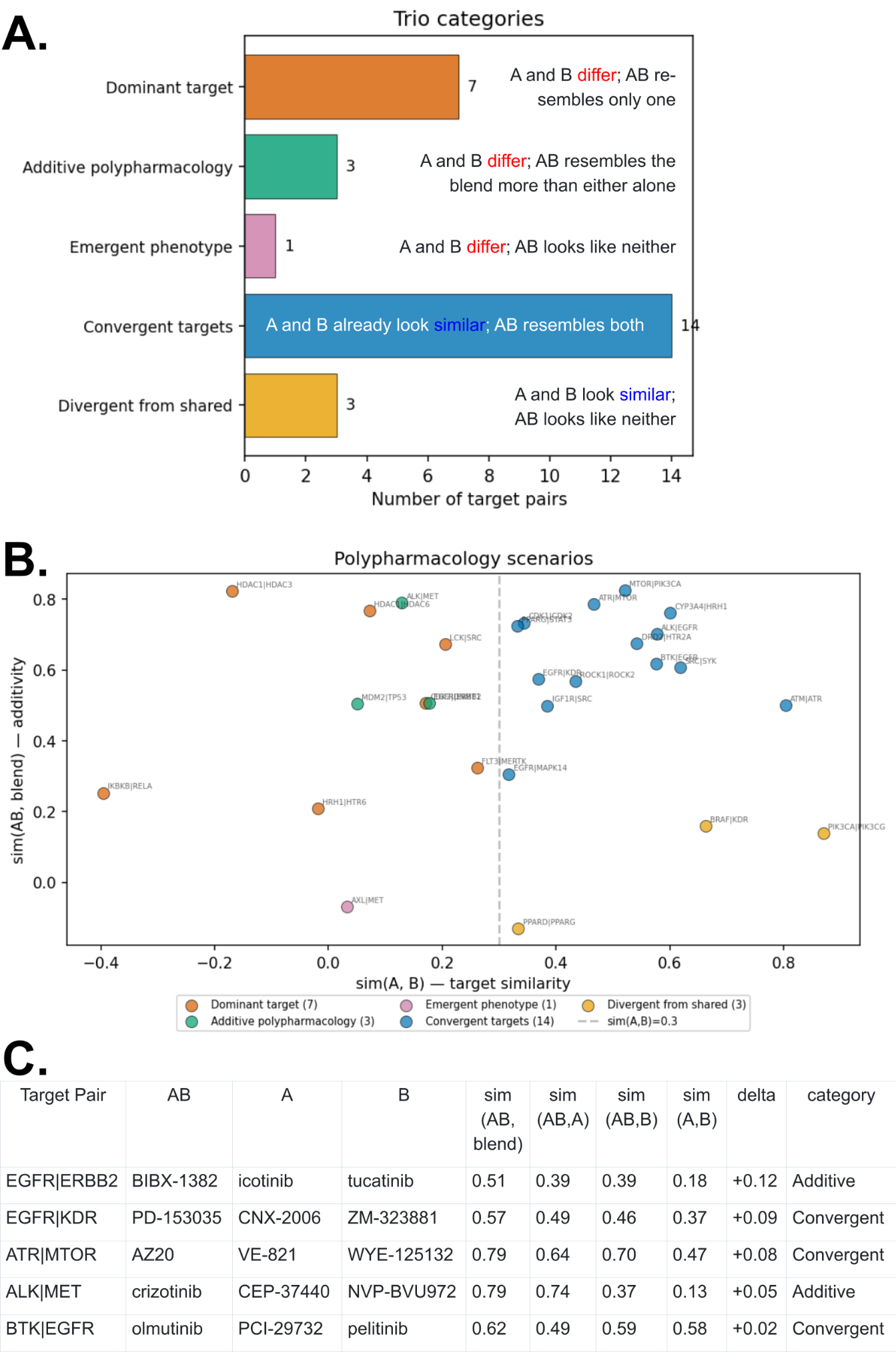

#### Supplementary Figure 13: Polypharmacologic compound profiles more often strongly matched one of their annotated targets rather than a blend of the two.

We selected trios of compounds where one is annotated as being a clean, dual-target compound (annotated as hitting exactly 2 targets (A, B) in the repurposing hub) plus two single-target compounds (each annotated as hitting only one of those targets, A or B), with all three compounds expressing a Cell Painting phenotype. Trios were then classified into five scenarios by comparing (using a cosine similarity threshold of 0.3 to mark as “similar”) the dual-target profile to the individual single-target profiles and to their computational blend; only the best trio per target pair is shown (28 pairs total; we dropped 4 pairs in which any of the three trio members had a nMAP < 0.5). (A) Counts of target pairs in each category: Emergent phenotype, Dominant target, Additive polypharmacology, Divergent from shared, and Convergent targets, as defined on each bar. (B) Scatter of each trio in the two-dimensional space used for classification, with sim(A, B) on the x-axis and sim(AB, blend) on the y-axis; the dashed vertical line at sim(A, B) = 0.3 separates the cases where the two single-target phenotypes are already similar (right) from the regime where the two single-target phenotypes are distinct (left). Points are colored by category and labeled with the two targets. (C) The top five trios with the largest delta (difference between sim(AB, blend) and the closer of sim(AB, A) and sim(AB, B)) are shown. The clearest additive cases are EGFR|ERBB2 and ALK|MET, where sim(A,B) is low and sim(AB, blend) exceeds the larger of sim(AB, A) and sim(AB, B); the remaining high-delta pairs (EGFR|KDR, ATR|MTOR, BTK|EGFR) are convergent, i.e. their two single-target profiles are already similar.

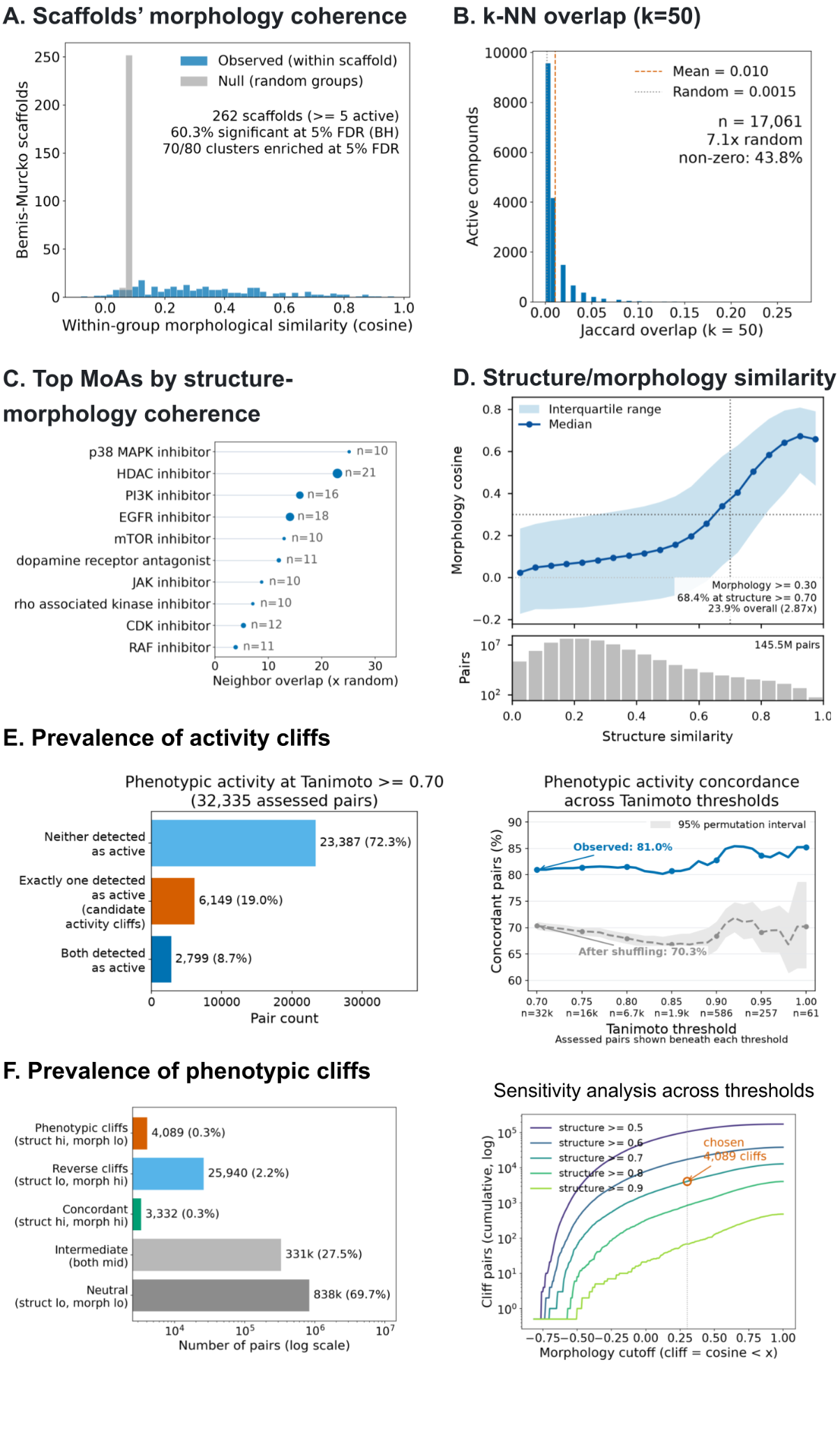

#### **Supplementary Figure** 14**: Compounds with similar chemical structure are more likely to share a similar morphological impact, with the exception of activity and phen**otypic cliffs**.**

(A) Compounds with shared scaffolds tend to yield similar morphological impact. For each of the 262 Bemis-Murcko scaffolds represented by at least 5 phenotypically active compounds, the mean pairwise morphological similarity among its members (observed, blue) is shown versus a size-matched permutation null (gray). 60.3% of scaffolds show greater self-similarity than expected by chance at 5% FDR using Benjamini-Hochberg correction. In a separate analysis not shown, 70 of 80 morphology-based KMeans clusters contain at least one over-represented Bemis-Murcko scaffold at 5% FDR. (B) Structure and morphology neighbor overlap more than expected by chance. For each of the 17,061 phenotypically active compounds, the Jaccard overlap is shown between its 50 nearest structural neighbors and its 50 nearest morphological neighbors (copairs mAP, FDR < 0.10). The overlap is small in absolute terms (mean 0.010 (dashed line), median 0, and only 43.8% of compounds share any neighbor), though 7x the chance expectation of 0.0015 (dotted red line). (C) Mechanism-of-action classes with the strongest structure-morphology coherence (neighbor-overlap enrichment over chance). Drug Repurposing Hub classes with n ≥ 10; dot area encodes the number of active compounds in the class. Note that the absolute values are small. (D) Morphological similarity increases with structural similarity across the 145,530,330 pairs among 17,061 phenotypically active compounds. The line and shading show the median and interquartile range of morphological cosine similarity within bins of Morgan ECFP4 cosine similarity; bars show pair counts. Pairs with structural similarity >= 0.70 were 2.87-fold enriched for morphological similarity >=0.3 (68.4% vs. 23.9% overall). (E) Activity cliffs. Structurally similar compounds had more concordant phenotypic activity status (both active or both inactive) than expected by chance (label shuffling), consistent with a general structure-activity relationship. Among 112,524 compounds having both a valid source-7-excluded perturbation-level Cell Painting profile and a nonzero Morgan ECFP4 fingerprint, all compound pairs were considered. We retained 32,366 pairs with exact Morgan Tanimoto >= 0.7; 32,335 of these had an activity assessment (compound-level mean normalized Average Precision above or below 10% FDR using Benjamini-Hochberg correction) for both compounds; these form the denominator in both plots. Left plot: Proportion of pairs in each category, with the orange middle bar showing 6,149 activity cliff pairs. Right plot: Sensitivity analysis across Tanimoto thresholds. At each threshold from 0.70 to 1.00, the blue curve gives the observed percentage of assessed pairs with concordant phenotypic activity status; the count printed below the threshold is the denominator. To establish the shuffled-label expectation, we kept the exact pair graph and the number of active compounds fixed, randomly reassigned the active labels among participating compounds, and recalculated concordance 2,000 times (seed 20260730). The gray dashed curve is the shuffled-label expectation and the gray band is its pointwise central 95% permutation interval. Bemis-Murcko scaffold membership was not used in the analysis, but we note that 3,855 activity cliffs share a scaffold, 2,291 have different scaffolds, and 3 lack a scaffold for at least one member. (F) Phenotypic cliffs. Left plot: The 1,202,120 unique pairs in the union of all per-compound structural k-nearest-neighbor lists (k = 100) among phenotypically active compounds are shown on a log scale and classified as phenotypic cliffs (high structural similarity, low morphological similarity), reverse cliffs (low-high), concordant (high-high), intermediate (both mid), or neutral (low-low). Structural similarity uses the Morgan-fingerprint cosine Tanimoto proxy, and morphological similarity uses Cell Painting profile cosine similarity. Right plot: Threshold sensitivity within the same k-nearest-neighbor pair set. Cumulative cliff counts are shown across morphological cosine cutoffs, with one curve per structural-similarity cutoff. The selected definition yielding 4,089 cliffs is marked; this number is threshold-dependent and approximate (structural proxy >= 0.70 and morphology cosine < 0.30; open marker).

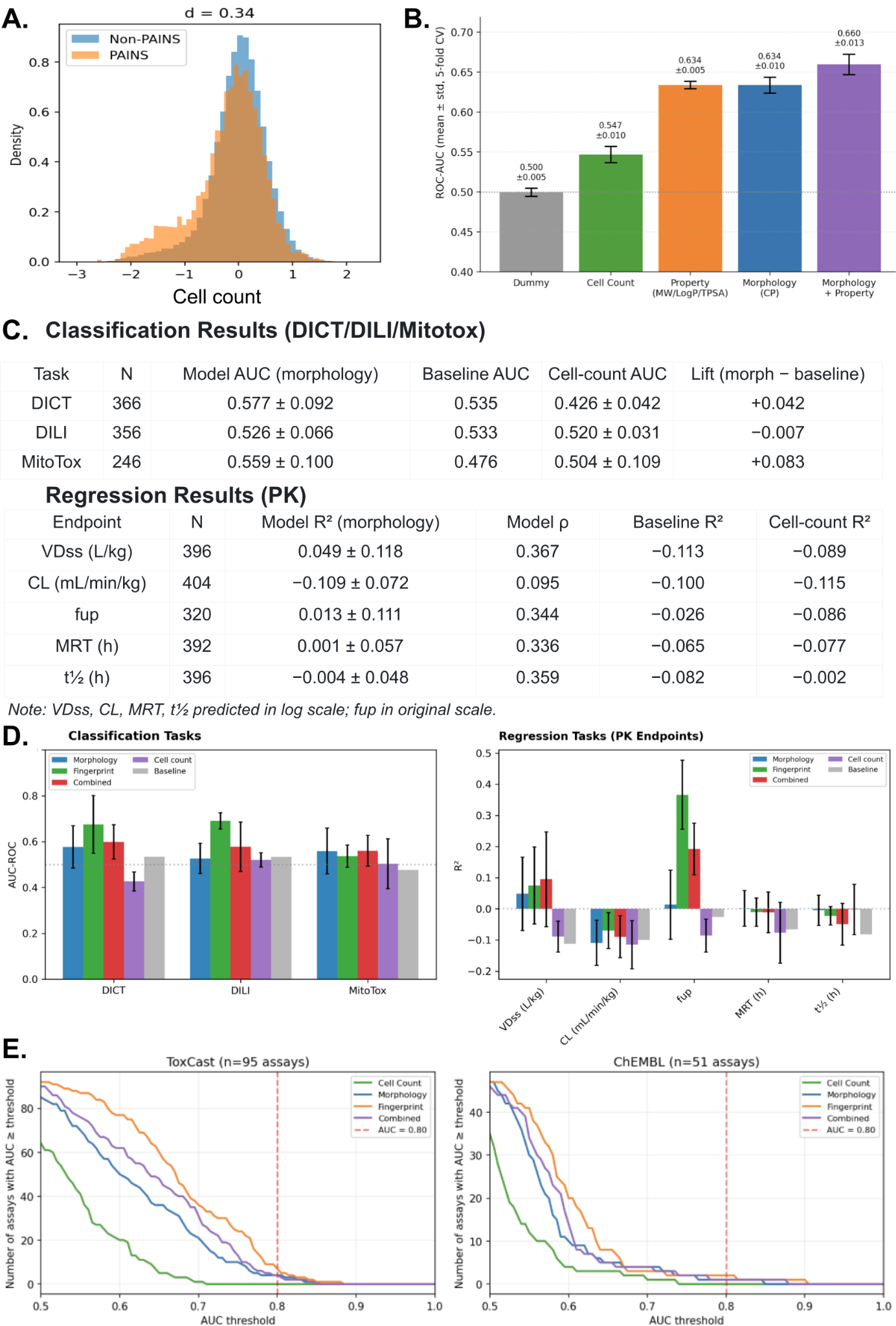

#### Supplementary Figure 15: Predicting chemical activities using JUMP Cell Painting profiles.

(A) Distribution of cell count across 112,524 compounds in the dataset for PAINS compounds (5,032 compounds flagged as PAINS by structural-alert matching against RDKit's built-in PAINS filter catalogue substructure set), and the 107,492 non-PAINS compounds. (B) AUC-ROC score for models predicting PAINS categories using cell count, physicochemical properties, or Cell Painting descriptors. PAINS prediction signal is not driven entirely by cytotoxicity. (C-D) Compounds were matched to JUMP by InChIKey connectivity (first 14 characters). Models are histogram gradient-boosting (HistGradientBoostingClassifier/Regressor; max_iter = 100, max_depth = 6) trained on Cell Painting morphology profiles, Morgan fingerprints (ECFP4, 2048-bit, radius 2), or their concatenation ("Combined"), under 5-fold (stratified, for classification) cross-validation with shared folds across feature sets (random_state = 42). Two reference baselines are shown: a label-shuffled stratified dummy classifier ("Baseline") and a model trained on the single median-cell-count feature ("Cell count"). Values are mean ± s.d. across folds. (C) Performance tables. Classification Results for each toxicity endpoint (DICT, drug-induced cardiotoxicity; DILI, drug-induced liver injury; MitoTox, mitochondrial toxicity): number of matched compounds (N), morphology model AUC, stratified-baseline AUC, cell-count-baseline AUC, and Lift (morphology − stratified baseline). Regression Results (PK) for each human pharmacokinetic endpoint, showing morphology model coefficient of determination (R²), Spearman rank correlation (ρ) between predicted and observed values, baseline and cell count R². VDss, CL, MRT and t½ were modeled in log scale; fup in original scale. (D) Performance bar charts. Classification shows cross-validated ROC-AUC for each feature set (Morphology, Fingerprint, Combined, Cell count) and the stratified baseline across the three toxicity endpoints; dotted line marks chance (AUC = 0.5). Regression Tasks (PK Endpoints) shows cross-validated R² for each feature set and the baseline across the five PK endpoints; dotted line marks R² = 0. Error bars represent ± s.d. (E) Binary activity predicted for ToxCast assays (n=95) and ChEMBL single-protein targets (n=51) using a histogram gradient-boosting classifier under 5-fold stratified cross-validation. Curves show, for each feature set, cell count (single feature), Cell Painting morphology, Morgan fingerprint (ECFP4, 2048-bit), and their concatenation. The number of assays whose mean cross-validated AUC exceeds each threshold is shown (dashed line: AUC = 0.80). The few ChEMBL endpoints with AUC greater than 0.80 were extremely class-imbalanced, and cell count alone also performed well on these tasks, suggesting that their apparent predictability may reflect systematic differences between the minority compounds and the remainder of the assayed set rather than target-specific biological signal.

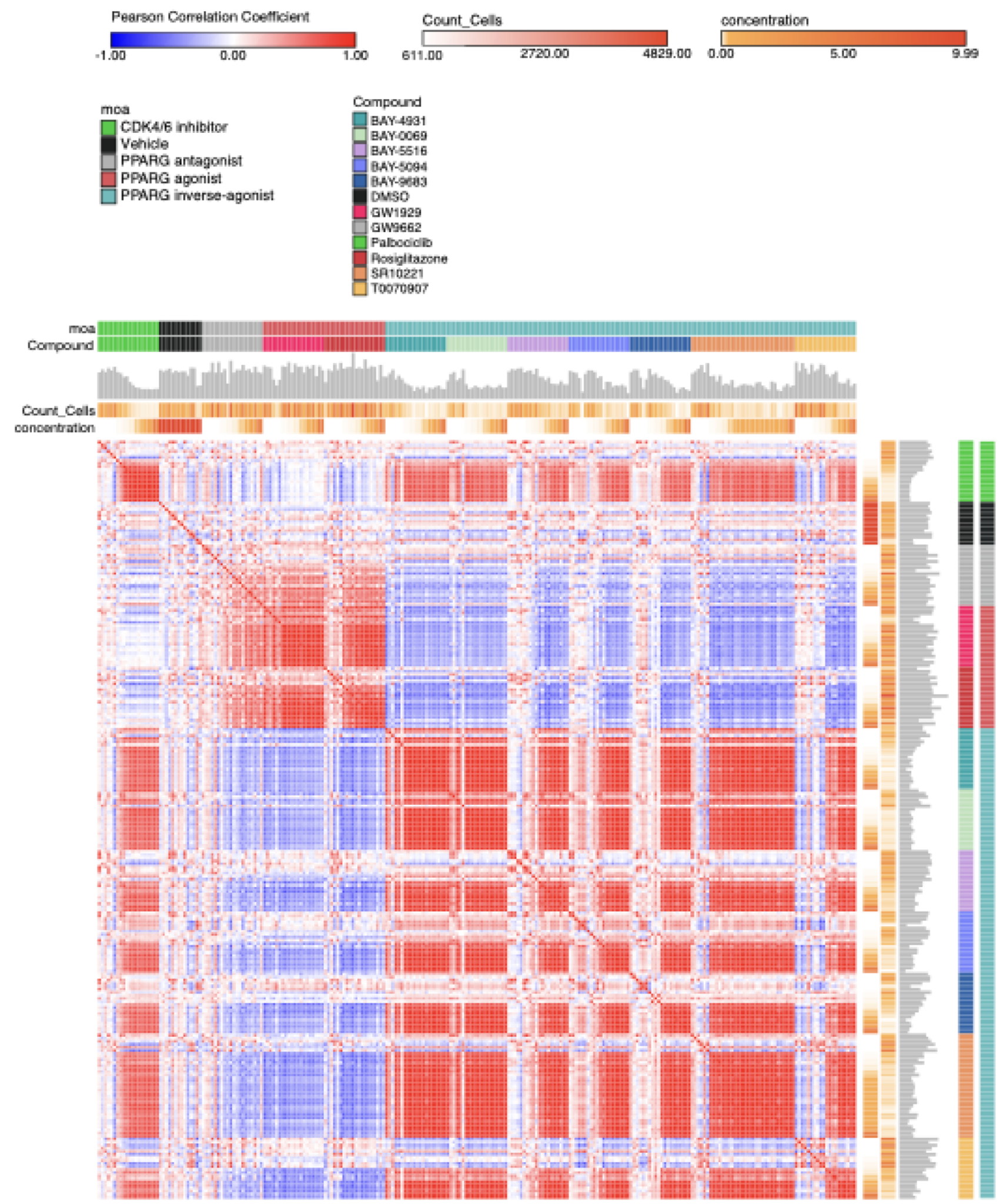

#### Supplementary Figure 16: Phenotypic effects of PPARG modulators are dose-dependent.

Pearson correlation coefficients were generated using the Similarity Matrix tool in Morpheus (Broad Institute, https://software.broadinstitute.org/morpheus/). Columns and rows are sorted by mechanism of action (moa), compound, and its concentration. Count_Cells indicates the cells counted at the end of treatment.

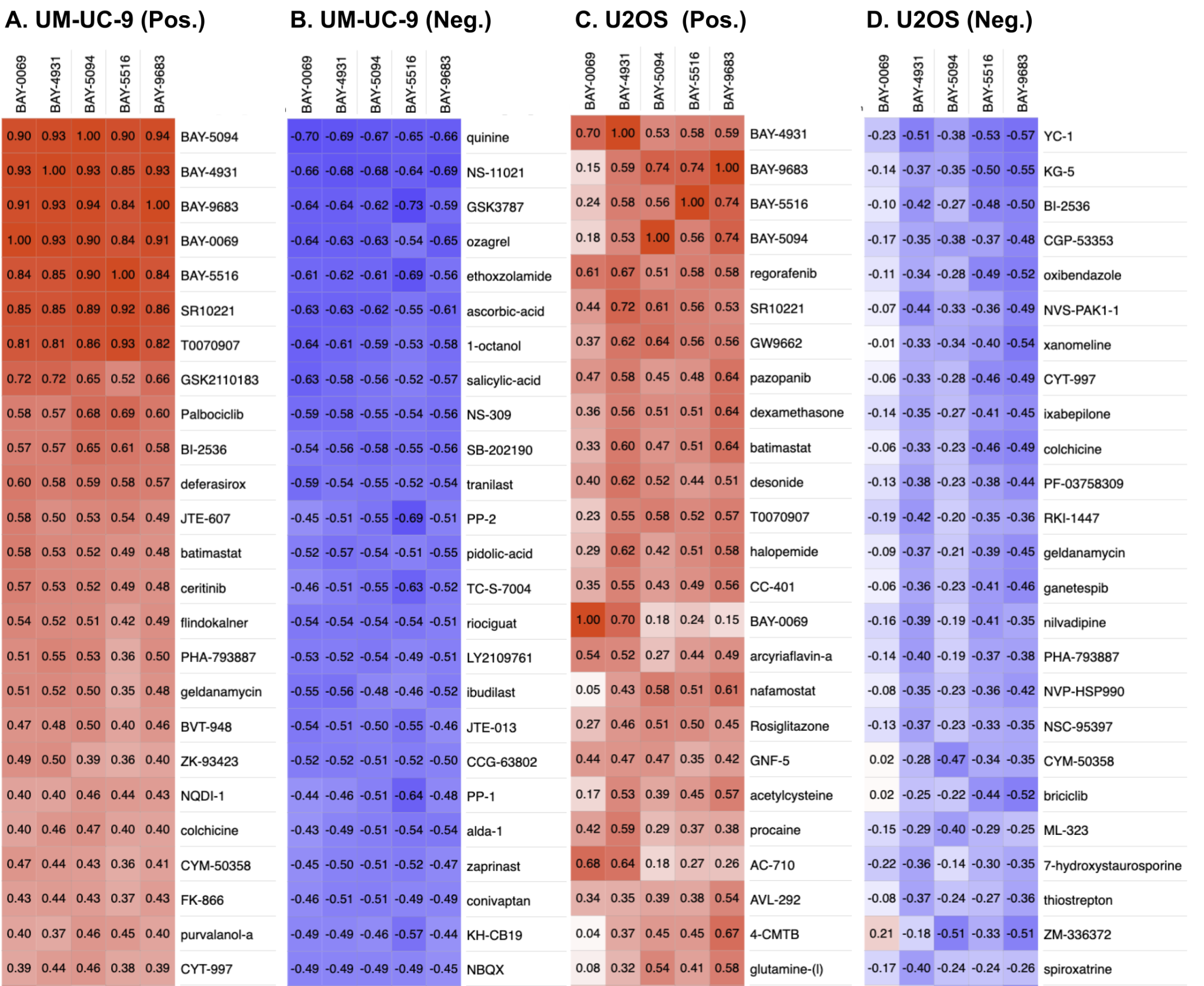

#### Supplementary Figure 17: Heatmap of BAY- compounds’ correlation to PPARG Cell Painting plate and Target-2 plate.

Correlation matrix of the top 30 compounds by mean similarity/anti-similarity (Pos. = positive Pearson correlation; Neg. = negative correlation). PPARG inverse agonists (A & B: UM-UC-9; C & D: U2OS). Palbociclib, purvalanol-a, and PHA-793887 are examples of compounds that target CDK activity and correlate with PPARG inverse agonists in UM-UC-9 (panel A). The BAY- compounds are represented by their max dose (2.5 μM). BAY query profiles are means of two wells per condition (n = 2 technical replicate wells); Target-2 reference profiles generally derive from a single well on the concurrently processed Target-2 plate.

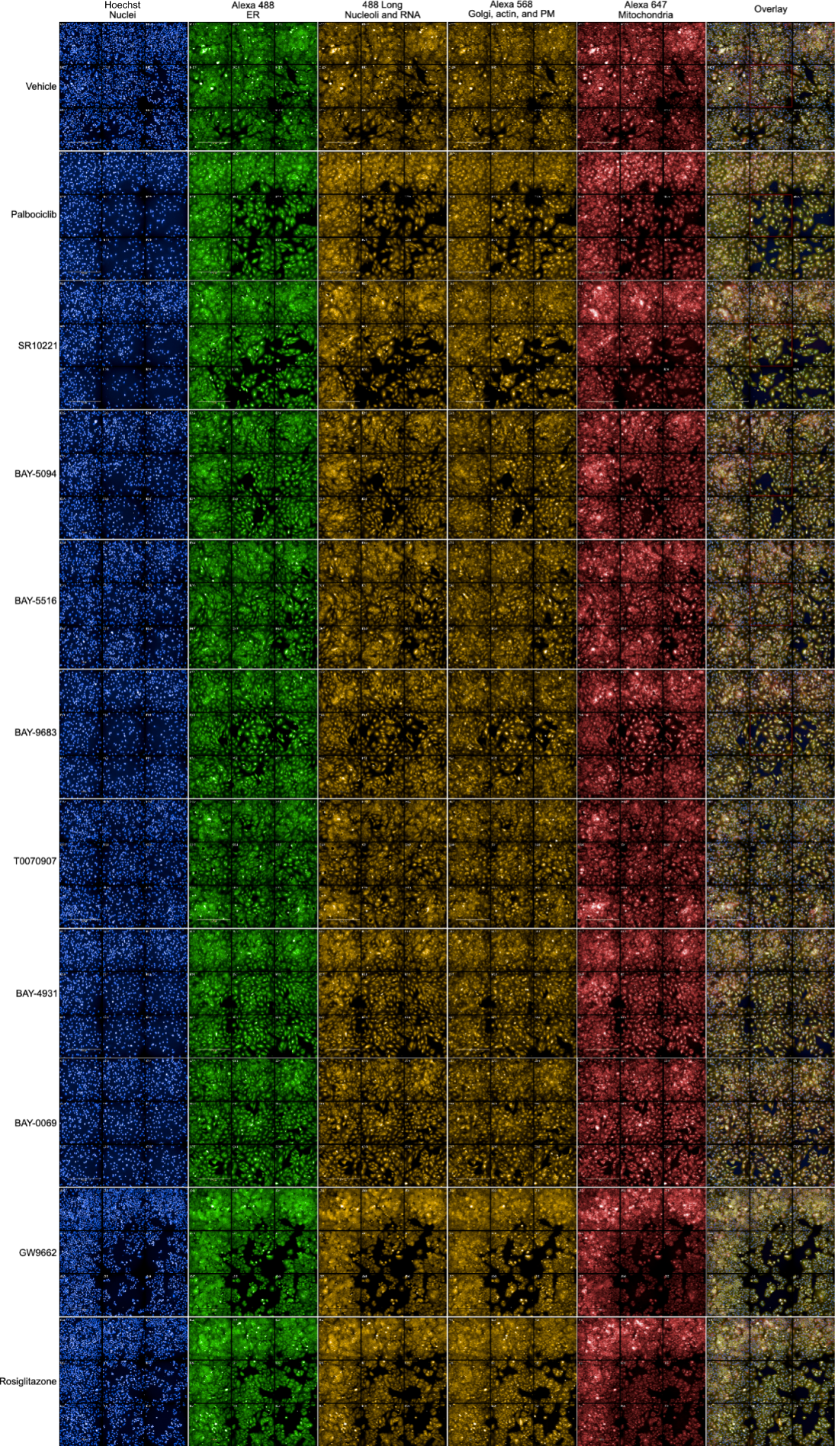

#### Supplementary Figure 18: Cell Painting images show that cells treated with PPARG inverse agonists adopt a senescent morphology with fewer, more dispersed cells.

Fluorescent images from 5 separate channels and full merged overlay of UM-UC-9 cells treated with indicated compounds at 900 nM (or vehicle control) for 72 hours. Channels are blue: Hoechst 33342 (nuclear DNA), Alexa 488 green: concanavalin A (endoplasmic reticulum), 488 Long yellow: PhenoVue 512 Nucleic Acid (nucleoli and cytoplasmic RNA), Alexa 568 yellow: phalloidin (actin) and WGA (Golgi and plasma membrane), and Alexa 647 red: MitoTracker Deep Red (mitochondria). Scale bars shown at 500 μm in bottom left image of most panels of 9 fields of view. Higher-resolution images are shown in Figure 4B.

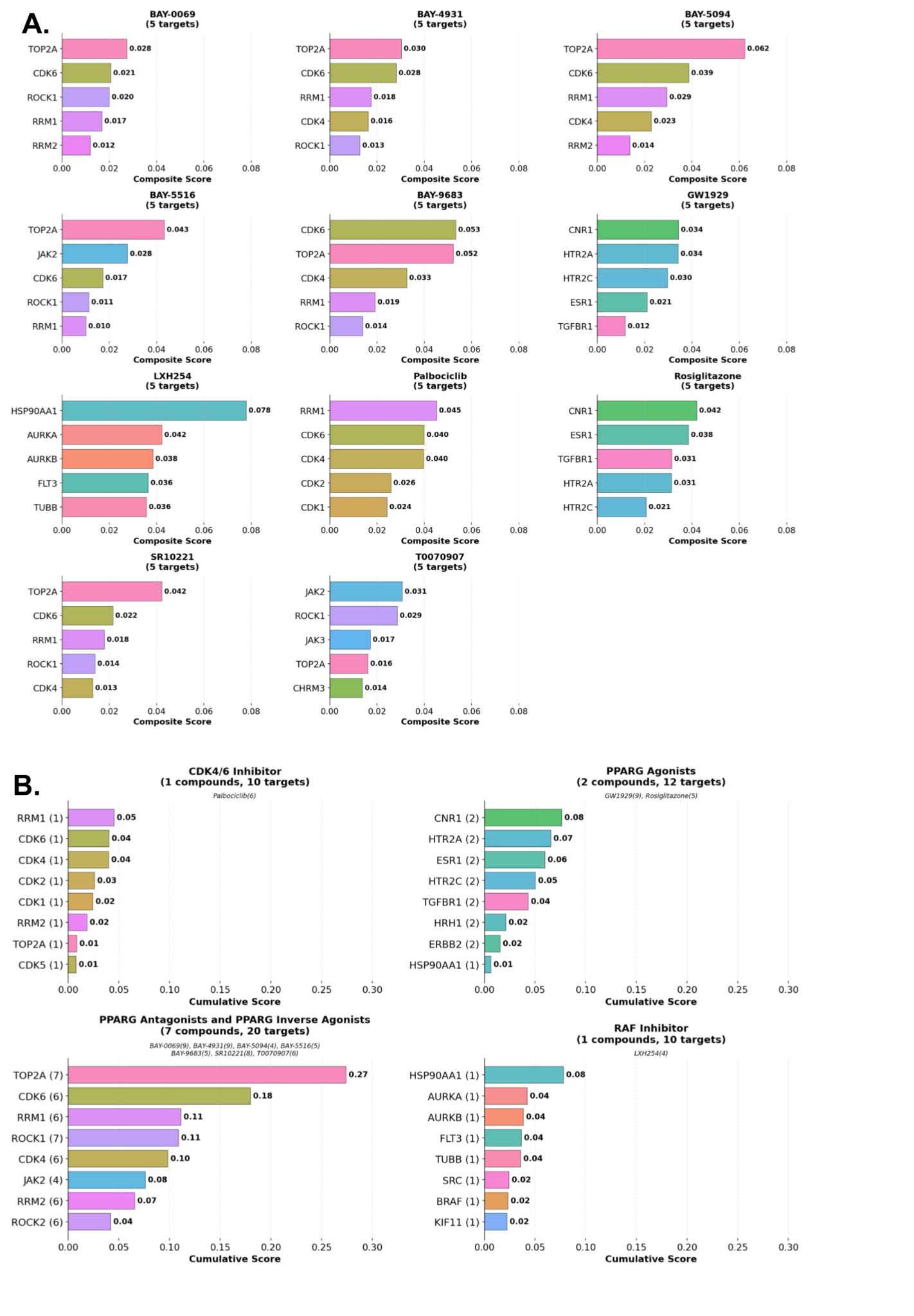

#### **Supplementary Figure** 19**: Top-ranked targets for compounds and compound groups.**

(A) Annotated targets for the compounds from the JUMP dataset that best match key compounds on the PPARG Cell Painting plate, including the novel PPARG inverse agonists. (B) Summarized top matching targets across compounds for each compound group, showing the cumulative target average precision (AP) score for each compound group.

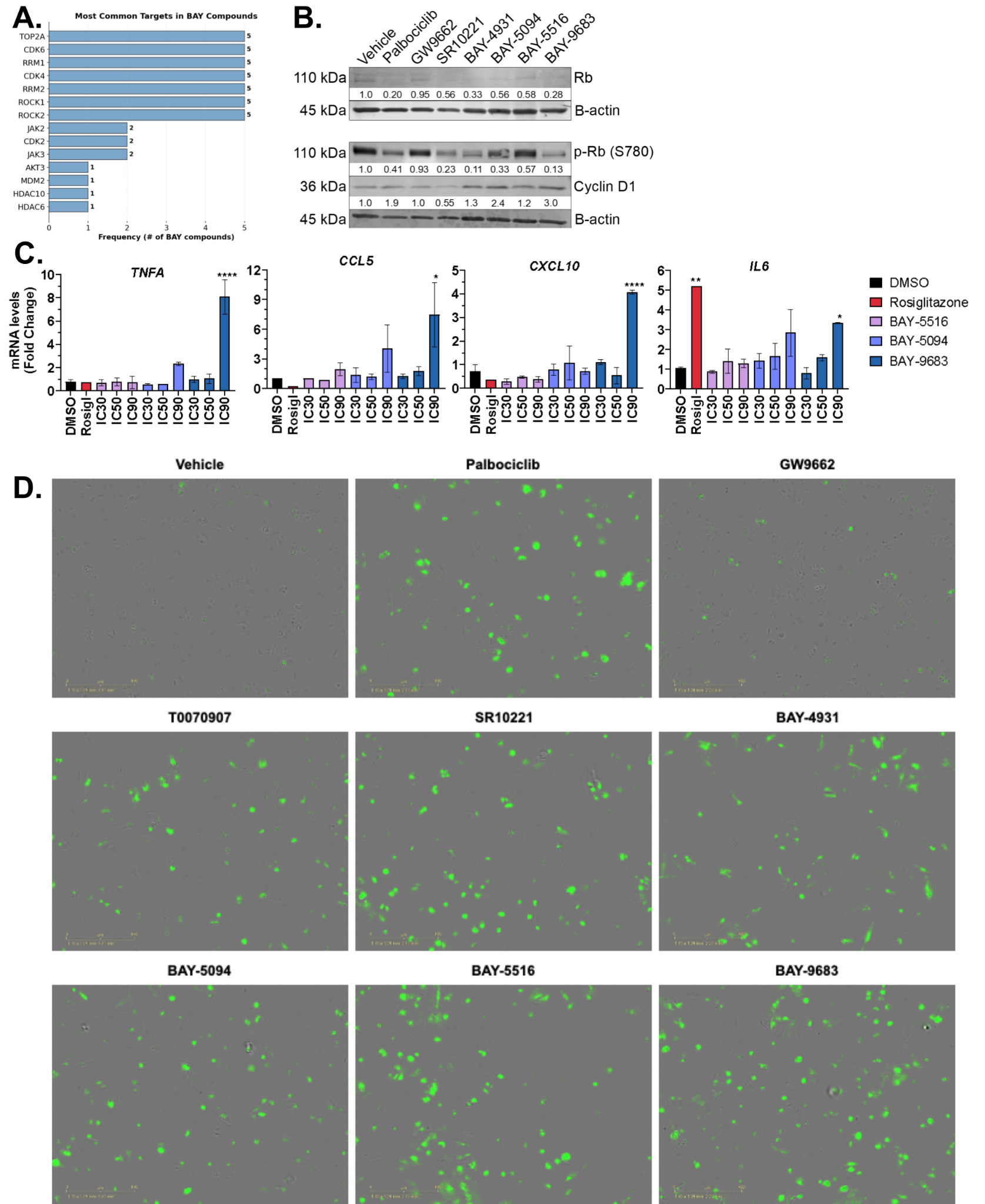

#### Supplementary Figure 20: PPARG inverse agonists produce morphology profiles similar to drugs targeting CDKs, RRM, TOP2A and induce senescence.

**(**A) Annotated targets are shown for the compounds from the JUMP dataset that best match to the set of five query BAY- compounds in UM-UC-9 cells (absolute counts as compared to the composite score shown in Figure 4C). See Methods for details. (B) Western blot analysis of cell lysates after 8 days of treatment at 500 nM revealed a marked reduction in phosphorylated Retinoblastoma Protein (p-Rb) (S780) levels in cells treated with palbociclib, SR10221, BAY-4931, BAY-9683, and BAY-5094 although not with BAY-5516, consistent with the G1 arrest observed by flow cytometry. (C) Gene expression analysis by RT-qPCR for senescence-associated secretory phenotype (SASP) genes TNF (TNFA), CCL5, CXCL10 and IL6. UM-UC-9 cells were treated for 5 days with indicated compounds at their anti-proliferative IC30, IC50, or IC90 doses (BAY-5516 at 13nM, 18nM, and 39nM; BAY-5094 at 8nM, 25nM, and 483nM; BAY-9683 at 42nM, 98nM, and 872nM, rosiglitazone at 100nM). The relative mRNA expression expressed as fold change is shown for 2 independent experiments ± S.E.M. Statistical analysis by one-way ANOVA with Dunnett's multiple comparisons test. * p<0.05, ** p<0.01, *** p<0.001, **** p<0.0001. An increase in expression was only statistically significant for BAY-9683. (D) PPARG inverse agonists lead to senescence-associated β-galactosidase activity and phenocopy palbociclib. Fluorescence images of UM-UC-9 cells that were treated with indicated compounds at 500nM for 8 days to induce senescence, followed by fixation and staining for senescence marker β-galactosidase activity. Cells were passaged after 4 days based on normalization to vehicle to maintain adequate space for growth and equivalent cell counts for each treatment condition. Images were captured using IncuCyte S3 Essen Bioscience/Sartorius.

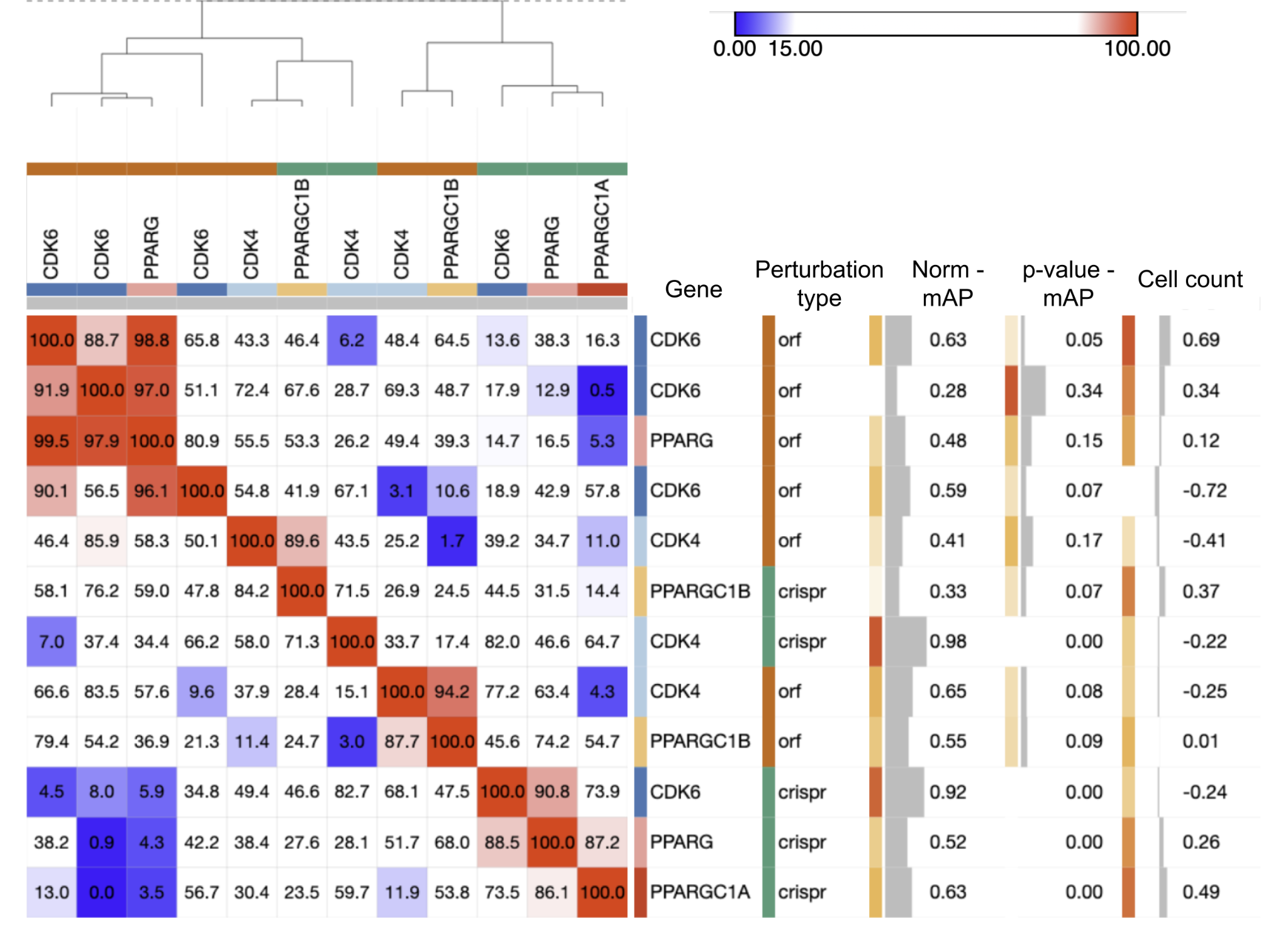

#### Supplementary Figure 21: Similarities between CDK4/6 and PPARG gene family member perturbations in the JUMP Cell Painting ORF/CRISPR datasets.

Clustermap showing the percentile rank of similarity between each gene-gene combination, reflected in a red/blue colormap (red = most similar; blue = least similar). Some genes had multiple reagents, which are listed separately and not combined. The numerical value shows the relative percentile matching score for each of the selected gene-reagent of the columns against all other gene-reagents in JUMP-ORF/CRISPR set, filtered to only show the gene-reagents in the CDK4/6 and PPARG* set. Other data shown for each gene include the normalized mean average precision (mAP) and its associated p-value for each gene; this is a measure of phenotypic activity - that is, how distinguishable is each gene-reagent from negative controls per the Cell Painting profiles (higher = stronger phenotype). Cell count is z-scored so it represents the number of standard deviations above/below the mean of negative control.
