## Supplementary_File_PHENOTYPIC_CONSISTENCY for "JUMP Cell Painting chemical dataset: Morphological map of 115,796 compounds in human cells"

### chemical probes

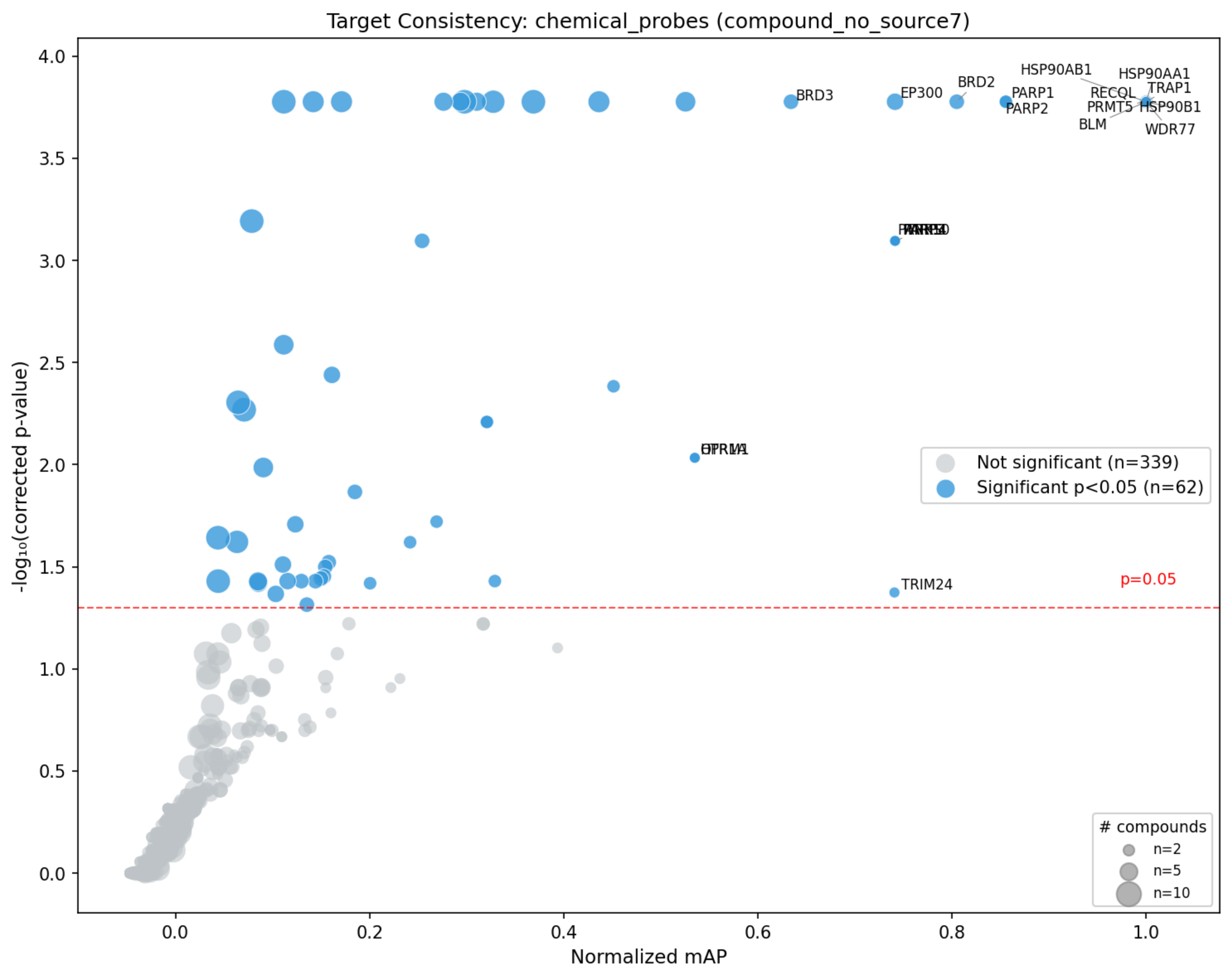
Target Consistency: chemical_probes (compound_no_source7)
-log₁₀(corrected p-value)
Normalized mAP
BRD3
EP300
BRD2
HSP90AB1
HSP90AA1
PARP1
RECQL
TRAP1
PARP2
PRMT5
HSP90B1
BLM
WDR77
TRIM24
p=0.05
Not significant (n=339)
Significant p<0.05 (n=62)
### compounds
n=2
n=5
n=10

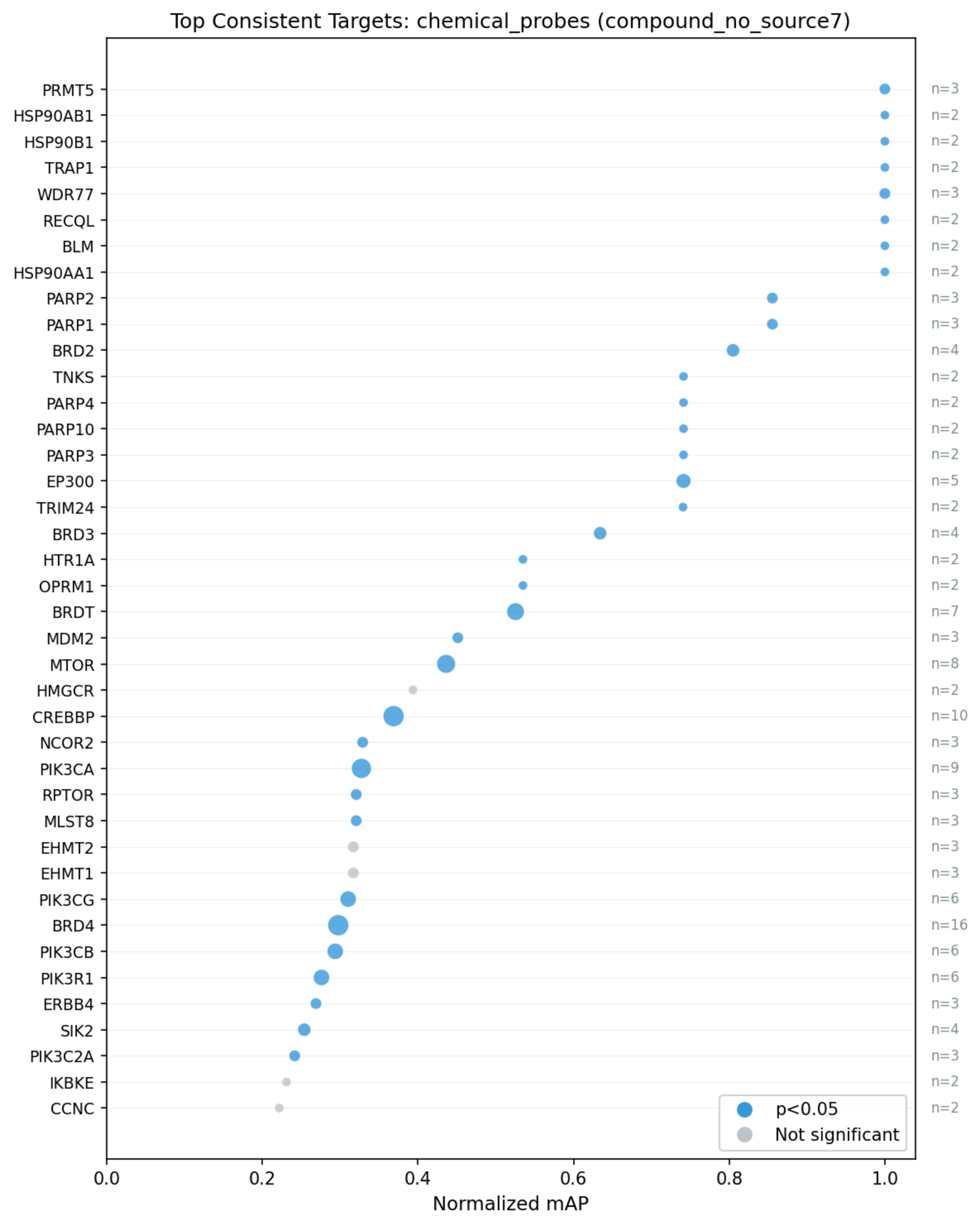

Top Consistent Targets: chemical_probes (compound_no_source7)
PRMT5
HSP90AB1
HSP90B1
TRAP1
WDR77
RECQL
BLM
HSP90AA1
PARP2
PARP1
BRD2
TNKS
PARP4
PARP10
PARP3
EP300
TRIM24
BRD3
HTR1A
OPRM1
BRDT
MDM2
MTOR
HMGCR
CREBBP
NCOR2
PIK3CA
RPTOR
MLST8
EHMT2
EHMT1
PIK3CG
BRD4
PIK3CB
PIK3R1
ERBB4
SIK2
PIK3C2A
IKBKE
CCNC
n=3
n=2
n=2
n=2
n=3
n=2
n=2
n=2
n=3
n=3
n=4
n=2
n=2
n=2
n=2
n=5
n=2
n=4
n=2
n=2
n=7
n=3
n=8
n=2
n=10
n=3
n=9
n=3
n=3
n=3
n=3
n=6
n=16
n=6
n=6
n=3
n=4
n=3
n=2
n=2
p<0.05
Not significant
0.0
0.2
0.4
0.6
0.8
1.0
Normalized mAP

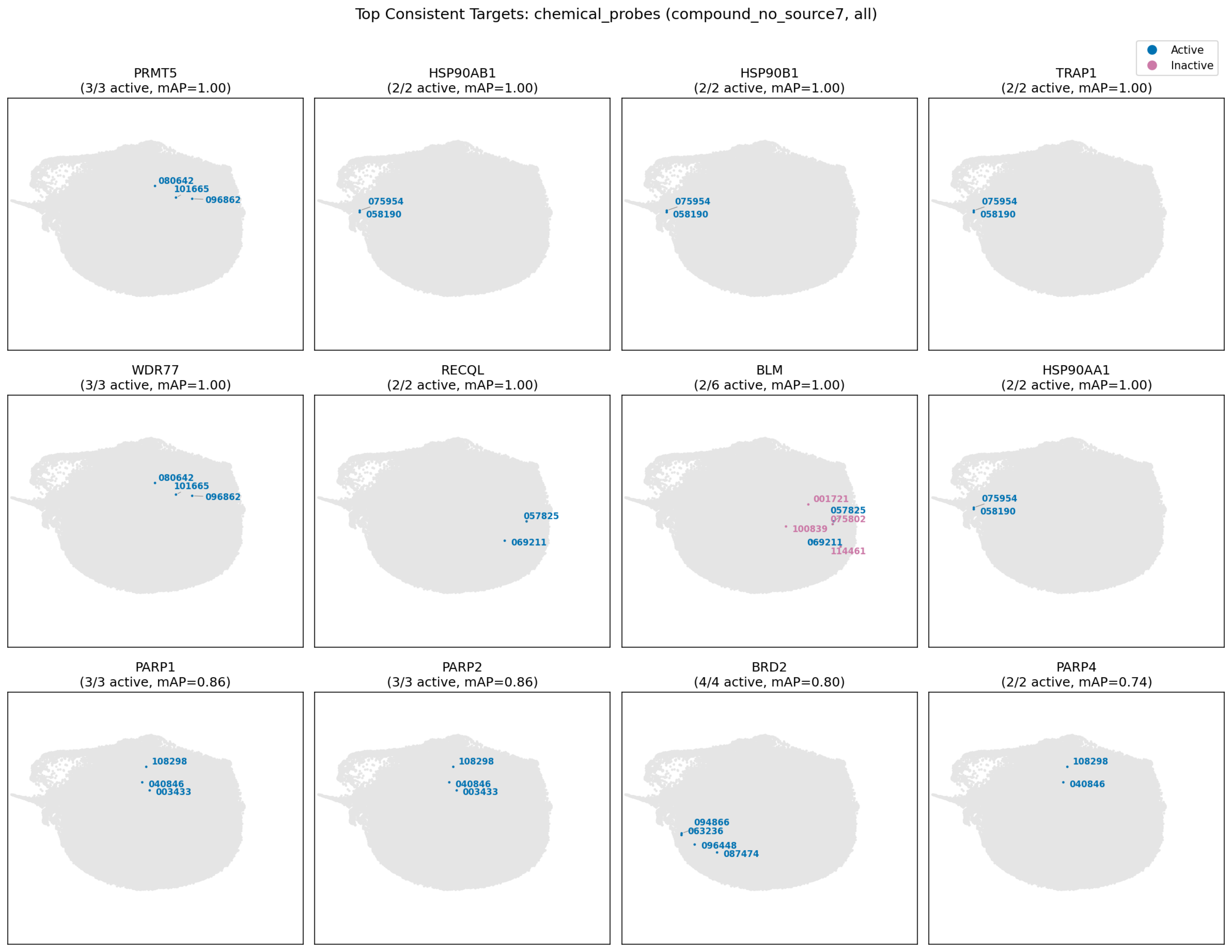
Top Consistent Targets: chemical_probes (compound_no_source7, all)
Active
Inactive
PRMT5
(3/3 active, mAP=1.00)
080642
101665
096862
HSP90AB1
(2/2 active, mAP=1.00)
075954
058190
HSP90B1
(2/2 active, mAP=1.00)
075954
058190
TRAP1
(2/2 active, mAP=1.00)
075954
058190
WDR77
(3/3 active, mAP=1.00)
080642
101665
096862
RECQL
(2/2 active, mAP=1.00)
057825
069211
BLM
(2/6 active, mAP=1.00)
001721
057825
075802
100839
069211
114461
HSP90AA1
(2/2 active, mAP=1.00)
075954
058190
PARP1
(3/3 active, mAP=0.86)
108298
040846
003433
PARP2
(3/3 active, mAP=0.86)
108298
040846
003433
BRD2
(4/4 active, mAP=0.80)
094866
063236
096448
087474
PARP4
(2/2 active, mAP=0.74)
108298
040846

Top Consistent Targets: chemical_probes (compound_no_source7, active)

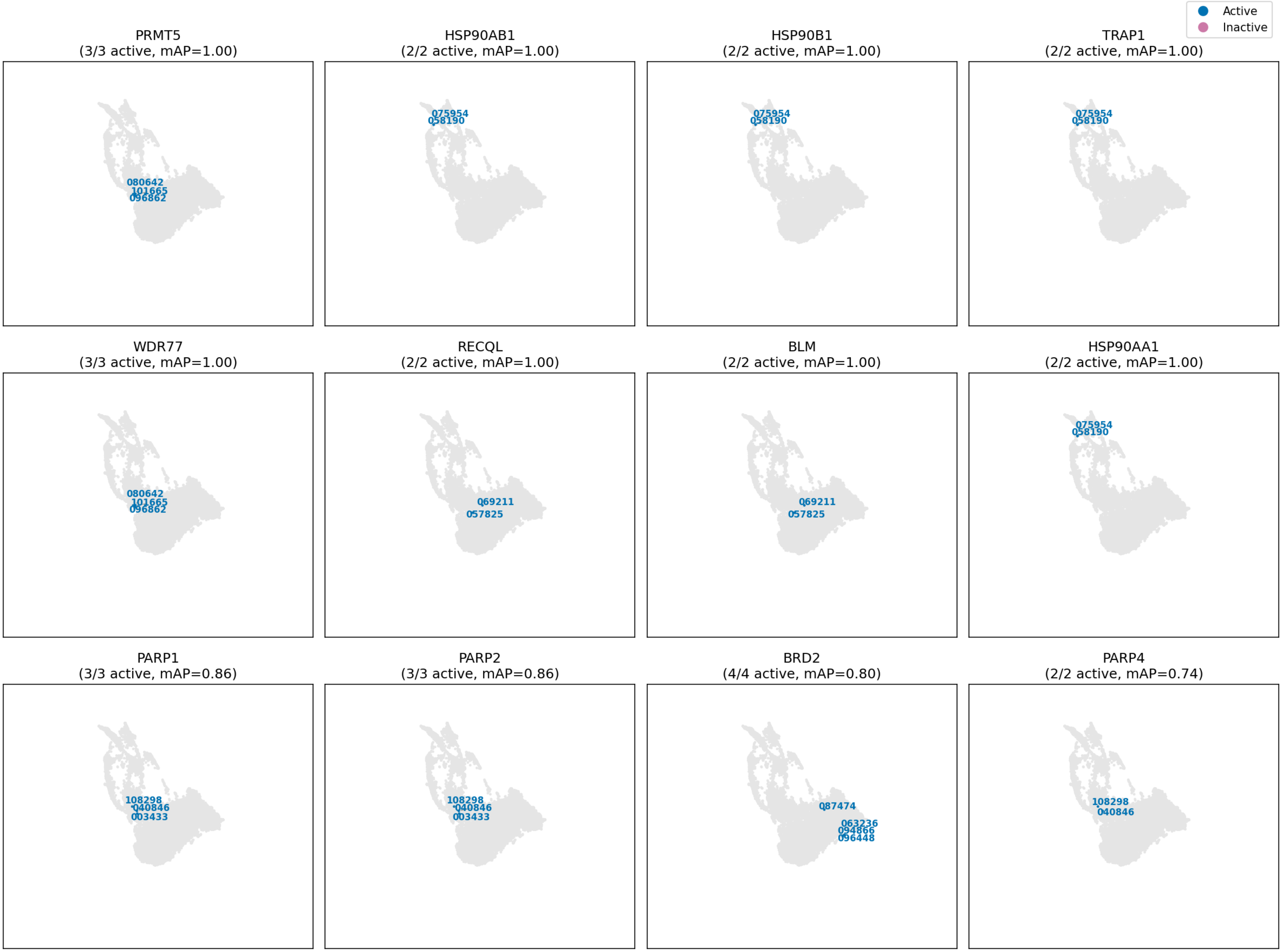

disease area

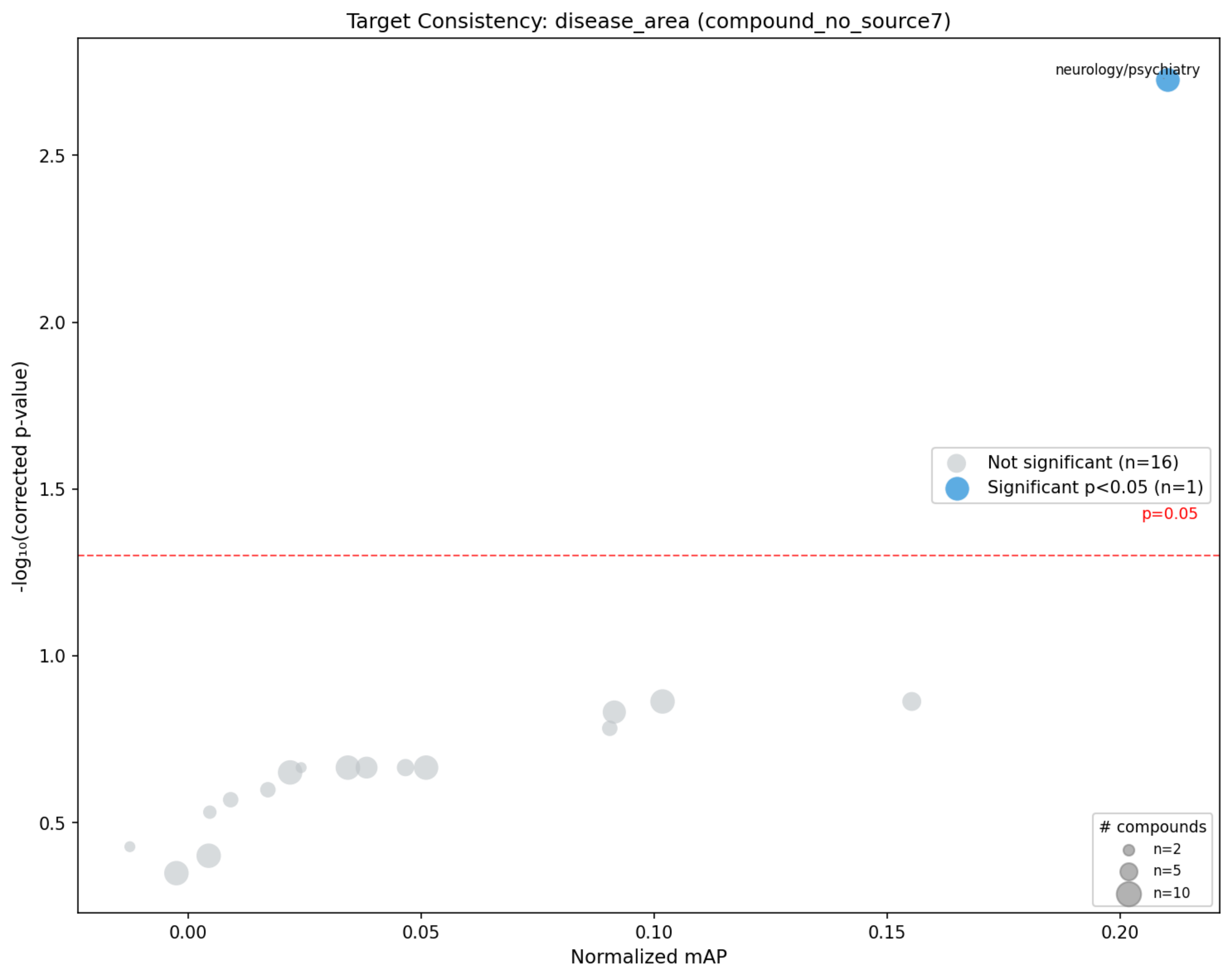

Target Consistency: disease_area (compound_no_source7)
neurology/psychiatry
Not significant (n=16)
Significant p<0.05 (n=1)
p=0.05
### compounds
n=2
n=5
n=10
-log₁₀(corrected p-value)
Normalized mAP
0.5
1.0
1.5
2.0
2.5
0.00
0.05
0.10
0.15
0.20

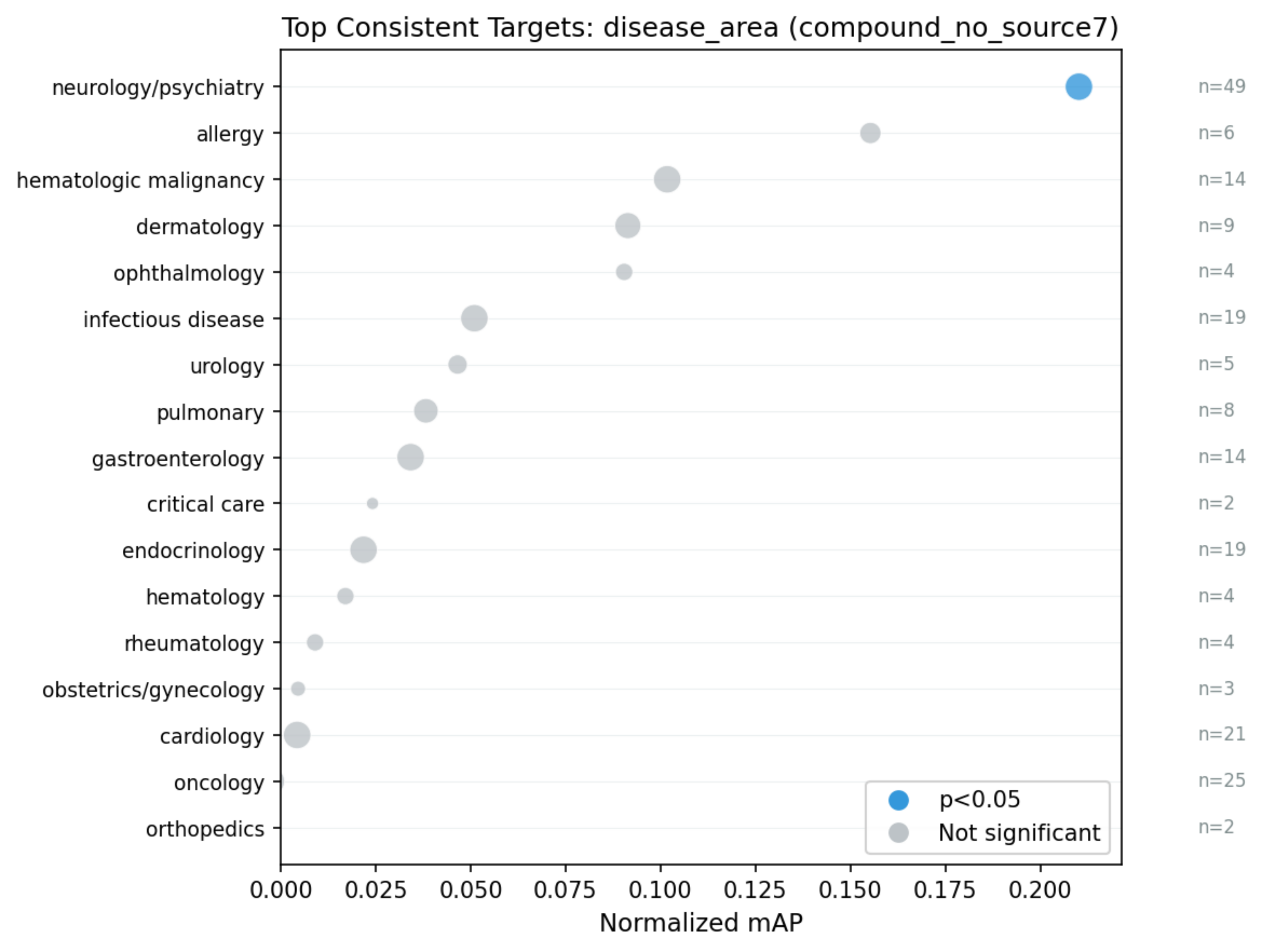

Top Consistent Targets: disease_area (compound_no_source7)
neurology/psychiatry
allergy
hematologic malignancy
dermatology
ophthalmology
infectious disease
urology
pulmonary
gastroenterology
critical care
endocrinology
hematology
rheumatology
obstetrics/gynecology
cardiology
oncology
orthopedics
n=49
n=6
n=14
n=9
n=4
n=19
n=5
n=8
n=14
n=2
n=19
n=4
n=4
n=3
n=21
n=25
n=2
p<0.05
Not significant
0.000
0.025
0.050
0.075
0.100
0.125
0.150
0.175
0.200
Normalized mAP

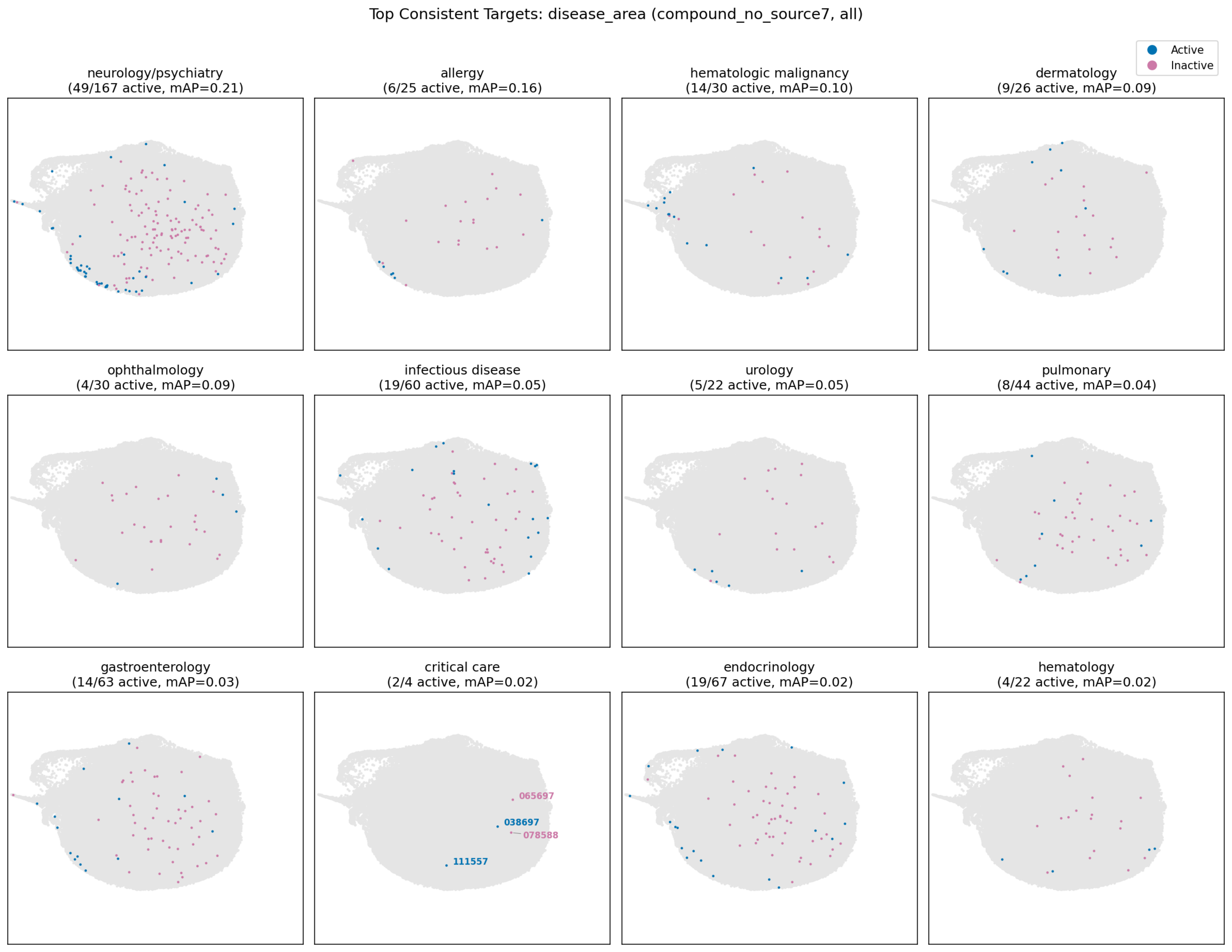
Top Consistent Targets: disease_area (compound_no_source7, all)
Active
Inactive
neurology/psychiatry
(49/167 active, mAP=0.21)
allergy
(6/25 active, mAP=0.16)
hematologic malignancy
(14/30 active, mAP=0.10)
dermatology
(9/26 active, mAP=0.09)
ophthalmology
(4/30 active, mAP=0.09)
infectious disease
(19/60 active, mAP=0.05)
urology
(5/22 active, mAP=0.05)
pulmonary
(8/44 active, mAP=0.04)
gastroenterology
(14/63 active, mAP=0.03)
critical care
(2/4 active, mAP=0.02)
065697
038697
078588
111557
endocrinology
(19/67 active, mAP=0.02)
hematology
(4/22 active, mAP=0.02)

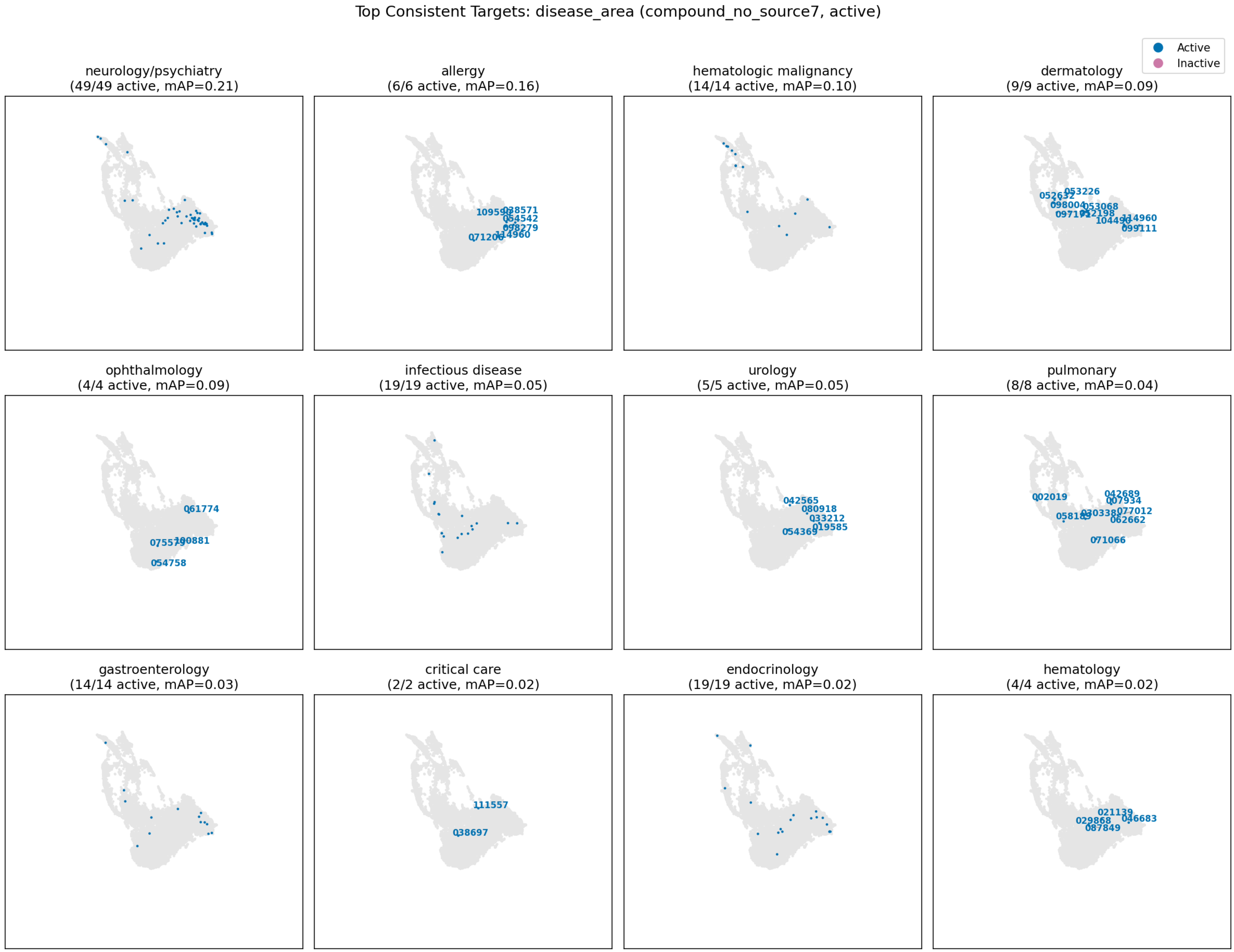

Top Consistent Targets: disease_area (compound_no_source7, active)
Active
Inactive
neurology/psychiatry
(49/49 active, mAP=0.21)
allergy
(6/6 active, mAP=0.16)
038571
054542
098279
114960
071206
hematologic malignancy
(14/14 active, mAP=0.10)
dermatology
(9/9 active, mAP=0.09)
053226
098004
053068
114960
099111
ophthalmology
(4/4 active, mAP=0.09)
061774
100881
054758
infectious disease
(19/19 active, mAP=0.05)
urology
(5/5 active, mAP=0.05)
042565
080918
033212
019585
054369
pulmonary
(8/8 active, mAP=0.04)
002019
042689
007934
077012
062662
071066
gastroenterology
(14/14 active, mAP=0.03)
critical care
(2/2 active, mAP=0.02)
111557
038697
endocrinology
(19/19 active, mAP=0.02)
hematology
(4/4 active, mAP=0.02)
021139
029868
046683
087849

moa

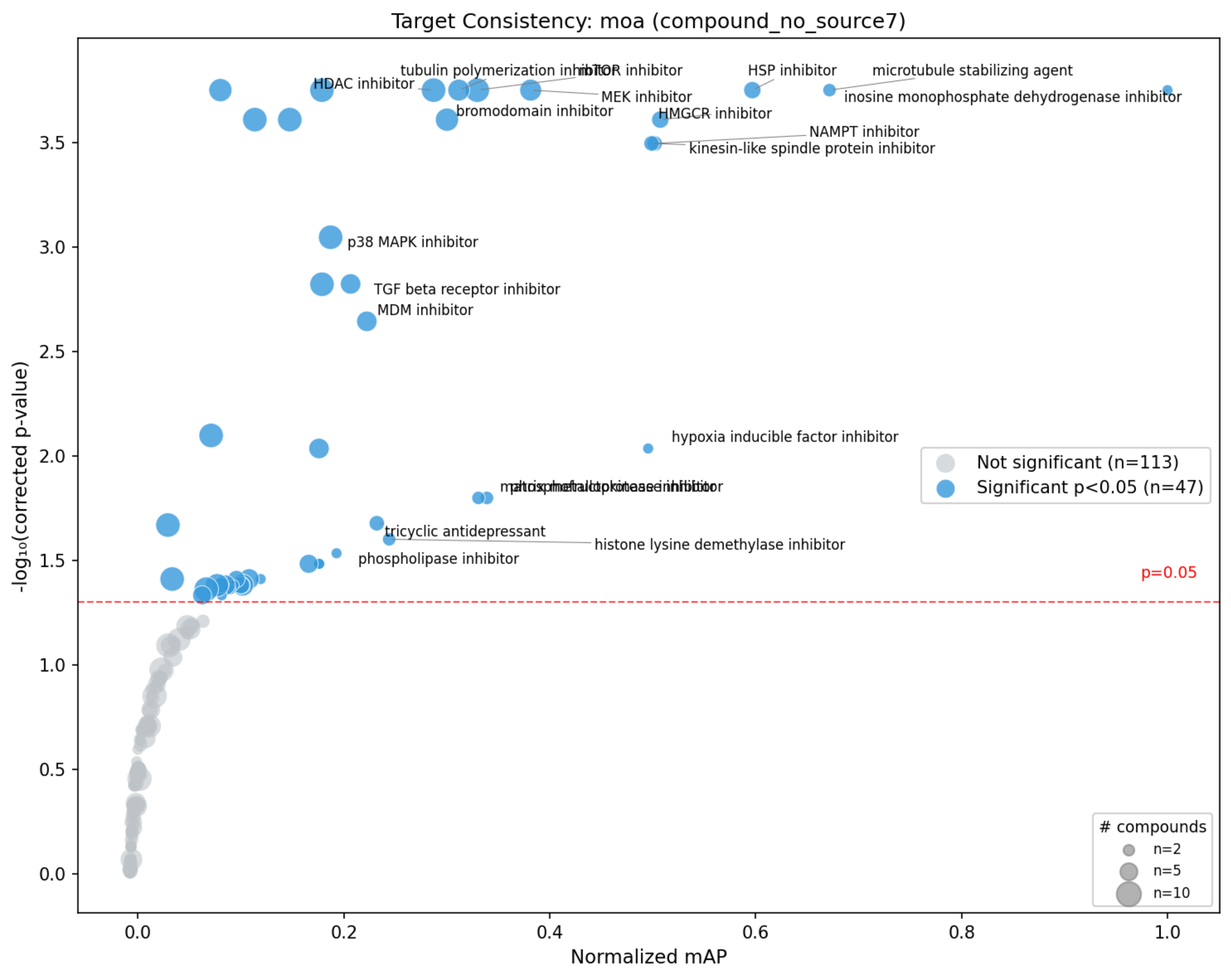

Target Consistency: moa (compound_no_source7)
-log₁₀(corrected p-value)
Normalized mAP
HDAC inhibitor
tubulin polymerization inhibitor
mTOR inhibitor
MEK inhibitor
bromodomain inhibitor
HSP inhibitor
microtubule stabilizing agent
inosine monophosphate dehydrogenase inhibitor
HMGCR inhibitor
NAMPT inhibitor
kinesin-like spindle protein inhibitor
p38 MAPK inhibitor
TGF beta receptor inhibitor
MDM inhibitor
hypoxia inducible factor inhibitor
tricyclic antidepressant
histone lysine demethylase inhibitor
phospholipase inhibitor
p=0.05
Not significant (n=113)
Significant p<0.05 (n=47)
### compounds
n=2
n=5
n=10
0.0
0.2
0.4
0.6
0.8
1.0
0.5
1.0
1.5
2.0
2.5
3.0
3.5

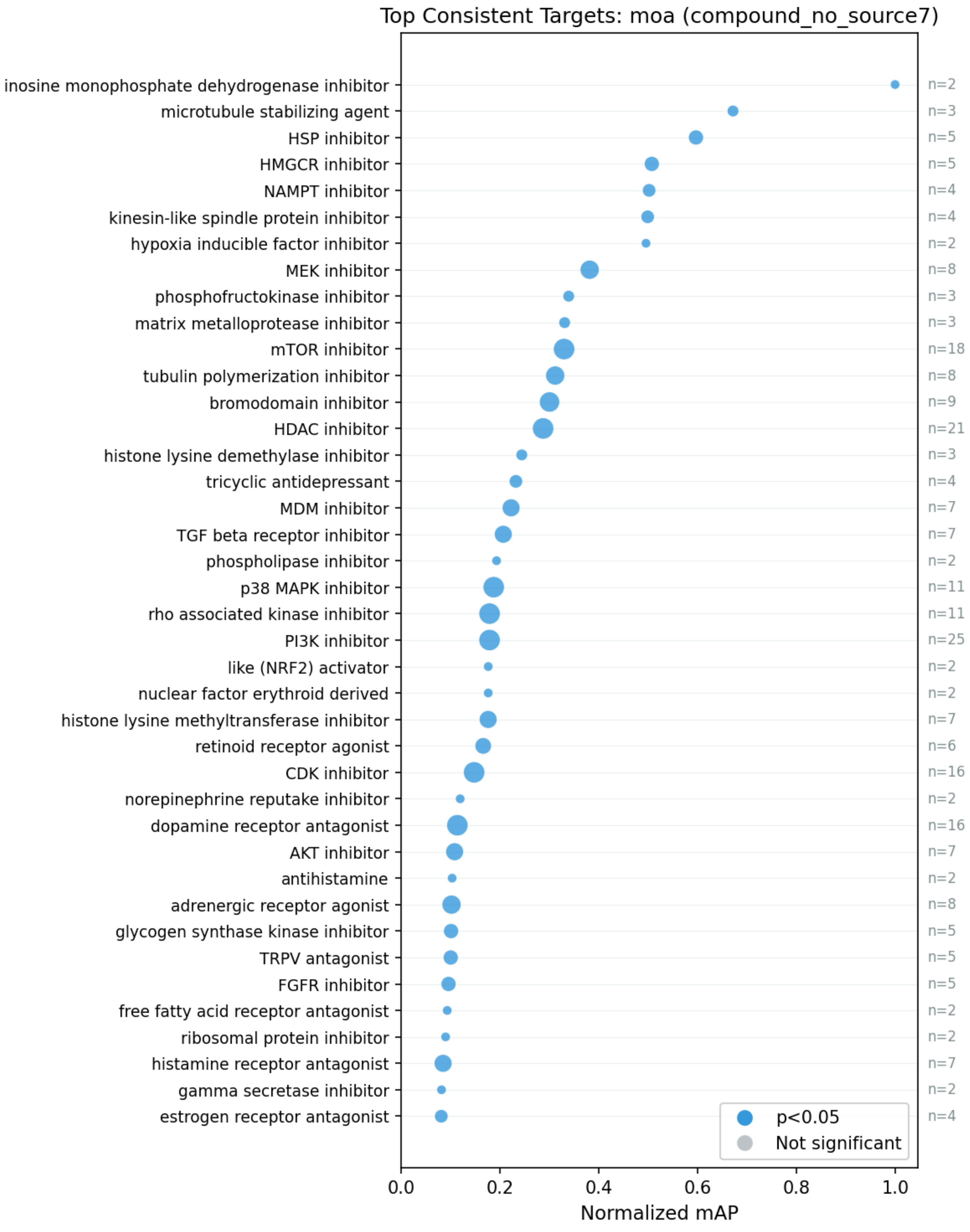

Top Consistent Targets: moa (compound_no_source7)
inosine monophosphate dehydrogenase inhibitor
microtubule stabilizing agent
HSP inhibitor
HMGCR inhibitor
NAMPT inhibitor
kinesin-like spindle protein inhibitor
hypoxia inducible factor inhibitor
MEK inhibitor
phosphofructokinase inhibitor
matrix metalloprotease inhibitor
mTOR inhibitor
tubulin polymerization inhibitor
bromodomain inhibitor
HDAC inhibitor
histone lysine demethylase inhibitor
tricyclic antidepressant
MDM inhibitor
TGF beta receptor inhibitor
phospholipase inhibitor
p38 MAPK inhibitor
rho associated kinase inhibitor
PI3K inhibitor
like (NRF2) activator
nuclear factor erythroid derived
histone lysine methyltransferase inhibitor
retinoid receptor agonist
CDK inhibitor
norepinephrine reputake inhibitor
dopamine receptor antagonist
AKT inhibitor
antihistamine
adrenergic receptor agonist
glycogen synthase kinase inhibitor
TRPV antagonist
FGFR inhibitor
free fatty acid receptor antagonist
ribosomal protein inhibitor
histamine receptor antagonist
gamma secretase inhibitor
estrogen receptor antagonist
n=2
n=3
n=5
n=5
n=4
n=4
n=2
n=8
n=3
n=3
n=18
n=8
n=9
n=21
n=3
n=4
n=7
n=7
n=2
n=11
n=11
n=25
n=2
n=2
n=7
n=6
n=16
n=2
n=16
n=7
n=2
n=8
n=5
n=5
n=5
n=2
n=2
n=7
n=2
n=4
p<0.05
Not significant
0.0
0.2
0.4
0.6
0.8
1.0
Normalized mAP

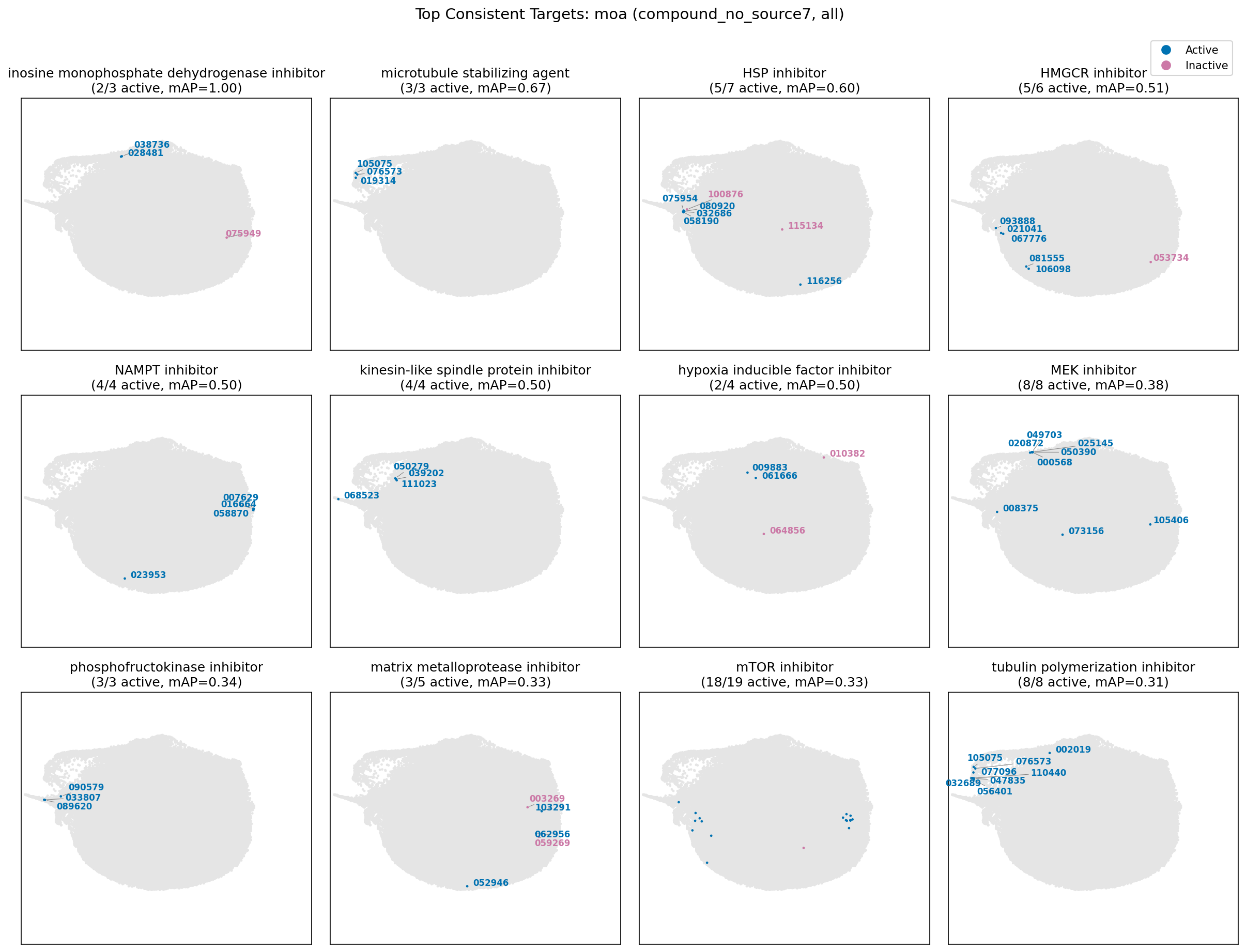

Top Consistent Targets: moa (compound_no_source7, all)
Active
Inactive
inosine monophosphate dehydrogenase inhibitor
(2/3 active, mAP=1.00)
038736
028481
075949
microtubule stabilizing agent
(3/3 active, mAP=0.67)
105075
076573
019314
HSP inhibitor
(5/7 active, mAP=0.60)
100876
075954
080920
032686
058190
115134
116256
HMGCR inhibitor
(5/6 active, mAP=0.51)
093888
021041
067776
081555
106098
053734
NAMPT inhibitor
(4/4 active, mAP=0.50)
007629
016664
058870
023953
kinesin-like spindle protein inhibitor
(4/4 active, mAP=0.50)
050279
039202
111023
068523
hypoxia inducible factor inhibitor
(2/4 active, mAP=0.50)
010382
009883
061666
064856
MEK inhibitor
(8/8 active, mAP=0.38)
049703
020872
025145
050390
000568
008375
105406
073156
phosphofructokinase inhibitor
(3/3 active, mAP=0.34)
090579
033807
089620
matrix metalloprotease inhibitor
(3/5 active, mAP=0.33)
003269
103291
062956
059269
052946
mTOR inhibitor
(18/19 active, mAP=0.33)
tubulin polymerization inhibitor
(8/8 active, mAP=0.31)
002019
105075
076573
077096
110440
032689
047835
056401

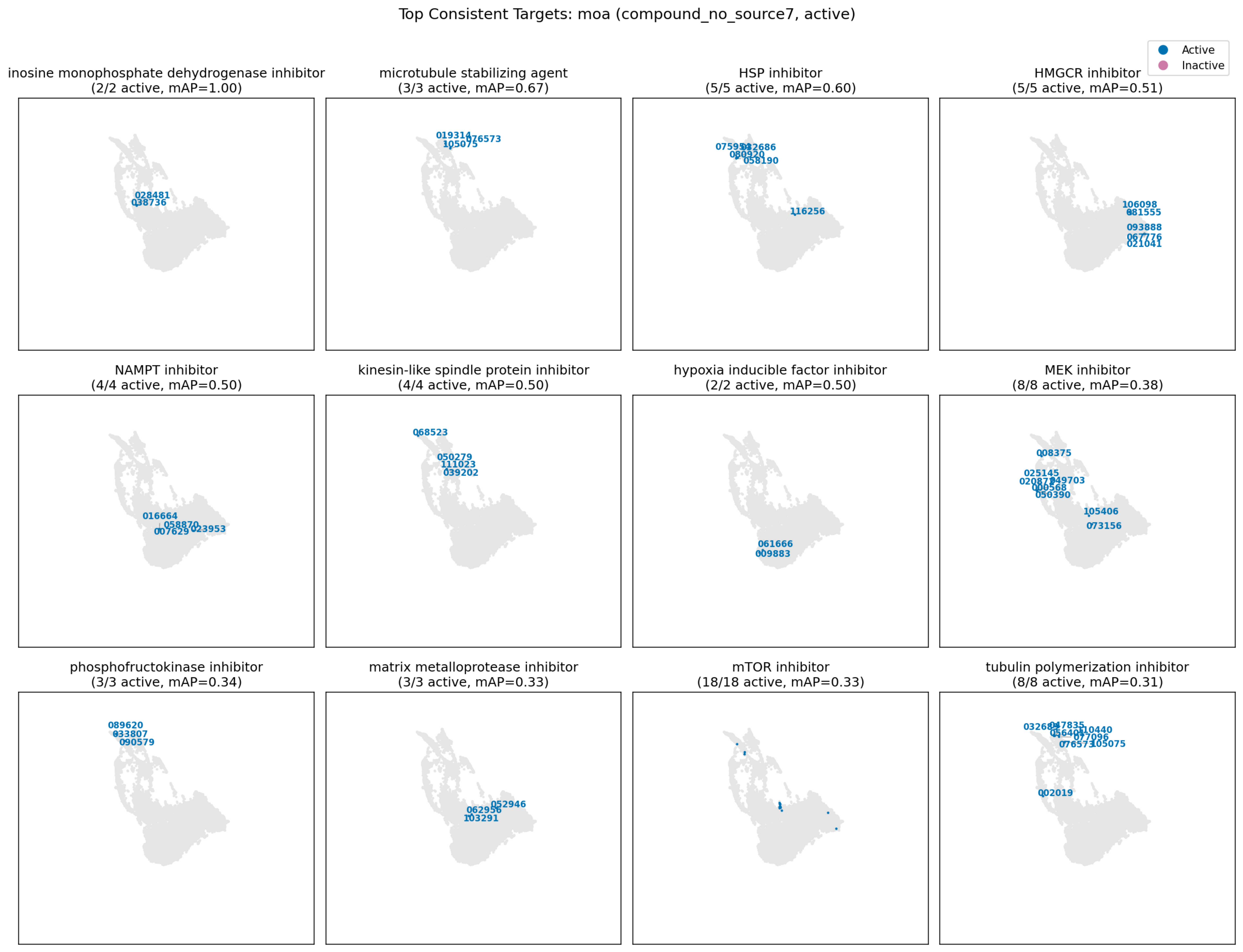
Top Consistent Targets: moa (compound_no_source7, active)
Active
Inactive
inosine monophosphate dehydrogenase inhibitor (2/2 active, mAP=1.00)
028481
038736
microtubule stabilizing agent (3/3 active, mAP=0.67)
019314
076573
105075
HSP inhibitor (5/5 active, mAP=0.60)
075958
032686
080920
058190
116256
HMGCR inhibitor (5/5 active, mAP=0.51)
106098
081555
093888
067776
021041
NAMPT inhibitor (4/4 active, mAP=0.50)
016664
058870
007629
023953
kinesin-like spindle protein inhibitor (4/4 active, mAP=0.50)
068523
050279
111023
039202
hypoxia inducible factor inhibitor (2/2 active, mAP=0.50)
061666
009883
MEK inhibitor (8/8 active, mAP=0.38)
008375
025145
049703
000568
050390
105406
073156
phosphofructokinase inhibitor (3/3 active, mAP=0.34)
089620
033807
090579
matrix metalloprotease inhibitor (3/3 active, mAP=0.33)
052946
062956
103291
mTOR inhibitor (18/18 active, mAP=0.33)
tubulin polymerization inhibitor (8/8 active, mAP=0.31)
047835
110440
077096
076573
105075
002019

motive biokg

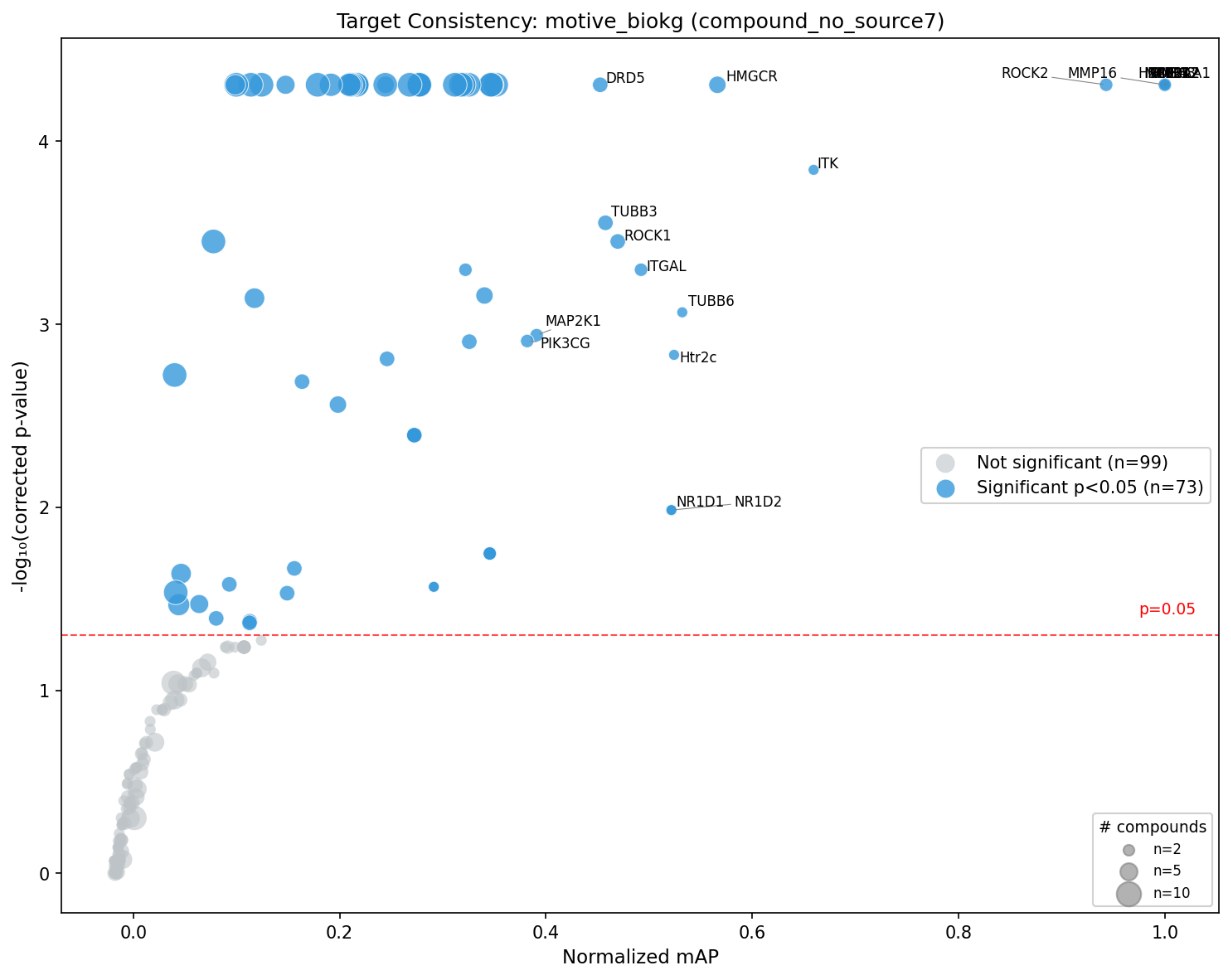
Target Consistency: motive_biokg (compound_no_source7)
DRD5
HMGCR
ROCK2
MMP16
ITK
TUBB3
ROCK1
ITGAL
TUBB6
MAP2K1
PIK3CG
Htr2c
NR1D1
NR1D2
Not significant (n=99)
Significant p<0.05 (n=73)
p=0.05
### compounds
n=2
n=5
n=10
-log₁₀(corrected p-value)
Normalized mAP
0
1
2
3
4
0.0
0.2
0.4
0.6
0.8
1.0

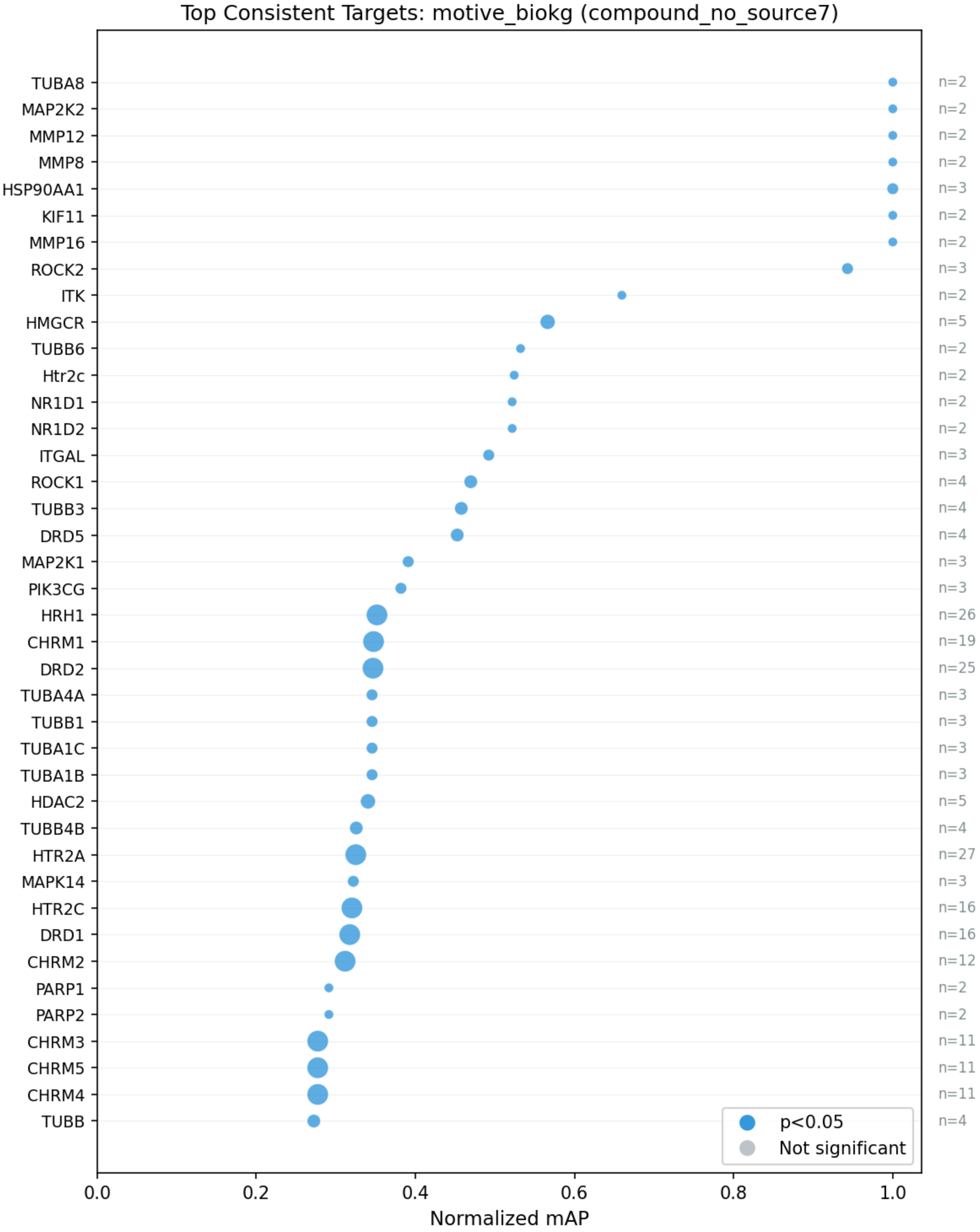

Top Consistent Targets: motive_biokg (compound_no_source7)
TUBA8
MAP2K2
MMP12
MMP8
HSP90AA1
KIF11
MMP16
ROCK2
ITK
HMGCR
TUBB6
Htr2c
NR1D1
NR1D2
ITGAL
ROCK1
TUBB3
DRD5
MAP2K1
PIK3CG
HRH1
CHRM1
DRD2
TUBA4A
TUBB1
TUBA1C
TUBA1B
HDAC2
TUBB4B
HTR2A
MAPK14
HTR2C
DRD1
CHRM2
PARP1
PARP2
CHRM3
CHRM5
CHRM4
TUBB
n=2
n=2
n=2
n=2
n=3
n=2
n=2
n=3
n=2
n=5
n=2
n=2
n=2
n=2
n=3
n=4
n=4
n=4
n=3
n=3
n=26
n=19
n=25
n=3
n=3
n=3
n=3
n=5
n=4
n=27
n=3
n=16
n=16
n=12
n=2
n=2
n=11
n=11
n=11
n=4
p<0.05
Not significant
0.0
0.2
0.4
0.6
0.8
1.0
Normalized mAP

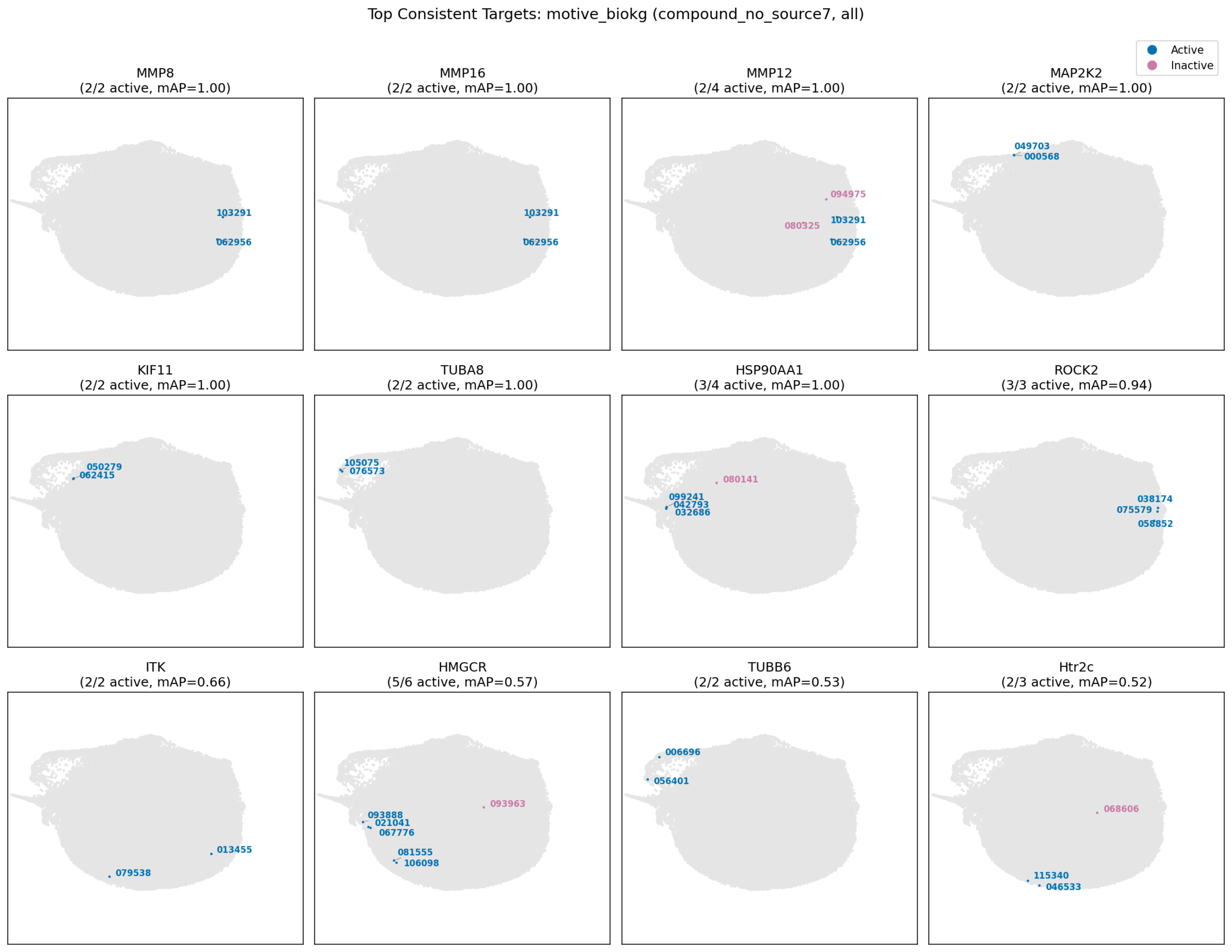

Top Consistent Targets: motive_biokg (compound_no_source7, all)
Active
Inactive
MMP8
(2/2 active, mAP=1.00)
103291
062956
MMP16
(2/2 active, mAP=1.00)
103291
062956
MMP12
(2/4 active, mAP=1.00)
094975
080325
103291
062956
MAP2K2
(2/2 active, mAP=1.00)
049703
000568
KIF11
(2/2 active, mAP=1.00)
050279
062415
TUBA8
(2/2 active, mAP=1.00)
105075
076573
HSP90AA1
(3/4 active, mAP=1.00)
080141
099241
042793
032686
ROCK2
(3/3 active, mAP=0.94)
038174
075579
058852
ITK
(2/2 active, mAP=0.66)
013455
079538
HMGCR
(5/6 active, mAP=0.57)
093963
093888
021041
067776
081555
106098
TUBB6
(2/2 active, mAP=0.53)
006696
056401
Htr2c
(2/3 active, mAP=0.52)
068606
115340
046533

Top Consistent Targets: motive_biokg (compound_no_source7, active)

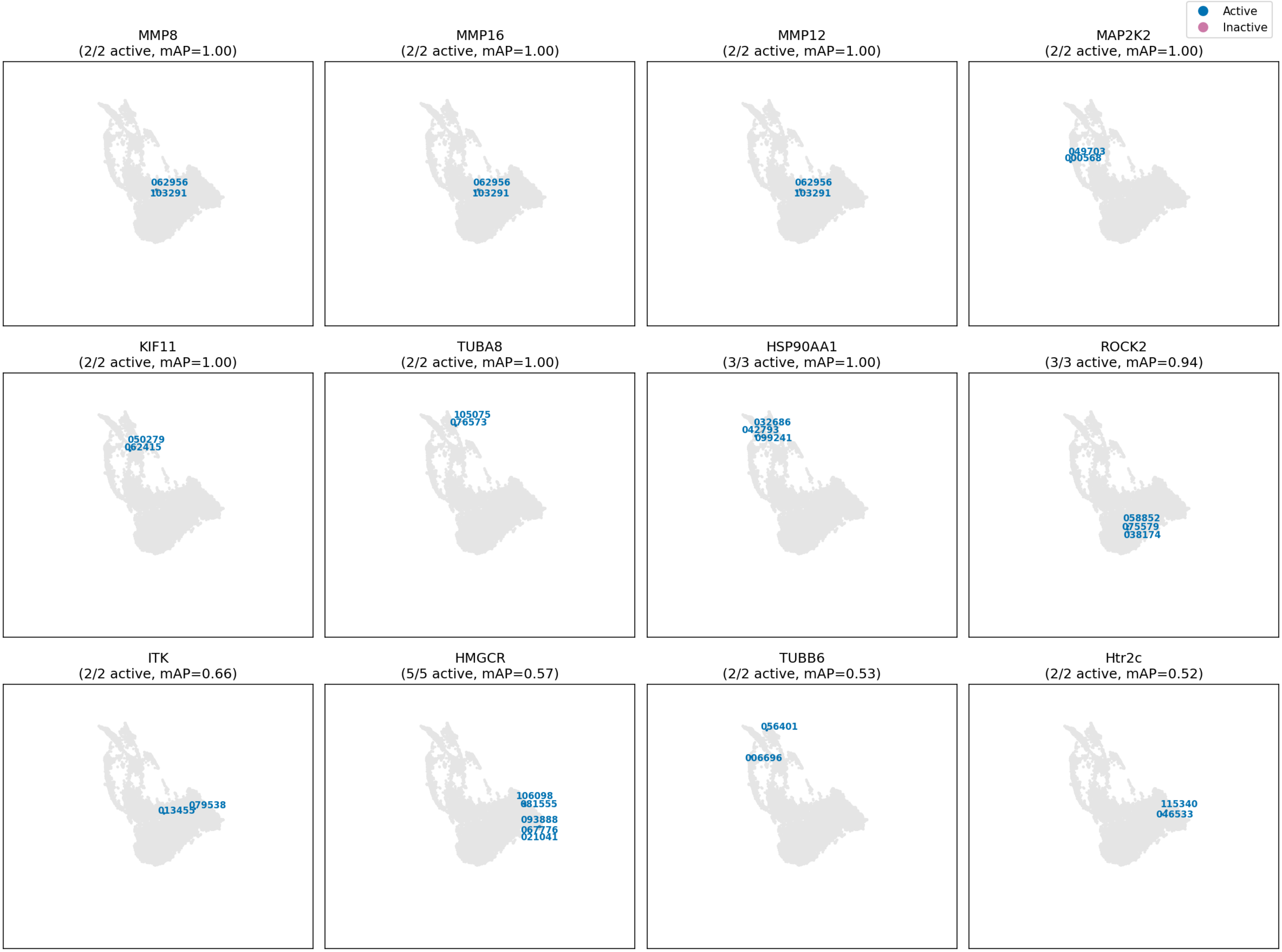

motive opentargets

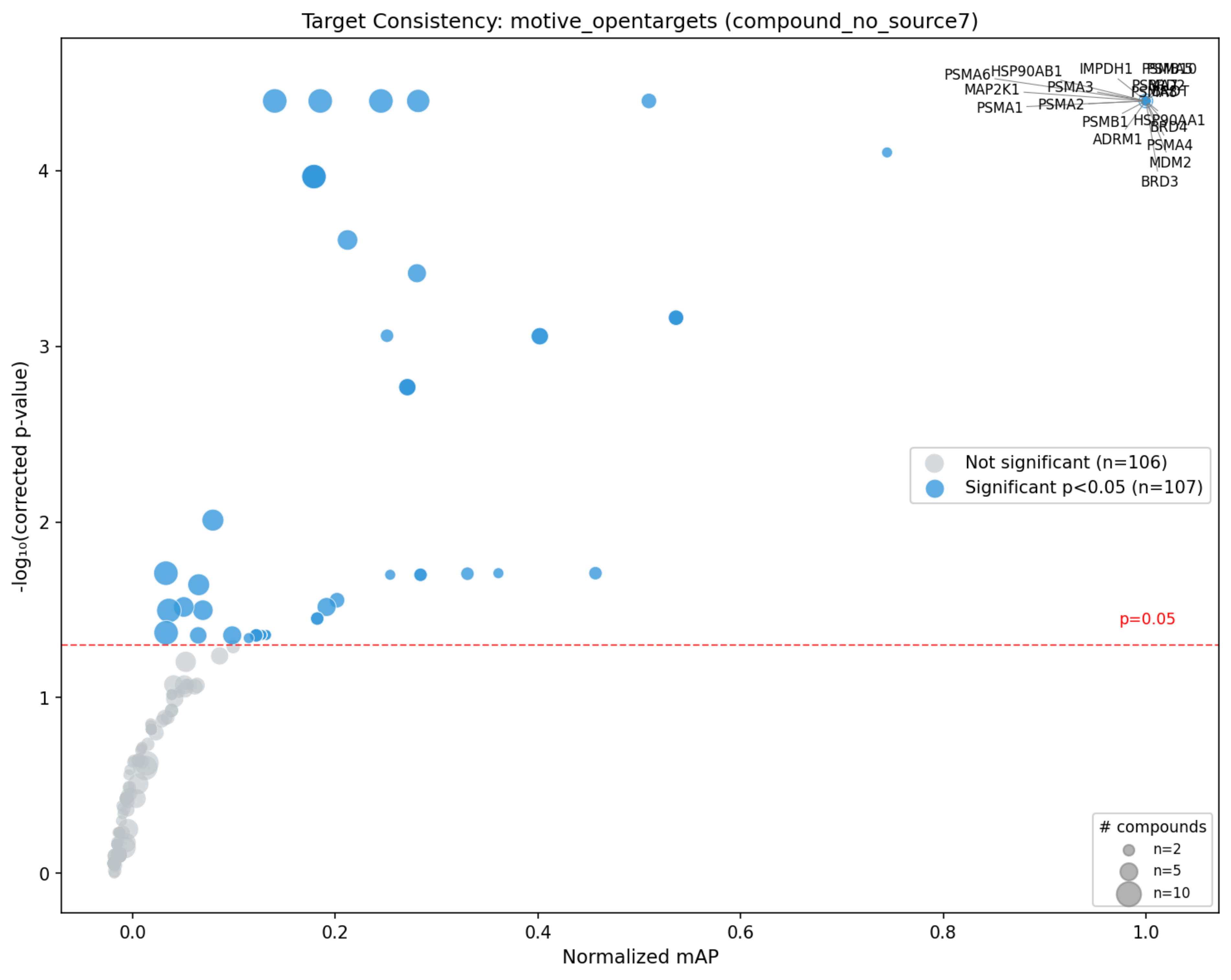
Target Consistency: motive_opentargets (compound_no_source7)
-log₁₀(corrected p-value)
Normalized mAP
0
1
2
3
4
0.0
0.2
0.4
0.6
0.8
1.0
PSMA6
HSP90AB1
IMPDH1
MAP2K1
PSMA3
PSMA1
PSMA2
PSMB1
HSP90AA1
BRD4
ADRM1
PSMA4
MDM2
BRD3
p=0.05
Not significant (n=106)
Significant p<0.05 (n=107)
### compounds
n=2
n=5
n=10

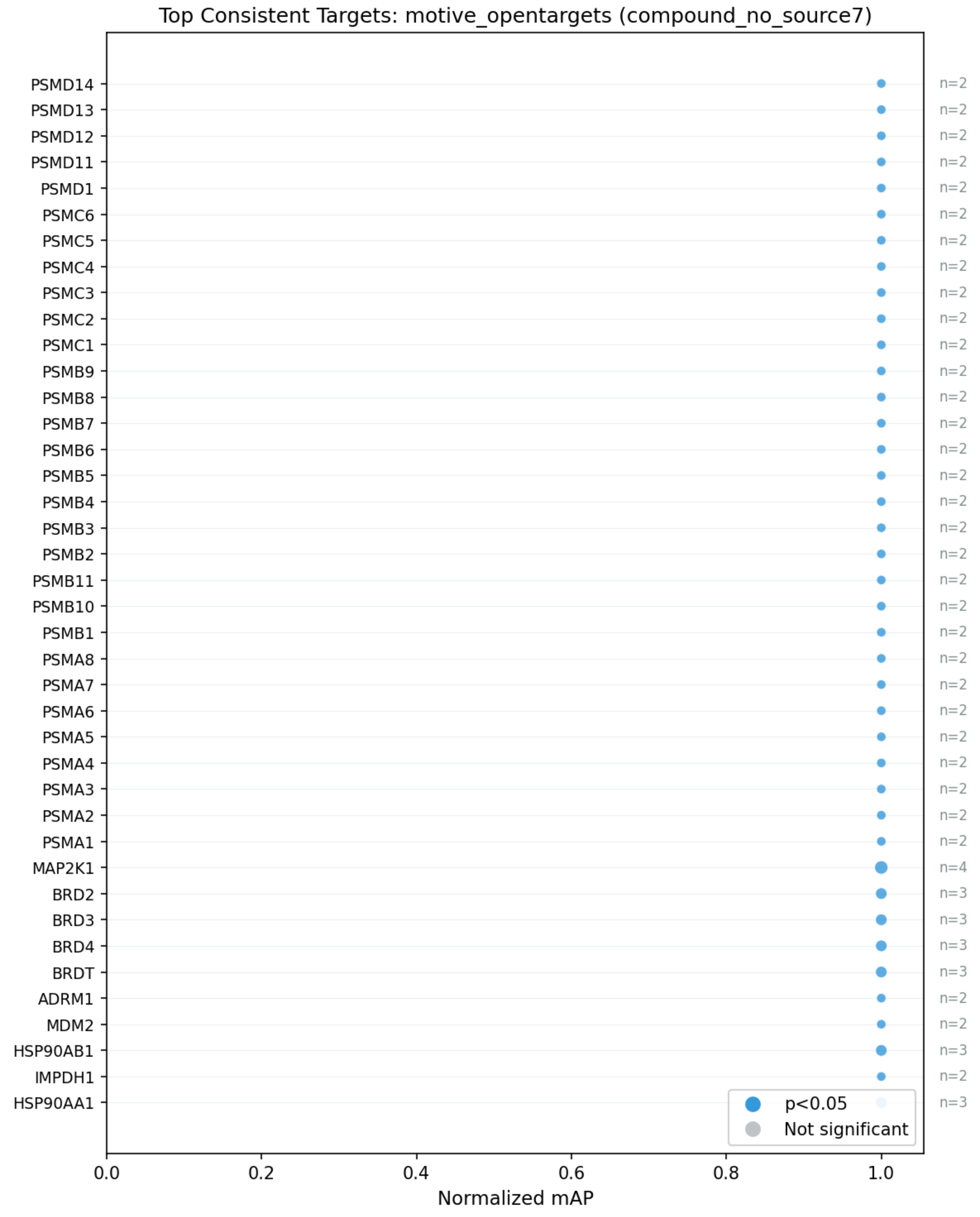
Top Consistent Targets: motive_opentargets (compound_no_source7)
PSMD14 n=2
PSMD13 n=2
PSMD12 n=2
PSMD11 n=2
PSMD1 n=2
PSMC6 n=2
PSMC5 n=2
PSMC4 n=2
PSMC3 n=2
PSMC2 n=2
PSMC1 n=2
PSMB9 n=2
PSMB8 n=2
PSMB7 n=2
PSMB6 n=2
PSMB5 n=2
PSMB4 n=2
PSMB3 n=2
PSMB2 n=2
PSMB11 n=2
PSMB10 n=2
PSMB1 n=2
PSMA8 n=2
PSMA7 n=2
PSMA6 n=2
PSMA5 n=2
PSMA4 n=2
PSMA3 n=2
PSMA2 n=2
PSMA1 n=2
MAP2K1 n=4
BRD2 n=3
BRD3 n=3
BRD4 n=3
BRDT n=3
ADRM1 n=2
MDM2 n=2
HSP90AB1 n=3
IMPDH1 n=2
HSP90AA1 n=3
p<0.05
Not significant
0.0
0.2
0.4
0.6
0.8
1.0
Normalized mAP

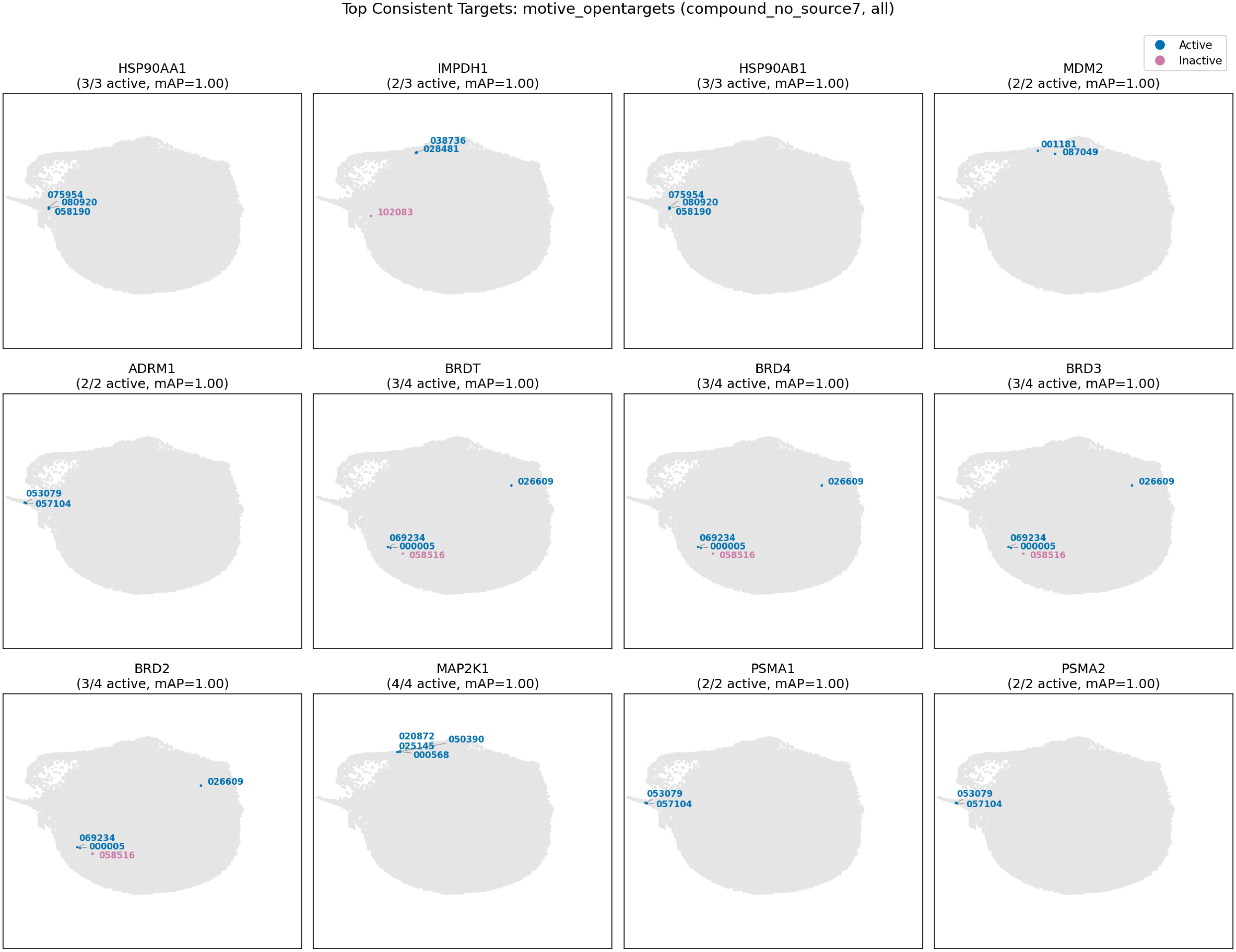
Top Consistent Targets: motive_opentargets (compound_no_source7, all)
Active
Inactive
HSP90AA1
(3/3 active, mAP=1.00)
075954
080920
058190
IMPDH1
(2/3 active, mAP=1.00)
038736
028481
102083
HSP90AB1
(3/3 active, mAP=1.00)
075954
080920
058190
MDM2
(2/2 active, mAP=1.00)
001181
087049
ADRM1
(2/2 active, mAP=1.00)
053079
057104
BRDT
(3/4 active, mAP=1.00)
026609
069234
000005
058516
BRD4
(3/4 active, mAP=1.00)
026609
069234
000005
058516
BRD3
(3/4 active, mAP=1.00)
026609
069234
000005
058516
BRD2
(3/4 active, mAP=1.00)
026609
069234
000005
058516
MAP2K1
(4/4 active, mAP=1.00)
020872
050390
025145
000568
PSMA1
(2/2 active, mAP=1.00)
053079
057104
PSMA2
(2/2 active, mAP=1.00)
053079
057104

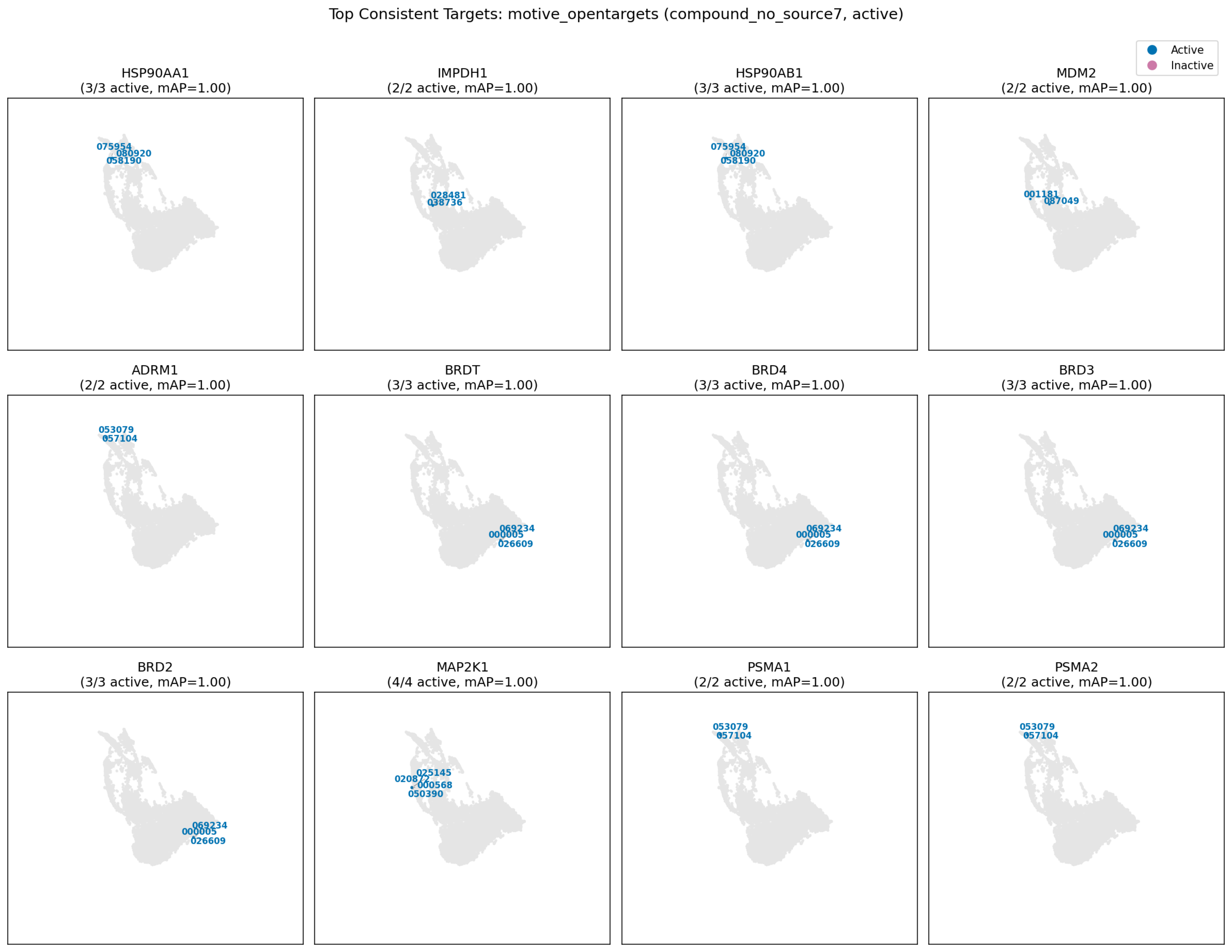

Top Consistent Targets: motive_opentargets (compound_no_source7, active)
Active
Inactive
HSP90AA1
(3/3 active, mAP=1.00)
075954
080920
058190
IMPDH1
(2/2 active, mAP=1.00)
028481
038736
HSP90AB1
(3/3 active, mAP=1.00)
075954
080920
058190
MDM2
(2/2 active, mAP=1.00)
001181
087049
ADRM1
(2/2 active, mAP=1.00)
053079
057104
BRDT
(3/3 active, mAP=1.00)
069234
000005
026609
BRD4
(3/3 active, mAP=1.00)
069234
000005
026609
BRD3
(3/3 active, mAP=1.00)
069234
000005
026609
BRD2
(3/3 active, mAP=1.00)
069234
000005
026609
MAP2K1
(4/4 active, mAP=1.00)
025145
020872
000568
050390
PSMA1
(2/2 active, mAP=1.00)
053079
057104
PSMA2
(2/2 active, mAP=1.00)
053079
057104

motive primekg

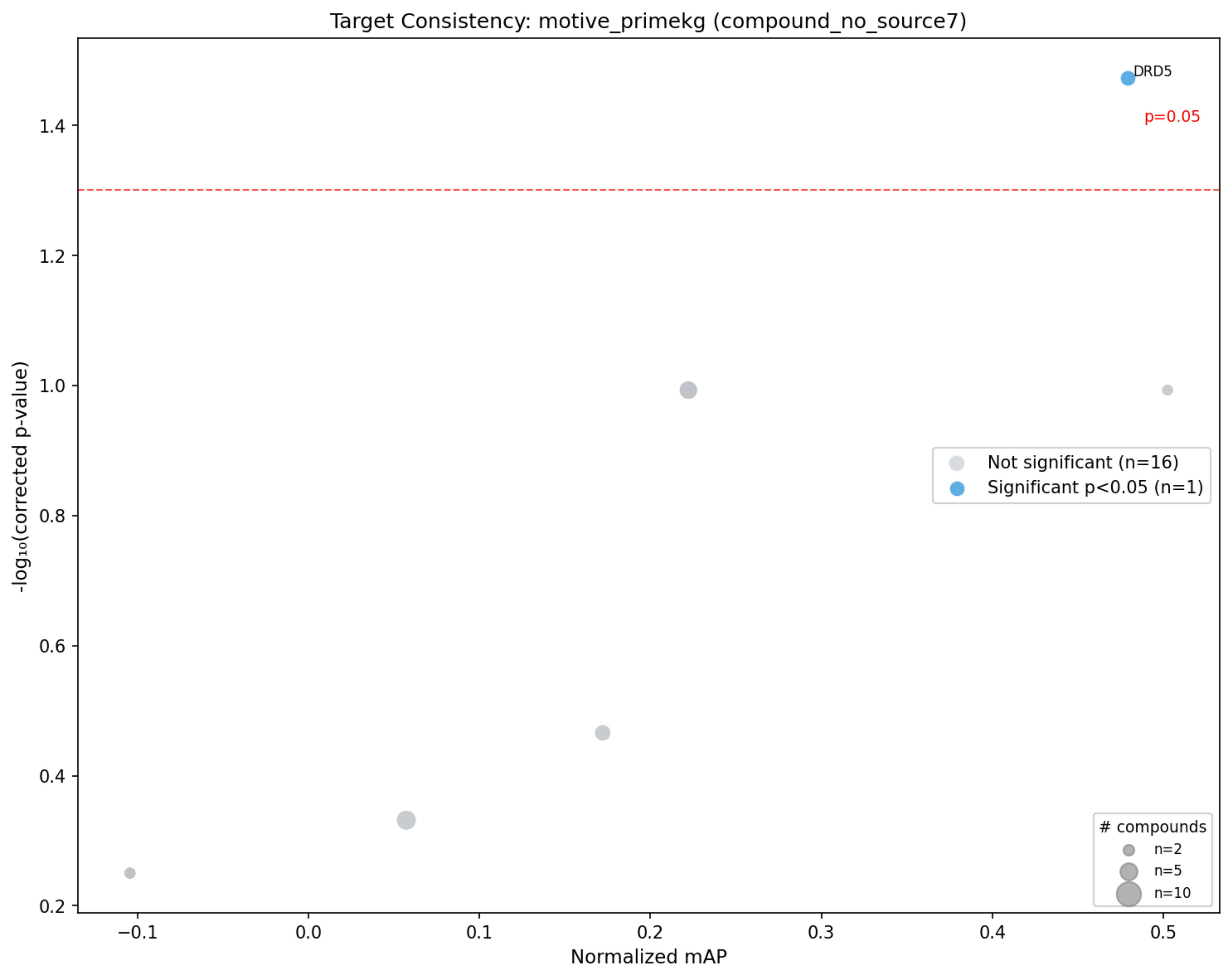
Target Consistency: motive_primekg (compound_no_source7)
DRD5
p=0.05
Not significant (n=16)
Significant p<0.05 (n=1)
### compounds
n=2
n=5
n=10
-log₁₀(corrected p-value)
Normalized mAP
1.4
1.2
1.0
0.8
0.6
0.4
0.2
−0.1
0.0
0.1
0.2
0.3
0.4
0.5

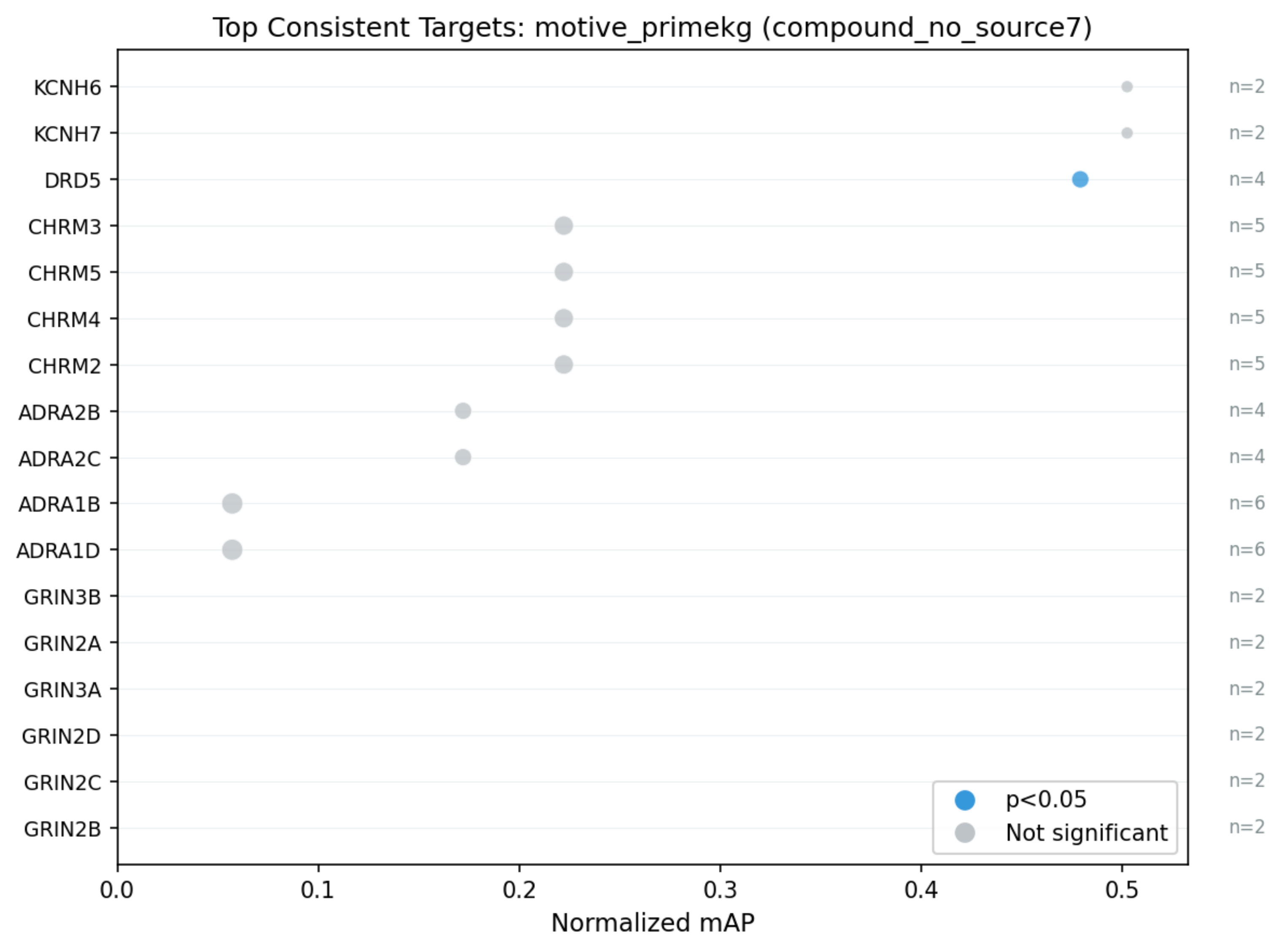

Top Consistent Targets: motive_primekg (compound_no_source7)
KCNH6
KCNH7
DRD5
CHRM3
CHRM5
CHRM4
CHRM2
ADRA2B
ADRA2C
ADRA1B
ADRA1D
GRIN3B
GRIN2A
GRIN3A
GRIN2D
GRIN2C
GRIN2B
n=2
n=2
n=4
n=5
n=5
n=5
n=5
n=4
n=4
n=6
n=6
n=2
n=2
n=2
n=2
n=2
n=2
p<0.05
Not significant
0.0
0.1
0.2
0.3
0.4
0.5
Normalized mAP

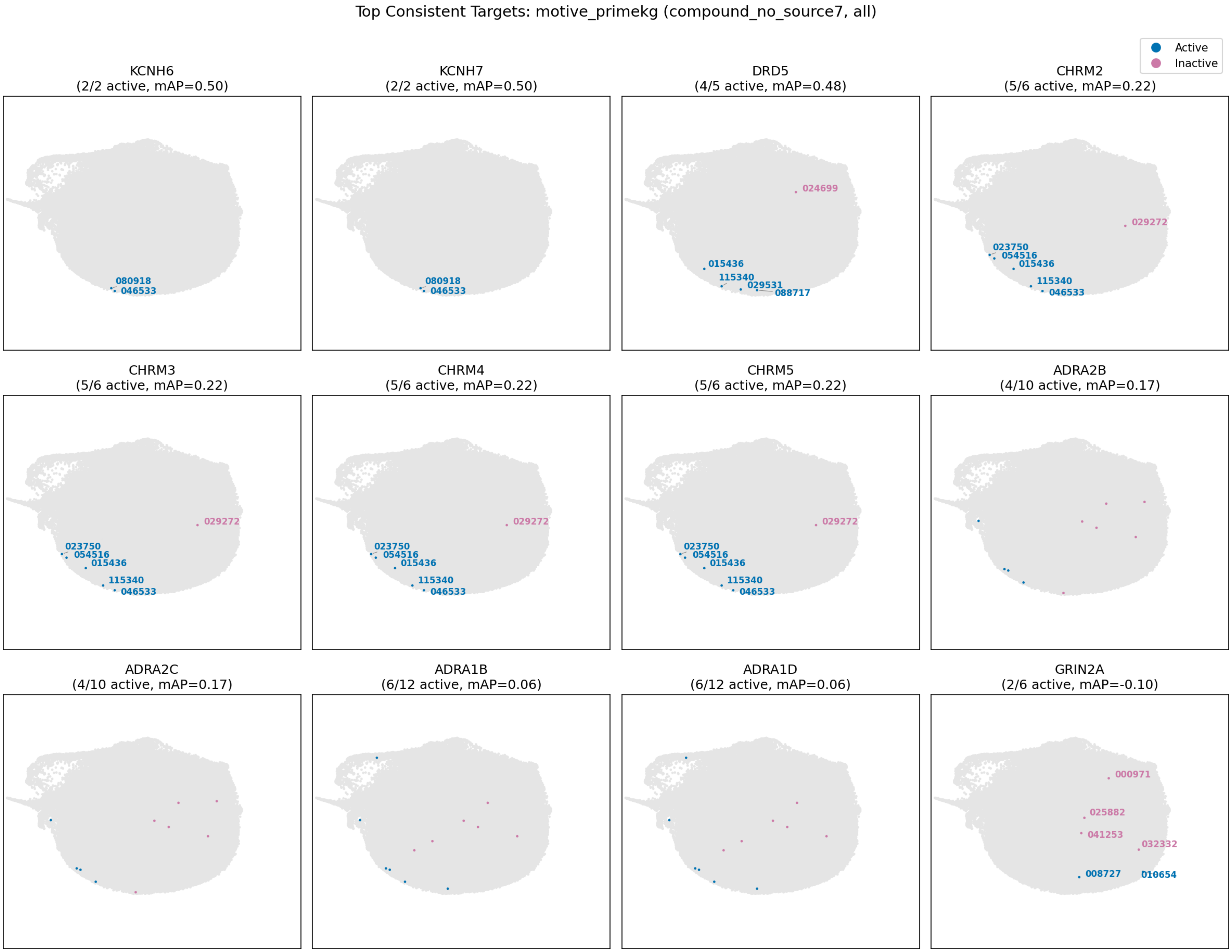
Top Consistent Targets: motive_primekg (compound_no_source7, all)
Active
Inactive
KCNH6
(2/2 active, mAP=0.50)
080918
046533
KCNH7
(2/2 active, mAP=0.50)
080918
046533
DRD5
(4/5 active, mAP=0.48)
024699
015436
115340
029531
088717
CHRM2
(5/6 active, mAP=0.22)
029272
023750
054516
015436
115340
046533
CHRM3
(5/6 active, mAP=0.22)
029272
023750
054516
015436
115340
046533
CHRM4
(5/6 active, mAP=0.22)
029272
023750
054516
015436
115340
046533
CHRM5
(5/6 active, mAP=0.22)
029272
023750
054516
015436
115340
046533
ADRA2B
(4/10 active, mAP=0.17)
ADRA2C
(4/10 active, mAP=0.17)
ADRA1B
(6/12 active, mAP=0.06)
ADRA1D
(6/12 active, mAP=0.06)
GRIN2A
(2/6 active, mAP=-0.10)
000971
025882
041253
032332
008727
010654

Top Consistent Targets: motive_primekg (compound_no_source7, active)

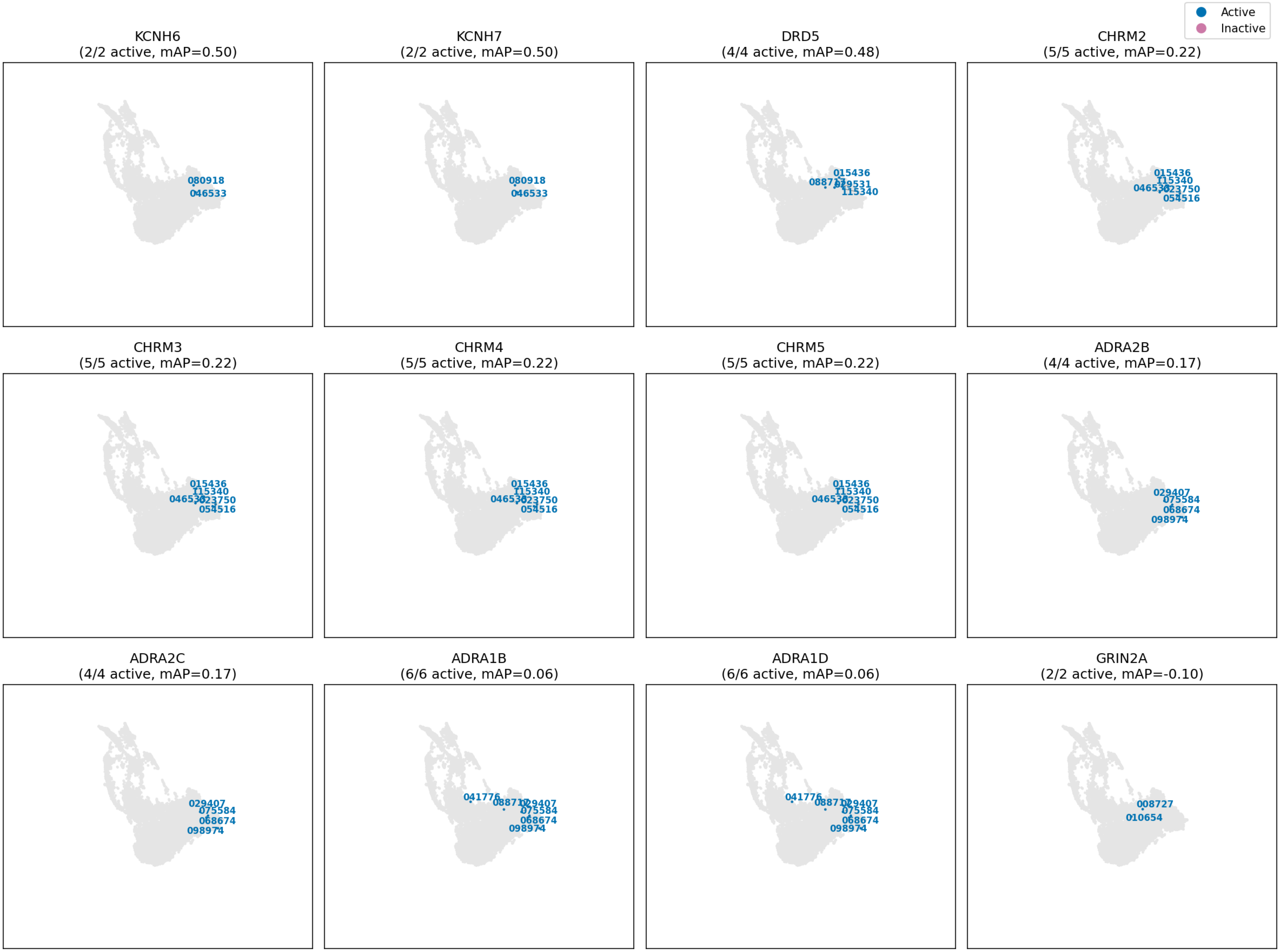

repurposing

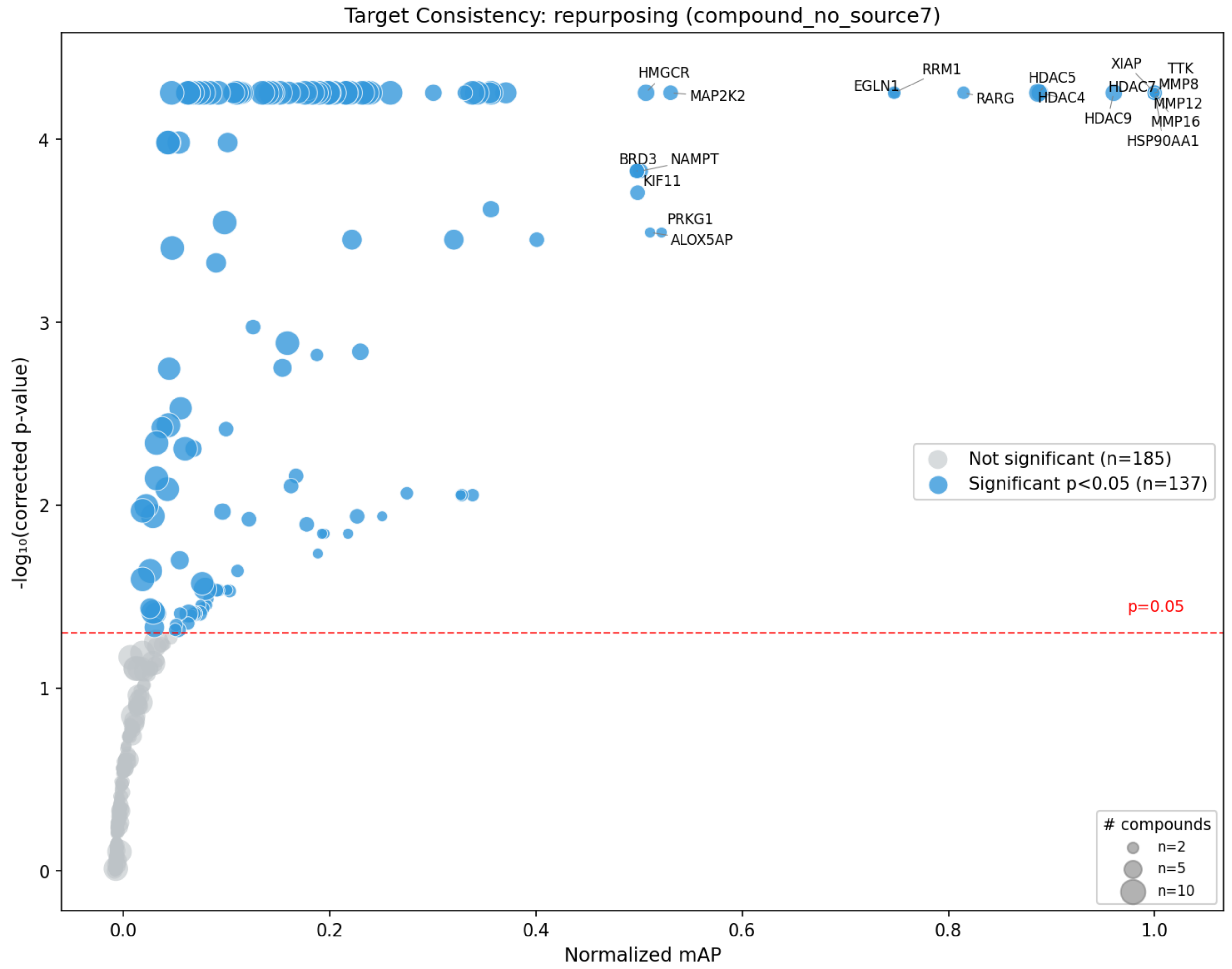

Target Consistency: repurposing (compound_no_source7)
-log₁₀(corrected p-value)
Normalized mAP
HMGCR
MAP2K2
EGLN1
RRM1
RARG
HDAC5
HDAC4
HDAC9
XIAP
HDAC7
TTK
MMP8
MMP12
MMP16
HSP90AA1
BRD3
NAMPT
KIF11
PRKG1
ALOX5AP
p=0.05
Not significant (n=185)
Significant p<0.05 (n=137)
### compounds
n=2
n=5
n=10

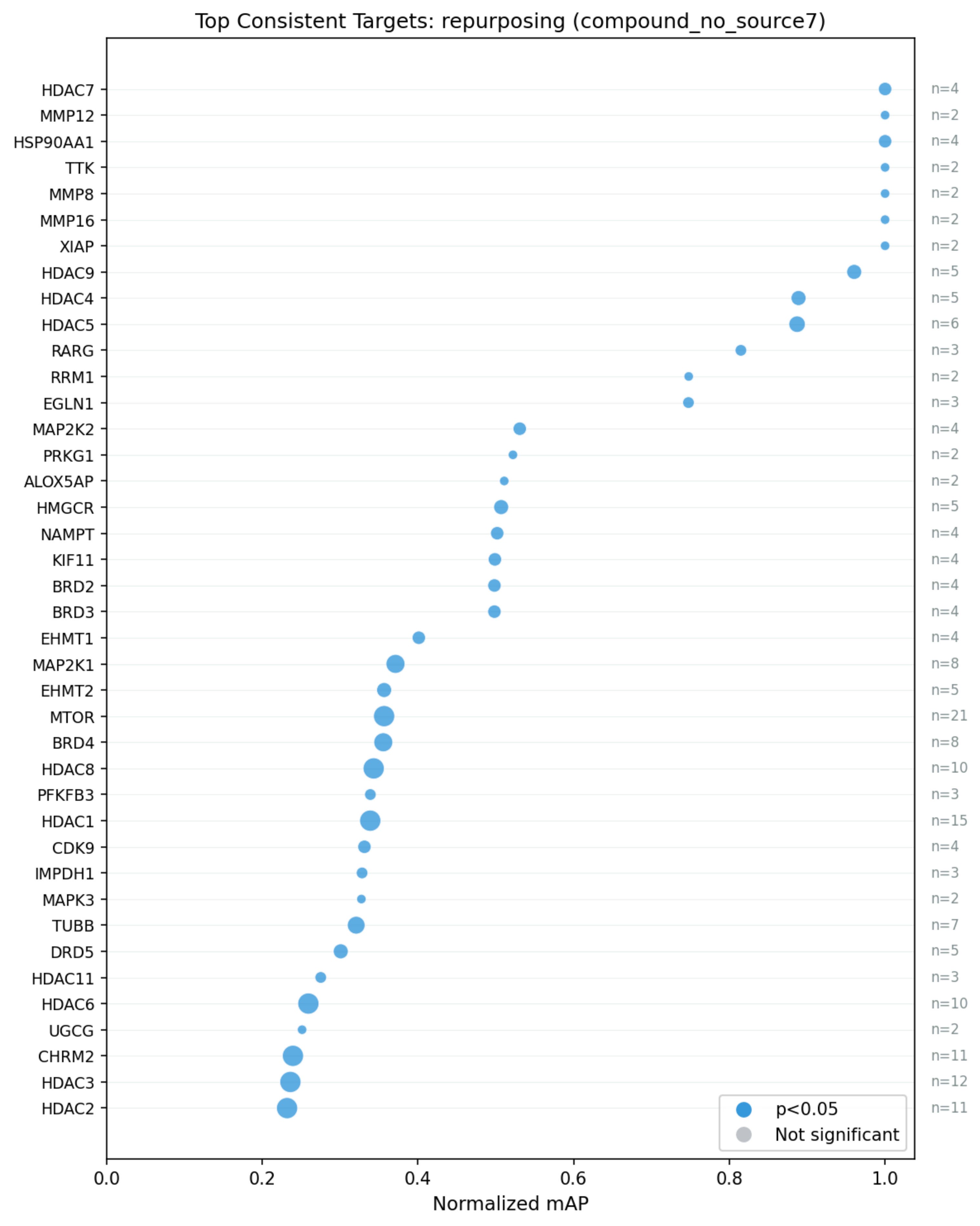
Top Consistent Targets: repurposing (compound_no_source7)
HDAC7
MMP12
HSP90AA1
TTK
MMP8
MMP16
XIAP
HDAC9
HDAC4
HDAC5
RARG
RRM1
EGLN1
MAP2K2
PRKG1
ALOX5AP
HMGCR
NAMPT
KIF11
BRD2
BRD3
EHMT1
MAP2K1
EHMT2
MTOR
BRD4
HDAC8
PFKFB3
HDAC1
CDK9
IMPDH1
MAPK3
TUBB
DRD5
HDAC11
HDAC6
UGCG
CHRM2
HDAC3
HDAC2
n=4
n=2
n=4
n=2
n=2
n=2
n=2
n=5
n=5
n=6
n=3
n=2
n=3
n=4
n=2
n=2
n=5
n=4
n=4
n=4
n=4
n=4
n=8
n=5
n=21
n=8
n=10
n=3
n=15
n=4
n=3
n=2
n=7
n=5
n=3
n=10
n=2
n=11
n=12
n=11
p<0.05
Not significant
0.0
0.2
0.4
0.6
0.8
1.0
Normalized mAP

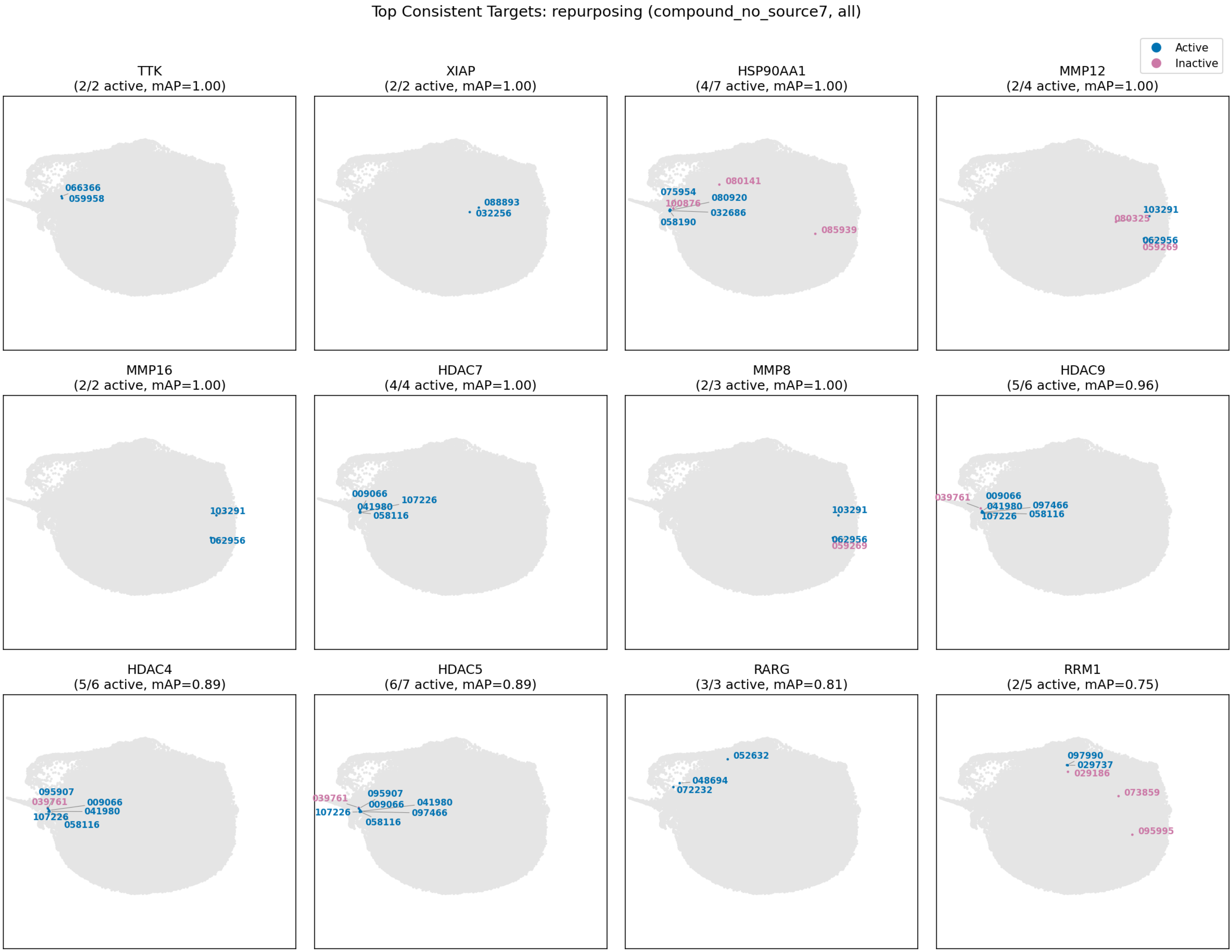

Top Consistent Targets: repurposing (compound_no_source7, all)
Active
Inactive
TTK
(2/2 active, mAP=1.00)
066366
059958
XIAP
(2/2 active, mAP=1.00)
088893
032256
HSP90AA1
(4/7 active, mAP=1.00)
080141
075954
100876
080920
032686
058190
085939
MMP12
(2/4 active, mAP=1.00)
103291
080325
062956
059269
MMP16
(2/2 active, mAP=1.00)
103291
062956
HDAC7
(4/4 active, mAP=1.00)
009066
107226
041980
058116
MMP8
(2/3 active, mAP=1.00)
103291
062956
059269
HDAC9
(5/6 active, mAP=0.96)
039761
009066
041980
097466
107226
058116
HDAC4
(5/6 active, mAP=0.89)
095907
039761
009066
041980
107226
058116
HDAC5
(6/7 active, mAP=0.89)
039761
095907
009066
041980
107226
097466
058116
RARG
(3/3 active, mAP=0.81)
052632
048694
072232
RRM1
(2/5 active, mAP=0.75)
097990
029737
029186
073859
095995

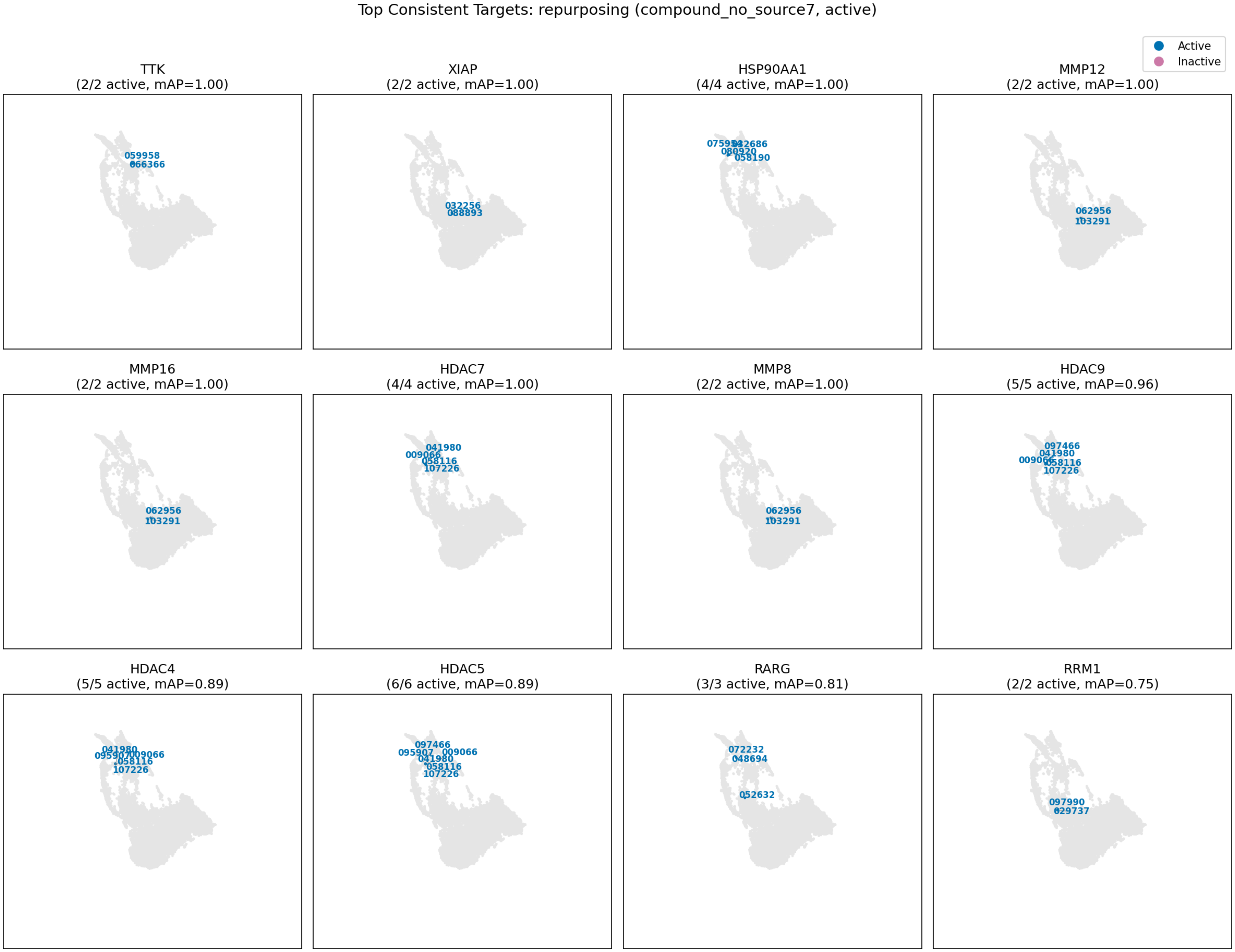
Top Consistent Targets: repurposing (compound_no_source7, active)
Active
Inactive
TTK
(2/2 active, mAP=1.00)
059958
XIAP
(2/2 active, mAP=1.00)
032256
088893
HSP90AA1
(4/4 active, mAP=1.00)
058190
MMP12
(2/2 active, mAP=1.00)
062956
MMP16
(2/2 active, mAP=1.00)
062956
HDAC7
(4/4 active, mAP=1.00)
041980
009066
107226
MMP8
(2/2 active, mAP=1.00)
062956
HDAC9
(5/5 active, mAP=0.96)
097466
041980
107226
HDAC4
(5/5 active, mAP=0.89)
107226
HDAC5
(6/6 active, mAP=0.89)
097466
095907
009066
041980
107226
RARG
(3/3 active, mAP=0.81)
072232
048694
RRM1
(2/2 active, mAP=0.75)
097990

toxcast assay

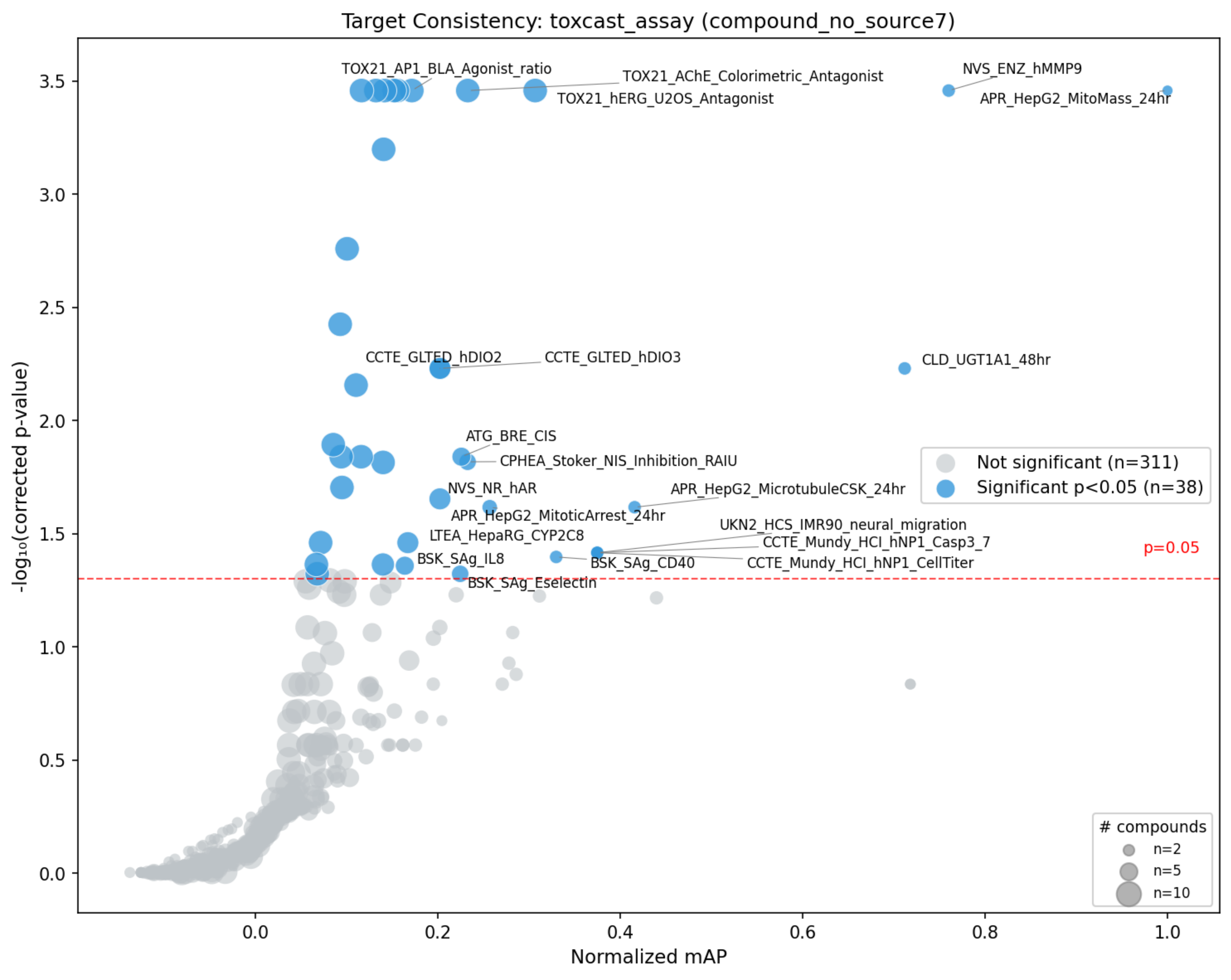
Target Consistency: toxcast_assay (compound_no_source7)
TOX21_AP1_BLA_Agonist_ratio
TOX21_AChE_Colorimetric_Antagonist
TOX21_hERG_U2OS_Antagonist
NVS_ENZ_hMMP9
APR_HepG2_MitoMass_24hr
CCTE_GLTED_hDIO2
CCTE_GLTED_hDIO3
CLD_UGT1A1_48hr
ATG_BRE_CIS
CPHEA_Stoker_NIS_Inhibition_RAIU
NVS_NR_hAR
APR_HepG2_MicrotubuleCSK_24hr
APR_HepG2_MitoticArrest_24hr
UKN2_HCS_IMR90_neural_migration
LTEA_HepaRG_CYP2C8
CCTE_Mundy_HCI_hNP1_Casp3_7
BSK_SAg_IL8
BSK_SAg_CD40
CCTE_Mundy_HCI_hNP1_CellTiter
BSK_SAg_Eselectin
p=0.05
Not significant (n=311)
Significant p<0.05 (n=38)
### compounds
n=2
n=5
n=10
-log₁₀(corrected p-value)
Normalized mAP
0.0
0.5
1.0
1.5
2.0
2.5
3.0
3.5
0.0
0.2
0.4
0.6
0.8
1.0

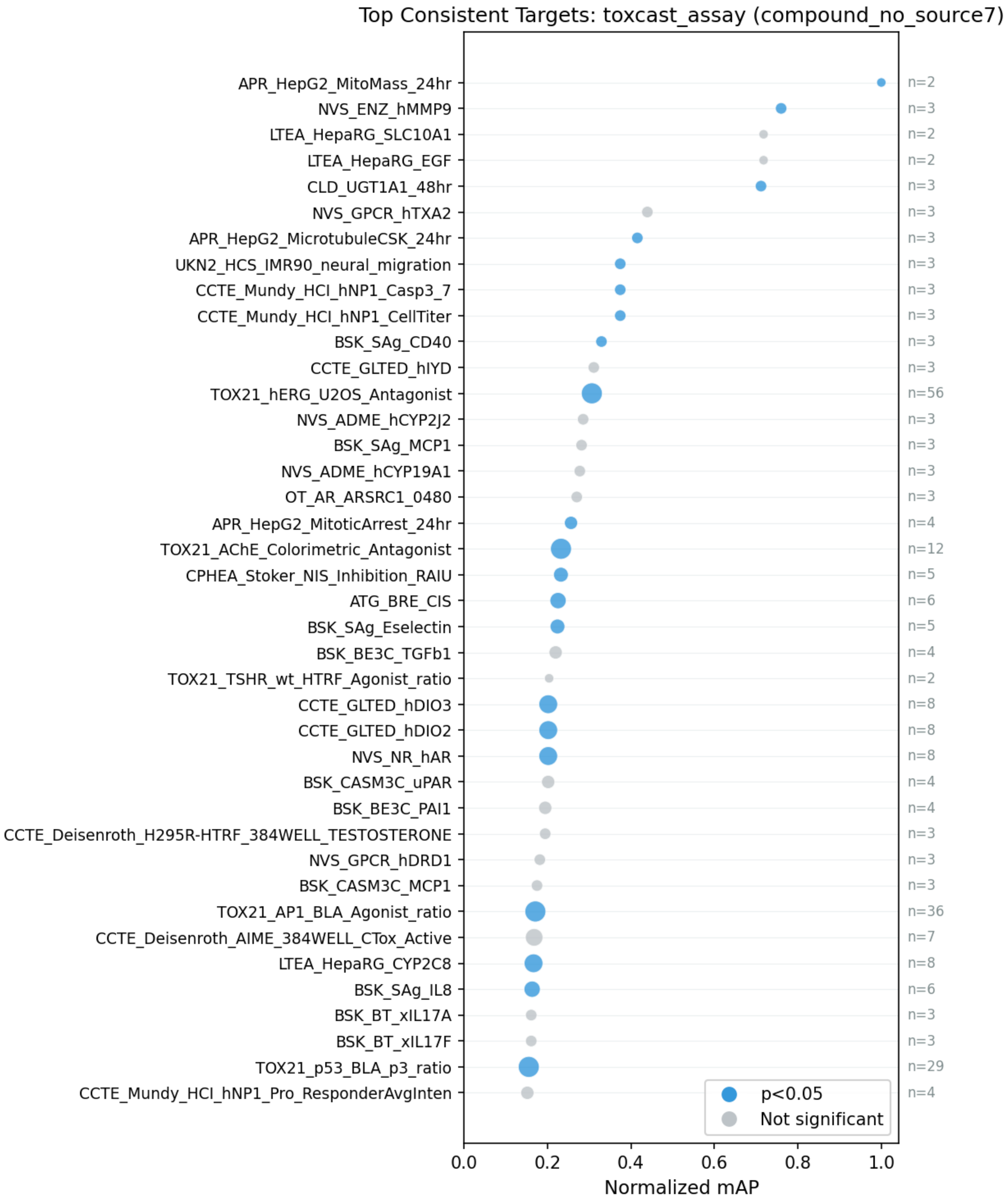
Top Consistent Targets: toxcast_assay (compound_no_source7)
APR_HepG2_MitoMass_24hr
NVS_ENZ_hMMP9
LTEA_HepaRG_SLC10A1
LTEA_HepaRG_EGF
CLD_UGT1A1_48hr
NVS_GPCR_hTXA2
APR_HepG2_MicrotubuleCSK_24hr
UKN2_HCS_IMR90_neural_migration
CCTE_Mundy_HCI_hNP1_Casp3_7
CCTE_Mundy_HCI_hNP1_CellTiter
BSK_SAg_CD40
CCTE_GLTED_hIYD
TOX21_hERG_U2OS_Antagonist
NVS_ADME_hCYP2J2
BSK_SAg_MCP1
NVS_ADME_hCYP19A1
OT_AR_ARSRC1_0480
APR_HepG2_MitoticArrest_24hr
TOX21_AChE_Colorimetric_Antagonist
CPHEA_Stoker_NIS_Inhibition_RAIU
ATG_BRE_CIS
BSK_SAg_Eselectin
BSK_BE3C_TGFb1
TOX21_TSHR_wt_HTRF_Agonist_ratio
CCTE_GLTED_hDIO3
CCTE_GLTED_hDIO2
NVS_NR_hAR
BSK_CASM3C_uPAR
BSK_BE3C_PAI1
CCTE_Deisenroth_H295R-HTRF_384WELL_TESTOSTERONE
NVS_GPCR_hDRD1
BSK_CASM3C_MCP1
TOX21_AP1_BLA_Agonist_ratio
CCTE_Deisenroth_AIME_384WELL_CTox_Active
LTEA_HepaRG_CYP2C8
BSK_SAg_IL8
BSK_BT_xIL17A
BSK_BT_xIL17F
TOX21_p53_BLA_p3_ratio
CCTE_Mundy_HCI_hNP1_Pro_ResponderAvgInten
n=2
n=3
n=2
n=2
n=3
n=3
n=3
n=3
n=3
n=3
n=3
n=3
n=56
n=3
n=3
n=3
n=3
n=4
n=12
n=5
n=6
n=5
n=4
n=2
n=8
n=8
n=8
n=4
n=4
n=3
n=3
n=3
n=36
n=7
n=8
n=6
n=3
n=3
n=29
n=4
p<0.05
Not significant
0.0
0.2
0.4
0.6
0.8
1.0
Normalized mAP

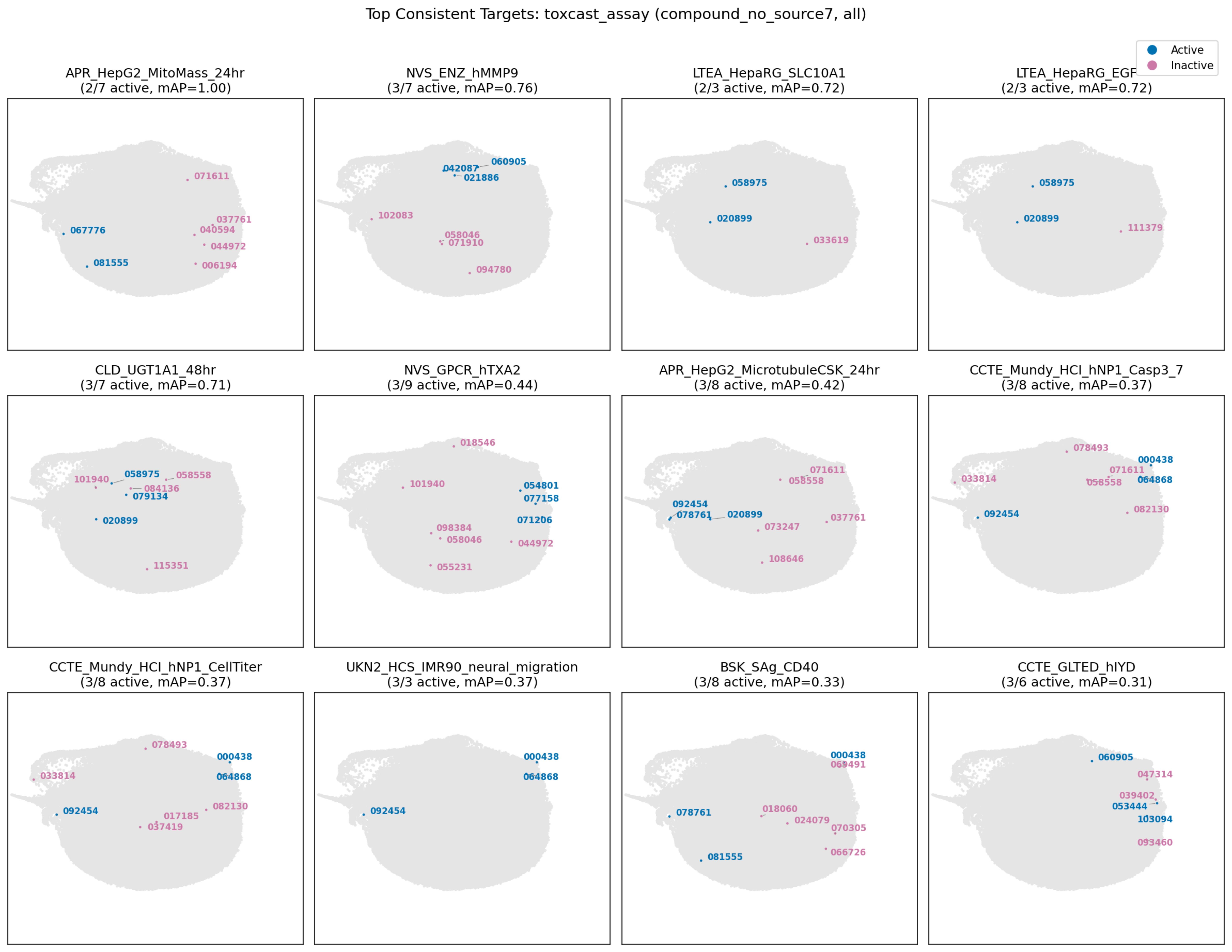
Top Consistent Targets: toxcast_assay (compound_no_source7, all)
Active
Inactive
APR_HepG2_MitoMass_24hr
(2/7 active, mAP=1.00)
071611
037761
040594
067776
044972
081555
006194
NVS_ENZ_hMMP9
(3/7 active, mAP=0.76)
060905
042087
021886
102083
058046
071910
094780
LTEA_HepaRG_SLC10A1
(2/3 active, mAP=0.72)
058975
020899
033619
LTEA_HepaRG_EGF
(2/3 active, mAP=0.72)
058975
020899
111379
CLD_UGT1A1_48hr
(3/7 active, mAP=0.71)
058975
058558
101940
084136
079134
020899
115351
NVS_GPCR_hTXA2
(3/9 active, mAP=0.44)
018546
101940
054801
077158
071206
098384
058046
044972
055231
APR_HepG2_MicrotubuleCSK_24hr
(3/8 active, mAP=0.42)
071611
058558
092454
078761
020899
037761
073247
108646
CCTE_Mundy_HCI_hNP1_Casp3_7
(3/8 active, mAP=0.37)
078493
000438
071611
033814
058558
064868
092454
082130
CCTE_Mundy_HCI_hNP1_CellTiter
(3/8 active, mAP=0.37)
078493
000438
033814
064868
082130
092454
017185
037419
UKN2_HCS_IMR90_neural_migration
(3/3 active, mAP=0.37)
000438
064868
092454
BSK_SAg_CD40
(3/8 active, mAP=0.33)
000438
069491
018060
078761
024079
070305
066726
081555
CCTE_GLTED_hIYD
(3/6 active, mAP=0.31)
060905
047314
039402
053444
103094
093460

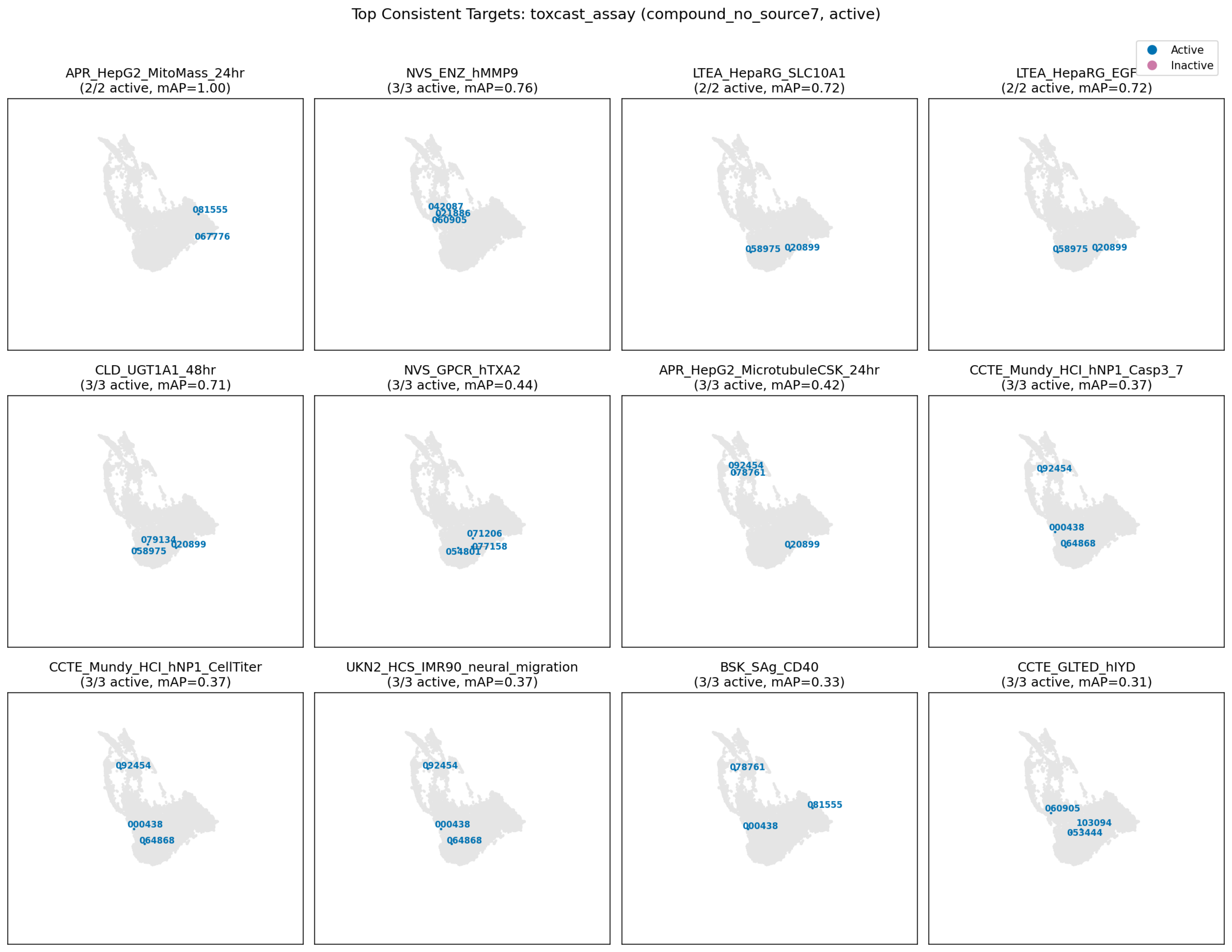

Top Consistent Targets: toxcast_assay (compound_no_source7, active)
Active
Inactive
APR_HepG2_MitoMass_24hr
(2/2 active, mAP=1.00)
081555
067776
NVS_ENZ_hMMP9
(3/3 active, mAP=0.76)
042087
021886
060905
LTEA_HepaRG_SLC10A1
(2/2 active, mAP=0.72)
058975
020899
LTEA_HepaRG_EGF
(2/2 active, mAP=0.72)
058975
020899
CLD_UGT1A1_48hr
(3/3 active, mAP=0.71)
079134
058975
020899
NVS_GPCR_hTXA2
(3/3 active, mAP=0.44)
071206
054801
077158
APR_HepG2_MicrotubuleCSK_24hr
(3/3 active, mAP=0.42)
092454
078761
020899
CCTE_Mundy_HCI_hNP1_Casp3_7
(3/3 active, mAP=0.37)
092454
000438
064868
CCTE_Mundy_HCI_hNP1_CellTiter
(3/3 active, mAP=0.37)
092454
000438
064868
UKN2_HCS_IMR90_neural_migration
(3/3 active, mAP=0.37)
092454
000438
064868
BSK_SAg_CD40
(3/3 active, mAP=0.33)
078761
081555
000438
CCTE_GLTED_hIYD
(3/3 active, mAP=0.31)
060905
103094
053444

uniprot

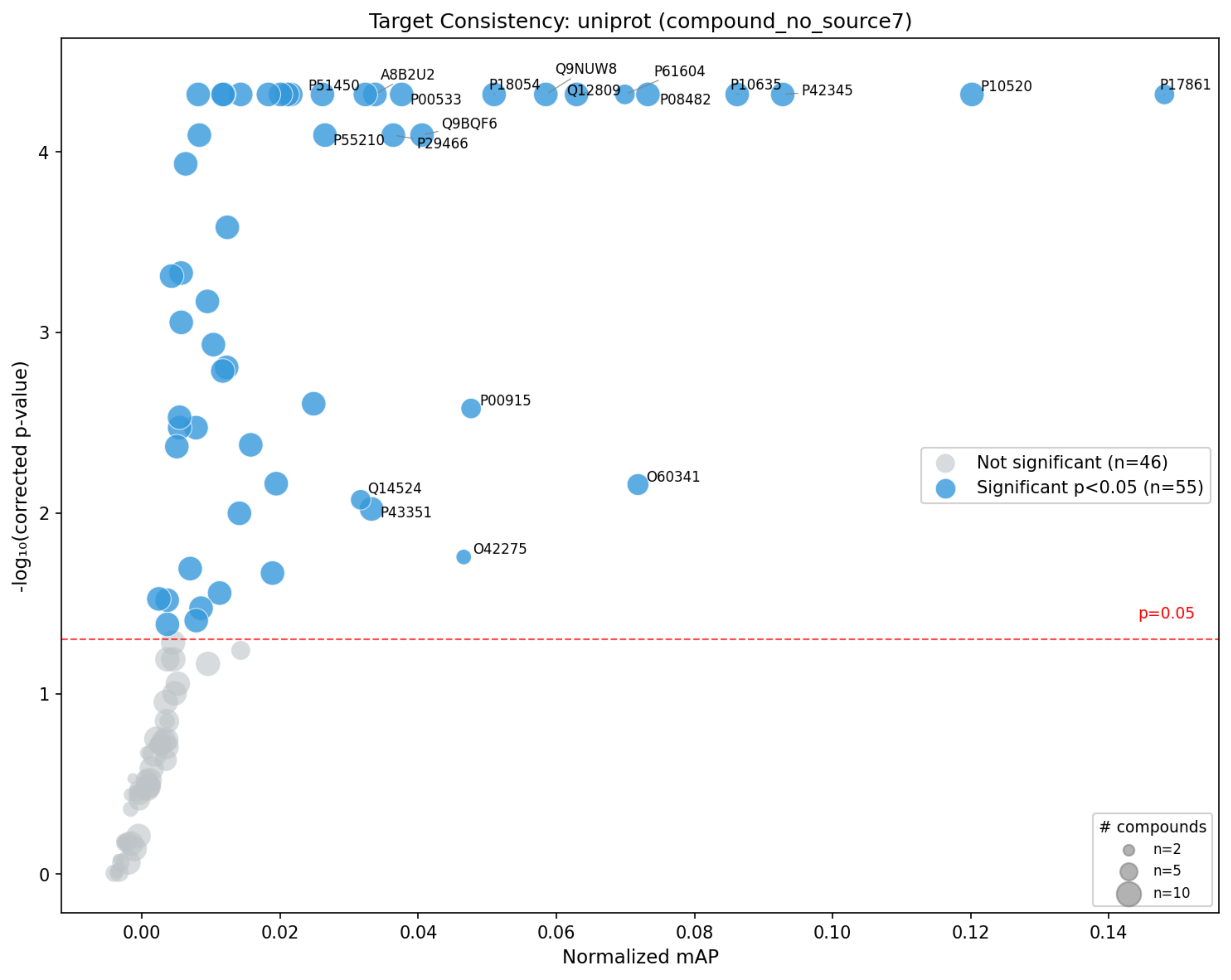

Target Consistency: uniprot (compound_no_source7)
-log₁₀(corrected p-value)
Normalized mAP
P51450
A8B2U2
P00533
P18054
Q9NUW8
Q12809
P61604
P08482
P10635
P42345
P10520
P17861
Q9BQF6
P55210
P29466
P00915
Q14524
P43351
O60341
O42275
p=0.05
Not significant (n=46)
Significant p<0.05 (n=55)
### compounds
n=2
n=5
n=10
0
1
2
3
4
0.00
0.02
0.04
0.06
0.08
0.10
0.12
0.14

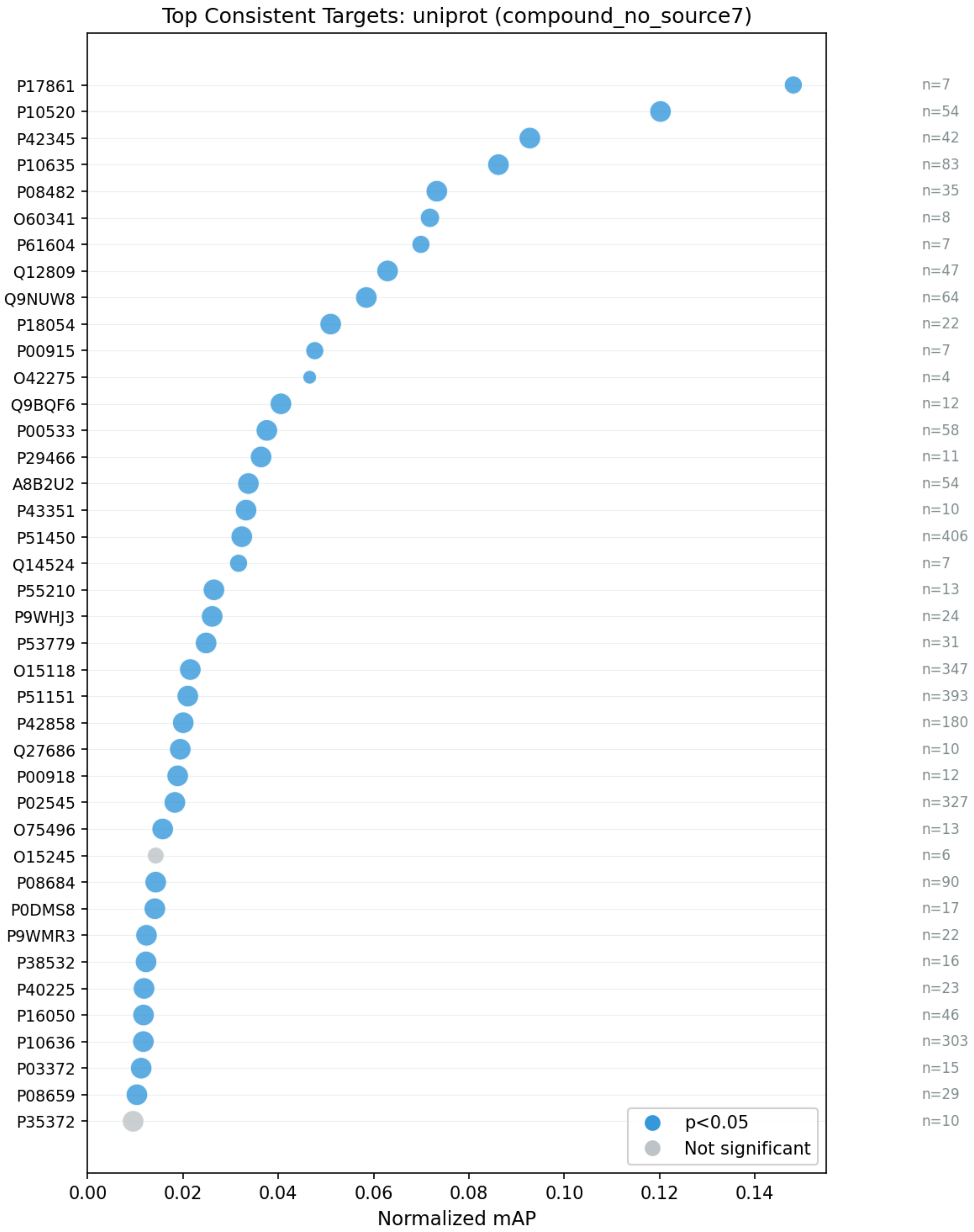
Top Consistent Targets: uniprot (compound_no_source7)
P17861
P10520
P42345
P10635
P08482
O60341
P61604
Q12809
Q9NUW8
P18054
P00915
O42275
Q9BQF6
P00533
P29466
A8B2U2
P43351
P51450
Q14524
P55210
P9WHJ3
P53779
O15118
P51151
P42858
Q27686
P00918
P02545
O75496
O15245
P08684
P0DMS8
P9WMR3
P38532
P40225
P16050
P10636
P03372
P08659
P35372
n=7
n=54
n=42
n=83
n=35
n=8
n=7
n=47
n=64
n=22
n=7
n=4
n=12
n=58
n=11
n=54
n=10
n=406
n=7
n=13
n=24
n=31
n=347
n=393
n=180
n=10
n=12
n=327
n=13
n=6
n=90
n=17
n=22
n=16
n=23
n=46
n=303
n=15
n=29
n=10
p<0.05
Not significant
0.00
0.02
0.04
0.06
0.08
0.10
0.12
0.14
Normalized mAP

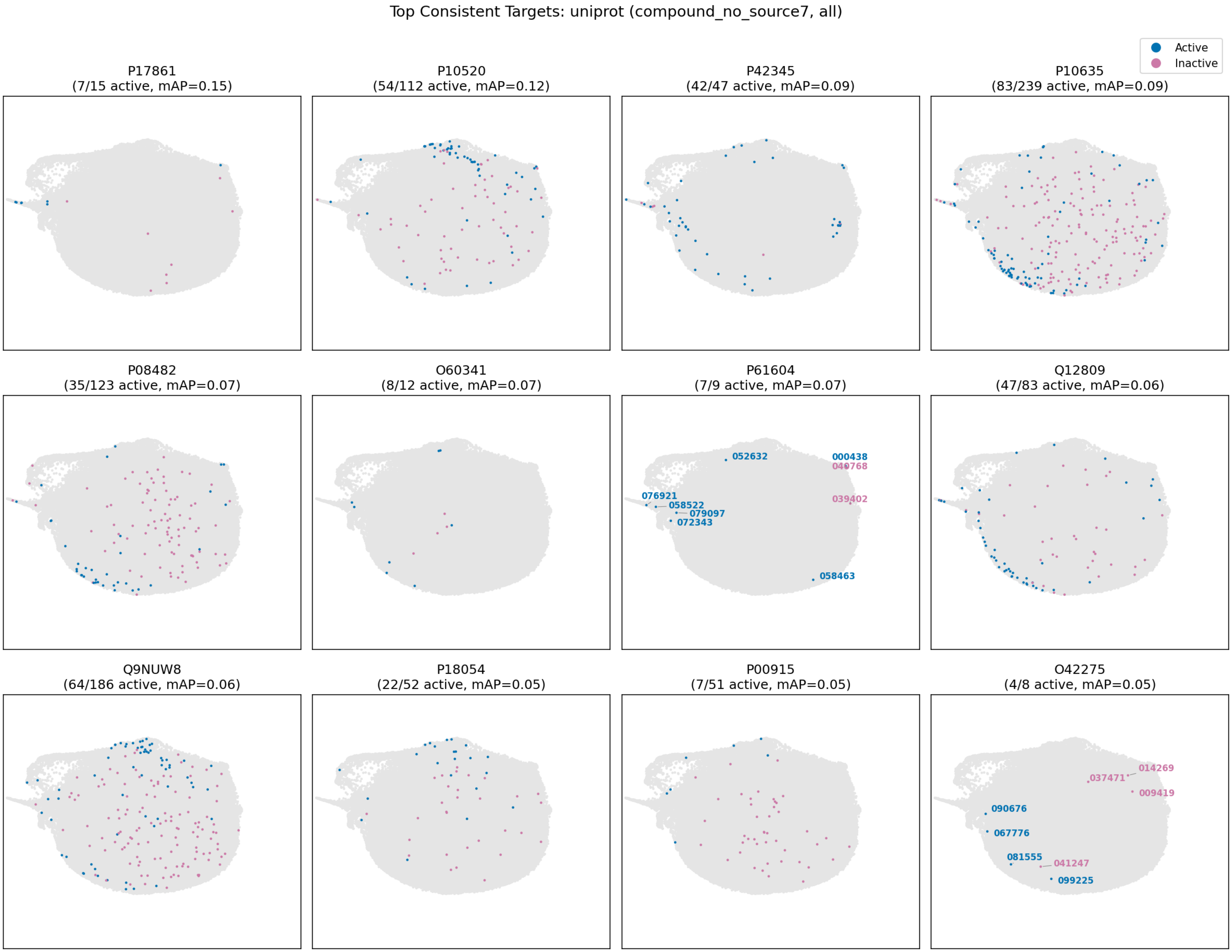

Top Consistent Targets: uniprot (compound_no_source7, all)
Active
Inactive
P17861
(7/15 active, mAP=0.15)
P10520
(54/112 active, mAP=0.12)
P42345
(42/47 active, mAP=0.09)
P10635
(83/239 active, mAP=0.09)
P08482
(35/123 active, mAP=0.07)
O60341
(8/12 active, mAP=0.07)
P61604
(7/9 active, mAP=0.07)
052632
000438
040768
039402
076921
058522
079097
072343
058463
Q12809
(47/83 active, mAP=0.06)
Q9NUW8
(64/186 active, mAP=0.06)
P18054
(22/52 active, mAP=0.05)
P00915
(7/51 active, mAP=0.05)
O42275
(4/8 active, mAP=0.05)
014269
037471
009419
090676
067776
081555
041247
099225

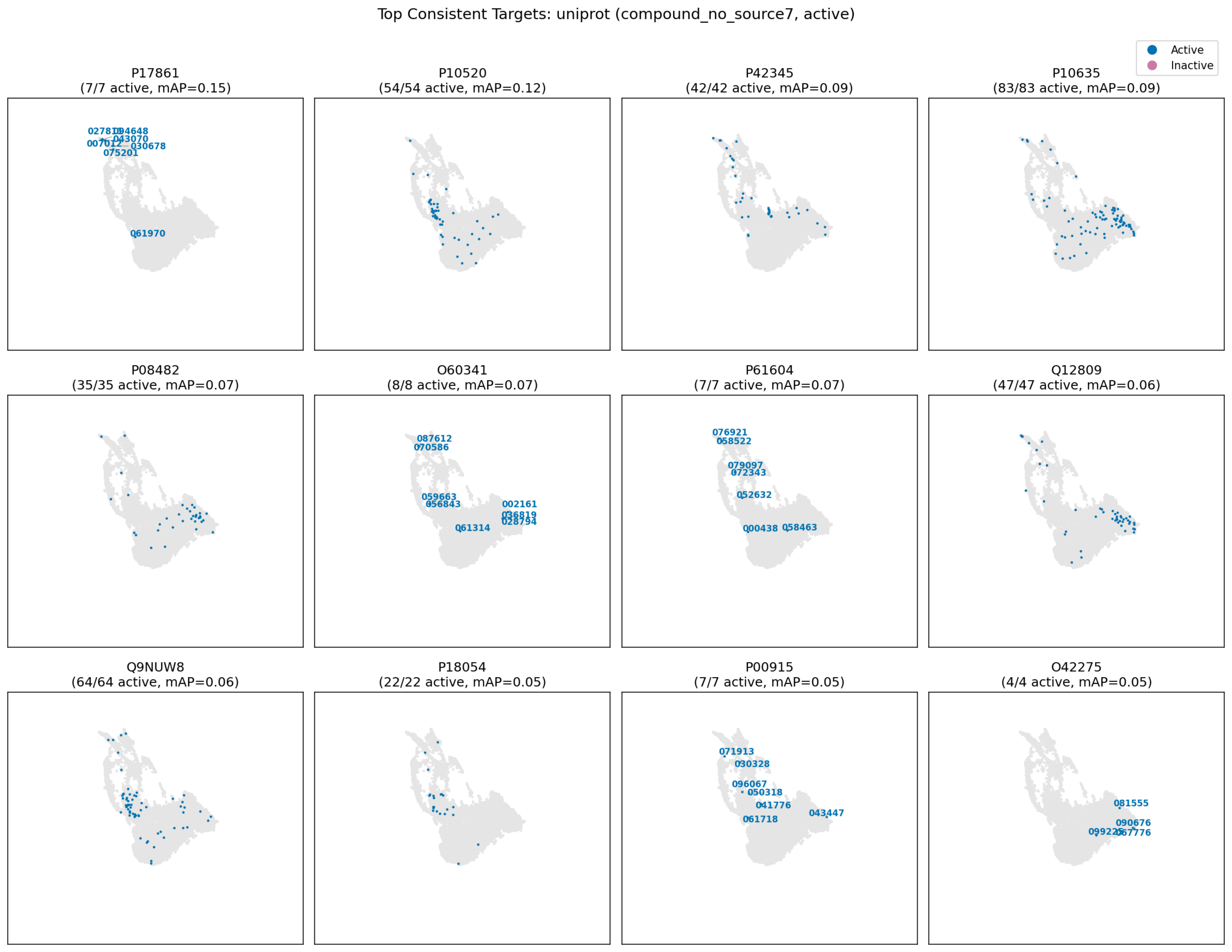
Top Consistent Targets: uniprot (compound_no_source7, active)
Active
Inactive
P17861
(7/7 active, mAP=0.15)
043070
007012
030678
075201
061970
P10520
(54/54 active, mAP=0.12)
P42345
(42/42 active, mAP=0.09)
P10635
(83/83 active, mAP=0.09)
P08482
(35/35 active, mAP=0.07)
O60341
(8/8 active, mAP=0.07)
087612
070586
059663
056843
002161
061314
P61604
(7/7 active, mAP=0.07)
076921
058522
079097
072343
052632
000438
058463
Q12809
(47/47 active, mAP=0.06)
Q9NUW8
(64/64 active, mAP=0.06)
P18054
(22/22 active, mAP=0.05)
P00915
(7/7 active, mAP=0.05)
071913
030328
096067
050318
041776
043447
061718
O42275
(4/4 active, mAP=0.05)
081555
090676
